# Recent Non-LTR Retrotransposon Activity Predicts Cancer Prevalence in Mammals

**DOI:** 10.1101/2025.08.30.673284

**Authors:** Vahid Nikoonejad Fard, Marc Tollis

## Abstract

Non-long terminal repeat retrotransposons (nLTRs), including long and short interspersed nuclear elements (L1 and SINEs), are the most abundant and active mobile elements in mammals. NLTRs play critical mutagenic and regulatory roles during oncogenesis in humans and model species. However, it is not known whether recent nLTR activity in the genome is related to the lifetime cancer risk of a species beyond humans and conventional model organisms. We examined whether recent nLTR activity predicts cancer prevalence across mammals using comparative analyses of *de novo* whole-genome repeat annotations from 55 species, each with over 20 published zoo pathology records. We quantified nLTR activity as the number of potentially active elements, their proximity to protein-coding genes and cancer gene orthologs (CGOs), and insertions within these genes. Across all three metrics, neoplasia prevalence was associated with both L1 and combined L1-SINE activity, while malignancy was linked exclusively to the L1-SINE predictors. This pattern suggests a complementary and escalating trajectory, where L1s contribute to early tumorigenic events, while SINE activity, driven by L1s, amplifies their impact and fuels the transition to malignancy. Moreover, genomes harboring more CGOs tended to exhibit higher neoplasia prevalence, and the number of fusion cancer genes was strongly correlated with the number of potentially active L1s across species. Our results further revealed a pattern wherein species with minimal cancer prevalence exhibit restricted activity of at least one major nLTR superfamily, suggesting that preserving genome stability through limited retrotransposition may serve as a protective mechanism against cancer.

## Introduction

Cancer has persisted throughout evolutionary history as an inherent tradeoff of multicellularity, with its presence now documented across nearly all metazoan lineages (Aktipis et al. 2015; Albuquerque et al. 2018). Malignant tumors occasionally arise from inherited mutations in critical oncogenes or tumor suppressor genes (TSGs), or more commonly, from the accumulation of somatic mutations, often accompanied by epigenetic reprogramming and widespread gene expression dysregulation during an individual’s lifetime. These aberrations collectively compromise core cellular programs, including growth suppression, proliferative restraint, apoptotic signaling, and other regulatory mechanisms that underlie the emergence of hallmark cancer traits (Hanahan and Weinberg 2011; Hanahan 2022).

Non-long terminal repeat retrotransposons (nLTRs), including autonomous long (L1) and nonautonomous short (SINEs) interspersed nuclear elements, are the most abundant and active transposable elements (TEs) in mammals. These elements have played a pivotal role in shaping the architecture, function, and evolutionary dynamics of mammalian genomes (Chalopin et al. 2015; Wells and Feschotte 2020; Colonna Romano and Fanti 2022). However, their activity also has well-documented mutagenic and regulatory consequences for the initiation and progression of various cancer types in humans and model organisms (Burns 2017; Lynch-Sutherland et al. 2020; Grundy et al. 2022). Aberrant nLTR activity can generate oncogenic structural and copy number variations (SVs and CNVs), ranging from small-scale insertions, deletions, and duplications to extensive chromosomal rearrangements, including chromoanagenesis events (Lee et al. 2012; Helman et al. 2014; Rodić et al. 2015; Rodriguez-Martin et al. 2020; Mendez-Dorantes et al. 2024). Not merely agents of mutation, nLTRs contribute to the TE repository of regulatory elements, providing alternative promoters, enhancers, and silencers, and serve as sources of noncoding RNA (ncRNAs) precursors, thereby reshaping transcriptional and post-transcriptional landscapes (Jacques et al. 2013; Johnson and Guigó 2014; Chuong et al. 2017; Trizzino et al. 2017; Lynch-Sutherland et al. 2020; Park et al. 2022; Lee et al. 2024). These multifaceted effects are often preceded by widespread TE derepression driven by extensive DNA hypomethylation and chromatin accessibility (Jansz 2019; Choudhary et al. 2023). Collectively, these alterations drive tumorigenesis by activating or amplifying oncogenes and inactivating or deleting TSGs (Lee et al. 2012; Tubio et al. 2014; Jang et al. 2019; Rodriguez-Martin et al. 2020).

In mammals, nLTRs are particularly active in the germline and during early embryonic development, where relaxed epigenetic constraints permit their transcription and mobilization (DiRusso & Clark, 2023; Fort et al., 2014; Oomen & Torres-Padilla, 2024; Zamudio & Bourc’his, 2010). Accordingly, novel oncogenic insertions can be passed on to offspring, contributing to the species’ inherited cancer risk. Such events are not rare, as any two human haploids, for example, differ by over 1,000 TE insertions, predominantly from nLTRs activity (Stewart et al. 2011; Sudmant et al. 2015). Consistently, *de novo* L1 insertions are estimated to occur in approximately 1% of human births and >12% of mouse births (Hancks and Kazazian 2012; Richardson et al. 2017). Manifesting this legacy, over 100 inherited human diseases have been linked to nLTR insertions that disrupt critical gene functions (Hancks and Kazazian 2016; Han et al. 2023).

At the somatic level, while nLTRs are often epigenetically repressed in healthy, terminally differentiated cells to safeguard genome stability, this control usually collapses in cancer cells. It is well recognized that L1 overexpression is a hallmark of many human cancers (Rodić et al. 2014). Approximately 1% of L1 insertions in certain human cancers are tumor-initiating events, and up to 50% of human tumors harbor somatic L1 retrotranspositions (Lee et al. 2012; Helman et al. 2014; Tubio et al. 2014; Cajuso et al. 2019; Rodriguez-Martin et al. 2020). Notably, L1’s ORF1 and ORF2 proteins, along with nLTR-derived transcripts, including those from SINEs, and nLTR-associated antigens, are recurrently expressed in malignant cells and are now increasingly recognized as diagnostic biomarkers and immunotherapeutic targets across a broad range of tumors (Kim et al. 2013; Laumont et al. 2018; Kong et al. 2019; Ponomaryova et al. 2020; Shah et al. 2023; Chaaban et al. 2025). SINEs can also be highly active upon derepression (Kramerov and Vassetzky 2011; Vassetzky and Kramerov 2013). For instance, Alu insertions within human *BRCA1* and *BRCA2* have been causally implicated in ovarian and breast cancer predisposition (Teugels et al. 2005; Talhouet et al. 2020).

The extensive mutagenic and regulatory potential also provides cancer cells with a rich source of variations that can be harnessed to develop novel oncogenic traits. Interestingly, cancer cells exploit this diversity, i.e., oncoexaptation, to evolve oncogenic traits such as sustained oncogenic transcription, immune evasion, cellular plasticity, resistance to cell death, and metastatic capacity (Jang et al. 2019; Payer and Burns 2019; Lynch-Sutherland et al. 2020; Grundy et al. 2022). In this broader context, nLTR activity can promote tumor evolution by fostering genome instability and chronic inflammation, two major enabling processes in somatic cancer evolution (Belgnaoui et al. 2006; Hancks and Kazazian 2016; Simon et al. 2019; Marasca et al. 2020; Gorbunova et al. 2021). Thus, genomes enriched in active nLTRs may be inherently more vulnerable to destabilization and oncogenesis, unless buffered by robust lineage-specific genome stabilizing and cancer defense mechanisms.

Despite the extensive body of research detailing nLTR contributions in cancer, their impact on cancer risk beyond humans and conventional model species remains unknown. Mammalian genomes exhibit broadly similar TE content and diversity, with L1 and SINEs representing the dominant elements, accounting for, on average, 22.6% (range: 8.2 to 52.8%) and 10.5% (range: 0.4 to 32.1%) of the genome, respectively. Despite this overarching similarity, recent nLTR activity and expansions differ substantially across the clade, as does cancer prevalence (Osmanski et al. 2023; Compton et al. 2024). Building on these observations and the wide-ranging and profound impacts of nLTRs in human cancers, we hypothesize that interspecific differences in recent nLTR activities may underlie the variation in cancer prevalence observed across mammals. To investigate this hypothesis, we developed a comparative analytical framework of four distinct yet interconnected models linking neoplasia and malignancy prevalence to (1) recent intact insertion counts, as proxies for active nLTRs, in an “Abundance model”; (2) the proximity of these elements to protein-coding (PC) genes and cancer gene orthologs (CGOs) in a “Proximity model”; (3) insertions within these genes in a “Genic-Insertion model”; and (4) CGO copy number and its functional subsets, including somatic, germline, oncogene, tumor suppressor, and fusion genes, in a “Cancer Gene Load model” (Table 1). Longevity was also included in analyses given its relevance to both nLTR activity and cancer risk (Cruickshanks et al. 2013; Gorbunova et al. 2021; Giordani et al. 2021; Merenciano et al. 2025). By systematically evaluating these models, we aim to clarify how nLTR-driven genomic dynamics may shape cancer risk across mammals, offering insights for human oncology and biodiversity conservation.

**Table 1.** Overview of statistical models used in the study.

| Model Name | Specifications<br>(Predictors) | Predictions<br>(Response Variables) |
| --- | --- | --- |
| Abundance Model | Number of either potentially active (i.e., full-length, recently inserted) L1s or combined L1+SINEs;<br>The multiple regression form includes longevity. | Neoplasia prevalence or<br>Malignancy prevalence |
| Proximity Model<br>(Average/Median) | Mean or median distance of potentially active L1s or combined L1+SINEs to the nearest protein-coding gene or cancer gene orthologs;<br>The multiple regression form includes longevity. | Neoplasia prevalence or<br>Malignancy prevalence |
| Genic-Insertion<br>Model | Number of insertions by potentially active L1s or combined L1+SINEs within protein-coding gene or cancer gene orthologs;<br>The multiple regression form includes longevity. | Neoplasia prevalence or<br>Malignancy prevalence |
| Cancer Gene Load<br>Model | Number of human cancer gene orthologs across various functional categories, including total number, somatic, germline, oncogene, tumor suppressor, and fusion genes per species;<br>The multiple regression form includes longevity | Neoplasia prevalence or<br>Malignancy prevalence |

## Materials and Methods

### Cancer Data

We retrieved cancer data from Compton et al. (2024). This dataset comprises 16,049 necropsy records across 292 species maintained under human care. Neoplasia and malignancy prevalence for each species in this dataset were calculated by dividing the number of necropsies reporting tumors (i.e., benign or malignant) by the total number of necropsies, and are presented as percentages (Supplementary Data S1). We restricted this dataset to species with at least 20 necropsy records to mitigate the impact of sampling error and statistical bias. We further excluded species with zero recorded neoplasia or malignancy, as cancer has been reported in almost all mammals (Aktipis et al. 2015; Rifkin et al. 2017; Albuquerque et al. 2018), and such zeros most likely reflect insufficient sampling rather than actual biological absence. This filtering approach removes likely undersampling artifacts and reduces bias in data distribution that otherwise violates regression assumptions.

### Genomic Data

We downloaded genome assemblies, annotations, and proteomes for 55 mammalian species spanning eleven orders, including Artiodactyla, Carnivora, Chiroptera, Cingulata, Diprotodontia, Hyracoidea, Lagomorpha, Pilosa, Primates, Proboscidea, and Rodentia from NCBI Datasets (Sayers et al. 2024). We restricted our analyses to assemblies with contig N50 values exceeding 10 kb, i.e., more than 1.5 times the 6.1 kb length of the L1 consensus sequence (Babushok and Kazazian 2007; Beck et al. 2011). We set this threshold to balance between assembly quality and species inclusion. While stricter thresholds could improve TE annotation accuracy, they would disproportionately exclude species with less-refined genomes, thereby reducing taxonomic representation and limiting the comparative and statistical power of analyses. We prioritized contig N50 as the primary quality metric for assemblies, as it better captures assembly continuity relevant to TE detection compared to BUSCO completeness scores. BUSCO scores were additionally considered to confirm the assembly completeness, with results indicating that more than 80% of the assemblies used in these analyses had BUSCO completeness scores exceeding 90% (i.e., 45 assemblies), while over 90% of assemblies had scores above 80% (i.e., 50 assemblies) (See Supplementary Data S1).

Annotation files were available for 31 of the mammalian species on NCBI. For the remaining species lacking annotations or proteomes, we transferred gene models from closely related, well-annotated species with LiftOff v1.6.3 (Shumate and Salzberg 2021) (Supplementary Data S1). Proteomes were either downloaded directly from NCBI or generated from annotation files using the convert function in GFFtK v23.11.2 (Palmer, 2024). To ensure adequate proteome completeness for downstream analyses, we excluded species with peptide BUSCO scores (Simão et al. 2015) below 60% from the Proximity, Genic-Insertion, and Cancer Gene Load models, as PC genes and CGOs were identified based on these annotations and proteome datasets.

### Longevity Data

Longevity data were obtained from the Myhrvold Amniote Database (Myhrvold et al. 2015) (Supplementary Data S1). Although the African elephant (*Loxodonta africana*, 960 months) and the chimpanzee (*Pan troglodytes*, 888 months) were identified as statistical outliers in the dataset by Grubbs’ test, we retained them as biologically meaningful data points. These species are naturally long-lived, and their longevity values reflect well-documented biological realities rather than errors or anomalies. Notably, their inclusion did not alter the significance of the models or violate any regression assumptions. Thus, they were retained as influential but valid observations in all analyses (see Supplementary Scripts).

### Repeat Annotation and nLTR Data Extraction

We employed RepeatModeler 2.0 (Flynn et al. 2020) to perform *de novo* identification of repetitive elements and to create species-specific repeat libraries. Next, we used RepeatMasker 4.1 (Smit et al. 2013) to query each species library against the respective reference genome to generate repeat annotations. The RepeatMasker output files were then converted to BED files using rmsk2bed from BEDOPS 2.4.41 (Neph et al. 2012). We used various in-house Python and Shell scripts to parse these BED files and: (1) quantify potentially active L1 and SINE elements for the Abundance model; (2) measure the average and median distances from each element to the nearest PC gene and CGO using BEDTools’ closest function (Quinlan and Hall 2010) for the Proximity model; and (3) identify and count recent insertions within PC genes and potential CGOs using BEDTools’ intersect function for the Genic-Insertion model (see Data Availability for code repositories).

Elements were classified as potentially active if L1s measured within ±10% of the 6.1 kb consensus length (Babushok and Kazazian 2007; Beck et al. 2011) and SINEs between 100 and 400 bp (Kramerov and Vassetzky 2011; Vassetzky and Kramerov 2013), both with ≤5% sequence divergence, indicating recent intact insertions. To estimate the proximity of elements to functional regions, we calculated the median and the mean within the interquartile range (IQR) of the distances from L1 and SINE elements to their nearest PC genes and CGOs. IQR means were used because the average distance in each species is heavily influenced by extreme values, which can distort typical proximity measures. These IQR means were computed by excluding outliers outside the bounds defined as Q1 − 1.5 × IQR and Q3 + 1.5 × IQR, where IQR is the difference between the third (Q3) and first (Q1) quartiles.

### Human Cancer Gene Orthologs In Mammals

To identify potential CGOs in each species, we first downloaded a list of 743 human cancer genes from the Cancer Gene Census within the COSMIC database (Sondka et al. 2018; Tate et al. 2019) (last accessed April 2024). We then retrieved their protein sequences from Ensembl v112 using BioMart (Kinsella et al. 2011), resulting in protein sequences for 721 cancer genes. For each cancer gene, we retrieved 1) gene symbols, 2) gene names, and 3) whether cancer-related mutations were germline, somatic, or both, and their role in cancer (i.e., oncogene, TSG, or both), as well as whether or not they were fusion genes. Next, we used this peptide file along with the complete proteomes of 46 mammals, with peptide BUSCO scores of over 60% in OrthoFinder (Emms and Kelly 2019). Then, we parsed the OrthoFinder output files (i.e., Orthogroups.GeneCount.tsv and Orthogroups.tsv) via in-house Shell, Python, and R scripts to count the number of orthologs for each cancer gene in species, as well as classify them into somatic, germline, oncogene, TSGs, and fusion genes (see Data Availability for code repositories). Notably, mouse (*Mus Musculus*, n=3298) and rat (*Rattus norvegicus*, n=3128) were identified as outliers in the total CGO counts, and mouse was also an outlier in the oncogene counts (i.e., 1607) according to Grubbs’ tests results. However, we retained them as valid influential observations because previous studies have confirmed their exceptionally high number of cancer genes (Tollis et al. 2020), and, importantly, because their inclusion neither altered the significance of our results nor violated regression assumptions (see Supplementary Scripts).

### Statistical Analyses

We conducted regression analyses using a modified Phylogenetic Generalized Least Squares (PGLS) function developed by Compton et al. (2024) to account for the non-independence of cancer prevalence among related species. This function, pglsSEyPagel, extends the phytools pglsSEy method by incorporating an estimated Pagel’s *λ* (Pagel 1999) into the regression model, rather than assuming a fixed value of one (i.e., Brownian motion). All analyses were performed in R v4.4.2 using a suite of packages available in the supplementary analytical scripts. A phylogenetic tree comprising 55 species was retrieved from TimeTree (https://timetree.org, Kumar et al., 2022; last accessed October 2024). Estimates of neoplasia and malignancy prevalence are inherently more reliable in species with larger necropsy sample sizes. To account for variation in sampling effort, species necropsy data were weighted by the inverse square root of each species’ total necropsy count (i.e., 1/√n).

We report the phylogenetic signal (*λ*) alongside the *p*-value and R² for all analyses. In all three nLTR modeling frameworks, we tested two distinct predictors of retrotransposon activity: (1) autonomous L1 elements alone, and (2) the combined burden of autonomous L1 and nonautonomous SINE elements. Because SINEs lack the enzymatic machinery required for mobilization and depend entirely on proteins encoded by autonomous elements, modeling SINEs alone as an independent predictor is biologically unjustified. However, we included SINE counts in the combined L1-SINE measurements for species in which no full-length L1s were detected. These SINE counts represent genuine insertions with divergence rates below 5%, indicating recent mobilization events potentially facilitated by unannotated or fragmented but transcriptionally active L1 remnants, or, to a lesser extent, by other TEs such as retrotransposable elements (e.g., RTE-BovB in Bovids) (Ohshima and Okada 2005; Almojil et al. 2021). This approach allowed us to more comprehensively capture recent retrotranspositional activity that may contribute to genome instability and cancer risk across species.

All response and predictor variables were moderately to highly right-skewed, particularly the nLTR data due to multiple zeros. This extreme skewness results in masking and swamping effects, whereby influential points mask each other, non-outliers are incorrectly flagged as outliers, or violations of model assumptions. Using Tukey’s Ladder of Power, we power-transformed all response and predictor variables to address these issues. Additionally, all predictor data were min– max normalized to a [0, 1] range to facilitate consistent effect size interpretation across models and to ensure visual comparability of results. For all PGLS analyses, we assessed model assumptions prior to inference, including normality of residuals, homoscedasticity, independence of errors, and collinearity in multiple regressions. To control for multiple testing, we applied a 5% False Discovery Rate (FDR) correction using the Benjamini-Hochberg method (Benjamini and Hochberg 1995).

### Comparing Statistical Models

We also conducted an Akaike Information Criterion test to see whether PGLS models outperformed the standard linear regression, as the phylogenetic signals in our results were negligible in some cases. The AIC test is a metric used to evaluate and compare statistical models based on their goodness of fit and complexity, and equals AIC = −2*ln* (Likelihood) + 2*k*, where *k* is the number of the model’s estimated parameters (i.e., complexity). The model with the lowest AIC is considered the best-fitting model. Results for comparing models are presented in Supplementary Tables S9A and S9B.

## Results

We examined whether the recent activity of two genomically pervasive nLTR superfamilies is associated with the prevalence of neoplasia and malignancy across mammals. To account for the non-independence of data among related species, we employed PGLS models using various nLTR activity measures as predictors. NLTR activity was evaluated based either on autonomous L1 elements alone or the combined burden of L1 and SINE elements, as modeling SINEs alone is biologically unjustified due to their reliance on L1-encoded enzymes for mobilization.

### Baseline Patterns of Cancer Prevalence and nLTRs Burden

Our dataset comprised 55 species from all four major eutherian superorders encompassing Laurasiatheria (n = 33), Euarchontoglires (n = 16), Afrotheria (n = 2), and Xenarthra (n = 2), as well as two marsupial species (Supplementary Data S1, Fig. 1). This representation of species across clades reflects the limited number of taxa for which both sufficient necropsy records and genome assemblies with adequate contiguity and annotation were available. Across the species, the highest neoplasia prevalence was observed in the ferret (63%), Asian elephant (40%), and mouse (39%), while the lowest rates appeared in the naked mole rat (1.3%), tammar wallaby (2.3%), and straw-colored fruit bat (3.2%). Malignancy prevalence was highest in the ferret (40%), snow leopard (30%), and mouse (27%), and lowest in the naked mole rat (0.7%), armadillo (2.5%), and Jamaican fruit bat (3.7%). Active L1 counts were highest in the rat (11,856), common marmoset (7,530), and mouse (7,419). In contrast, nine species, including raccoon, meerkat, and the straw-colored fruit bat, showed no detectable full-length L1 elements inferred to be transpositionally active. Active SINE counts were highest in the ruffed lemur (91,383), Hamadryas baboon (58,956), and colobus monkey (57,498), and lowest in the Baringo giraffe (1), Addra gazelle (3), and Guinea hog (3) (Supplementary Data S1, Fig. 1).

**Figure 1.**
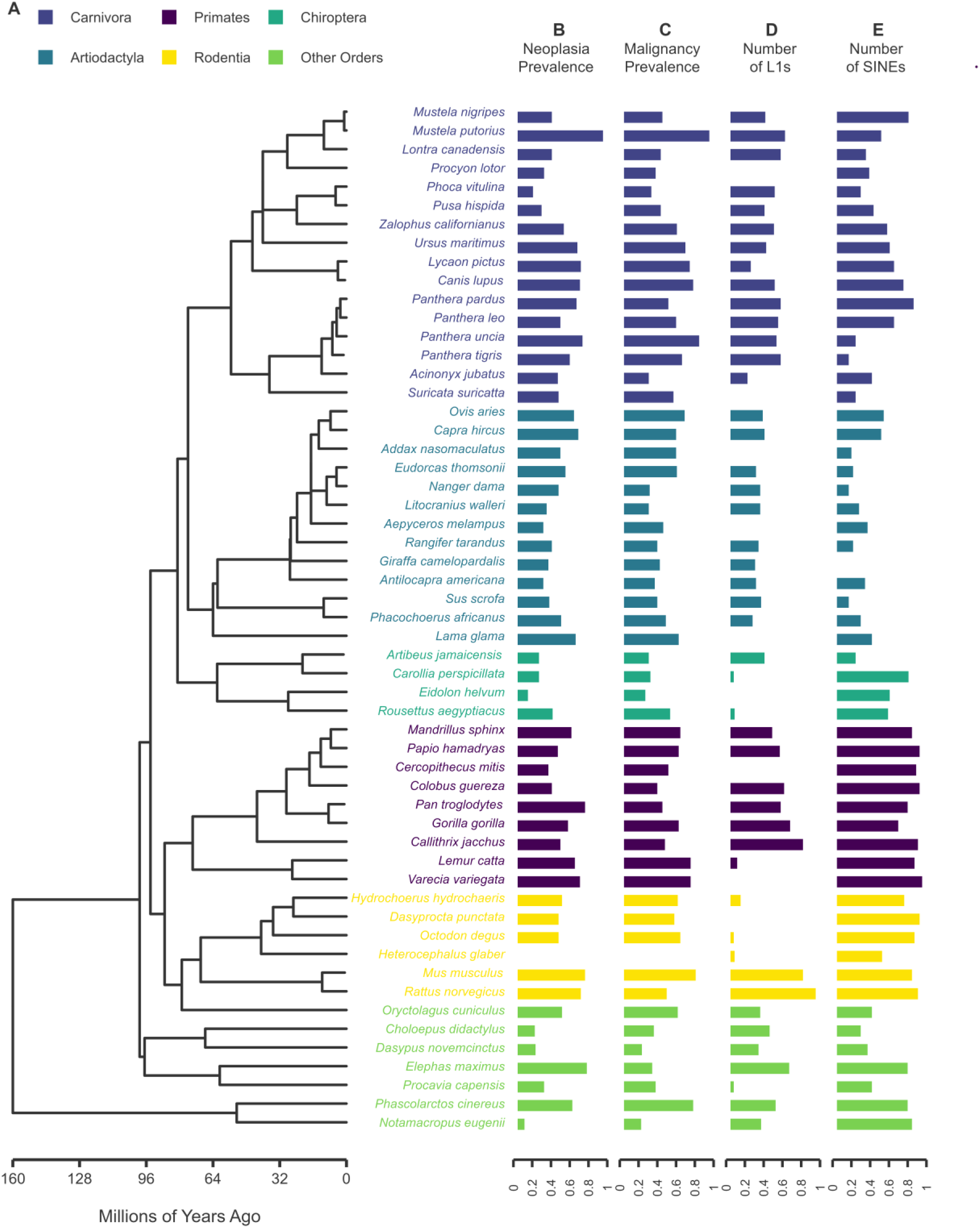
Ladderized phylogenetic tree of the 55 mammalian species analyzed in this study. The main tree was derived from TimeTree (Kumar et al., 2022; https://timetree.org). Species are color-coded by taxonomic order. Horizontal bar plots to the right of the tree display neoplasia prevalence, malignancy prevalence, the number of active L1s, and the number of active SINEs identified in each genome. All values are Tukey-transformed and normalized to the [0, 1] range to enhance cross-species comparability.

### The Abundance Model

We first tested whether the number of potentially active nLTRs predicts cancer prevalence across mammals. We found a significant association between neoplasia prevalence and both the number of active L1s (*p* = 0.003, R² = 0.16) and the combined L1-SINE burden (*p* = 0.016, R² = 0.09) across 55 species (Supplementary Table S1A; Fig. 2A-B). Malignancy prevalence was also significantly linked with the combined L1-SINE burden (*p* = 0.045, R² = 0.07; Fig. 2C), whereas the number of L1s alone was not predictive (*p* = 0.17, R² = 0.04; Supplementary Table S1A; Supplementary Fig. S1). When longevity was included as a covariate, both nLTR predictors remained significantly associated with neoplasia (L1: FDR-adjusted *p* = 0.006, adjusted R² = 0.14; L1 and SINEs: FDR-adjusted *p* = 0.02, adjusted R² = 0.08; Supplementary Table S1B). However, neither predictor showed a significant correlation with malignancy, and longevity did not emerge as a significant factor in any multiple PGLS model.

**Figure 2.**
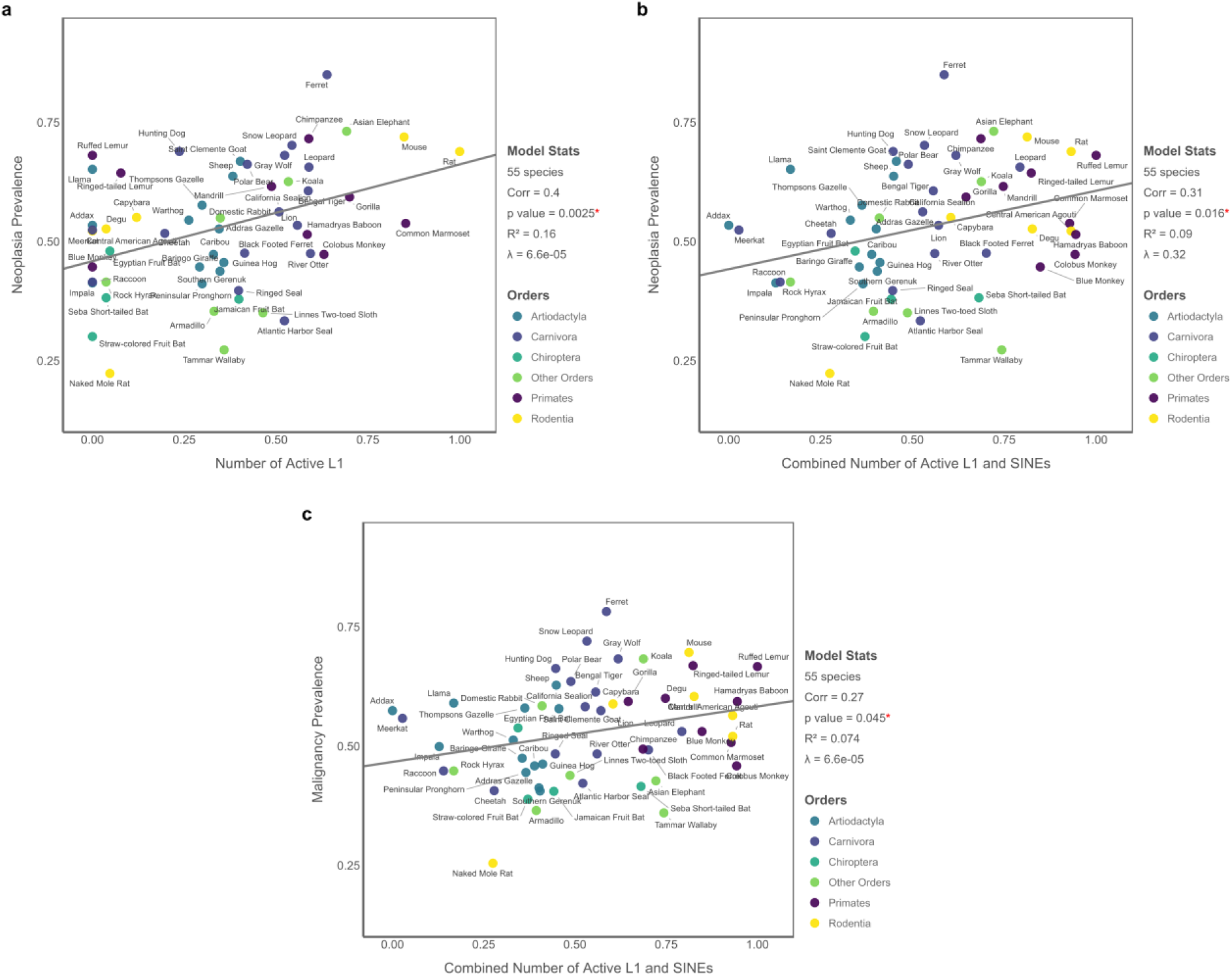
PGLS results for the Abundance Model showing significant associations between active nLTR abundance and cancer prevalence across 55 mammalian species. (A) Neoplasia prevalence is positively associated with the number of active L1 elements. (B) The combined number of active L1s and SINEs also predicts neoplasia prevalence. (C) Malignancy prevalence is significantly associated with the combined number of active L1s and SINEs. All variables are Tukey-transformed; predictors are additionally normalized to the [0, 1] range for consistent effect size interpretation and visualization.

### The Proximity Model

Next, we evaluated whether the genomic proximity of active nLTR elements to PC genes and CGOs is associated with variation in mammal cancer prevalence. Among 45 species examined, neoplasia prevalence was positively associated with the average distance between active L1 elements and PC genes, a counterintuitive pattern relative to initial expectations. This association was significant in both simple (*p* = 0.018, R² = 0.12; Fig. 3A; Supplementary Table S2A) and multiple (FDR-adjusted *p* = 0.032, adjusted R² = 0.08; Supplementary Table S2B) PGLS models. In contrast, the average proximity of combined L1s and SINEs to PC genes showed no association with neoplasia prevalence (*p* = 0.50, R² = 0.01), and neither proximity measure was significantly related to malignancy prevalence (L1: *p* = 0.16, R² = 0.05; L1 and SINEs: *p* = 0.62, R² = 0.006; Supplementary Table S2A; Supplementary Fig. S2). Adding longevity as a covariate yielded the same results, and longevity itself did not emerge as a significant contributor (Supplementary Table S2B). Analyses using median distances produced qualitatively similar results and are presented in Supplementary Tables S3A-S3B and Supplementary Figs. S3 and S4.

**Figure 3.**
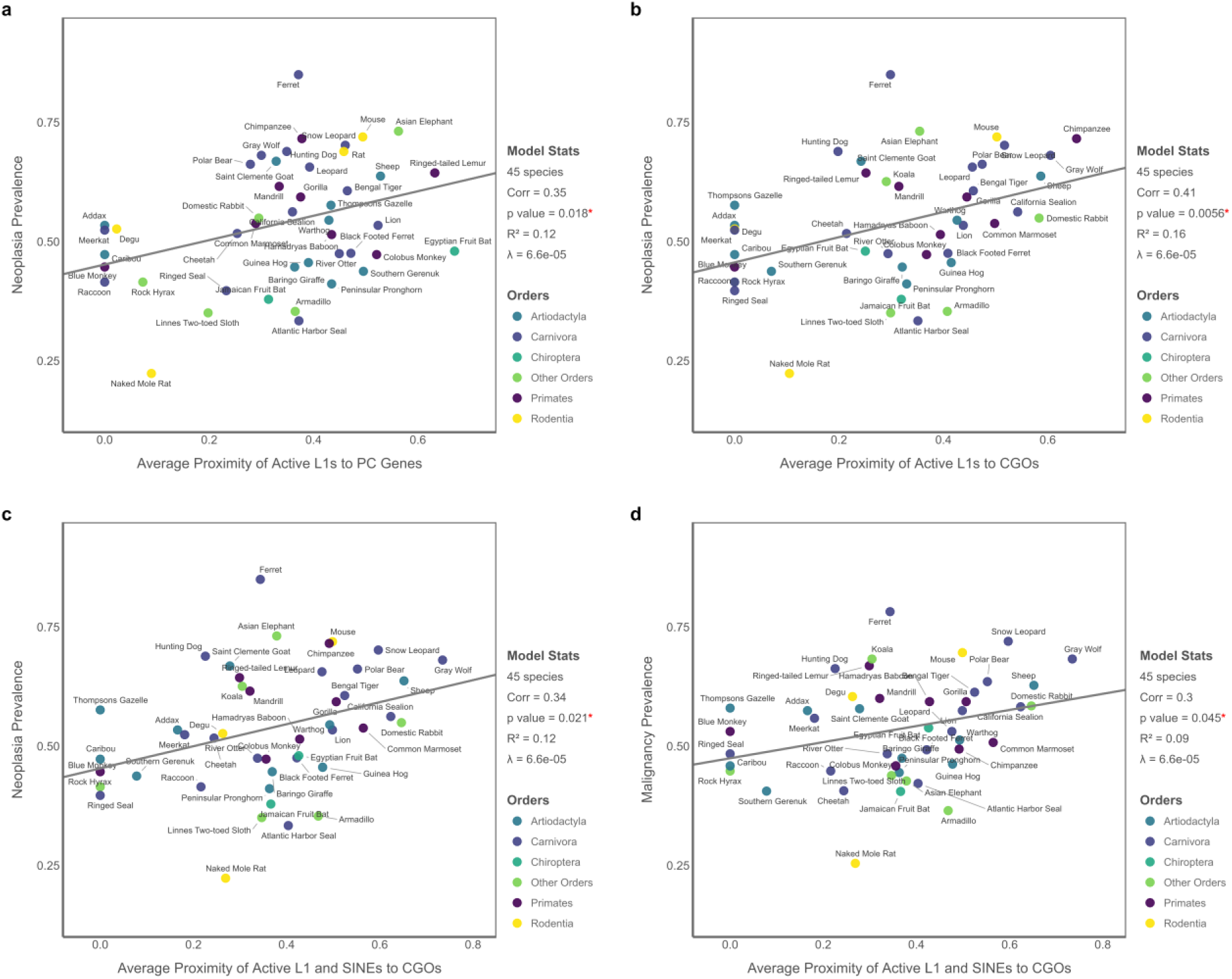
PGLS results for the Proximity Model in 45 mammals based on the average distance of active L1s and combined L1 and SINEs to PC genes and CGOs. (A) Neoplasia prevalence is significantly associated with the average proximity of active L1 elements to PC genes. (B) Neoplasia prevalence is also associated with the average proximity of L1 elements to CGOs. (C) A similar association is observed using the combined proximity of L1s and SINEs to CGOs. (D) Malignancy prevalence is significantly associated with the average proximity of L1s and SINEs to CGOs. All variables are transformed using Tukey’s ladder of powers, while predictors are additionally scaled to the [0, 1] range to allow consistent interpretation and visualization of effect sizes across models.

We also observed significant links between the proximity of active nLTRs to CGOs and cancer prevalence. Neoplasia prevalence was positively associated with both the average proximity of L1s (*p* = 0.006, R² = 0.16; Fig. 3B) and the combined L1s and SINEs (*p* = 0.02, R² = 0.12; Fig. 3C; Supplementary Table S4A). These links remained significant when controlling for longevity (FDR-adjusted *p* = 0.01 and *p* = 0.04, respectively; Supplementary Table S4B). Additionally, malignancy prevalence was significantly correlated with the combined proximity of L1s and SINEs to CGOs (*p* = 0.045, FDR-adjusted p = 0.049, R² = 0.09; Fig. 3D), while the L1-only model was not predictive (*p* = 0.07, R² = 0.09; Supplementary Fig. S2D). Longevity did not significantly affect any of these models, and median-based proximity models yielded similar but slightly weaker patterns, with significant associations limited to neoplasia prevalence only (Supplementary Tables S5A-S5B; Supplementary Fig. S4).

### The Genic-Insertion Model

To assess whether the insertion of active elements within functional regions is linked to cancer susceptibility, we examined the number of active L1 and combined L1-SINE insertions intersecting PC genes and CGOs. In models based on 46 species, we found that neoplasia prevalence was significantly associated with both the number of active L1 (*p* = 0.003, R² = 0.18) and the combined number of L1 and SINE insertions within PC genes (*p* = 0.01, R² = 0.13; Supplementary Table S6A; Fig. 4A-B). These associations remained statistically significant after adjusting for species longevity (FDR-adjusted *p* = 0.006, adjusted R² = 0.14, and FDR-adjusted *p* = 0.03, adjusted R² = 0.09, respectively; Supplementary Table S6B). In contrast, malignancy prevalence was not associated with the number of L1 insertions within PC genes alone (*p* = 0.15, R² = 0.05), but showed a marginal significant positive relationship with the combined L1-SINE insertion count (*p* = 0.049, R² = 0.08; Fig. 4C), which remained significant after controlling for Longevity (FDR-adjusted *p* = 0.044, adjusted R² = 0.08; Supplementary Table S6B). Longevity itself was not a significant predictor in any of the Genic-Insertion models.

**Figure 4.**
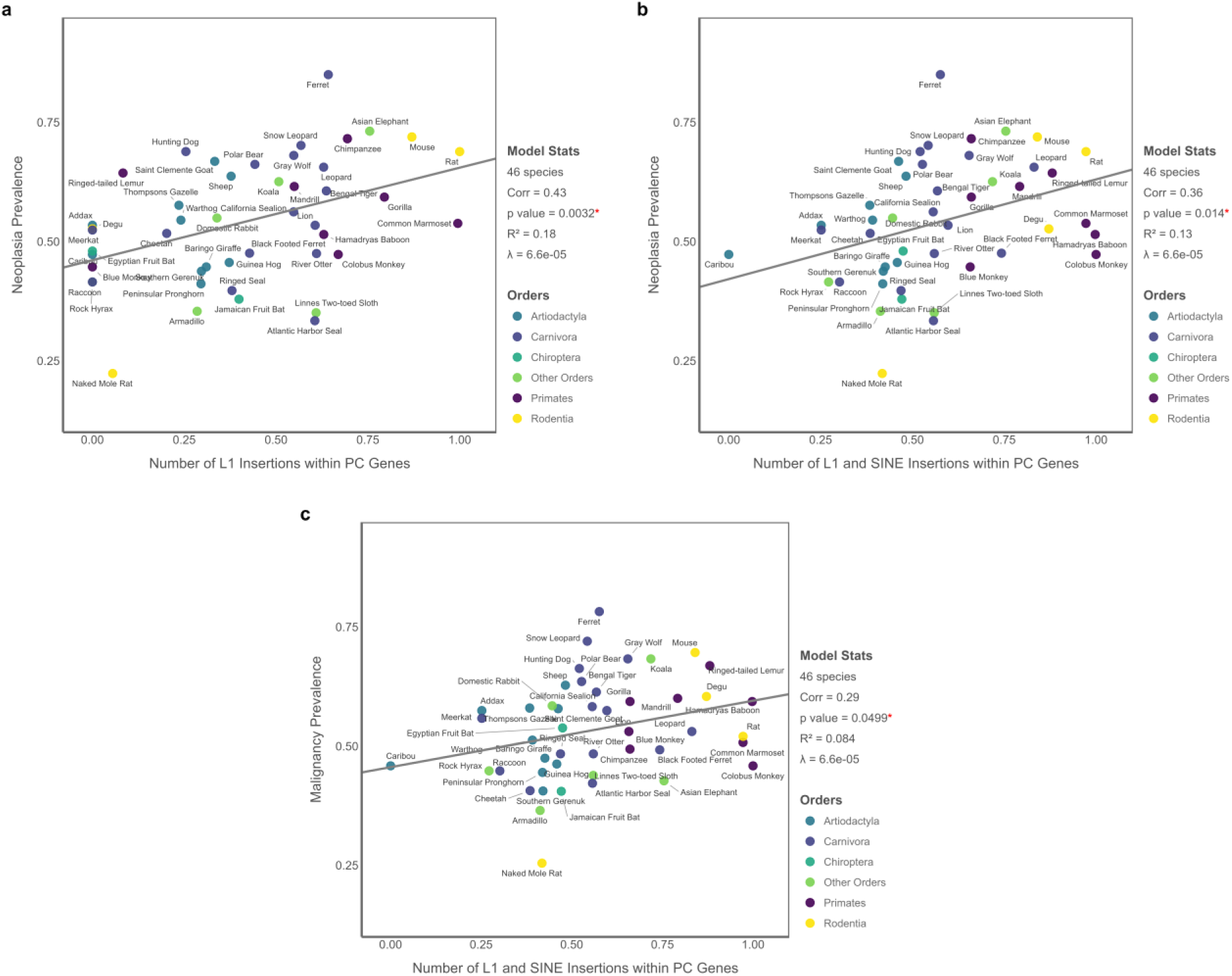
PGLS results for the Genic-Insertion Model based on the number of active nLTR insertions within PC genes (n = 46 species). (A) Neoplasia prevalence is significantly associated with the number of L1 insertions. (B) The combined number of L1 and SINE insertions also predicts neoplasia prevalence. (C) Malignancy prevalence is significantly associated with the combined number of L1 and SINE insertions within PC genes. All variables are transformed using Tukey’s ladder of powers. Predictor variables were further scaled to the [0, 1] range to ensure consistent interpretation and visualization of effect sizes across models.

**Figure 5.**
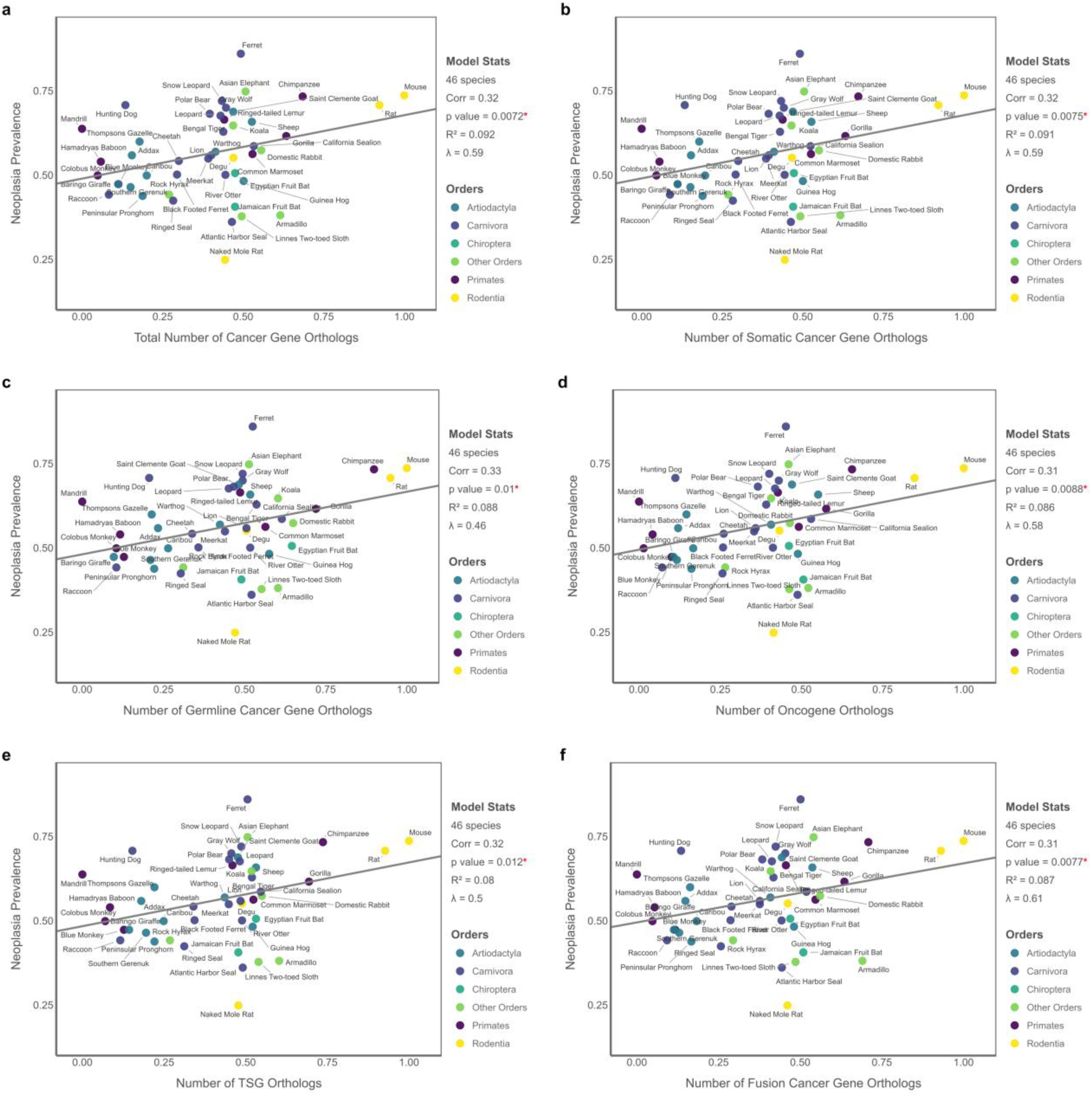
PGLS results for the Potential Cancer Gene Load Model showing associations between neoplasia prevalence and the number of cancer gene homologs (n = 46 species). Each panel displays results for a distinct subset of CGOs: (A) total number of CGOs, (B) somatic CGOs, (C) germline CGOs, (D) oncogene orthologs, (E) TSG orthologs, and (F) fusion gene orthologs. All variables are transformed using Tukey’s ladder of powers. Predictor variables are further scaled to the [0, 1] range to ensure consistent interpretation and visualization of effect sizes across models.

When insertions into CGOs were considered, no significant associations were observed with either neoplasia or malignancy prevalence (Supplementary Tables S7A-S7B). This lack of signal may be partly attributable to the high number of species with few detected insertions in these genes, which resulted in poor model fits. To assess whether the genic insertion results are orthogonal to overall TE abundance, we further evaluated the correlation between the number of L1 insertions within PC genes and total active L1 counts across species. This correlation was extremely high (*r* = 0.98). Similarly, the number of combined L1 and SINE insertions within PC genes was highly correlated with their overall genomic abundance (*r* = 0.92). When both total TE counts and their genic insertion counts were included as predictors in multiple PGLS models, neither variable remained statistically significant, and variance inflation factors exceeded the threshold (i.e., >5) in all models, indicating substantial collinearity (Supplementary Table S14).

### The Cancer Gene Load Model

Beyond retrotransposon metrics, we also tested whether interspecies variation in the number of detected CGOs in different categories, including total number of CGOs, somatic, germline, oncogene, TSG, and fusion genes, is associated with cancer prevalence. Following the approach described by Matthews et al. (2025), we employed Orthofinder to detect CGOs in species based on their proteome files, which excluded retrogenes and pseudogenes and thus focused solely on PC gene family expansions and contractions. Therefore, CGO counts reported here may differ from those in previous studies (Caulin and Maley 2011; Sulak et al. 2016; Tollis et al. 2020; Vazquez and Lynch 2021). Among the 46 species examined, mouse, rat, and chimpanzee had the highest total counts of CGOs of all types, whereas mandrill, colobus monkey, and Hamadryas baboon had the lowest counts. These species also showed the same pattern for TSG and oncogene ortholog counts.

Using simple PGLS models, we found that neoplasia prevalence is significantly associated with the total number of CGOs (*p* = 0.007, R² = 0.09), as well as with each functional category, comprising somatic (*p* = 0.007, R² = 0.09), germline (*p* = 0.01, R² = 0.09), oncogene (*p* = 0.009, R² = 0.09), TSGs (*p* = 0.01, R² = 0.08), and fusion genes (*p* = 0.008, R² = 0.09) (Supplementary Table S8A). In contrast, malignancy prevalence showed no significant correlation with any of the CGO categories. These associations were robust to the inclusion of longevity as a covariate, with FDR-adjusted *p*-values ranging from 0.013 to 0.044 and adjusted R² values ranging from 0.05 to 0.074 (Supplementary Table S8B). We also identified an almost perfect association between the number of oncogenes and TSGs across species (*r* = 0.98), which aligns with a prior report by Tollis et al. (2020).

### Interrelationships Among Predictors

Cancer is an age-dependent disease, meaning species with longer lifespans accumulate more mutations over time, which in turn, increases cancer risk as they age (Seluanov et al. 2018). However, across the 55 mammals examined, we found no significant link between longevity as a response and neoplasia (*p* = 0.58, R² = 0.41) or malignancy prevalence (*p* = 0.78, R² = 0.4) as predictors (Supplementary Table S13), although a weak negative trend was observed for malignancy (Fig. S10). We further examined whether nLTR abundance and insertion are associated with two additional biological features across species, including longevity and the number of annotated fusion genes. We identified a strong positive association between the number of active L1 elements and the number of fusion genes (*p* = 0.00017, R² = 0.38; Supplementary Table S12, Fig. 6), congruent with the recognized capacity of L1s to promote genomic rearrangements that give rise to fusion gene formation (Moran et al. 1999; Rodriguez-Martin et al. 2020; Mendez-Dorantes et al. 2024). The combined L1-SINEs burden also showed a weaker positive trend, though not statistically significant (*p* = 0.07, R² = 0.23), reinforcing the idea that autonomous retroelements play a pivotal role in genome rearrangement leading to gene fusions, likely due to mechanisms such as mediating recombination or 3′ transductions (Xing et al. 2006; Tubio et al. 2014). Furthermore, we surveyed the relationship between nLTR activity and species longevity. Neither the total number of active L1s (*p* = 0.2, R² = 0.42) nor the combined L1-SINE burden (*p* = 0.53, R² = 0.41) could predict lifespan across the 55 mammalian species analyzed (Supplementary Table S10). Similarly, the number of nLTR genic-insertions within coding regions was not predictive of longevity (*p* = 0.06, R² = 0.4 for L1s, *p* = 0.92, R² = 0.36 for L1 and SINEs; Supplementary Table S11).

**Figure 6.**
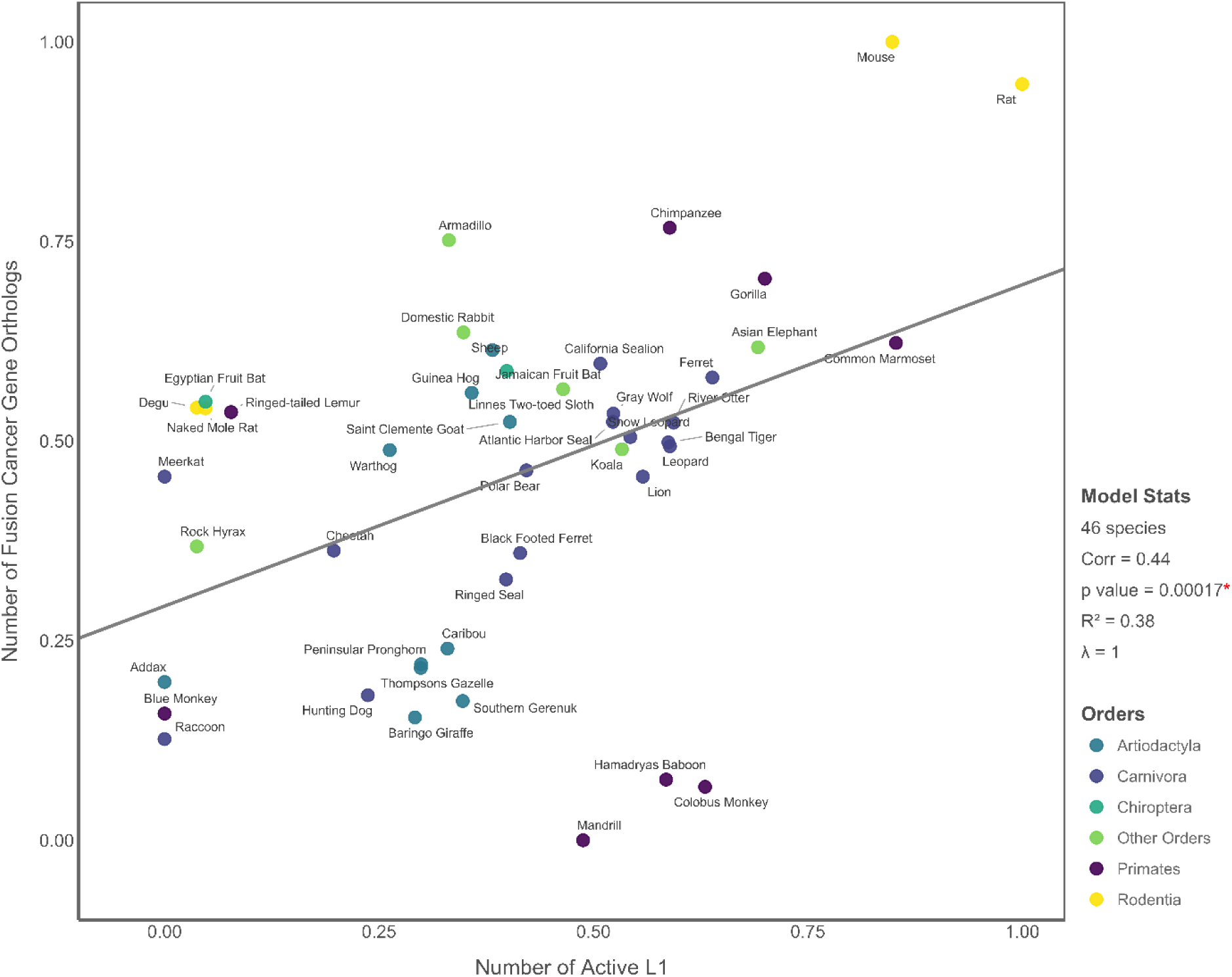
PGLS result showing a strongly significant positive association between the number of active L1 elements and the number of fusion cancer gene orthologs across 46 mammalian species. All variables are transformed using Tukey’s ladder of powers. The predictor and response variables are also scaled to the [0, 1] range to ensure consistent interpretation and visualization of effect sizes across models.

### Phylogenetic Signal and Model Goodness-of-Fit

Because the underlying phylogenetic relatedness between species can affect comparative analyses, we tested whether incorporating shared ancestry has improved model performance by comparing Akaike Information Criterion (AIC) scores across models with different levels of phylogenetic signal. For most of our simple and multiple PGLS models, the AIC values for *λ* = 0 and the estimated *λ* were identical or nearly so, reflecting the negligible *λ* estimates in many models. Models using *λ* = 1 consistently yielded substantially higher AIC scores than *λ* = 0 or estimated-*λ* models, indicating poorer fits (Supplementary Table S9A and S9B).

## Discussion

Whether nLTRs act as active drivers or passive passengers during oncogenesis remains unclear. However, decades of research have firmly established their diverse mutagenic and regulatory contributions to the initiation and progression of various cancers in human and model species (Burns 2017; Lynch-Sutherland et al. 2020; Grundy et al. 2022), where the imprint of nLTR activity is a hallmark of the most common and deadly human malignancies, including lung, colorectal, breast, and prostate cancers (Lee et al. 2012; Rodić and Burns 2013; Cajuso et al. 2019; Grundy et al. 2022). Here, leveraging a comparative oncology framework spanning up to 55 species, we found significant associations between recent nLTR activity and neoplasia and malignancy prevalence. Our findings extend the current knowledge beyond humans and conventional model organisms to a broad range of mammals, suggesting that genomes enriched in active nLTRs tend to be inherently more susceptible to germline and somatic oncogenic alterations, as widespread derepression of fully competent or structurally eroded yet transcriptionally active elements compromises genome integrity and creates a permissive environment for neoplastic transformation.

Across all nLTR models, we found that neoplasia prevalence is associated with both L1 abundance and combined L1 and SINE burden, indicating that autonomous retrotransposition alone can be sufficient to trigger early tumor development. In contrast, malignancy prevalence was linked only to the combined load of L1s and SINEs, pointing to a compounding effect that amplifies genome instability beyond the impact of L1 elements alone. This pattern suggests that L1 elements may play a predominant role in tumor initiation, while SINE activation, spurred by elevated L1 activity, may further accelerate cancer progression and malignant transformation. Together, these findings reveal a macroevolutionary pattern in which the interplay between autonomous and nonautonomous nLTR activity has shaped cancer predisposition during mammalian evolution, highlighting retroelements as a pervasive and long-term force in genome-cancer dynamics.

Nevertheless, the relationship between nLTR abundance and cancer prevalence is not universally linear. Some cancer-resistant species invest heavily in genome surveillance and stability through mechanisms like enhanced apoptosis, stringent cell cycle regulation, efficient DNA repair, and robust immune surveillance (Tollis et al. 2017b, a; Boddy et al. 2020a, b). These mechanisms enable rapid detection and neutralization of oncogenic damage, including that associated with nLTR activity. Elephants represent a prime example of such an evolutionary cancer resistance mechanism; despite their massive body size, long lifespan, and relatively high neoplasia rate, the malignant conversion of neoplastic lesions remains strikingly rare, a pattern attributed to their multiple copies of *TP53* that provide robust tumor suppressive functions (Abegglen et al. 2015; Tollis et al. 2017a, 2021; Vazquez and Lynch 2021). Interestingly, this well-established pattern is manifested in our results. The Asian elephant genome contains 4,143 potentially active L1s and 6,783 SINEs, reflecting a substantial nLTR burden. However, despite ranking second in highest neoplasia prevalence (i.e., 0.4), only 11% of these cases advance to malignancy (i.e., 0.045 malignancy prevalence). This result reinforces the idea that strong lineage-specific cancer defenses can counterbalance even high retrotranspositional pressure and modulate the TE-cancer dynamics beyond simple abundance-driven expectations.

We also found that the proximity of recent nLTR insertions to coding regions is correlated with cancer prevalence across mammals, but in an unexpected direction, with greater distances correlating positively with both neoplasia and malignancy prevalence. The biological significance of the insertions used in our analyses is nuanced, as these elements have been co-opted or tolerated during the recent evolutionary history of species, yet their presence, even when far from genes, does not imply they are harmless. Distal insertions can remain mobilization-competent or be transcomplemented to seed new insertions elsewhere. They may also influence gene regulation through long-range enhancer or promoter activity, supply ORF2p enzymes, producing aberrant transcripts and ncRNAs with oncogenic potential (Helman et al. 2014; Johnson and Guigó 2014; Chuong et al. 2017; Lynch-Sutherland et al. 2020; Park et al. 2022; Lee et al. 2024). Furthermore, their sequence homology can promote non-allelic and ectopic recombination events, particularly when nested or clustered, induce genome-wide hypomethylation, remodel chromatin architecture that can bring them into contact with cancer genes, and ultimately contribute to genome destabilization (Kent et al. 2017; Fueyo et al. 2022; Choudhary et al. 2023; Mendez-Dorantes et al. 2024).

Most importantly, selection may exert different pressures on germline-heritable versus somatic insertions arising in tumors. In somatic cancer evolution, gene-proximal insertions that would be disadvantageous in the germline may confer a selective advantage to cancerous cells by enhancing their oncogenic fitness. Contrastingly, at the population and species level, strong purifying selection typically purges germline insertions near gene-rich regions that disrupt essential functions. TE studies also consistently confirm that insertions are disproportionately skewed toward intergenic and gene-poor regions, reflecting the long-term retention of elements in genomic “safe havens” subject to weaker selective constraints (Charlesworth et al. 1994b, a; Zamudio and Bourc’his 2010; Levin and Moran 2011; Marsano and Dimitri 2022). Thus, deleterious cancer-associated insertions that persist in present-day genomes are generally expected to be farther from coding regions, likely resulting in a gradient of deleterious potential that increases with distance from critical genes when comparing species genomes. While usually repressed, these elements can be reactivated when epigenetic regulation collapses in cancer. Consequently, species with intact nLTRs concentrated farther from genes may retain a larger reservoir of potentially oncogenic elements that have escaped strong selection, which could help explain their higher cancer prevalence. Nonetheless, further molecular and population-level studies will be required to determine whether this pattern reflects a genuine biological signal or results from structural artifacts in the data.

Results from the Genic-Insertion model for PC genes mirrored those from the Abundance model, with the combined intragenic L1-SINE load predicting both neoplasia and malignancy, while L1 insertions were associated only with neoplasia. This pattern did not extend to CGOs, likely due to poor model fit driven by multiple species with few detectable nLTR insertions in these genes. Just as with intergenic insertions, the persistence of intragenic elements is also likely due to co-option or evolutionary tolerance, whereby insertions confer antagonistic pleiotropic benefits or their harmful effects manifest post-reproductively (Medawar 1957; Williams 1957; Campisi 2013; Boddy et al. 2020b; Merenciano et al. 2025). Remarkably, germline mutations account for up to 20% of the known mutations in human cancer genes (Sondka et al. 2018), contributing to cancer predisposition. Some well-established nLTR-derived examples include *de novo* Alu insertions into *BRCA1* and *BRCA2*, which predispose carriers to hereditary breast and ovarian cancers (Batzer and Deininger 2002; Teugels et al. 2005), and germline L1 insertions into the *APC* TSG, which underlie familial adenomatous polyposis and increase colorectal cancer risk (Miki et al. 1992).

Notably, a recent comparative study found that species-level germline mutation rates predict cancer mortality across 37 vertebrates (Kapsetaki et al. 2023). Our results for both Abundance and Genic-Insertion models consistently suggest that nLTR-derived insertions may represent an additional source of heritable mutational burden contributing to the broader mutation-cancer relationship reported in that study. However, we also found strong correlations between genic-insertion counts and overall nLTR abundance (*r* = 0.97 for L1s; *r* = 0.92 for L1-SINEs). Such multicollinearity complicates causal inference, making it unclear whether intragenic insertions independently elevate cancer risk or reflect the impact of broader genomic nLTR load. Active nLTRs exhibit minimal insertional preference compared to other TE types, resulting in a broadly dispersed insertional pattern across mammalian genomes (Richardson et al. 2015). This pattern likely explains why species with more recent nLTR insertions tend to have more intragenic insertions. Future studies, including population-level polymorphism surveys and insertional site preference assays, can disentangle the specific contribution of intragenic insertions from the broader effects of nLTR abundance.

Results from the cancer gene load models indicate that the total number of CGOs and each functional category, including somatic, germline, oncogenes, TSGs, and fusion genes, is significantly associated with neoplasia across species, yet not with malignancy. This pattern implies that a larger CGO repertoire increases the insertional target for disruption by nLTR activity and other mutagenic processes, raising the probability of oncogenesis. We also observed an almost perfect positive correlation between counts of TSGs and oncogenes across species (*r* = 0.98), consistent with a prior report by Tollis et al. (2020). This close correspondence suggests that the evolutionary expansion of TSGs and oncogenes has been tightly coordinated, likely driven by selective pressures to balance proliferative capacity with genomic safeguards. Taken together, these patterns are consistent with a larger mutational target for cancer initiation coupled with strengthened defenses that constrain malignant progression, which may explain the lack of correlation with malignancy.

Furthermore, we identified a highly significant association between the number of active L1 elements and fusion gene counts. Fusion genes can be generated directly by nLTR activities through insertion, deletion, and 3′ transduction, or indirectly by providing cryptic splice sites or promoters that drive read-through transcripts encompassing two adjacent genes. Moreover, sequence homology between active or even degenerated elements can facilitate non-allelic homologous and ectopic recombination, leading to the formation of fusion genes (Tubio et al. 2014; Goodier 2016; Makałowski et al. 2019; Gasparotto et al. 2023; Prokopov et al. 2025). Further experimental surveys, such as RNA-seq and retrotransposition assays, can confirm these findings and determine the extent to which active L1 elements contribute to fusion gene formation across species.

Evaluation of cancer prevalence relative to longevity revealed no predictive association for either neoplasia or malignancy. Likewise, longevity did not emerge as a significant covariate in any of our nLTR or CGO models, suggesting that lifespan may evolve independently from the cancer risk across mammals. These findings align with a recent report showing no association between longevity and cancer prevalence, total PC gene copy numbers, or the expansion and contraction of individual PC gene families across 94 mammalian species examined (Matthews et al. 2025). However, they contrast with the results of several previous studies (Sulak et al., 2016; Tollis et al., 2020; Vazquez & Lynch, 2021). For instance, Tollis et al. (2020) reported a significant link between cancer gene duplications and longevity quotient across 63 mammals. Similarly, several studies of elephant lineage have reported correlations between *TP53* copy numbers, body size, and Longevity (Caulin and Maley 2011; Caulin et al. 2015; Abegglen et al. 2015; Vazquez and Lynch 2021).

Likewise, none of the nLTR metrics could predict longevity in models using longevity as the response variable. Numerous studies have shown that TE activity increases with age due to loss of epigenetic control, and this activation in turn exacerbates aging and reduces lifespan by inducing DNA damage and chronic inflammation, which contribute to age-related diseases such as cancer (Belgnaoui et al. 2006; De Cecco et al. 2013a, 2019; Simon et al. 2019; Gorbunova et al. 2021). However, it has also been shown that effective suppression of TE activity extends the lifespan of select model animals (De Cecco et al. 2013b; Wood et al. 2016; Giordani et al. 2021; Yushkova and Moskalev 2023). This effective TE suppression, together with species-specific cancer defense mechanisms in some species, may therefore decouple longevity from both cancer prevalence and nLTR metrics in our dataset. Overall, the relationships between longevity, cancer prevalence, and other related factors such as TEs activity at the macroevolutionary level remain controversial and inconsistently resolved, as lifespan is a multifactorial, context-dependent trait shaped by species-specific adaptations, further complicated by methodological heterogeneity and data limitations inherent in comparative studies.

Among all the species examined, the naked mole rat exhibited the lowest prevalence of neoplasia and malignancy, reinforcing its status as an exceptional model for cancer resistance studies. Naked mole rat cells produce very high molecular weight hyaluronan (vHMW-HA), which triggers hypersensitive contact inhibition through CD44-mediated upregulation of *p16^Ink4a* gene. This process limits cell proliferation and protects cells against DNA damage (Tian et al. 2013; Takasugi et al. 2020). A previous study has also reported that this species carries more TSG copies than any other rodents (Tollis et al., 2020). However, it did not rank among the top species in our dataset, likely because our analyses were restricted to PC gene orthologs only. Notably, we detected only two active L1 elements in the naked mole rat’s genome, which may indicate an additional layer of defense that protects genome integrity by diminishing the parasitic activity or mitigating genomic instability caused by nLTRs, thereby contributing to the species’ exceptionally low cancer incidence.

Bats also exhibit remarkably low cancer rates despite their small body sizes and high metabolic demands (Wang et al. 2011; Tollis et al. 2017b; Gonzalez et al. 2018), a pattern also reflected in cancer data. Strikingly, Seba’s short-tailed bat, straw-colored fruit bat, and Egyptian fruit bat showed extremely low numbers of potentially active L1s (i.e., 1, 0, and 2, respectively), suggesting reduced retrotransposon activity as a possible contributor to genome stability in these species. Interestingly, while these bats had varying nontrivial SINE counts, the Jamaican fruit bat presented the opposite pattern, with a relatively high number of potentially active L1s (i.e., 859) but remarkably few active SINEs (i.e., 6). Although the impact and consequences of these contrasting autonomous and nonautonomous nLTR landscapes are different, they may indicate species-specific strategies to prevent genomic instability caused by retroelements.

Previous studies have shown that bats exhibit positive selection on numerous TSG and DNA repair genes, alongside unique immune adaptations such as the modifications in the Interferon I gene family (Zhang et al. 2013; Foley et al. 2018; Scheben et al. 2023), which together likely underpin their ability to restrict genome instability and enhanced cancer resistance. Similarly, the genome of the nine-banded armadillo included only 23 active SINE counts despite 506 full-length L1 elements, aligning with its low malignancy prevalence observed in this species (Vazquez et al., 2022). Taken together, the pattern of reduced activity of at least one major nLTR superfamily across these species supports the idea that species-specific adaptations to respond to DNA damage and maintain genome integrity may be an essential form of cancer prevention.

Nevertheless, the results of this study should be interpreted with caution. Primarily, necropsy-based cancer prevalence data were derived from animals under human care, which may not fully represent disease burdens in the wild. Besides, environmental exposures vary widely among species, and extended lifespans in captivity can reveal cancer susceptibilities that might remain unexpressed in natural conditions. Although we adjusted for variation in necropsy sample sizes, uneven sampling may also still influence the results, and statistical power may have been limited by the relatively small number of species meeting inclusion criteria.

Limitations in genomic data are another consideration. Despite applying quality filters, fragmented assemblies can underestimate nLTR counts, particularly for L1s, and inflate distance estimates. Our classification of potentially active elements, based on length thresholds and ≤5% divergence, is not a perfect criterion, as it may exclude degraded elements that remain transcriptionally active. Unlike studies considering pseudogenes and retrogenes, we restricted CGO counts to PC genes. These counts can be affected by annotation and proteome quality, ascertainment bias from human peptide queries, and evolutionary divergence that results in undetected orthologs. Cancer genes annotated in the human genome may not serve identical roles in other species, and species-specific cancer genes not implicated in human cancers may exist. Finally, while comparative correlations yield valuable evolutionary insights, they do not establish causality, and further studies are necessary to confirm these patterns. Even so, the corroborative results across Abundance, Proximity, and Genic-Insertion models provide a coherent signal supporting our inferences.

## Conclusion and Future Directions

This study provides the first macroevolutionary analysis linking recent nLTR activity to cancer prevalence across diverse mammals. Our results suggest reframing nLTRs not merely as stochastic mutagenic and regulatory agents contributing to somatic cancer evolution but also as a species-level evolutionary force that shapes long-term cancer vulnerability and the evolution of genome-stabilizing mechanisms. Consistent signals across all nLTR models indicate a predominant role for L1s in tumor-initiating events, while their convergent activity with SINEs intensifies their impact and boosts the malignant transformation. We also found that a larger CGO repertoire coincides with higher neoplasia prevalence, yet the coordinated expansion of TSGs and oncogenes, along with additional cancer defense mechanisms, particularly in cancer-resistant species, appears to alleviate the risk of malignancy. Interestingly, genomic L1 load exhibited a strong association with fusion gene burden, yet experimental surveys are needed to confirm these results and determine the extent to which L1s contribute to fusion gene formation. Furthermore, cancer-resistant species such as the naked mole rat and several bats showed reduced activity in at least one major nLTR superfamily, highlighting that cancer resistance in these species may partly depend on suppressing nLTR mobilization as potent agents of genome instability.

Advances in long-read sequencing now enable precise annotation of full-length TE insertions, especially within complex or heterochromatic regions, and the development of comprehensive TE catalogs with improved annotations. Combined with integrative multi-omics approaches, these platforms can reveal the precise transcriptional activity, regulatory roles, and epigenetic modifications of TEs in cancer-related pathways, particularly during the early stages of tumorigenesis. TE polymorphism surveys can further provide clearer insight into inherited variation in retrotransposition activity and the associated mutational burden contributing to cancer susceptibility by extending these insights to the population scale.

Future research in comparative oncology should also increasingly rely on the growing wealth of genomic resources and analytical frameworks to complement the traditional focus on ecological and life history traits. Expanding the scope of this research to include other structural and regulatory variants may yield further insights into the evolutionary dynamics of cancer predisposition and resistance. These efforts may target immunogenomic surveys, such as examining variation in immune gene repertoires, to examine the coevolution of immunity and cancer risk, and comparative epigenomic analyses, encompassing cancer-resistant lineages, to reveal the extent and complexity of lineage-specific TE silencing mechanisms contributing to genome stability, such as DNA methylation patterns. Expanding the taxonomic scope to comprise a wider range of mammals and other vertebrates would offer a broader evolutionary perspective. By expanding cancer research to include non-model species, comparative oncology promises to overcome limitations of traditional rodent models, deepen our understanding of genomic determinants of cancer risk, and develop a universal theory of cancer. These insights into the evolutionary basis of cancer prevalence and species-specific mechanisms for cancer suppression will ultimately inform human cancer research and support biodiversity conservation efforts.

## Supporting information

Supplementary Scripts

Supplementary Variable Descriptions

README file

## Acknowledgments

We gratefully acknowledge Dr. Simone Gable for her research contributions, which have enriched our understanding of transposable elements and genome evolution and informed aspects of this study. Computational analyses were performed using the Monsoon high-performance computing cluster at Northern Arizona University. This work was supported in part by the Division of Environmental Biology (National Science Foundation) under grant DEB 2323124 to M.T., a State of Arizona Technology Research Initiative Fund (TRIF) Faculty Support Grant awarded to M.T., and the National Institutes of Health U54 CA217376 (M.T.).

## Authors’ Contribution Statement

**V. N. Fard:** Conceptualization, Data Curation, Methodology, Software, Formal Analysis, Investigation, Visualization, Writing – Original Draft, Writing – Review & Editing.

**M. Tollis:** Conceptualization, Methodology, Supervision, Project Administration, Validation, and Writing – Review & Editing.

## Conflict of Interests

The authors declare no competing interests.

## Data Archiving

All scripts and datasets generated, analyzed, and used in this study are publicly available on GitHub (https://github.com/VNF1981/Non-LTR-Retrotransposons-Cancer-Project). A README.txt file provides detailed instructions on the data structure, analysis steps, and how to reproduce the results.

## Research Ethics Statement

This study did not involve live animals, human participants, or new sample collection. All analyses were conducted on data obtained from publicly available databases and published sources.

## Notes

https://github.com/VNF1981/Non-LTR-Retrotransposons-Cancer-Project/tree/main

### Competing Interest Statement

The authors have declared no competing interest.

