## Supplementary Tables and Figures for "Recent Non-LTR Retrotransposon Activity Predicts Cancer Prevalence in Mammals"

**Supplementary Table S1.** PGLS results for the Abundance Model. S1A reports associations between neoplasia or malignancy prevalence and the number of potentially active L1 elements and the combined number of L1s and SINEs in univariate PGLSs. S1B shows the same models with longevity included as a covariate in multivariate PGLSs. All variables are Tukey-transformed, with predictor variables also min-max normalized to the [0, 1] range to enable consistent effect size interpretation and visualization across models.

**Supplementary Table S1A**

| Response Variable | Predictor | Number of Species | Corr | Slope | Standard Error (SE) | *t-value* | *p-value* | $R^2$ | $\lambda$ |
|---|---|---|---|---|---|---|---|---|---|
| Neoplasia | Number of L1 | 55 | 0.4 | 0.008 | 0.002 | 3.17 | 0.003* | 0.16 | 0.00006 |
| | Number of L1 and SINEs | 55 | 0.31 | 0.2 | 0.08 | 2.49 | 0.016* | 0.09 | 0.32 |
| Malignancy | Number of L1 | 55 | 0.19 | 0.08 | 0.05 | 1.38 | 0.17 | 0.04 | 0.00006 |
| | Number of L1 and SINEs | 55 | 0.27 | 0.12 | 0.06 | 2.05 | 0.045* | 0.07 | 0.00006 |

**Supplementary Table S1B**

| Response Variable | Predictor | Number of Species | Corr | Slope | Standard Error (SE) | *t-value* | Adj. *p-value* (FDR) | Adj. $R^2$ | $\lambda$ |
|---|---|---|---|---|---|---|---|---|---|
| Neoplasia | Number of L1 | 55 | 0.4 | 0.2 | 0.07 | 3.01 | 0.006* | 0.14 | 0.00006 |
| | Longevity | | 0.17 | 0.00008 | 0.00009 | 0.89 | 0.37 | | |

| | | | | | | | | | |
|---|---|---|---|---|---|---|---|---|---|
| | Number of L1 and SINEs | 55 | 0.31 | 0.2 | 0.08 | 2.52 | 0.02* | 0.08 | 0.39 |
| | Longevity | | 0.17 | 0.0001 | 0.0001 | 1.27 | 0.21 | | |
| Malignancy | Number of L1 | 55 | 0.19 | 0.08 | 0.06 | 1.42 | 0.24 | 0.0005 | 0.00006 |
| | Longevity | | -0.03 | 0.00003 | 0.00008 | -0.38 | 0.7 | | |
| | Number of L1 and SINEs | 55 | 0.27 | 0.12 | 0.06 | 2.06 | 0.07 | 0.04 | 0.00006 |
| | Longevity | | -0.03 | -0.0003 | 0.00008 | -0.37 | 0.7 | | |

**Supplementary Table S2.** PGLS results for the Proximity Model based on the average distances of nLTRs to the PC Genes. S2A reports associations between neoplasia or malignancy prevalence and the average distance of potentially active L1 elements or the combined L1s and SINEs to the nearest PC Genes. S2B includes longevity as a covariate in multivariate models. All variables are Tukey-transformed, with predictor variables further min-max normalized to the [0, 1] scale for consistent effect size comparison and visualization across models.

**Supplementary Table S2A**

| Response Variable | Predictor | Number of Species | Corr | Slope | Standard Error (SE) | *t-value* | *p-value* | $R^2$ | $\lambda$ |
|---|---|---|---|---|---|---|---|---|---|
| Neoplasia | Proximity of L1 (Ave) | 45 | 0.35 | 0.26 | 0.1 | 2.45 | 0.018* | 0.12 | 0.00006 |
| | Proximity of L1 and SINEs (Ave) | 45 | 0.1 | 0.05 | 0.07 | 0.67 | 0.5 | 0.01 | 0.00006 |

| | | | | | | | | | |
|---|---|---|---|---|---|---|---|---|---|
| Malignancy | Proximity of L1 (Ave) | 45 | 0.22 | 0.13 | 0.09 | 1.45 | 0.16 | 0.05 | 0.00006 |
| | Proximity of L1 and SINEs (Ave) | 45 | 0.08 | 0.03 | 0.06 | 0.5 | 0.62 | 0.006 | 0.00006 |

**Supplementary Table S2B**

| Response Variable | Predictor | Number of Species | Corr | Slope | Standard Error (SE) | *t-value* | Adj. *p-value* (FDR) | Adj. $R^2$ | $\lambda$ |
|---|---|---|---|---|---|---|---|---|---|
| Neoplasia | Proximity of L1 (Ave) | 45 | 0.35 | 0.25 | 0.1 | 8.78 | 0.032* | 0.08 | 0.00006 |
| | Longevity | | 0.08 | 0.00003 | 0.0001 | 0.27 | 0.8 | | |
| | Proximity of L1 and SINEs (Ave) | 45 | 0.1 | 0.05 | 0.07 | 0.67 | 0.62 | -0.03 | 0.00006 |
| | Longevity | | 0.08 | 0.00005 | 0.0001 | 0.5 | 0.62 | | |
| Malignancy | Proximity of L1 (Ave) | 45 | 0.22 | 0.14 | 0.09 | 1.54 | 0.19 | 0.02 | 0.00006 |
| | Longevity | | -0.12 | -0.00008 | 0.00008 | -0.98 | 0.33 | | |
| | Proximity of L1 and SINEs (Ave) | 45 | 0.08 | 0.03 | 0.06 | 0.49 | 0.62 | -0.03 | 0.00006 |
| | Longevity | | -0.12 | -0.00007 | 0.00008 | -0.79 | 0.62 | | |

**Supplementary Table S3.** PGLS results for Proximity Model based on the median distance of nLTRs to PC Genes. S3A presents models relating neoplasia or malignancy prevalence to the median proximity of either L1 elements or L1s and SINEs combined. S3B includes longevity as an additional covariate. All variables are transformed using the Tukey method. Predictors are also min-max normalized to the [0, 1] scale to support standardized effect size comparisons and visualization across analyses.

**Supplementary Table S3A**

| Response Variable | Predictor | Number of Species | Corr | Slope | Standard Error (SE) | *t-value* | *p-value* | $R^2$ | $\lambda$ |
|---|---|---|---|---|---|---|---|---|---|
| Neoplasia | Proximity of L1 (Med) | 44 | 0.35 | 0.28 | 0.11 | 2.46 | 0.018* | 0.13 | 0.00006 |
| | Proximity of L1 and SINEs (Med) | 44 | 0.08 | 0.04 | 0.07 | 0.5 | 0.6 | 0.006 | 0.00006 |
| Malignancy | Proximity of L1 (Med) | 44 | 0.19 | 0.12 | 0.1 | 1.27 | 0.21 | 0.04 | 0.00006 |
| | Proximity of L1 and SINEs (Med) | 44 | 0.09 | 0.03 | 0.06 | 0.58 | 0.56 | 0.008 | 0.00006 |

**Supplementary Table S3B**

| Response Variable | Predictor | Number of Species | Corr | Slope | Standard Error (SE) | *t-value* | Adj. *p-value* (FDR) | Adj. $R^2$ | $\lambda$ |
|---|---|---|---|---|---|---|---|---|---|
| Neoplasia | Proximity of L1 (Med) | 44 | 0.35 | 0.28 | 0.0001 | 0.1 | 0.03* | 0.08 | 0.00006 |
| | Longevity | | 0.07 | 0.00001 | 0.0001 | 0.1 | 0.92 | | |

| | | | | | | | | | |
|---|---|---|---|---|---|---|---|---|---|
| | Proximity of L1 and SINEs (Med) | 44 | 0.08 | 0.04 | 0.07 | 0.5 | 0.65 | -0.04 | 0.00006 |
| | Longevity | | 0.07 | 0.00005 | 0.0001 | 0.45 | 0.65 | | |
| Malignancy | Proximity of L1 (Med) | 44 | 0.19 | 0.14 | 0.1 | 1.42 | 0.24 | 0.015 | 0.00006 |
| | Longevity | | -0.12 | -0.00009 | 0.00008 | -1.03 | 0.31 | | |
| | Proximity of L1 and SINEs (Med) | 44 | 0.09 | 0.035 | 0.06 | 0.6 | 0.55 | -0.02 | 0.00006 |
| | Longevity | | -0.12 | -0.00007 | 0.00008 | -0.8 | 0.55 | | |

**Supplementary Table S4.** PGLS results for the Proximity Model based on the average distance of active nLTRs to CGOs. S4A shows associations between neoplasia or malignancy prevalence and the average proximity of L1s or combined L1s and SINEs to CGOs. S4B includes longevity as a covariate. All variables are Tukey-transformed, and predictor variables are min-max normalized to the [0, 1] range to facilitate comparison of effect sizes and visualizations across models.

**Supplementary Table S4A**

| Response Variable | Predictor | Number of Species | Corr | Slope | Standard Error (SE) | *t-value* | *p-value* | $R^2$ | $\lambda$ |
|---|---|---|---|---|---|---|---|---|---|
| Neoplasia | Proximity of L1 (Ave) | 45 | 0.41 | 0.46 | 0.03 | 13.93 | 0.006* | 0.16 | 0.00006 |
| | Proximity of L1 and SINEs (Ave) | 45 | 0.34 | 0.23 | 0.1 | 2.39 | 0.02* | 0.12 | 0.00006 |

| | | | | | | | | | |
|---|---|---|---|---|---|---|---|---|---|
| Malignancy | Proximity of L1 (Ave) | 45 | 0.28 | 0.15 | 0.08 | 1.88 | 0.07 | 0.08 | 0.00006 |
| | Proximity of L1 and SINEs (Ave) | 45 | 0.3 | 0.17 | 0.08 | 2.07 | 0.045* | 0.09 | 0.00006 |

**Supplementary Table S4B**

| Response Variable | Predictor | Number of Species | Corr | Slope | Standard Error (SE) | *t-value* | Adj. *p-value* (FDR) | Adj. $R^2$ | $\lambda$ |
|---|---|---|---|---|---|---|---|---|---|
| Neoplasia | Proximity of L1 (Ave) | 45 | 0.41 | 0.26 | 0.09 | 2.77 | 0.01* | 0.13 | 0.00006 |
| | Longevity | | 0.11 | 0.00001 | 0.0001 | 0.17 | 0.86 | | |
| | Proximity of L1 and SINEs (Ave) | 45 | 0.34 | 0.225 | 0.1 | 2.29 | 0.04* | 0.08 | 0.00006 |
| | Longevity | | 0.11 | 0.00005 | 0.0001 | 0.52 | 0.6 | | |
| Malignancy | Proximity of L1 (Ave) | 45 | 0.28 | 0.18 | 0.08 | 2.18 | 0.052 | 0.08 | 0.00006 |
| | Longevity | | -0.14 | -0.0001 | 0.00008 | -1.44 | 0.15 | | |
| | Proximity of L1 and SINEs (Ave) | 45 | 0.3 | 0.18 | 0.08 | 2.2 | 0.049* | 0.08 | 0.00006 |
| | Longevity | | -0.14 | -0.0001 | 0.00008 | -1.22 | 0.23 | | |

**Supplementary Table S5.** PGLS results for the Proximity Model based on the median distance of active nLTRs to CGOs. S5A presents associations between neoplasia or malignancy prevalence and the median proximity of L1s or combined L1s and SINEs to CGOs in univariate models. S5B includes longevity as a covariate. All variables are Tukey-transformed, and predictor variables are min-max normalized to the [0, 1] range to maintain consistency in effect size interpretation and visualization across models.

**Supplementary Table S5A**

| Response Variable | Predictor | Number of Species | Corr | Slope | Standard Error (SE) | *t-value* | *p-value* | $R^2$ | $\lambda$ |
|---|---|---|---|---|---|---|---|---|---|
| Neoplasia | Proximity of L1 (Med) | 45 | 0.37 | 0.24 | 0.09 | 2.62 | 0.01* | 0.14 | 0.00006 |
| | Proximity of L1 and SINEs (Med) | 45 | 0.3 | 0.2 | 0.1 | 2.06 | 0.045* | 0.09 | 0.00006 |
| Malignancy | Proximity of L1 (Med) | 45 | 0.23 | 0.13 | 0.08 | 1.54 | 0.13 | 0.05 | 0.00006 |
| | Proximity of L1 and SINEs (Med) | 45 | 0.27 | 0.15 | 0.08 | 1.82 | 0.075 | 0.07 | 0.00006 |

**Supplementary Table S5B**

| Response Variable | Predictor | Number of Species | Corr | Slope | Standard Error (SE) | *t-value* | Adj. *p-value* (FDR) | Adj. $R^2$ | $\lambda$ |
|---|---|---|---|---|---|---|---|---|---|
| Neoplasia | Proximity of L1 (Med) | 45 | 0.37 | 0.24 | 0.1 | 2.47 | 0.026* | 0.1 | 0.00006 |
| | Longevity | | 0.11 | 0.00002 | 0.0001 | 0.15 | 0.88 | | |

| | | | | | | | | | |
|---|---|---|---|---|---|---|---|---|---|
| | Proximity of L1 and SINEs (Med) | 45 | 0.3 | 0.19 | 0.1 | 1.97 | 0.08 | 0.05 | 0.00006 |
| | Longevity | | 0.11 | 0.00005 | 0.0001 | 0.53 | 0.59 | | |
| Malignancy | Proximity of L1 (Med) | 45 | 0.23 | 0.15 | 0.08 | 1.86 | 0.1 | 0.05 | 0.00006 |
| | Longevity | | -0.14 | -0.0001 | 0.00008 | -1.4 | 0.17 | | |
| | Proximity of L1 and SINEs (Med) | 45 | 0.27 | 0.16 | 0.08 | 1.96 | 0.08 | 0.06 | 0.00006 |
| | Longevity | | -0.14 | -0.0001 | 0.00008 | -1.19 | 0.24 | | |

**Supplementary Table S6.** PGLS results for the Genic-Insertion Model based on the number of active nLTR insertions within PC Genes. S6A reports associations between neoplasia or malignancy prevalence and the number of L1 or combined L1 and SINE insertions intersecting PC Genes. S6B includes longevity as a covariate in multivariate PGLSs. All variables are Tukey-transformed, and predictor variables are min-max normalized to the [0, 1] range to ensure consistency in effect size interpretation and visualization across models.

**Supplementary Table S6A**

| Response Variable | Predictor | Number of Species | Corr | Slope | Standard Error (SE) | *t-value* | *p-value* | $R^2$ | $\lambda$ |
|---|---|---|---|---|---|---|---|---|---|

| | | | | | | | | | |
|---|---|---|---|---|---|---|---|---|---|
| Neoplasia | Number of L1 Insertions within PC Genes | 46 | 0.43 | 0.19 | 0.06 | 3.11 | 0.003* | 0.18 | 0.00006 |
| | Number of L1 and SINE Insertions within PC Genes | 46 | 0.36 | 0.21 | 0.08 | 2.55 | 0.01* | 0.13 | 0.00006 |
| Malignancy | Number of L1 Insertions within PC Genes | 46 | 0.21 | 0.08 | 0.05 | 1.46 | 0.15 | 0.05 | 0.00006 |
| | Number of L1 and SINE Insertions within PC Genes | 46 | 0.29 | 0.14 | 0.07 | 2.02 | 0.0499* | 0.08 | 0.00006 |

**Supplementary Table S6B**

| Response Variable | Predictor | Number of Species | Corr | Slope | Standard Error (SE) | *t-value* | Adj. *p-value* (FDR) | Adj. $R^2$ | $\lambda$ |
|---|---|---|---|---|---|---|---|---|---|
| Neoplasia | Number of L1 Insertions within PC Genes | 46 | 0.43 | 0.2 | 0.06 | 3.05 | 0.006* | 0.14 | 0.00006 |
| | Longevity | | 0.07 | -0.00003 | 0.0001 | -0.27 | 0.78 | | |

| | | | | | | | | | |
|---|---|---|---|---|---|---|---|---|---|
| | Number of L1 and SINE Insertions within PC Genes | 46 | 0.36 | 0.21 | 0.08 | 2.47 | 0.03* | 0.09 | 0.00006 |
| | Longevity | | 0.07 | 0.0000002 | 0.0001 | 0.002 | 0.99 | | |
| Malignancy | Number of L1 Insertions within PC Genes | 46 | 0.21 | 0.1 | 0.07 | 1.75 | 0.13 | 0.04 | 0.00006 |
| | Longevity | | -0.13 | -0.0001 | -0.00008 | -1.32 | 0.19 | | |
| | Number of L1 and SINE Insertions within PC Genes | 46 | 0.29 | 0.16 | 0.07 | 2.25 | 0.044* | 0.08 | 0.00006 |
| | Longevity | | -0.13 | -0.0001 | 0.00008 | -1.34 | 0.18 | | |

**Supplementary Table S7.** PGLS results for the Genic-Insertion Model based on the number of active nLTR insertions within CGOs. S7A presents associations between neoplasia or malignancy prevalence and the number of L1 or combined L1 and SINE insertions overlapping CGOs. S7B includes longevity as a covariate. All variables are Tukey-transformed, and predictor variables are min-max normalized to the [0, 1] range to ensure consistency in effect size interpretation and visualization across models.

**Supplementary Table S7A**

| Response Variable | Predictor | Number of Species | Corr | Slope | Standard Error (SE) | *t-value* | *p-value* | $R^2$ | *λ* |
|---|---|---|---|---|---|---|---|---|---|
| Neoplasia | Number of L1 Insertions within PC Genes | 46 | 0.08 | 0.03 | 0.06 | 0.51 | 0.61 | 0.006 | 0.00006 |
| | Number of L1 and SINE Insertions within PC Genes | 46 | 0.13 | 0.07 | 0.07 | 0.9 | 0.37 | 0.02 | 0.00006 |
| Malignancy | Number of L1 Insertions within PC Genes | 46 | -0.04 | -0.01 | 0.05 | -0.27 | 0.79 | 0.001 | 0.00006 |
| | Number of L1 and SINE Insertions within PC Genes | 46 | 0.11 | 0.05 | 0.06 | 0.76 | 0.45 | 0.01 | 0.00006 |

**Supplementary Table S7B**

| Response Variable | Predictor | Number of Species | Corr | Slope | Standard Error (SE) | *t-value* | Adj. *p-value* (FDR) | Adj. $R^2$ | *λ* |
|---|---|---|---|---|---|---|---|---|---|

| | | | | | | | | | |
|---|---|---|---|---|---|---|---|---|---|
| Neoplasia | Number of L1 Insertions within PC Genes | 46 | 0.08 | 0.02 | 0.07 | 0.35 | 0.78 | -0.04 | 0.00006 |
| | Longevity | | 0.07 | 0.00003 | 0.0001 | 0.28 | 0.78 | | |
| | Number of L1 and SINE Insertions within PC Genes | 46 | 0.13 | 0.06 | 0.08 | 0.77 | 0.66 | -0.03 | 0.00006 |
| | Longevity | | 0.07 | 0.00001 | 0.0001 | 0.14 | 0.89 | | |
| Malignancy | Number of L1 Insertions within PC Genes | 46 | -0.04 | 0.005 | 0.055 | 0.09 | 0.93 | -0.03 | 0.00006 |
| | Longevity | | -0.13 | -0.00008 | 0.00009 | -0.84 | 0.6 | | |
| | Number of L1 and SINE Insertions within PC Genes | 46 | 0.11 | 0.08 | 0.07 | 1.18 | 0.24 | 0.004 | 0.00006 |
| | Longevity | | -0.13 | -0.0001 | 0.00009 | -1.27 | 0.24 | | |

**Supplementary Table S8.** PGLS results for the Potential Cancer Gene Load Model based on the number of CGOs per species. S8A shows associations between neoplasia or malignancy prevalence and the total number of CGOs, as well as the number of somatic, germline, oncogene, TSG, and fusion gene orthologs. S8B includes longevity as a covariate. All variables are Tukey-transformed, and predictor variables are min-max normalized to the [0, 1] range to ensure consistency in effect size interpretation and visualization across models.

**Supplementary Table S8A**

| Response Variable | Predictor | Number of Species | Corr | Slope | Standard Error (SE) | *t-value* | *p-value* | $R^2$ | $\lambda$ |
|---|---|---|---|---|---|---|---|---|---|
| Neoplasia | Total Number of CGOs | 46 | 0.32 | 0.25 | 0.09 | 2.81 | 0.007* | 0.09 | 0.58 |
| | Somatic | 46 | 0.32 | 0.26 | 0.09 | 2.8 | 0.007* | 0.09 | 0.59 |
| | Germline | 46 | 0.33 | 0.23 | 0.08 | 2.67 | 0.01* | 0.09 | 0.46 |
| | Oncogene | 46 | 0.31 | 0.25 | 0.09 | 2.74 | 0.009* | 0.09 | 0.58 |
| | TSG | 46 | 0.32 | 0.24 | 0.09 | 2.62 | 0.01* | 0.08 | 0.5 |
| | Fusion | 46 | 0.31 | 0.25 | 0.09 | 2.8 | 0.008* | 0.09 | 0.61 |
| Malignancy | Total Number of CGOs | 46 | 0.16 | 0.08 | 0.08 | 1.05 | 0.3 | 0.03 | 0.00006 |
| | Somatic | 46 | 0.15 | 0.08 | 0.08 | 1.03 | 0.31 | 0.02 | 0.00006 |
| | Germline | 46 | 0.18 | 0.09 | 0.07 | 1.21 | 0.23 | 0.03 | 0.00006 |

| | | | | | | | | | |
|---|---|---|---|---|---|---|---|---|---|
| | Oncogene | 46 | 0.16 | 0.08 | 0.08 | 1.06 | 0.29 | 0.025 | 0.00006 |
| | TSG | 46 | 0.16 | 0.08 | 0.08 | 1.09 | 0.28 | 0.03 | 0.00006 |
| | Fusion | 46 | 0.13 | 0.06 | 0.07 | 0.84 | 0.4 | 0.015 | 0.00006 |

**Supplementary Table S8B**

| Response Variable | Predictor | Number of Species | Corr | Slope | Standard Error (SE) | *t-value* | Adj. *p-value* (FDR) | Adj. $R^2$ | $\lambda$ |
|---|---|---|---|---|---|---|---|---|---|
| Neoplasia | Total Number of CGOs | 46 | 0.32 | 0.25 | 0.09 | 2.75 | 0.013* | 0.06 | 0.56 |
| | Longevity | | 0.07 | 0.00005 | 0.0001 | 0.52 | 0.6 | | |
| | Somatic | 46 | 0.32 | 0.25 | 0.09 | 2.73 | 0.013* | 0.055 | 0.57 |
| | Longevity | | 0.07 | 0.00005 | 0.0001 | 0.53 | 0.59 | | |
| | Germline | 46 | 0.33 | 0.19 | 0.08 | 2.32 | 0.037* | 0.074 | 0.00006 |
| | Longevity | | 0.07 | 0.00005 | 0.0001 | 0.55 | 0.58 | | |
| | Oncogene | 46 | 0.31 | 0.25 | 0.09 | 2.67 | 0.016* | 0.05 | 0.56 |

| | | | | | | | | | |
|---|---|---|---|---|---|---|---|---|---|
| | Longevity | | 0.07 | 0.00005 | 0.0001 | 0.5 | 0.62 | | |
| | TSG | 46 | 0.32 | 0.19 | 0.08 | 2.25 | 0.044* | 0.07 | 0.00006 |
| | Longevity | | 0.07 | 0.00006 | 0.00009 | 0.62 | 0.53 | | |
| | Fusion | 46 | 0.31 | 0.25 | 0.09 | 2.71 | 0.014* | 0.05 | 0.59 |
| | Longevity | | 0.07 | 0.00005 | 0.0001 | 0.5 | 0.61 | | |
| Malignancy | Total Number of CGOs | 46 | 0.16 | 0.07 | 0.08 | 0.97 | 0.41 | -0.0055 | 0.00006 |
| | Longevity | | -0.13 | -0.00007 | 0.00008 | -0.82 | 0.41 | | |
| | Somatic | 46 | 0.15 | 0.07 | 0.08 | 0.95 | 0.41 | -0.006 | 0.00005 |
| | Longevity | | -0.13 | -0.00007 | 0.00008 | -0.81 | 0.41 | | |
| | Germline | 46 | 0.18 | 0.08 | 0.07 | 1.18 | 0.37 | 0.0045 | 0.00006 |
| | Longevity | | -0.13 | -0.00007 | 0.00008 | -0.86 | 0.39 | | |
| | Oncogene | 46 | 0.16 | 0.08 | 0.08 | 0.1 | 0.41 | -0.0045 | 0.00007 |

| | | | | | | | | | |
|---|---|---|---|---|---|---|---|---|---|
| | Longevity | | -0.13 | -0.00007 | 0.00008 | -0.82 | 0.41 | | |
| | TSG | 46 | 0.16 | 0.08 | 0.08 | 1.03 | 0.41 | -0.003 | 0.00006 |
| | Longevity | | -0.13 | -0.00007 | 0.00008 | -0.83 | 0.41 | | |
| | Fusion | 46 | 0.13 | 0.06 | 0.07 | 0.78 | 0.44 | -0.013 | 0.00006 |
| | Longevity | | -0.13 | -0.00007 | 0.00009 | -0.84 | 0.44 | | |

**Supplementary Table S9.** Model comparison results based on the Akaike Information Criterion (AIC) for all PGLS models. S9A summarizes AIC values for univariate regression models using λ = 0, λ = 1, and λ estimated from the data. S9B presents AIC values for the corresponding multivariate regression models, including longevity as a covariate. Lower AIC values indicate better model fit.

**Supplementary Table S9A**

| Model | Predictor Variable | Neoplasia | | | Malignancy | | |
|---|---|---|---|---|---|---|---|
| | | $\lambda = 0$ | $\lambda$ = Estimated | $\lambda = 1$ | $\lambda = 0$ | $\lambda$ = Estimated | $\lambda = 1$ |
| Abundance | L1s | -71.51 | -71.51 | -11.88 | -88.18 | -88.18 | -26.84 |
| | L1s and SINEs | -67.45 | -67.16 | 3.71 | -90.43 | -90.43 | -19.33 |
| Proximity (Average to PC Genes) | L1s | -57.58 | -57.57 | 6.45 | -57.58 | -73.22 | -11.31 |
| | L1s and SINEs | -52.15 | -52.14 | 1.63 | -52.15 | -71.35 | -24.9 |
| Proximity (Average to CGOs) | L1s | -60.73 | -60.73 | 11.73 | -60.73 | -72.6 | -7.59 |
| | L1s and SINEs | -58.23 | -58.23 | 11.89 | -58.23 | -73.3 | -7.14 |
| | L1s | -62.68 | -62.68 | -0.71 | -62.68 | -73.86 | -16.36 |

| | | | | | | | |
|---|---|---|---|---|---|---|---|
| Genic-Insertion (PC Genes) | L1s and SINEs | -59.84 | -59.84 | 7.44 | -59.84 | -75.76 | -14.45 |
| Genic-Insertion (CGOs) | L1s | -53.78 | -53.77 | -4.95 | -53.78 | -71.77 | -20.16 |
| | L1s and SINEs | -54.33 | -54.33 | -16.28 | -54.33 | -72.3 | -35.56 |
| Potential Cancer Gene Load Model | Total Number of CGOs | -61.01 | -60.52 | -11.43 | -70.21 | -60.52 | -19.29 |
| | Somatic | -60.93 | -60.44 | -11.88 | -70.18 | -70.17 | -19.66 |
| | Germline | -61.38 | -60.28 | -5.03 | -70.59 | -70.58 | -14.49 |
| | Oncogene | -60.81 | -60.18 | -10.03 | -70.25 | -70.24 | -17.65 |
| | TSG | -60.96 | -59.89 | -6.24 | -70.31 | -70.3 | -15.82 |
| | Fusion | -60.56 | -60.26 | -13.16 | -69.81 | -69.8 | -20.24 |

**Supplementary Table S9B**

| Model | Predictor Variables | Neoplasia | | | Malignancy | | |
|---|---|---|---|---|---|---|---|
| | | $\lambda = 0$ | $\lambda$ = Estimated | $\lambda = 1$ | $\lambda = 0$ | $\lambda$ = Estimated | $\lambda = 1$ |
| Abundance | L1s + Longevity | -70.35 | -70.36 | -9.19 | -86.33 | -86.33 | -25.12 |
| | L1s and SINEs + Longevity | -66.66 | -66.8 | 4.89 | -88.58 | -90.43 | -17.35 |
| Proximity (Average to PC Genes) | L1s + Longevity | -55.66 | -55.65 | 7.74 | -72.25 | -72.25 | -9.32 |
| | L1s and SINEs + Longevity | -50.42 | -50.41 | 3.29 | -70.03 | -71.35 | -23 |
| Proximity (Average to CGOs) | L1s + Longevity | -58.76 | -58.76 | 12.83 | -72.78 | -72.78 | -5.61 |
| | L1s and SINEs + Longevity | -56.52 | -56.52 | 13.14 | -72.87 | -72.88 | -5.14 |

| | | | | | | | |
|---|---|---|---|---|---|---|---|
| Genic-Insertion (PC Genes) | L1s + Longevity | -60.76 | -60.76 | 1.23 | -73.68 | -73.68 | -15.26 |
| | L1s and SINEs + Longevity | -57.84 | -57.84 | 8.72 | -75.66 | -75.76 | -12.47 |
| Genic-Insertion (CGOs) | L1s + Longevity | -51.86 | -51.86 | -3.79 | -70.53 | -70.52 | -18.16 |
| | L1s and SINEs + Longevity | -52.35 | -52.35 | -14.88 | -71.99 | -72.3 | -33.6 |
| Potential Cancer Gene Load Model | Total Number of CGOs + Longevity | -59.46 | -58.81 | -9.69 | -68.93 | -68.92 | -18.95 |
| | Somatic + Longevity | -59.39 | -58.74 | -10.12 | -68.89 | -68.89 | -19.27 |
| | Germline + Longevity | -59.39 | -59.7 | -10.12 | -69.39 | -69.38 | -14.14 |
| | Oncogene + Longevity | -59.23 | -58.44 | -8.26 | -68.98 | -68.97 | -17.15 |
| | TSG + Longevity | -59.37 | -59.37 | 4.41 | -69.04 | -69.04 | -15.31 |
| | Fusion + Longevity | -58.95 | -58.52 | -11.49 | -68.57 | -68.56 | -20.06 |

**Supplementary Table S10.** PGLS results for the association between species longevity and nLTR abundance. The table reports models using either the number of active L1s or the combined number of L1s and SINEs as predictors. Both the response and predictor variables are Tukey-transformed and min-max normalized to the [0, 1] range to ensure consistency in effect size interpretation and visualization across models.

| Response Variable | Predictor | Number of Species | Corr | Slope | Standard Error (SE) | *t-value* | *p-value* | $R^2$ | $\lambda$ |
|---|---|---|---|---|---|---|---|---|---|
| Longevity | Number of L1 | 55 | 0.02 | 0.1 | 0.08 | 1.28 | 0.2 | 0.42 | 1 |

| | | | | | | | | | |
|---|---|---|---|---|---|---|---|---|---|
| | Number of L1 and SINEs | 55 | 0 | -0.06 | 0.1 | -0.63 | 0.53 | 0.41 | 1 |

**Supplementary Table S11.** PGLS results for the association between species longevity and the number of active nLTR insertions within PC Genes. Models use either the number of L1 insertions or the combined number of L1 and SINE insertions as predictors. Both the response and predictor variables are Tukey-transformed and min-max normalized to the [0, 1] range to ensure consistency in effect size interpretation and visualization across models.

| Response Variable | Predictor | Number of Species | Corr | Slope | Standard Error (SE) | *t-value* | *p-value* | $R^2$ | $\lambda$ |
|---|---|---|---|---|---|---|---|---|---|
| Longevity | Number of L1 Insertions within PC Genes | 46 | 0.12 | 0.15 | 0.08 | 1.91 | 0.06 | 0.4 | 1 |
| | Number of L1 and SINE Insertions within PC Genes | 46 | 0.09 | -0.01 | 0.12 | -0.09 | 0.92 | 0.36 | 1 |

**Supplementary Table S12.** PGLS results for the association between the number of fusion genes and nLTR abundance. Models use either the number of active L1s or the combined number of L1s and SINEs as predictors. Both the response and predictor variables are Tukey-transformed and min-max normalized to the [0, 1] range to ensure consistency in effect size interpretation and visualization across models.

| Response Variable | Predictor | Number of Species | Corr | Slope | Standard Error (SE) | *t-value* | *p-value* | $R^2$ | $\lambda$ |
|---|---|---|---|---|---|---|---|---|---|

| | | | | | | | | | |
|---|---|---|---|---|---|---|---|---|---|
| Longevity | Number of L1 | 46 | 0.44 | 0.42 | 0.1 | 4.11 | 0.00017* | 0.38 | 1 |
| | Number of L1 and SINEs | 46 | 0.2 | 0.25 | 0.13 | 1.85 | 0.07 | 0.23 | 0.9 |

**Supplementary Table S13.** PGLS results for the association between longevity as a response and neoplasia and malignancy prevalence as predictors. Both the response and predictor variables are Tukey-transformed and min-max normalized to the [0, 1] range to ensure consistency in effect size interpretation and visualization across models.

| Response Variable | Predictor | Number of Species | Corr | Slope | Standard Error (SE) | *t-value* | *p-value* | $R^2$ | $\lambda$ |
|---|---|---|---|---|---|---|---|---|---|
| Longevity | Neoplasia | 55 | 0.05 | 0.06 | 0.1 | 0.56 | 0.58 | 0.41 | 1 |
| | Malignancy | 55 | -0.04 | 0.04 | 0.13 | 0.27 | 0.78 | 0.4 | 1 |

### Supplementary Figures

**Supplementary Figure S1.** PGLS result for the Abundance Model showing the association between malignancy prevalence and the number of active L1 elements across 55 mammalian species. The relationship is not statistically significant. All variables are Tukey-transformed; the predictor is additionally min-max normalized to the [0, 1] range for consistency in effect size interpretation and visualization with other models.

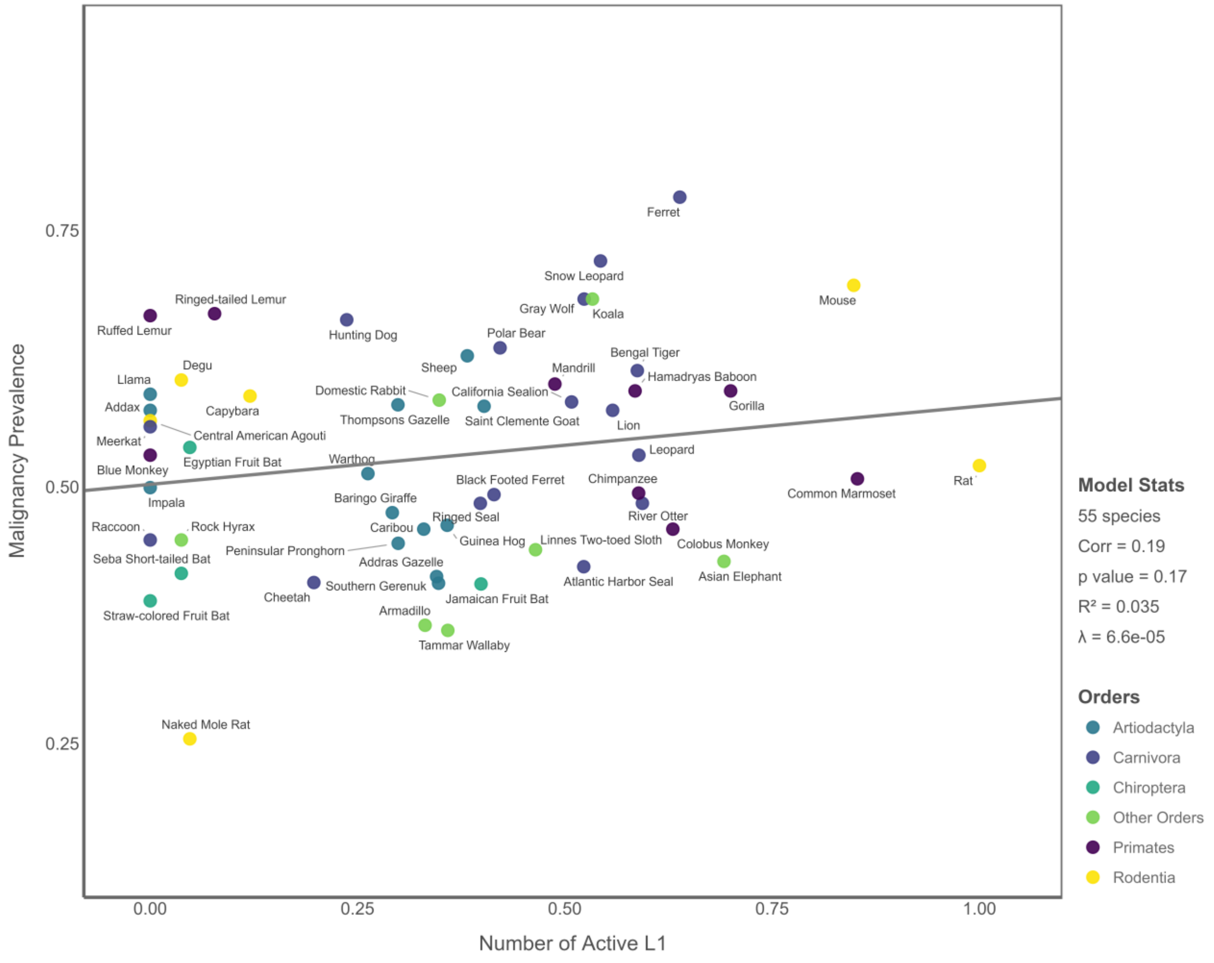

**Supplementary Figure S2.** PGLS results for the Proximity Model based on the average distance of active nLTRs to PC Genes and CGOs. None of the associations shown are statistically significant. Panels (A–C) show models testing neoplasia or malignancy prevalence in relation to the average proximity of L1s or combined L1s and SINEs to PC Genes (n = 45 species). Panel (D) shows the model testing malignancy prevalence against the average proximity of L1s to CGOs (n = 45 species). All variables are transformed using Tukey's ladder of powers. Predictors are additionally scaled to the [0, 1] range to allow consistent interpretation and visualization of effect sizes across models.

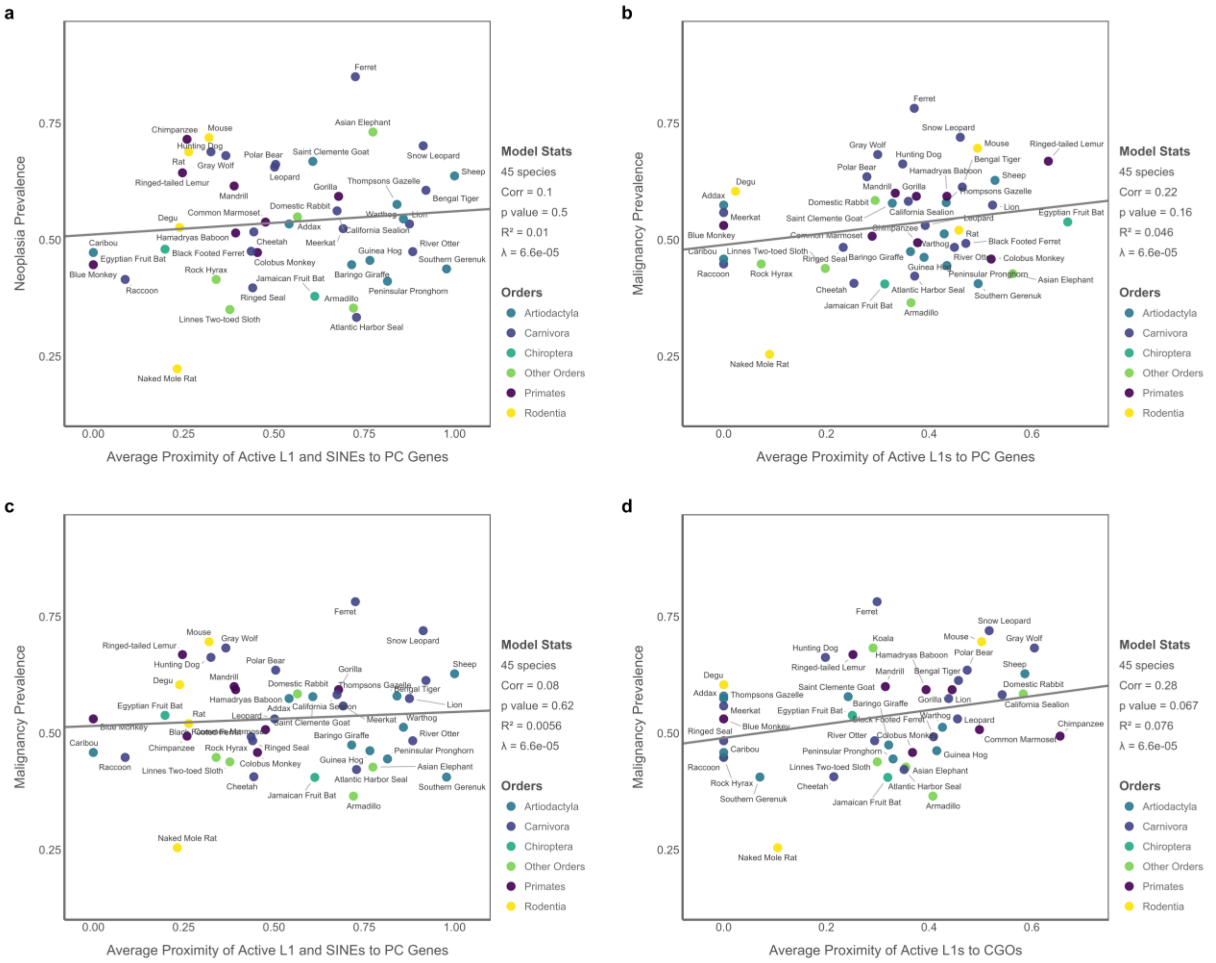

**Supplementary Figure S3.** PGLS results for the Proximity Model based on the median distance of active nLTRs to PC Genes (n = 44 species). (A) Neoplasia prevalence is significantly associated with the median proximity of active L1 elements to PC Genes. (B–D) show non-significant associations involving neoplasia and malignancy prevalences with the proximity of L1s or the combined proximity of L1s and SINEs. All variables are transformed using Tukey's ladder of powers. Predictors are additionally scaled to the [0, 1] range to allow consistent interpretation and visualization of effect sizes across models.

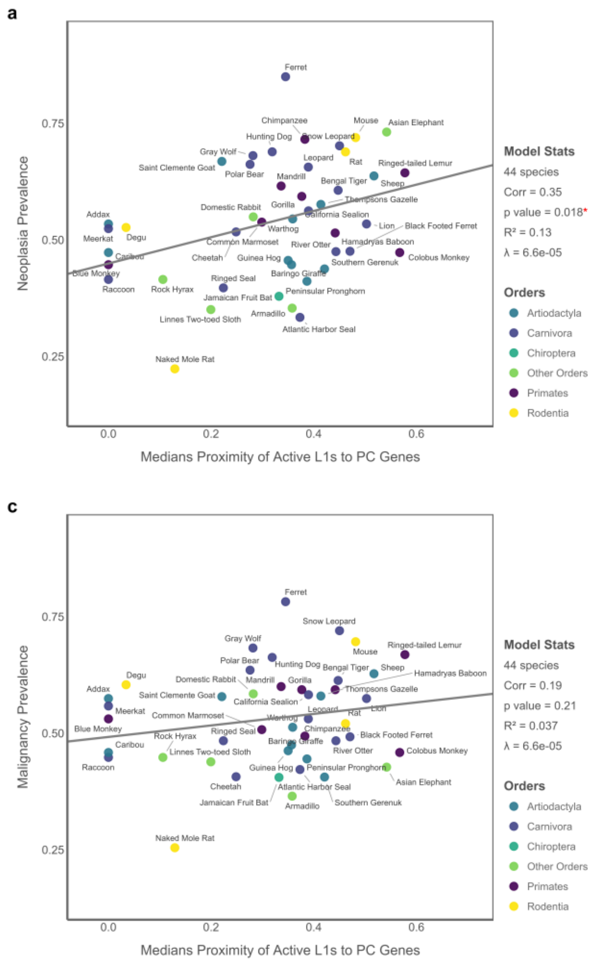

**Supplementary Figure S4.** PGLS results for the Proximity Model based on the median distance of active nLTRs to CGOs (n = 45 species). (A–B) show significant associations between neoplasia prevalence and the median proximity of either L1 elements or combined L1s and SINEs. (C–D) show non-significant associations involving malignancy prevalence. All variables are transformed using Tukey's ladder of powers, while predictors are also scaled to the [0, 1] range to ensure consistent interpretation and visualization of effect sizes across models.

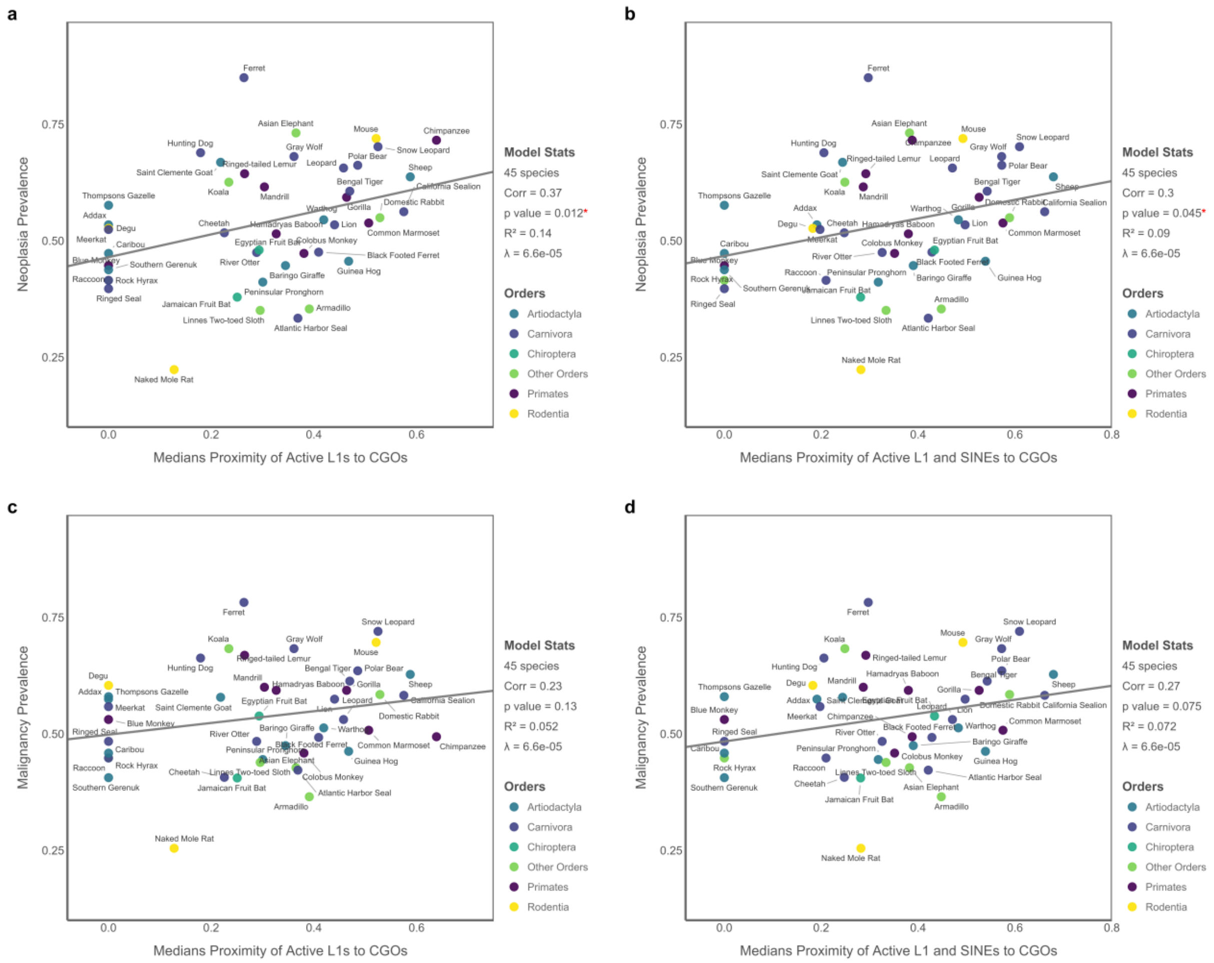

**Supplementary Figure S5.** PGLS result for the Genic-Insertion Model testing the association between malignancy prevalence and the number of active L1 insertions within PC Genes (n = 46 species). The relationship is not statistically significant. All variables are transformed using Tukey's ladder of powers. Predictor variables are also scaled to the [0, 1] range to ensure consistent interpretation and visualization of effect sizes across models.

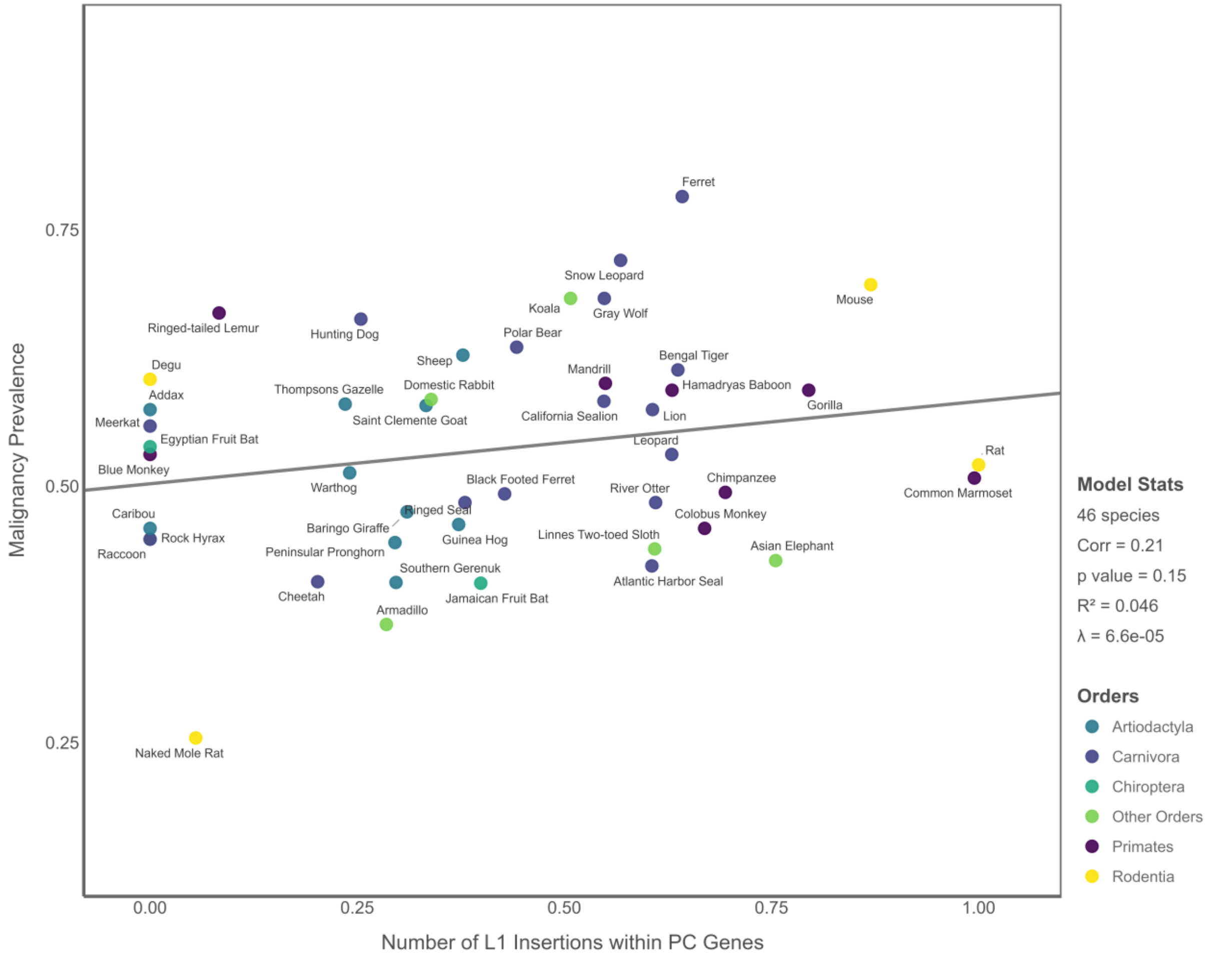

**Supplementary Figure S6.** PGLS results for the Genic-Insertion Model based on the number of active nLTR insertions within CGOs (n = 46 species). None of the associations is statistically significant. Panels (A–B) show models for neoplasia prevalence using L1 insertions alone or combined L1 and SINE insertions. Panels (C–D) show the corresponding models for malignancy prevalence. All variables are transformed using Tukey's ladder of powers. Predictor variables are also scaled to the [0, 1] range to ensure consistent interpretation and visualization of effect sizes across models.

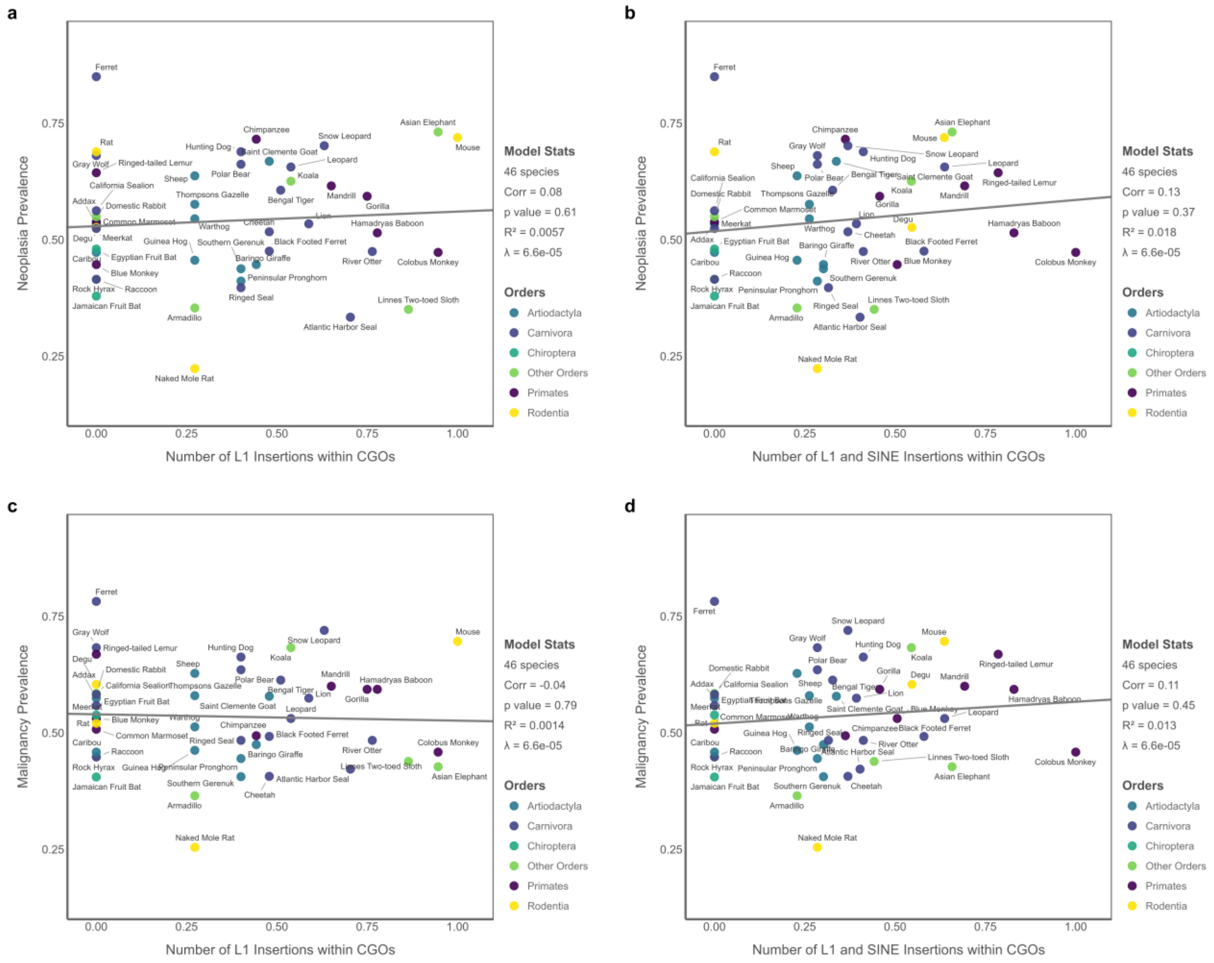

**Supplementary Figure S7.** PGLS results for the Potential Cancer Gene Load Model testing associations between malignancy prevalence and the number of cancer gene orthologs (n = 46 species). None of the associations were statistically significant. Panels (A–F) show results for the total number of CGOs, somatic CGOs, germline CGOs, oncogene orthologs, TSG orthologs, and fusion gene orthologs, respectively. All variables were transformed using Tukey's ladder of powers, and predictor variables were further scaled to the [0, 1] range to ensure consistent interpretation and visualization of effect sizes across models.

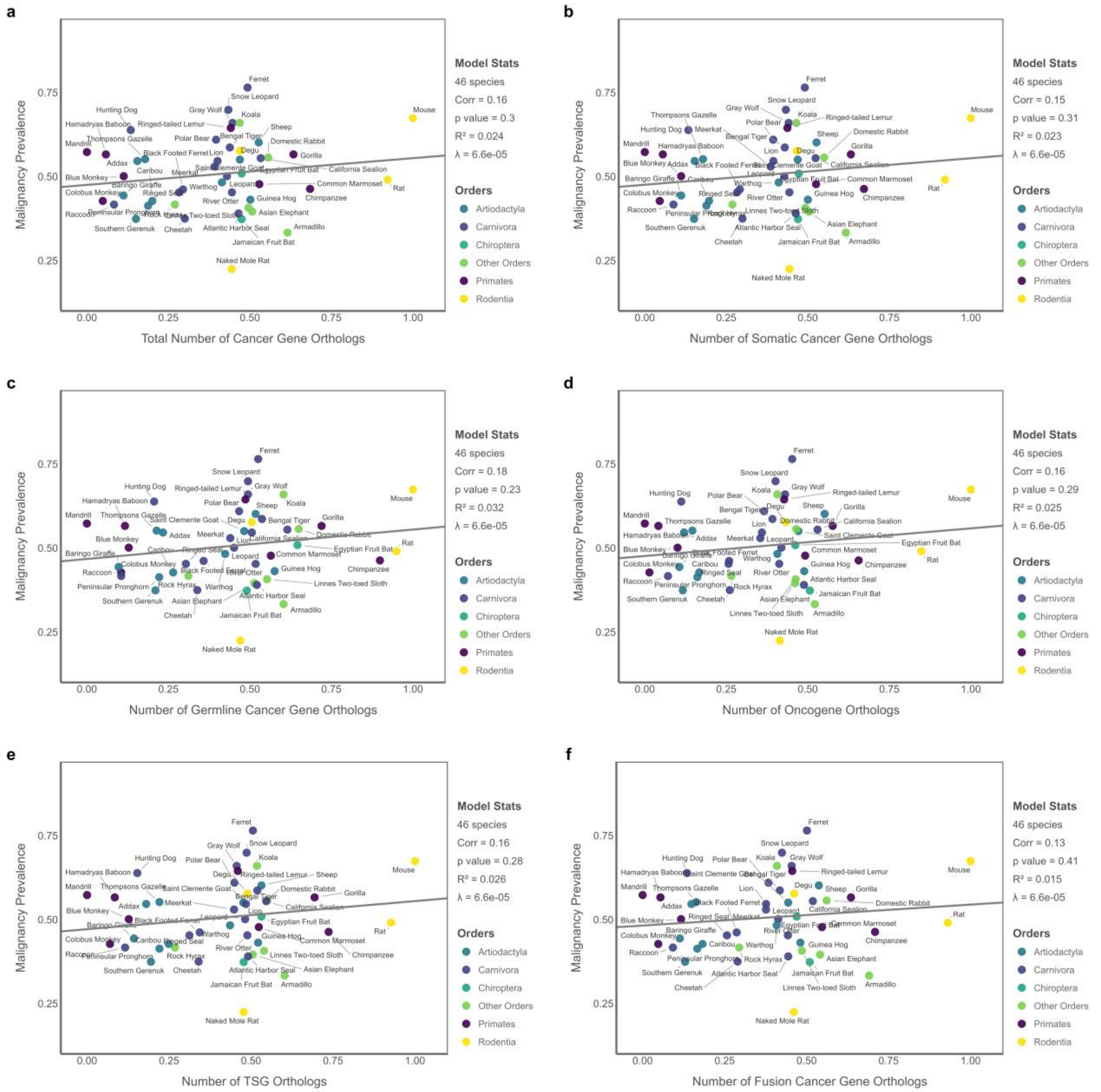

**Supplementary Figure S8.** PGLS results showing the relationships between species longevity and various nLTR-related predictors. None of the associations is statistically significant. Panels (A–B) show models using the number of active L1s and the combined number of L1s and SINEs (n = 55 species). Panels (C–D) show models using the number of active L1 or combined L1 and SINE insertions within PC Genes (n = 46 species). All variables are transformed using Tukey's ladder of powers and scaled to the [0, 1] range to ensure consistent interpretation and visualization of effect sizes across models.

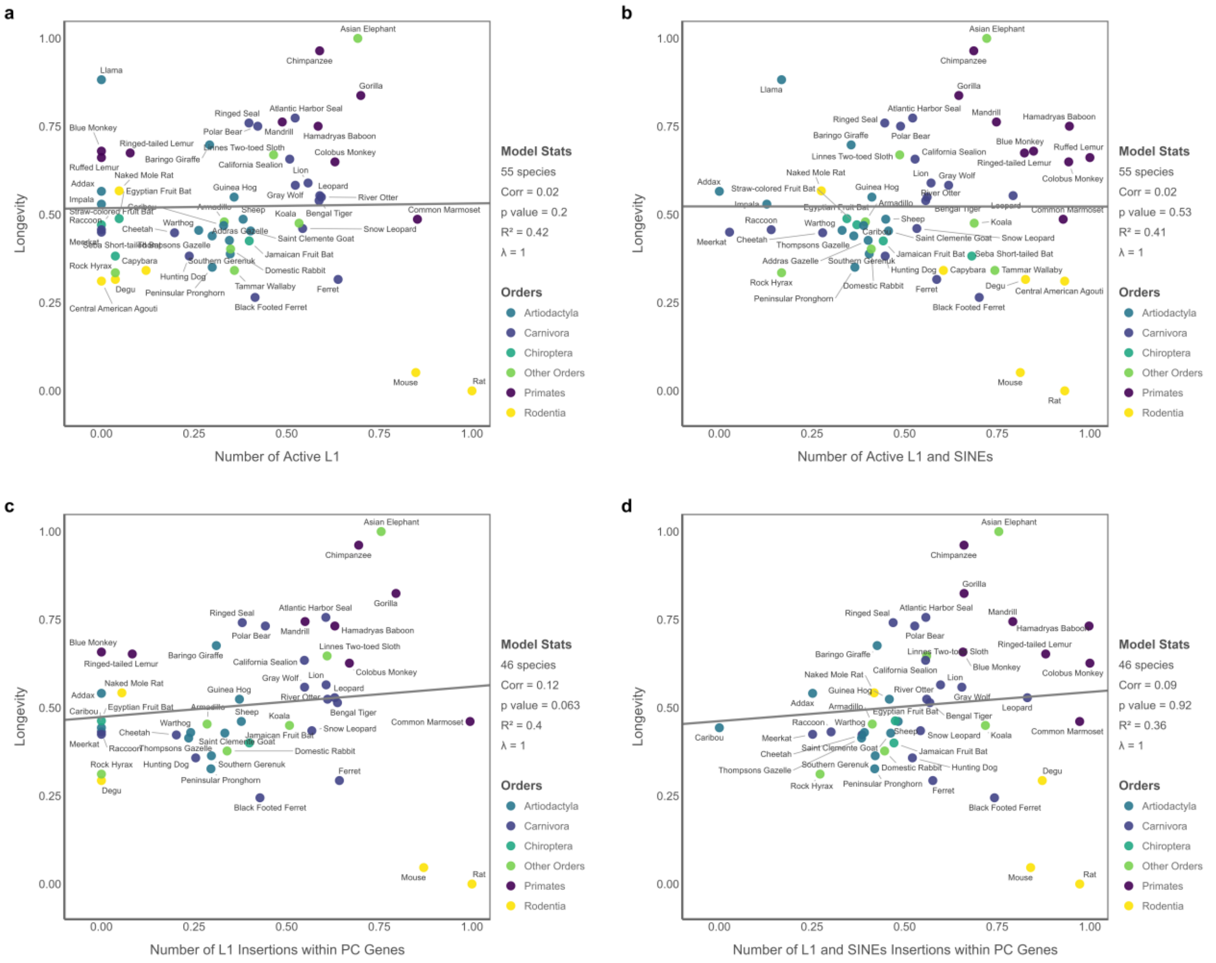

**Supplementary Figure S9.** PGLS result testing the association between the combined abundance of active L1s and SINEs and the number of fusion genes across 46 mammalian species. The relationship is not statistically significant. All variables are transformed using Tukey's ladder of powers and scaled to the [0, 1] range to ensure consistent interpretation and visualization of effect sizes across models.

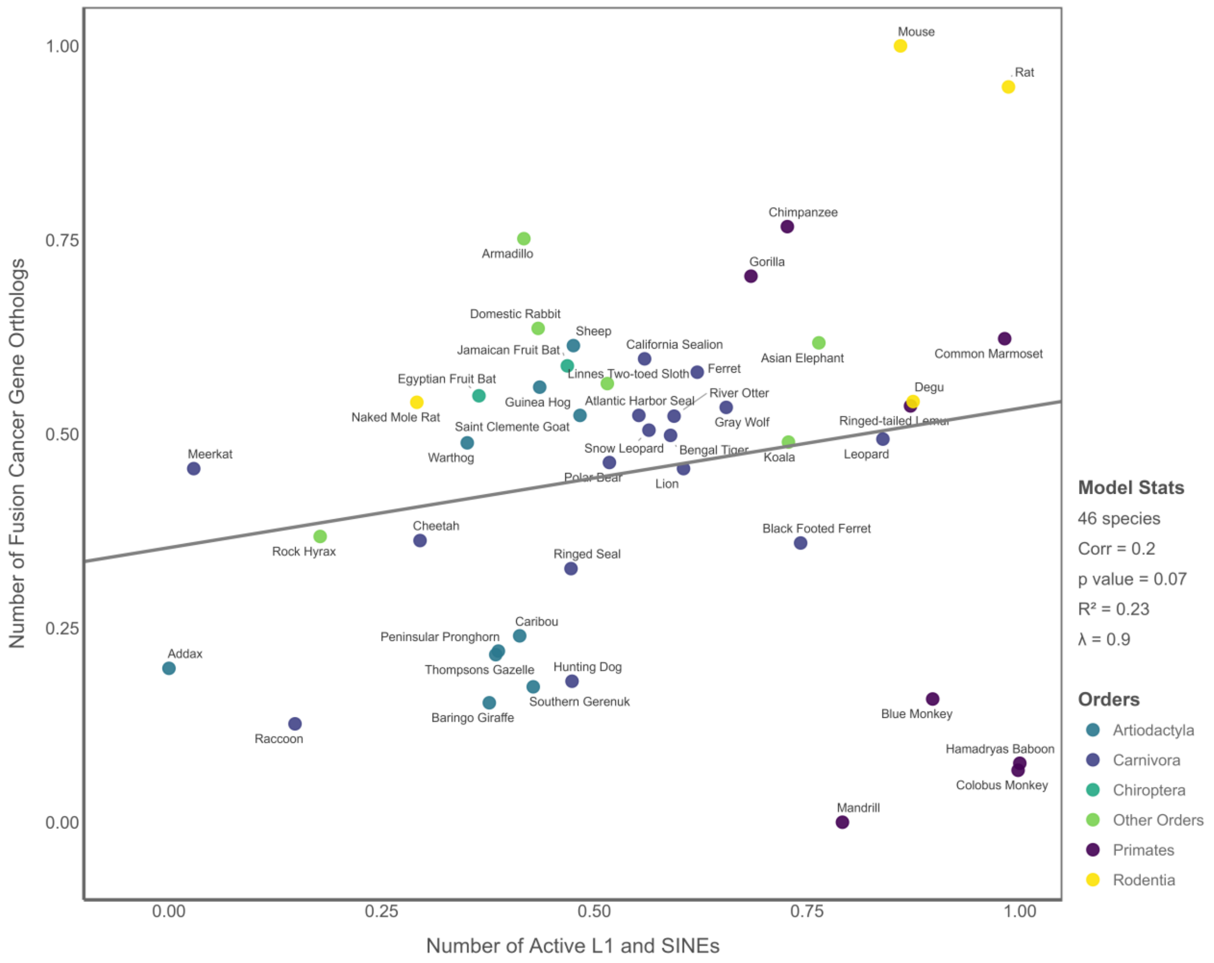

**Supplementary Figure S10.** PGLS results testing the association between longevity and (A) neoplasia and (B) malignancy prevalence across 55 mammals. The relationships are not statistically significant. All variables are transformed using Tukey's ladder of powers and scaled to the [0, 1] range to ensure consistent interpretation and visualization of effect sizes across models.

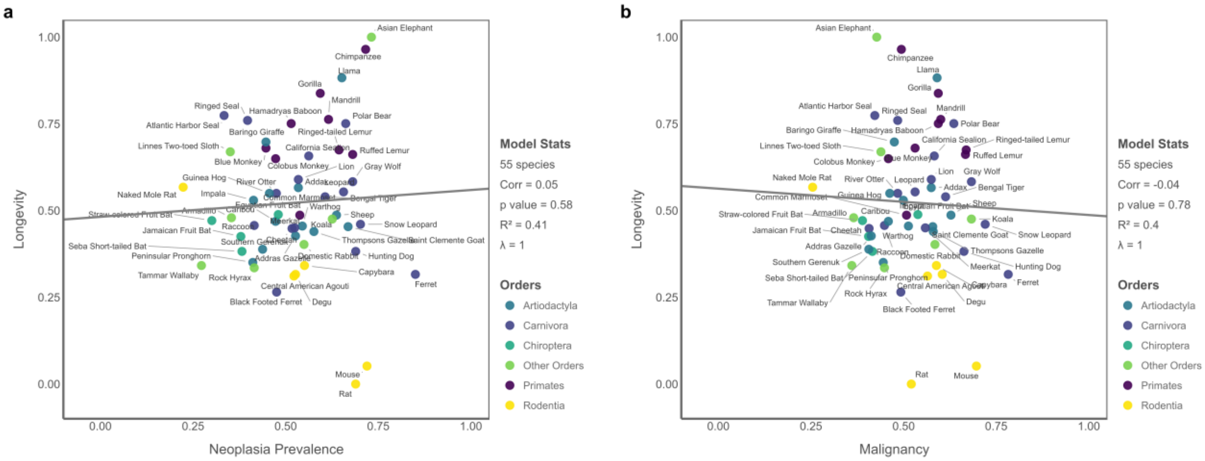


**Supplementary Figure S11.** (A) Relationship between number of L1s and number of L1 insertions within PC Genes, and (B) the number of L1 and SINEs versus the number of L1 and SINE insertions within PC Genes across 46 mammals. All variables are transformed using Tukey's ladder of powers and scaled to the [0, 1] range.

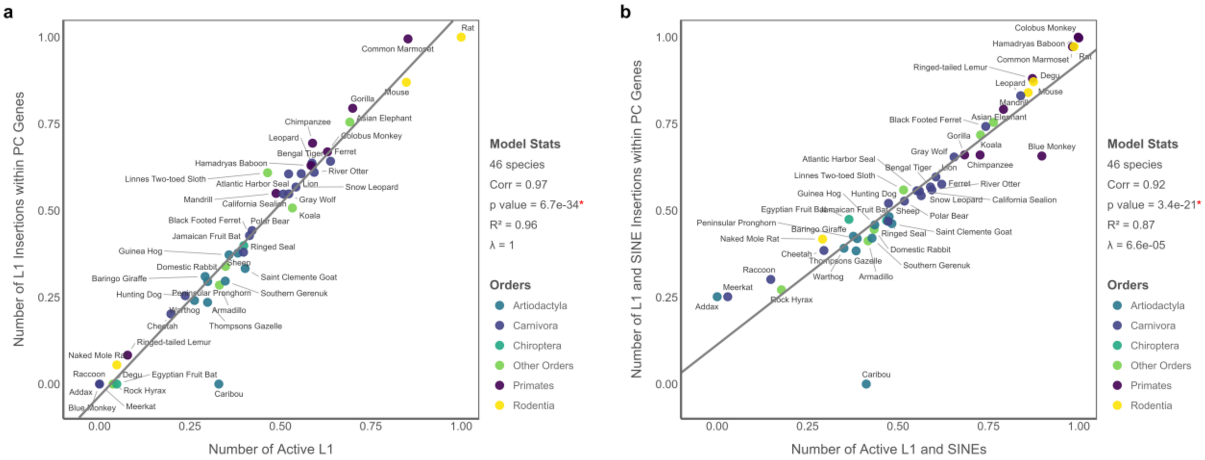

**Supplementary Figure S12.** Relationship between the number of TSGs and Oncogenes. Both variables are scaled to the [0, 1] range.

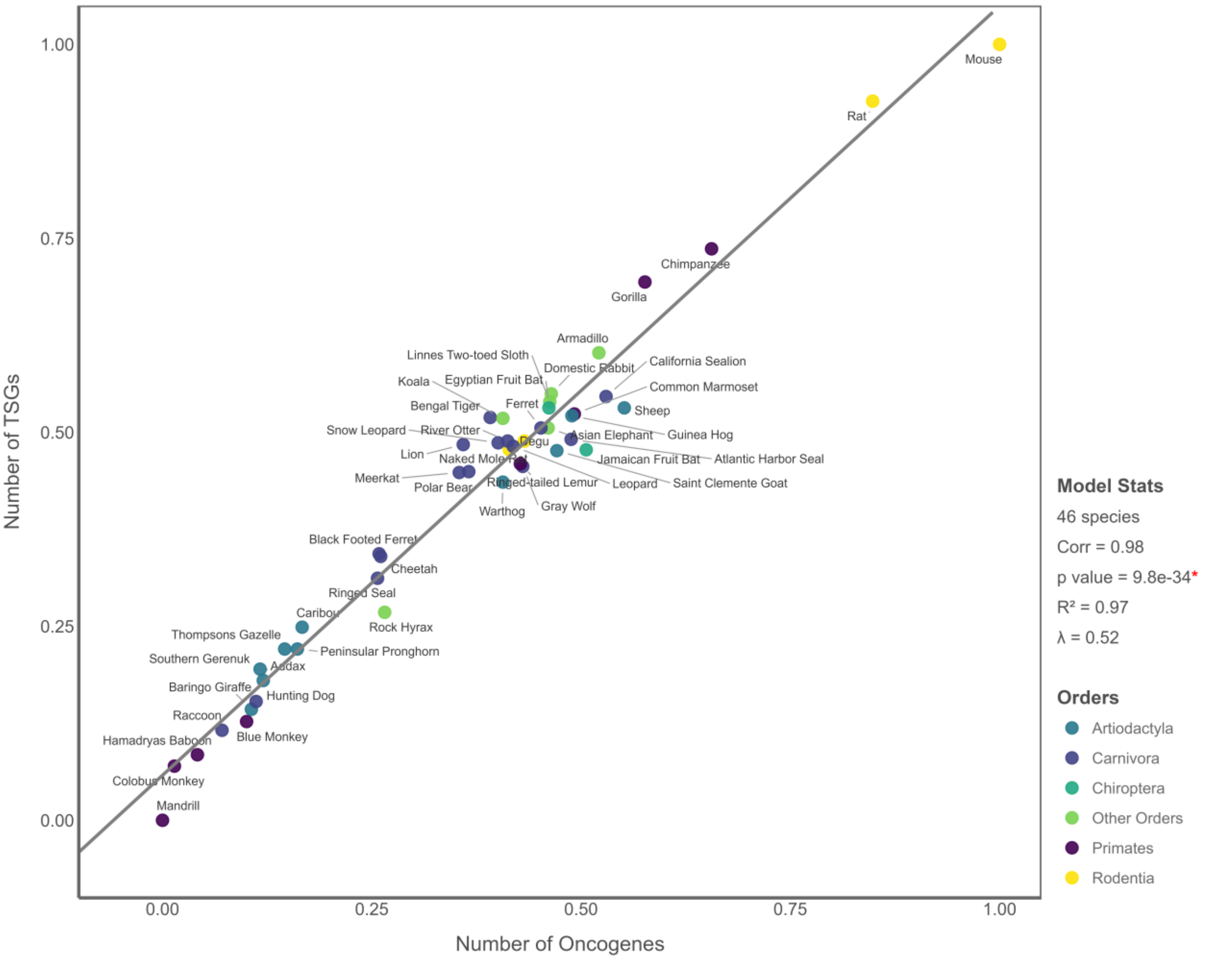
