## Supplementary Scripts for "Recent Non-LTR Retrotransposon Activity Predicts Cancer Prevalence in Mammals": 0_calculate_lengths.py

```
import os

# Specify the directory containing the BED files
directory = '/Path to the directory of BED files'

# Loop over each file in the directory
for filename in os.listdir(directory):
    if filename.endswith('.bed'):
        # Construct the full file path
        file_path = os.path.join(directory, filename)
        
        # Process each file
        with open(file_path, 'r') as file:
            lines = file.readlines()
        
        # Process each line
        with open(file_path, 'w') as file:
            for line in lines:
                if line.startswith('#'):  # Skip header or commented lines
                    continue
                parts = line.strip().split('\t')
                
                # Calculate the length of the element (abs(3rd col - 2nd col))
                if len(parts) >= 10:  # Ensure there are enough columns
                    length = abs(int(parts[2]) - int(parts[1]))
                    parts[9] = str(length)  # Replace the 10th column
                
                # Write the updated line back to file
                file.write('\t'.join(parts) + '\n')

print('Lengths calculated and files updated.')
```
