## Supplementary Scripts for "Recent Non-LTR Retrotransposon Activity Predicts Cancer Prevalence in Mammals": 1_subset_BED.sh

```
# Directory containing the BED files
directory="Path to the BED files directory"

# Loop over each BED file in the directory
for file in "$directory"/*.bed; do
    # Define the new filename with SINE suffix
    new_file="${file%.bed}_All_SINE.bed"
    
    # Subset entries where the 11th column is 'SINE\' and save to new file
    awk '$11 ~ /^SINE\//' "$file" > "$new_file"

done

echo "Subsetting complete. Files saved with specified suffix"

# NOTE: You need to do the same for LINE/L1 elements
```
