## Supplementary Scripts for "Recent Non-LTR Retrotransposon Activity Predicts Cancer Prevalence in Mammals": 2_length_Divergence_LINE1s.sh

```
# Redirect output to results.txt and write the header
echo -e "Species\tL1_Div_2\tL1_Div_5\tL1_Div_10\tL1_Div_20" > L1_results.txt

# Directory containing the _L1.bed files
directory="Path to the directory containing _L1.bed files"

# Loop over each _L1.bed file in the directory
for file in "$directory"/*_L1.bed; do
    # Extract species name from filename by removing path and _L1.bed extension
    species=$(basename "$file" _L1.bed)

    # Prepare a line to append to the results file
    output_line="$species"

    # Loop through divergence rate thresholds
    for divergence in 2 5 10 20; do
        # Count entries matching both length and divergence criteria (i.e., 6.1 kb for L1)
        count=$(awk -v div=$divergence '$10 > 5490 && $10 < 6710 && $7 <= div' "$file" | wc -l)

        # Append count to the output line
        output_line="$output_line\t$count"
    done

    # Write the complete line to results.txt
    echo -e "$output_line" >> L1_results.txt
done
```
