## Supplementary Scripts for "Recent Non-LTR Retrotransposon Activity Predicts Cancer Prevalence in Mammals": 3_length_Divergence_ALL_SINES.sh

```
# Redirect output to results.txt and write the header
echo -e "Species\tAllSINE_Div_2\tAllSINE_Div_5\tAllSINE_Div_10\tAllSINE_Div_20" > AllSINE_results.txt

# Directory containing the _All_SINE.bed files
directory="Path to the directory containing the _All_SINE.bed files"

# Loop over each  file in the directory
for file in "$directory"/*_All_SINE.bed; do
    # Extract species name from filename by removing path and  extension
    species=$(basename "$file" _All_SINE.bed)

    # Prepare a line to append to the results file
    output_line="$species"

    # Loop through divergence rate thresholds
    for divergence in 2 5 10 20; do
        # Count entries matching both length and divergence criteria (i.e., between 100 bp to 400 bp for SINEs)
        count=$(awk -v div=$divergence '$10 > 100 && $10 < 400 && $7 <= div' "$file" | wc -l)

        # Append count to the output line
        output_line="$output_line\t$count"
    done

    # Write the complete line to results.txt
    echo -e "$output_line" >> AllSINE_results.txt
done
```
