## Supplementary Scripts for "Recent Non-LTR Retrotransposon Activity Predicts Cancer Prevalence in Mammals": 4_TE_to_Genes_distance_script.sh

```
#!/bin/bash

#SBATCH --job-name=nLTRDistance
#SBATCH --array=1-55
#SBATCH --cpus-per-task=1
#SBATCH --time=12:00:00
#SBATCH --mail-type=all
#SBATCH --mail-user="Enter Email"
#SBATCH --output="Insert Path"
#SBATCH --error="Insert Path"
#SBATCH --mem=32000

##################################################################################################################################################################################
# "Description:"
#   This script takes the rmsk2bed output (RepeatMasker .out files converted to .bed) and a gene annotation file (essentially in .bed format)  
#   to calculate the distance of each TE insertion 
#   (e.g., L1, SINEs) to the nearest gene (i.e., PC gene or CGO). It also replaces the 10th column with element lengths, filters entries based on 
#   TE type (i.e., L1 or SINEs), divergence, and length thresholds, and outputs cleaned distance tables for downstream analyses.
#
#   Requirements:
#     - TE_file: RepeatMasker annotations (rmsk2bed output) in bed format (e.g., Mus_musculus_GRCm39_genomic.bed)
#     - gene_file: bed annotation containing only protein coding genes or only cancer gene orthologs
#     - species_list.txt: plain text list of species names (Genus_species), one per line, located in the same directory
#
#   Output:
#     - For each species, results are written to a dedicated subdirectory.
#     - Final files include two files (i.e., for L1 and SINEs) each one includes three columns representing the superfamily, family, 
#       and the distance to the nearest gene(e.g., 6_L1_subset.txt).
#
#   Notes:
#     - Distances are calculated with bedtools closest (-d -t all -io -D a).
#     - Divergence is taken from column 7; element lengths are recalculated as (end – start) and stored in column 10.
#     - Negative distances (upstream) are converted to absolute values in later steps.
#	  - This script is designed to submit as a job array on slurm
##################################################################################################################################################################################

export species_name=$(awk "NR==${SLURM_ARRAY_TASK_ID}" species_list.txt)
export TE_file=/"path to bed rmsk2bed outputs"/${species_name}\.bed
export gene_file=/"path to bed annotations"/${species_name}_CGOs.bed
export results=/"Path to the envisioned output directory"

cd ${results} && mkdir -p ${species_name} && cd ${species_name}

## Remove occasional extra columns including asterisks from TE annotation files (some files have 16th column that must be removed)
if awk '{ if (sub(/\*$/, "") && NF == 15) $0 = gensub(/[ \t]+$/, "", "g"); print }' "$TE_file" > TE_file_asterisks_removed; then
    echo "Removed asterisks from TE annotation file successfully."
else
    echo "Error: Failed to remove asterisks from TE annotation file."
    exit 1
fi

# Calculate gene-TE closest distances
if bedtools closest -d -a TE_file_asterisks_removed -b "${gene_file}" -t all -io -D a > gene_te_closest_distances.txt; then
    echo "Calculated gene-TE closest distances successfully."
else
    echo "Error: Failed to calculate gene-TE closest distances."
    exit 1
fi

# Step to calculate the length of each TE element and replace the 10th column with the calculated length
if awk '{OFS="\t"; $10 = $3 - $2; print}' gene_te_closest_distances.txt > 1_modified_gene_te_closest_distances.txt; then
    echo "Successfully calculated the length of each TE element and updated the 10th column."
else
    echo "Error: Failed to calculate and update the lengths."
    exit 1
fi

# Step to keep only specific entries (L1 and SINEs) based on the 11th column and save the result
if awk '$11 == "LINE/L1" || $11 ~ /^SINE\//' 1_modified_gene_te_closest_distances.txt > 2_modified_gene_te_closest_distances.txt; then
    echo "Successfully filtered and saved the specified entries."
else
    echo "Error: Failed to filter the entries."
    exit 1
fi

# Step to remove entries where the divergence (7th column) is greater than 5
if awk '$7 <= 5' 2_modified_gene_te_closest_distances.txt > 3_modified_gene_te_closest_distances.txt; then
    echo "Successfully filtered entries based on divergence and saved to 3_modified_gene_te_closest_distances.txt."
else
    echo "Error: Failed to filter based on divergence."
    exit 1
fi

## Step to filter entries based on the 11th column and length in the 10th column
if awk '($11 ~ /^SINE\// && $10 >= 100 && $10 <= 400) || \
        ($11 == "LINE/L1" && $10 >= 5450 && $10 <= 6750)' 3_modified_gene_te_closest_distances.txt > 4_final_filtered_gene_te_distances.txt; then
    echo "Successfully filtered based on specific TE types and lengths."
else
    echo "Error: Failed to filter based on TE types and lengths."
    exit 1
fi

# Step to extract subfamilies, families, and absolute values of distances (# also removing the minus signs for TEs in upstream (5' end) of the gens)
if awk '{
    OFS="\t"; 
    # Split the 11th column into family and subfamily
    split($11, parts, "/"); 
    family = parts[1]; 
    subfamily = (length(parts) > 1) ? parts[2] : "NA"; 
    # Take the absolute value of the last column
    abs_distance = ($NF < 0) ? -$NF : $NF;
    # Print family, subfamily, and absolute value in a new file
    print family, subfamily, abs_distance
}' 4_final_filtered_gene_te_distances.txt > 5_extracted_columns.txt; then
    echo "Successfully extracted families, subfamilies, and absolute distance values."
else
    echo "Error: Failed to extract the required columns."
    exit 1
fi

# Step to create L1 subset
if awk '$2 == "L1"' 5_extracted_columns.txt > 6_L1_subset.txt; then
    echo "Successfully created L1_subset.txt."
else
    echo "Error: Failed to create L1_subset.txt."
    exit 1
fi

# Step to create All_SINE subset
if awk '$1 == "SINE"' 5_extracted_columns.txt > 6_SINE_subset.txt; then
    echo "Successfully created Alu_subset.txt."
else
    echo "Error: Failed to create Alu_subset.txt."
    exit 1
fi
```
