## Supplementary Scripts for "Recent Non-LTR Retrotransposon Activity Predicts Cancer Prevalence in Mammals": 5_Combining_results.sh

```
#!/bin/bash

# Base directory where species' subdirectories are located
base_dir="Path to parent directory where species' directories exist"

# Loop through all subdirectories (named as Genus_species such as Panthera_leo) in the base directory
for sub_dir in "$base_dir"/*/; do
    # Extract the subdirectory name
    sub_dir_name=$(basename "$sub_dir")

    # Define the two text files
    file1="$sub_dir/6_L1_subset.txt"
    file2="$sub_dir/6_SINE_subset.txt"

    # Define the output file as the subdirectory name (e.g., Panthera_leo.txt)
    output_file="$sub_dir/$sub_dir_name.txt"

    # Remove the pre-existing output file if it exists
    if [[ -f "$output_file" ]]; then
        rm "$output_file"
    fi

    # Get the number of lines for each file (default to 0 if file is empty)
    len1=$(if [[ -s "$file1" ]]; then wc -l < "$file1"; else echo 0; fi)
    len2=$(if [[ -s "$file2" ]]; then wc -l < "$file2"; else echo 0; fi)

    # Get the maximum number of lines across all files
    max_len=$(echo -e "$len1\n$len2" | sort -nr | head -1)

    # Prepare empty columns with zeros for missing or empty files
    col1=$(if [[ -s "$file1" ]]; then cut -f3 "$file1"; else yes 0 | head -n "$max_len"; fi)
    col2=$(if [[ -s "$file2" ]]; then cut -f3 "$file2"; else yes 0 | head -n "$max_len"; fi)

    # Replace backslashes with underscores in the header
    header1="${sub_dir_name}_LINE1"
    header2="${sub_dir_name}_SINE"

    # Write the header to the output file
    echo -e "$header1\t$header2" > "$output_file"

    # Combine the columns and append them to the output file
    paste <(echo "$col1") <(echo "$col2") >> "$output_file"
done
```
