## Supplementary Scripts for "Recent Non-LTR Retrotransposon Activity Predicts Cancer Prevalence in Mammals": 6_intersects_counts_script.sh

```
#!/bin/bash

#SBATCH --job-name=TE-Genes-Intersections
#SBATCH --array=1-55
#SBATCH --cpus-per-task=1
#SBATCH --time=24:00:00
#SBATCH --mail-type=all
#SBATCH --mail-user=email
#SBATCH --output=/ Path to /hcg_Intersect_TE_GENE_%a.out
#SBATCH --error=/Path to/hcg_Intersect_TE_GENE_%a.err
#SBATCH --mem=32000

###################################################################################################################################################################
# "Description:"
#   This script takes the RepeatMasker bed file and a gene annotation bed file and counts, for each nLTR insertion
#   (e.g., L1, SINE), how many gene features it intersects. It is used for genic insertion analyses with
#   either protein coding genes (PC) or cancer gene orthologs (CGOs).
#
#   Requirements:
#     - TE_file: RepeatMasker annotations converted to bed (e.g., Genus_species.bed)
#     - gene_file: bed annotation containing only PC genes or only CGOs (e.g., Genus_species_PCgenes.bed or Genus_species_CGOs.bed)
#     - Both inputs should be coordinate-sorted
#
#   Core operation:
#     - Intersections are computed with: bedtools intersect -a TE_file_asterisks_removed -b "${gene_file}" -sorted -c
#       The -c flag appends a count column giving the number of overlapping gene features per TE record.
#       Records with count > 0 are considered genic insertions for the corresponding gene set.
#
#   Output:
#     - For each species, results are saved to a dedicated subdirectory in a file named as Genus_species.txt
#       (e.g., Ovis_aries.txt)
#
#     - This file has the following structure showing the number of active nLTR Insertions within either PC genes or CGOs
#
#   Notes:
#     - Version differences in awk and bedtools may affect behavior. To reproduce our results exactly, use the same versions as in this study.
#     - Divergence is taken from column 7; element lengths are recalculated as (end – start) and stored in column 10.
#	  - This script is designed to submit as a job array on slurm
###################################################################################################################################################################

export species_name=$(awk "NR==${SLURM_ARRAY_TASK_ID}" species_list.txt)
export TE_file=/"path to bed rmsk2bed outputs"/${species_name}\.bed
export gene_file=/"path to bed annotations"/${species_name}_CGOs.bed
export results=/"Path to the envisioned output directory"


cd ${results} && mkdir -p ${species_name} && cd ${species_name}

## Remove occasional extra columns including asterisks from TE annotation files (some files have 16th column that must be removed)
if awk '{ if (sub(/\*$/, "") && NF == 15) $0 = gensub(/[ \t]+$/, "", "g"); print }' "$TE_file" > TE_file_asterisks_removed; then
    echo "Removed asterisks from TE_file successfully."
else
    echo "Error: Failed to remove asterisks from TE_file."
    exit 1
fi

# Calculate gene-TE intersections
if bedtools intersect -a TE_file_asterisks_removed -b "${gene_file}" -sorted -c > TEs_to_genes_intersects.bed; then
  echo "Counted gene-TE intersections successfully."
else
    echo "Error: Failed to count gene-TE intersections."
    exit 1
fi

# Step to calculate the length of each TE element and replace the 10th column with the calculated length
if awk '{OFS="\t"; $10 = $3 - $2; print}' TEs_to_genes_intersects.bed > 1_modified_gene_te_intersections.txt; then
    echo "Successfully calculated the length of each TE element and updated the 10th column."
else
    echo "Error: Failed to calculate and update the lengths."
    exit 1
fi

# Step to keep only specific entries (L1 and SINEs) based on the 11th column and save the result
if awk '$11 == "LINE/L1" || $11 == "LINE/RTE-BovB" ||  $11 ~ /^SINE\//' 1_modified_gene_te_intersections.txt > 2_modified_gene_te_intersections.txt; then
    echo "Successfully filtered and saved the specified entries."
else
    echo "Error: Failed to filter the entries."
    exit 1
fi

# Step to remove entries where the divergence (7th column) is greater than 5
if awk '$7 <= 5' 2_modified_gene_te_intersections.txt > 3_modified_gene_te_intersections.txt; then
    echo "Successfully filtered entries based on divergence and saved to 3_modified_gene_te_closest_distances.txt."
else
    echo "Error: Failed to filter based on divergence."
    exit 1
fi

# Step to filter SINE subset
if awk '($11 ~ /^SINE\// && $10 >= 100 && $10 <= 400)' 3_modified_gene_te_intersections.txt > All_SINE.txt; then
    echo "Successfully created Alu.txt subset."
else
    echo "Error: Failed to create SINE.txt subset."
    exit 1
fi

# Step to filter L1 subset
if awk '($11 == "LINE/L1" && $10 >= 5450 && $10 <= 6750)' 3_modified_gene_te_intersections.txt > L1.txt; then
    echo "Successfully created L1.txt subset."
else
    echo "Error: Failed to create L1.txt subset."
    exit 1
fi

# Define the output file
output_file="${species_name}.txt"

# Step 1: Create the first row with column headers
echo -e "\tL1\tSINE" > "$output_file"

# Step 2: Calculate the sum of the 16th column for each file and write the second row
l1_count=$(awk '{sum += $16} END {print sum}' L1.txt)
SINE_count=$(awk '{sum += $16} END {print sum}' All_SINE.txt)

# Step 3: Write the species name and the counts to the second row
echo -e "${species_name}\t${l1_count:-0}\t${SINE_count:-0}" >> "$output_file"

# Step 4: Notify successful completion
echo "Successfully created ${species_name}.txt with counts."
```
