## Supplementary Scripts for "Recent Non-LTR Retrotransposon Activity Predicts Cancer Prevalence in Mammals": 7_single_final_intersections_file.sh

```
# Path to the parent directory containing species subdirectories with species result files and named as genus_species
parent_dir="/Path to Parent dir containing subdirs"

# Define the final output file that will store all species counts
final_output="${parent_dir}/all_species_gene_intersections.txt"

# Initialize a flag to handle header inclusion
header_included=false

# Loop through each species directory
for species_dir in "${parent_dir}"/*/; do
    # Extract the species name from the directory path
    species_name=$(basename "${species_dir}")
    
    # Define the species_name.txt file path
    species_file="${species_dir}${species_name}.txt"
    
    # Check if the file exists
    if [[ -f "${species_file}" ]]; then
        # If the header has not been included, add the first line (header) to the final output
        if ! $header_included; then
            head -n 1 "${species_file}" > "$final_output"
            header_included=true
        fi
        
        # Append the second row (counts) to the final output file
        sed -n '2p' "${species_file}" >> "$final_output"
    else
        echo "Warning: ${species_file} not found."
    fi
done

echo "Successfully concatenated all species counts into ${final_output}."
```
