## Supplementary Scripts for "Recent Non-LTR Retrotransposon Activity Predicts Cancer Prevalence in Mammals": 8_commands_to_extract_CGO_data.sh

```
# This script is used to extract the CGOs data from the OrthoFinder output

# Extract cancer gene names from the cancer ortholog peptide file generated with ENSEMBL BioMart
grep -oP '(?<=^>).*' HCG.faa > gene_names.txt

# Run extract_CG_orthogroups_IDs.sh to get cancer gene orthogroup IDs from Orthogroups.tsv. All we need to do is to extract the orthogroups that contain Homo sapiens data.
bash extract_CG_orthogroups_IDs.sh

# Create a subset of Orthogroups.tsv containing cancer gene orthogroups only.  
awk -F'\t' 'NR==FNR {ids[$1]; next} FNR==1 || $1 in ids' HCG_orthogroups_IDs.txt Orthogroups.tsv > HCG_orthogroups.tsv

# Create a subset of the gene count matrix (Orthogroups.GeneCount.tsv) 
# containing only HCG orthogroups, and save it to a new file
head -n 1 Orthogroups.GeneCount.tsv > HCG_Orthogroups.GeneCount.tsv
grep -Ff HCG_orthogroups_IDs.txt Orthogroups.GeneCount.tsv >> HCG_Orthogroups.GeneCount.tsv

# Remove OG0000000 from the gene count matrix. This orthogroup has highly inflated counts, 
# and excluding it is a common practice to avoid bias. One may choose to remove additional 
# inflated orthogroups depending on the analysis.
grep -v "^OG0000000" HCG_orthogroups.tsv > filtered_HCG_orthogroups.tsv
grep -v "^OG0000000" HCG_Orthogroups.GeneCount.tsv > filtered_HCG_Orthogroups.GeneCount.tsv

# Create a mapped_ID file linking each cancer gene to its corresponding orthogroup. 
# The output has two columns: orthogroup ID and gene name
awk -F'\t' 'NR == 1 {for (i = 1; i <= NF; i++) {if ($i == "Orthogroup") col1 = i; if ($i == "Homo_sapiens") col2 = i}} NR > \ 
1 && $col2 != "" {split($col2, genes, ", "); for (j in genes) print $col1 "\t" genes[j]}' filtered_HCG_orthogroups.tsv > mapped_IDs.txt

#sort the mapped file based on the orthogroup names in the first column
sort -k1,1 mapped_IDs.txt -o mapped_IDs.txt

# Remove redundancies 
cut -f1 mapped_IDs.txt | sort | uniq -d
cut -f2 mapped_IDs.txt | sort | uniq -d

# Prepare the input file for counting_orthologs_for_each_gene.py. 
# This file is used to generate the final gene count matrix via the Python script, with cancer gene names as rows 
# and species as columns, where each value represents the count of the corresponding ortholog in the corresponding species.
awk '{print $2, $1}' mapped_IDs.txt | sort | uniq | awk '{gene_to_orthogroups[$1] = gene_to_orthogroups[$1] " " $2} END {for \
(gene in gene_to_orthogroups) print gene, gene_to_orthogroups[gene]}' > gene_to_orthogroups.txt
```
