## Supplementary Scripts for "Recent Non-LTR Retrotransposon Activity Predicts Cancer Prevalence in Mammals": 10_counting_orthologs_for_each_gene.py

```
import pandas as pd

# Load orthogroup gene counts
gene_counts_df = pd.read_csv('filtered_HCG_Orthogroups.GeneCount.tsv', sep='\t', index_col=0)

# Load gene to orthogroups mapping
gene_to_orthogroups = {}
with open('gene_to_orthogroups.txt') as f:
    for line in f:
        gene, *orthogroups = line.strip().split()
        gene_to_orthogroups[gene] = orthogroups

# Initialize an empty DataFrame to store summed counts per gene
species_list = gene_counts_df.columns.tolist()
result_df = pd.DataFrame(0, index=species_list, columns=gene_to_orthogroups.keys())

# Sum counts for each gene across its orthogroups
for gene, orthogroups in gene_to_orthogroups.items():
    # Filter orthogroups to only those present in gene_counts_df
    valid_orthogroups = [og for og in orthogroups if og in gene_counts_df.index]
    if valid_orthogroups:
        # Sum across valid orthogroups for this gene
        gene_counts = gene_counts_df.loc[valid_orthogroups].sum()
        # Assign summed counts to the result DataFrame
        result_df[gene] = gene_counts
    else:
        print(f"No valid orthogroups found in counts file for gene {gene}")

# Transpose the DataFrame to match desired format (species as rows, genes as columns)
result_df = result_df.T
result_df.to_csv('HCG_counts.txt', sep='\t')
```
