## Supplementary Scripts for "Recent Non-LTR Retrotransposon Activity Predicts Cancer Prevalence in Mammals": 11_make_species_orthologs_list.sh

```
awk -F'\t' '
NR == 1 {
    # Save the headers (species names)
    for (i = 2; i <= NF; i++) species[i] = $i
}
NR > 1 {
    # For each species (columns starting from 2)
    for (i = 2; i <= NF; i++) {
        if ($i != "") {
            # Split the cell contents into individual orthologues
            split($i, orthologues, ", ")
            for (j in orthologues) {
                # Append each orthologue to the species-specific array
                species_orthologues[i] = (species_orthologues[i] ? species_orthologues[i] "\n" : "") orthologues[j]
            }
        }
    }
}
END {
    # Print species names as header
    for (i = 2; i <= NF; i++) {
        printf "%s%s", species[i], (i == NF ? "\n" : "\t")
    }
    # Print orthologues for each species column
    max_lines = 0
    for (i = 2; i <= NF; i++) {
        split(species_orthologues[i], lines, "\n")
        for (j in lines) species_orthologues_arr[i][j] = lines[j]
        max_lines = (length(lines) > max_lines ? length(lines) : max_lines)
    }
    for (row = 1; row <= max_lines; row++) {
        for (i = 2; i <= NF; i++) {
            printf "%s%s", (species_orthologues_arr[i][row] ? species_orthologues_arr[i][row] : ""), (i == NF ? "\n" : "\t")
        }
    }
}
' filtered_HCG_orthogroups.tsv > species_orthologs.txt
```
