## Supplementary Scripts for "Recent Non-LTR Retrotransposon Activity Predicts Cancer Prevalence in Mammals": 12_making_HCG_gff.sh

```
#!/bin/bash

#SBATCH --job-name=CGO_gff
#SBATCH --array=1-55
#SBATCH --cpus-per-task=1
#SBATCH --time=06:00:00
#SBATCH --mail-type=END
#SBATCH --mail-user=email
#SBATCH --output=/path to/CGO_gff_%A_%a.out
#SBATCH --error=/path to/CGO_gff_%A_%a.err
#SBATCH --mem=16000

# Read species from the list based on array task ID
species=$(awk "NR==${SLURM_ARRAY_TASK_ID}" species_list.txt)

PARENT_DIR="/Path to the parent directory containing speceis subdirectories, which contain main gff files"
species_dir="$PARENT_DIR/$species"
main_gff="$species_dir/${species}.gff"
output_gff="$species_dir/${species}_HCG_orthologs.gff"
not_found_file="$species_dir/not_found_orthologs.txt"
species_orthologs="species_orthologs.txt"

# Extract the column number for the species
col_num=$(head -n 1 "$species_orthologs" | tr '\t' '\n' | nl | awk -v species="$species" '$2 == species {print $1}')

if [ -z "$col_num" ]; then
    echo "Species $species not found in $species_orthologs"
    exit 1
fi

# Clear previous output files if they exist
> "$output_gff"
> "$not_found_file"

# Extract gene list for the species
gene_list=$(awk -v col="$col_num" 'NR > 1 {print $col}' "$species_orthologs" | grep -v "^$")

# Loop through each gene in the list
while IFS= read -r gene; do
    gene=$(echo $gene | tr -d '\r')  # Trim carriage returns
    gene_found=false

    # Find all entries related to the exact gene using regex
    gene_entries=$(awk -v gene="$gene" 'BEGIN {IGNORECASE=1} $9 ~ "(^|;)ID=(gene-|exon-|cds-)?" gene "(;|$)"' "$main_gff")

    if [ -n "$gene_entries" ]; then
        # Check for gene entries first
        echo "$gene_entries" | awk '$3 == "gene"' >> "$output_gff"
        if echo "$gene_entries" | awk '$3 == "gene"' | grep -q .; then
            gene_found=true
        fi

        # If no gene entry, check for the longest CDS
        if [ "$gene_found" = false ]; then
            longest_cds=$(echo "$gene_entries" | awk '$3 == "CDS" {print $5-$4+1, $0}' | sort -nr | head -n1 | cut -d' ' -f2-)
            if [ -n "$longest_cds" ]; then
                echo "$longest_cds" >> "$output_gff"
                gene_found=true
            fi
        fi

        # If no CDS, create a gene entry from exon coordinates
        if [ "$gene_found" = false ]; then
            exon_entries=$(echo "$gene_entries" | awk '$3 == "exon"')
            if [ -n "$exon_entries" ]; then
                start=$(echo "$exon_entries" | awk '{print $4}' | sort -n | head -n1)
                end=$(echo "$exon_entries" | awk '{print $5}' | sort -nr | head -n1)
                exemplar=$(echo "$exon_entries" | head -n1)
                echo "$exemplar" | awk -v start="$start" -v end="$end" 'BEGIN {OFS="\t"} {$3="gene"; $4=start; $5=end; print $0}' >> "$output_gff"
                gene_found=true
            fi
        fi
    fi

    # If no entry found at all, log the gene
    if [ "$gene_found" = false ]; then
        echo "$gene" >> "$not_found_file"
    fi

done <<< "$gene_list"
```
