## Supplementary Scripts for "Recent Non-LTR Retrotransposon Activity Predicts Cancer Prevalence in Mammals": 13_extract_PC_genes.sh

```
#!/bin/bash

# Parent directory containing subdirectories for each species
parent_directory="/Path to speceis subdirectories"

# Loop through species subdirectories named as Genus_species including complete gff files named as Genus_species.gff
for species_directory in "$parent_directory"/*; do
    species_name=$(basename "$species_directory")
    gff_file="$species_directory/$species_name.gff"
    output_file="$species_directory/${species_name}_protein_coding_genes.gff"

    if [ -f "$gff_file" ]; then
        # Extract protein-coding genes
        awk -F '\t' '$3 == "gene" && $9 ~ /gene_biotype=protein_coding/' "$gff_file" > "$output_file"
    fi
done

echo "Protein-coding genes have been extracted."
```
