## Supplementary Scripts for "Recent Non-LTR Retrotransposon Activity Predicts Cancer Prevalence in Mammals": 14_calculate_means_medians.R

```
###################################################################################################
########################################## Calculate IQR means and medians
#########################
# This script requires the path to a directory containing all Genus_species.txt files, 
# each representing proximity to either PC genes or CGOs
####
rm(list=ls(all=TRUE))
ls()

#####################################################################
# Required libraries
library(dplyr)

# Function to calculate the trimmed mean based on IQR with missing values removed
trimmed_mean_iqr <- function(column_data) {
  # Remove NA values
  column_data <- na.omit(column_data)
  
  # Check if the column is entirely zeros or empty
  if (all(column_data == 0) || length(column_data) == 0) {
    return(0)  # Return zero if all values are zero or column is empty
  }
  
  # Calculate the 1st and 3rd quartiles
  q1 <- quantile(column_data, 0.25, na.rm = TRUE)
  q3 <- quantile(column_data, 0.75, na.rm = TRUE)
  
  # Calculate IQR
  iqr <- q3 - q1
  
  # Define bounds for outlier removal
  lower_bound <- q1 - 1.5 * iqr
  upper_bound <- q3 + 1.5 * iqr
  
  # Remove outliers
  trimmed_data <- column_data[column_data >= lower_bound & column_data <= upper_bound]
  
  # Calculate and return the trimmed mean, rounded to 2 decimal places
  round(mean(trimmed_data, na.rm = TRUE), 2)
}

# Function to calculate the median, handling zeros or empty columns
calculate_median <- function(column_data) {
  # Remove NA values
  column_data <- na.omit(column_data)
  
  # Check if the column is entirely zeros or empty
  if (all(column_data == 0) || length(column_data) == 0) {
    return(0)  # Return zero if all values are zero or column is empty
  }
  
  # Calculate and return the median, rounded to 2 decimal places
  round(median(column_data, na.rm = TRUE), 2)
}

# Directory containing the txt files
file_directory <- "Path to the .txt files directory"

# Get the list of text files
file_list <- list.files(file_directory, pattern = "*.txt", full.names = TRUE)

# Initialize the final dataframe to store the results
combined_stats <- data.frame(
  species = character(),
  IQR_average = numeric(),
  median_all_elements = numeric(),
  stringsAsFactors = FALSE
)

# Loop through each file
for (file in file_list) {
  # Read the file into a dataframe
  df <- read.table(file, header = TRUE, sep = "\t")
  
  # Get the species name by removing the file extension
  species_name <- sub(".txt", "", basename(file))
  
  # Combine all columns into one vector
  all_elements <- unlist(df)
  all_elements <- all_elements[all_elements != 0]
  
  # Calculate IQR average and median for all elements combined
  iqr_avg <- trimmed_mean_iqr(all_elements)
  med_all <- calculate_median(all_elements)
  
  # Add the results to the combined_stats dataframe
  combined_stats <- rbind(combined_stats, data.frame(
    species = species_name,
    IQR_average = iqr_avg,
    median_all_elements = med_all,
    stringsAsFactors = FALSE
  ))
}

# Print the final dataframe
print(combined_stats)

library(writexl)
write_xlsx(combined_stats, "All_Elements_IQR_means_medians.xlsx")
```
