## Supplementary Scripts for "Recent Non-LTR Retrotransposon Activity Predicts Cancer Prevalence in Mammals": 18_CGO_data_processing.R

```
#######################################################################################
##################################################### 
# This script merge the 15_HCG_counts.csv generated in slurm with the cancer
# gene functional status in 16_CGO_Status.xlsx first, then process the  
# data and merge it to the final Supplementary Data S1 Sheet
#################################
rm(list=ls(all=TRUE))
ls()

#############################
# Load necessary libraries
library(readxl)
library(writexl)
library(dplyr)
library(tidyr)

#############################
# Loading counts data
# Set file paths (adjust to your actual paths)
file1 <- "C:Path to /15_CGO_counts.xlsx"
hcg_counts <- read_excel(file1)
head(hcg_counts)

##############################
# Loading cancer data  
file2 <- "Path to /16_CGO_Status.xlsx"
hcg_status <- read_excel(file2)
head(hcg_status)
nrow(hcg_status)

##################Step one > sorting counts file based on gene names
# Sort hcg_counts by the Gene Symbol (first column)
hcg_counts <- hcg_counts[order(hcg_counts[[1]]), ]

# View the first few rows to confirm sorting
head(hcg_counts)

################## Step two > filter hcg_status for those genes that exist in hcg_counts Gene_Name column
# Filter hcg_status to keep only genes present in hcg_counts
hcg_status_temp <- hcg_status %>%
  filter(`Gene Symbol` %in% hcg_counts$Gene_Name)

# View the first few rows to confirm
head(hcg_status_temp)

################## Step three > combining two files
nrow(hcg_status_temp)
nrow(hcg_counts) 

# Find genes in hcg_counts that are not in hcg_status
missing_genes <- setdiff(hcg_counts$Gene_Name, hcg_status$`Gene Symbol`)
# Display missing genes
print(missing_genes)

# Check for duplicates in hcg_counts
duplicates <- hcg_counts %>%
  group_by(Gene_Name) %>%
  filter(n() > 1)

# Display the duplicates
print(duplicates)

############### Let's see if gene names in their respective columns are matched  
# Check if gene names match row by row
matching <- all(hcg_counts$Gene_Name == hcg_status_temp$`Gene Symbol`)

# Print the result
if (matching) {
  print("Gene names match perfectly!")
} else {
  print("Mismatch detected!")
  mismatches <- hcg_counts$Gene_Name[hcg_counts$Gene_Name != hcg_status_temp$`Gene Symbol`]
  print(mismatches)
}

nrow(hcg_status_temp)
nrow(hcg_counts)

############### Now cbind them
# Combine the data frames directly
final_hcg_counts_status <- cbind(hcg_status_temp, hcg_counts)

# Save the final data frame as a record
write_xlsx(final_hcg_counts_status, "Path to save /17_CGO_counts_status.xlsx")

# Confirm the first few rows
head(final_hcg_counts_status)
tail(final_hcg_counts_status)

##################### calculating ortholog numbers in each functional category  
# Extract species names of the hcg_counts
species <- colnames(hcg_counts)[-1]
species <- species[species != "Homo_sapiens"]
length(species)
CGO_total_counts <- data.frame(species = species)

################################################# Counting Somatics
# Subset rows where Somatic == "yes"
temp1 <- final_hcg_counts_status[final_hcg_counts_status$Somatic == "yes", ]
head(temp1)
#View(temp1)
# Set k as the index after which to start summing (e.g., column 6)
k <- 11  
# Sum values across all columns after column k
temp2 <- colSums(temp1[, (k+1):ncol(temp1)], na.rm = TRUE)
length(temp2)

# Removing the record for Homo_sapiens
temp2 <- temp2[!names(temp2) %in% c("Homo_sapiens", "Total", "Average")]
length(temp2)

temp2_df <- data.frame(
  Species = names(temp2),
  Value = as.numeric(temp2),
  row.names = NULL
)

CGO_total_counts$Somatic <- as.numeric(temp2)
#View(CGO_total_counts)

################################################# Counting Germlines
# Subset rows where Germline == "yes"
temp1 <- final_hcg_counts_status[final_hcg_counts_status$Germline == "yes", ]
head(temp1)
#View(temp1)
# Set k as the index after which to start summing (e.g., column 6)
k <- 11  
# Sum values across all columns after column k
temp2 <- colSums(temp1[, (k+1):ncol(temp1)], na.rm = TRUE)
length(temp2)

temp2 <- temp2[!names(temp2) %in% c("Homo_sapiens", "Total", "Average")]
length(temp2)

temp2_df <- data.frame(
  Species = names(temp2),
  Value = as.numeric(temp2),
  row.names = NULL
)

CGO_total_counts$Germline <- as.numeric(temp2)
#View(CGO_total_counts)

################################################# Counting Oncogenes
# Subset rows where Oncogene == "yes"
temp1 <- final_hcg_counts_status[final_hcg_counts_status$Oncogene == "yes", ]
head(temp1)
#View(temp1)
# Set k as the index after which to start summing (e.g., column 6)
k <- 11  
# Sum values across all columns after column k
temp2 <- colSums(temp1[, (k+1):ncol(temp1)], na.rm = TRUE)
length(temp2)

temp2 <- temp2[!names(temp2) %in% c("Homo_sapiens", "Total", "Average")]
length(temp2)

temp2_df <- data.frame(
  Species = names(temp2),
  Value = as.numeric(temp2),
  row.names = NULL
)

CGO_total_counts$Oncogene <- as.numeric(temp2)
#View(CGO_total_counts)

################################################# Counting TSGs
# Subset rows where TSG == "yes"
temp1 <- final_hcg_counts_status[final_hcg_counts_status$TSG == "yes", ]
head(temp1)
#View(temp1)
# Set k as the index after which to start summing (e.g., column 6)
k <- 11  
# Sum values across all columns after column k
temp2 <- colSums(temp1[, (k+1):ncol(temp1)], na.rm = TRUE)
length(temp2)

temp2 <- temp2[!names(temp2) %in% c("Homo_sapiens", "Total", "Average")]
length(temp2)

temp2_df <- data.frame(
  Species = names(temp2),
  Value = as.numeric(temp2),
  row.names = NULL
)

CGO_total_counts$TSG <- as.numeric(temp2)
#View(CGO_total_counts)

################################################# Counting Fusions
# Subset rows where Fusion == "yes"
temp1 <- final_hcg_counts_status[final_hcg_counts_status$Fusion == "yes", ]
head(temp1)
#View(temp1)
# Set k as the index after which to start summing (e.g., column 6)
k <- 11  
# Sum values across all columns after column k
temp2 <- colSums(temp1[, (k+1):ncol(temp1)], na.rm = TRUE)
length(temp2)

temp2 <- temp2[!names(temp2) %in% c("Homo_sapiens", "Total", "Average")]
length(temp2)

temp2_df <- data.frame(
  Species = names(temp2),
  Value = as.numeric(temp2),
  row.names = NULL
)

CGO_total_counts$Fusion <- as.numeric(temp2)
#View(CGO_total_counts)

################################### Summin up the total number of CGOs in each species
temp3 <- final_hcg_counts_status
# Set k as the index after which to start summing (e.g., column 6)
k <- 11  
# Sum values across all columns after column k
temp4 <- colSums(temp3[, (k+1):ncol(temp3)], na.rm = TRUE)
length(temp4)

temp4 <- temp4[!names(temp4) %in% c("Homo_sapiens", "Total", "Average")]
length(temp4)

temp4_df <- data.frame(
  Species = names(temp4),
  Value = as.numeric(temp4),
  row.names = NULL
)

CGO_total_counts$Total_HCGHs <- as.numeric(temp4)
#View(CGO_total_counts)

################################## Saving the final data sheet
# Save the final dataframe to CGO_total_counts. Information in this file can be directly added to the Supplementary Data S1
write_xlsx(CGO_total_counts, "Path to /CGO_total_counts.xlsx")
write.csv(CGO_total_counts, "Path to /CGO_total_counts.csv", row.names = FALSE)
```
