## Supplementary Scripts for "Recent Non-LTR Retrotransposon Activity Predicts Cancer Prevalence in Mammals": 0_pglsSEyPagel.R

```
suppressPackageStartupMessages({
library(nlme)
library(phytools)
library(tidyverse)
library(geiger)
library(ape)
library(caper)
library(rms)
library(stringr)
})

#make sure to run all of this before you get to work.
#pgls sey base (just run all of this)
modPgls.SEy = function (model, data, corClass = corBrownian, tree, se = NULL, 
                        method = c("REML", "ML"), interval = c(0, 1000), corClassValue=1, sig2e=NULL, ...) 
{
  Call <- match.call()
  corfunc <- corClass
  spp <- rownames(data)
  data <- cbind(data, spp)
  if (is.null(se)) 
    se <- setNames(rep(0, Ntip(tree)), tree$tip.label)[spp]
  else se <- se[spp]
  
  lk <- function(sig2e, data, tree, model, ve, corfunc, spp) {
    tree$edge.length <- tree$edge.length * sig2e
    ii <- sapply(1:Ntip(tree), function(x, e) which(e == 
                                                      x), e = tree$edge[, 2])
    tree$edge.length[ii] <- tree$edge.length[ii] + ve[tree$tip.label]
    vf <- diag(vcv(tree))[spp]
    w <- varFixed(~vf)
    COR <- corfunc(corClassValue, tree, form = ~spp, ...)
    fit <- gls(model, data = cbind(data, vf), correlation = COR, 
               method = method, weights = w)
    -logLik(fit)
  }
  
  if (is.null(sig2e)) {
    fit <- optimize(lk, interval = interval, data = data, tree = tree, 
                    model = model, ve = se^2, corfunc = corfunc, spp = spp)
    sig2e=fit$minimum
  }
  
  tree$edge.length <- tree$edge.length * sig2e
  ii <- sapply(1:Ntip(tree), function(x, e) which(e == x), 
               e = tree$edge[, 2])
  tree$edge.length[ii] <- tree$edge.length[ii] + se[tree$tip.label]^2
  vf <- diag(vcv(tree))[spp]
  w <- varFixed(~vf)
  obj <- gls(model, data = cbind(data, vf), correlation = corfunc(corClassValue, 
                                                                  tree, form = ~spp, ...), weights = w, method = method)
  obj$call <- Call
  obj$sig2e <- sig2e
  obj
}

#Internal function
pglsSEyPagelToOptimizeLambda=function(lambda,model,data,tree,...) {
  -logLik(modPgls.SEy(model=model,data=data,tree=tree,corClassValue=lambda,corClass=corPagel,fixed=T,...)) #Returns -logLikelihood of the pgls.SEy model with lambda fixed to the value of the lambda argument. sig2e will be optimized within modPgls.SEy unless given as an argument here
}

#Function intended for users
pglsSEyPagel=function(model, data, tree, lambdaInterval=c(0,1),...){
  optimizedModel=optimize(pglsSEyPagelToOptimizeLambda,lambdaInterval,model=model,data=data,tree=tree,...) #Optimizes lambda in the lambdaInterval using the pglsSEyPagelToOptimizeLambda function
  return(modPgls.SEy(model=model,data=data,tree=tree,corClass=corPagel,fixed=T,corClassValue=optimizedModel$minimum,...)) #Returns the final model fit
}
```
