## Supplementary Scripts for "Recent Non-LTR Retrotransposon Activity Predicts Cancer Prevalence in Mammals": 01_nLTR_Abundance.R

```
###################################################################################################
####################################### The nLTR Abundance Analyses
#################
rm(list=ls(all=TRUE))
ls()

#####################################################################
suppressPackageStartupMessages({
  library(ggplot2)
  library(reshape2)
  library(plyr)
  library(tidyverse)
  library(dplyr)
  library(rcompanion)# Tukey Transformation
  library(car) # Durbin Watson Test
  library(phytools) # PGLS 
  library(olsrr) # Assess the normality of residuals
  library(MASS) # Transformation
  library(lmtest) # Breusch_Pagan test
  library(outliers) # To identify outliers (Grub's test)
  library(ggimage) # Plotting
  library(magick) # Plotting
  library(ggtree)
  library(cowplot) # For Plotting
  library(ggtext) # For legends in plots
  library(ggrepel) # To avoid overlaps between labels in plots
  library(viridis) # Color blind palette for plots
  library(caper) # For reading tree
  library(rr2) # For reading tree / Likelihood-based R2
  library(devtools) # For RPICR
  #library(ROBRT) # For RPICR > You need to use install_github from devtools
  library(geiger) # for RPICR
})
#####################################################################
########## Load custom PGLS functions into the current session
source("Path to /0_pglsSEyPagel.R")
#####################################################################
setwd("Path to the working directory")
getwd()

# Reading Excel data 
mydata <- read.csv("Path to the Supplementary Data S1/Supplementary Data S1.csv", header = TRUE)

#####################################################################
########## Load the ultrametric phylogenetic tree
tre <- read.tree("Mammals_tree.newick")

########## See the orders and superorders
#species_count <- as.data.frame(table(mydata$Order))
#colnames(species_count) <- c("Order","Count")  # Rename the columns
#print(species_count)
#View(mydata[mydata$Order == "Diprotodontia", ])

#species_count <- as.data.frame(table(mydata$SuperOrder))
#colnames(species_count) <- c("SuperOrder","Count")  # Rename the columns
#print(species_count)
#View(mydata[mydata$Order == "Diprotodontia", ])

#####################################################################
# Data preparation
#####################################################################
rownames(mydata) <- mydata$species

########## Setting SE
SE <- setNames(mydata$SE, mydata$species)[rownames(mydata)]
nrow(mydata)

########## Check for normality in cancer data
hist(mydata$NeoplasiaPrevalence)
hist(mydata$MalignancyPrevalence)

########## Transform data to mitigate skewness and make the data follow a normal distribution
### Tukey's Ladder of Powers transform 
mydata$NeoplasiaPrevalence <- transformTukey(mydata$NeoplasiaPrevalence)
mydata$MalignancyPrevalence <- transformTukey(mydata$MalignancyPrevalence)
hist(mydata$NeoplasiaPrevalence)
hist(mydata$MalignancyPrevalence)

########## Check for the outliers 
grubbs_result <- grubbs.test(mydata$NeoplasiaPrevalence)
print(grubbs_result)
grubbs_result <- grubbs.test(mydata$MalignancyPrevalence)
print(grubbs_result)

########## Define predictor variables (L1 counts and combined L1 + SINE counts)
mydata$Transformed_L1_Counts <- mydata$L1_Counts
mydata$Transformed_L1_SINEs_Counts <- mydata$L1_SINEs_Counts

########## Transform nLTRs data to mitigate skewness and make the data follow a normal distribution
hist(mydata$Transformed_L1_Counts)
mydata$Transformed_L1_Counts <- transformTukey(mydata$Transformed_L1_Counts)
# scale data to the 0,1 range
mydata$Transformed_L1_Counts <- (mydata$Transformed_L1_Counts - min(mydata$Transformed_L1_Counts)) / (max(mydata$Transformed_L1_Counts) - min(mydata$Transformed_L1_Counts))
hist(mydata$Transformed_L1_Counts)


hist(mydata$Transformed_L1_SINEs_Counts)
mydata$Transformed_L1_SINEs_Counts <- transformTukey(mydata$Transformed_L1_SINEs_Counts)
# scale data to the 0,1 range
mydata$Transformed_L1_SINEs_Counts <- (mydata$Transformed_L1_SINEs_Counts - min(mydata$Transformed_L1_SINEs_Counts)) / (max(mydata$Transformed_L1_SINEs_Counts) - min(mydata$Transformed_L1_SINEs_Counts))
hist(mydata$Transformed_L1_SINEs_Counts)


########### Check for the potential outliers in nLTR counts
grubbs_result <- grubbs.test(mydata$Transformed_L1_Counts)
print(grubbs_result)
grubbs_result <- grubbs.test(mydata$Transformed_L1_SINEs_Counts)
print(grubbs_result)

########## Pearson Correlations Between Neoplasia/Malignancy and NLTR Measurements
corrNeo <- round(cor(mydata$NeoplasiaPrevalence, mydata$Transformed_L1_Counts),2)
corrNeo
corrNeo2 <- round(cor(mydata$NeoplasiaPrevalence, mydata$Transformed_L1_SINEs_Counts),2)
corrNeo2

corrMal <- round(cor(mydata$MalignancyPrevalence, mydata$Transformed_L1_Counts),2)
corrMal
corrMal2 <- round(cor(mydata$MalignancyPrevalence, mydata$Transformed_L1_SINEs_Counts),2)
corrMal2

#######################################################################
### Run PGLS
#######################################################################

########### Simple PGLS (Cancer ~ (Number of L1 OR All nLTRs) + e)

### Neoplasia vs L1
NL1 <- pglsSEyPagel(NeoplasiaPrevalence ~ Transformed_L1_Counts, data = mydata, tree=tre, se=SE, method = "ML")
summary(NL1)
anova(NL1)
# Grab likelihood-based R2, Lambda, and p values for plotting
RelNL1 <- R2_lik(NL1)
RelNL1 <- format(RelNL1)
RelNL1 <- signif(as.numeric(RelNL1), digits = 2)
RelNL1
lambdaNL1 <- summary(NL1)$modelStruct$corStruct
lambdaNL1 <- signif(lambdaNL1[1], digits = 2)
lambdaNL1
NL1_pvalue <- summary(NL1)$tTable
NL1_pvalue <- signif(NL1_pvalue[2,4], digits = 2)
NL1_pvalue
# Check for normality of residuals 
ols_test_normality(NL1$residuals)
hist(NL1$residuals, main = "NL1 Residual Histogram")
qqnorm(NL1$residuals)
qqline(NL1$residuals)
# check for independence of errors (autocorrelation) 
# Autocorrelation occurs when the residuals of a regression model are correlated
# with each other, indicating that there is some pattern or dependence among the errors.
model <- lm(NeoplasiaPrevalence ~ Transformed_L1_Counts , data=mydata)
durbinWatsonTest(model) #Durbin-Watson test using car package
# Check for heteroscedasticity (Breusch-Pagan test)
# Checks if the variance of the errors is the same for every value of x (homoscedasticity). 
# Look for a random scatter of points around zero, with no discernible pattern or trend. 
# If the spread of residuals appears to vary systematically as the predicted
# values or predictor variable(s) change, it suggests potential heteroscedasticity.
plot(NL1, which = 1)  # Residuals vs. Fitted value plot
plot(NL1, which = 3)  # Scale-Location plot (square root of standardized residuals vs. Fitted values)
bptest(lm(NL1$residuals ~ fitted(NL1)))
### AIC Model Comparison
# Load the data and prepare comparative data object for AIC comparisons
comparative_data <- comparative.data(phy = tre, data = mydata, 
                                     names.col = "species", 
                                     vcv = TRUE, na.omit = FALSE)
# Lambda fixed at 1 (full phylogenetic signal)
model_lambda_1 <- pgls(NeoplasiaPrevalence ~ Transformed_L1_Counts, data = comparative_data, lambda = 1.00)
# Compare AIC values
cat("AIC with Lambda = 0:", round(AIC(model),2), "\n")
cat("AIC with Estimated Lambda:", round(AIC(NL1),2), "\n")
cat("AIC with Lambda = 1:", round(AIC(model_lambda_1),2), "\n")


#### Neoplasia vs (L1 + SINES)
NL1SINEs <- pglsSEyPagel(NeoplasiaPrevalence ~ Transformed_L1_SINEs_Counts, data = mydata, tree=tre, se=SE, method = "ML")
summary(NL1SINEs)
anova(NL1SINEs)
# Grab likelihood-based R2, Lambda, and p values for plotting
RelNL1SINEs <- R2_lik(NL1SINEs)
RelNL1SINEs <- format(RelNL1SINEs)
RelNL1SINEs <- signif(as.numeric(RelNL1SINEs), digits= 2)
RelNL1SINEs
lambdaNL1SINEs <- summary(NL1SINEs)$modelStruct$corStruct
lambdaNL1SINEs <- signif(lambdaNL1SINEs[1], digits = 2)
lambdaNL1SINEs
NL1SINEs_pvalue <- summary(NL1SINEs)$tTable
NL1SINEs_pvalue <- signif(NL1SINEs_pvalue[2,4], digits = 2)
NL1SINEs_pvalue
# Check for normality of residuals 
ols_test_normality(NL1SINEs$residuals)
hist(NL1SINEs$residuals, main = "NL1SINEs Residual Histogram")
qqnorm(NL1SINEs$residuals)
qqline(NL1SINEs$residuals)
# Check for independence of errors (autocorrelation) 
# Autocorrelation occurs when the residuals of a regression model are correlated
# with each other, indicating that there is some pattern or dependence among the errors.
model <- lm(NeoplasiaPrevalence ~ Transformed_L1_SINEs_Counts , data=mydata)
durbinWatsonTest(model) #Durbin-Watson test using car package
# Check for heteroscedasticity (Breusch-Pagan test)
# Checks if the variance of the errors is the same for every value of x (homoscedasticity). 
# Look for a random scatter of points around zero, with no discernible pattern or trend. 
# If the spread of residuals appears to vary systematically as the predicted
# values or predictor variable(s) change, it suggests potential heteroscedasticity.
plot(NL1SINEs, which = 1)  # Residuals vs. Fitted value plot
plot(NL1SINEs, which = 3)  # Scale-Location plot (square root of standardized residuals vs. Fitted values)
bptest(lm(NL1SINEs$residuals ~ fitted(NL1SINEs)))
### AIC Model Comparison
# Load the data and prepare comparative data object for AIC comparisons
# comparative_data <- comparative.data(phy = tre, data = mydata, 
#                                      names.col = "species", 
#                                      vcv = TRUE, na.omit = FALSE)
# Lambda fixed at 1 (full phylogenetic signal)
model_lambda_1 <- pgls(NeoplasiaPrevalence ~ Transformed_L1_Counts, data = comparative_data, lambda = 1.00)
# Compare AIC values
cat("AIC with Lambda = 0:", round(AIC(model),2), "\n")
cat("AIC with Estimated Lambda:", round(AIC(NL1SINEs),2), "\n")
cat("AIC with Lambda = 1:", round(AIC(model_lambda_1),2), "\n")

#### Malignancy vs L1
ML1 <- pglsSEyPagel(MalignancyPrevalence ~ Transformed_L1_Counts, data = mydata, tree=tre, se=SE, method = "ML")
summary(ML1)
anova(ML1)
# Grab likelihood-based R2, Lambda, and p values for plotting
RelML1 <- R2_lik(ML1)
RelML1 <- format(RelML1)
RelML1 <- signif(as.numeric(RelML1), digits= 2)
RelML1
lambdaML1 <- summary(ML1)$modelStruct$corStruct
lambdaML1 <- signif(lambdaML1[1], digits = 2)
lambdaML1
ML1_pvalue <- summary(ML1)$tTable
ML1_pvalue <- signif(ML1_pvalue[2,4], digits = 2)
ML1_pvalue
# Check for normality of residuals 
ols_test_normality(ML1$residuals)
hist(ML1$residuals, main = "ML1 Residual Histogram")
qqnorm(ML1$residuals)
qqline(ML1$residuals)
# Check for independence of errors (autocorrelation) 
# Autocorrelation occurs when the residuals of a regression model are correlated
# with each other, indicating that there is some pattern or dependence among the errors.
model <- lm(MalignancyPrevalence ~ Transformed_L1_Counts , data=mydata)
durbinWatsonTest(model) #Durbin-Watson test using car package
# Check for heteroscedasticity (Breusch-Pagan test)
# Checks if the variance of the errors is the same for every value of x (homoscedasticity). 
# Look for a random scatter of points around zero, with no discernible pattern or trend. 
# If the spread of residuals appears to vary systematically as the predicted
# values or predictor variable(s) change, it suggests potential heteroscedasticity.
plot(ML1, which = 1)  # Residuals vs. Fitted value plot
plot(ML1, which = 3)  # Scale-Location plot (square root of standardized residuals vs. Fitted values)
bptest(lm(ML1$residuals ~ fitted(ML1)))

### AIC Model Comparison
# Load the data and prepare comparative data object for AIC comparisons
# comparative_data <- comparative.data(phy = tre, data = mydata, 
#                                      names.col = "species", 
#                                      vcv = TRUE, na.omit = FALSE)
# Lambda fixed at 1 (full phylogenetic signal)
model_lambda_1 <- pgls(MalignancyPrevalence ~ Transformed_L1_Counts, data = comparative_data, lambda = 1.00)
# Compare AIC values
cat("AIC with Lambda = 0:", round(AIC(model),2), "\n")
cat("AIC with Estimated Lambda:", round(AIC(ML1),2), "\n")
cat("AIC with Lambda = 1:", round(AIC(model_lambda_1),2), "\n")

#### Malignancy vs (L1 + SINES)
ML1SINEs <- pglsSEyPagel(MalignancyPrevalence ~ Transformed_L1_SINEs_Counts, data = mydata, tree=tre, se=SE, method = "ML")
summary(ML1SINEs)
anova(ML1SINEs)
# Grab likelihood-based R2, Lambda, and p values for plotting
RelML1SINEs <- R2_lik(ML1SINEs)
RelML1SINEs <- format(RelML1SINEs)
RelML1SINEs <- signif(as.numeric(RelML1SINEs), digits= 2)
RelML1SINEs
lambdaML1SINEs <- summary(ML1SINEs)$modelStruct$corStruct
lambdaML1SINEs <- signif(lambdaML1SINEs[1], digits = 2)
lambdaML1SINEs
ML1SINEs_pvalue <- summary(ML1SINEs)$tTable
ML1SINEs_pvalue <- signif(ML1SINEs_pvalue[2,4], digits = 2)
ML1SINEs_pvalue
# Check for normality of residuals 
ols_test_normality(ML1SINEs$residuals)
hist(ML1SINEs$residuals, main = "ML1SINEs Residual Histogram")
qqnorm(ML1SINEs$residuals)
qqline(ML1SINEs$residuals)
# Check for independence of errors (autocorrelation) 
# Autocorrelation occurs when the residuals of a regression model are correlated
# with each other, indicating that there is some pattern or dependence among the errors.
model <- lm(MalignancyPrevalence ~ Transformed_L1_SINEs_Counts , data=mydata)
durbinWatsonTest(model) #Durbin-Watson test using car package
# Check for heteroscedasticity (Breusch-Pagan test)
# Checks if the variance of the errors is the same for every value of x (homoscedasticity). 
# Look for a random scatter of points around zero, with no discernible pattern or trend. 
# If the spread of residuals appears to vary systematically as the predicted
# values or predictor variable(s) change, it suggests potential heteroscedasticity.
plot(ML1SINEs, which = 1)  # Residuals vs. Fitted value plot
plot(ML1SINEs, which = 3)  # Scale-Location plot (square root of standardized residuals vs. Fitted values)
bptest(lm(ML1SINEs$residuals ~ fitted(ML1SINEs)))

### AIC Model Comparison
# Load the data and prepare comparative data object for AIC comparisons
# comparative_data <- comparative.data(phy = tre, data = mydata, 
#                                      names.col = "species", 
#                                      vcv = TRUE, na.omit = FALSE)
# Lambda fixed at 1 (full phylogenetic signal)
model_lambda_1 <- pgls(MalignancyPrevalence ~ Transformed_L1_SINEs_Counts, data = comparative_data, lambda = 1.00)
# Compare AIC values
cat("AIC with Lambda = 0:", round(AIC(model),2), "\n")
cat("AIC with Estimated Lambda:", round(AIC(ML1SINEs),2), "\n")
cat("AIC with Lambda = 1:", round(AIC(model_lambda_1),2), "\n")

########### Multiple PGLS Controlling for Longevity 
########### (Neoplasia or Malignancy ~ (Number of L1 OR All nLTRs) + Longevity + e) 

hist(mydata$maximum_longevity_m)

########### Check for the potential outliers in Longevity data
grubbs_result <- grubbs.test(mydata$maximum_longevity_m)
print(grubbs_result)

# We did not remove two influential points in our analyses. 
# See Methods for more detail ... 
########## Remove the most influential data point (elephant and Chimpanzee)
#mydata <- mydata[mydata$maximum_longevity_m != (max(mydata$maximum_longevity_m)), ]
#mydata <- mydata[mydata$maximum_longevity_m != (max(mydata$maximum_longevity_m)), ]
#nrow(mydata)

########### Pearson Correlation between Neoplasia/Malignancy and Longevity
cor(mydata$NeoplasiaPrevalence, mydata$maximum_longevity_m)
cor(mydata$MalignancyPrevalence, mydata$maximum_longevity_m)

# nLTR activities are also shown to be associated with longevity
# Test for any potential correlation 
#cor(mydata$maximum_longevity_m, mydata$Transformed_L1_Counts)
#cor(mydata$maximum_longevity_m, mydata$Transformed_L1_SINEs_Counts)

#### Neoplasia vs L1 + Longevity
NL1Lon <- pglsSEyPagel(NeoplasiaPrevalence ~ Transformed_L1_Counts +  maximum_longevity_m, data = mydata, tree=tre, se=SE, method = "ML")
summary(NL1Lon)
anova(NL1Lon)
# FDR test and extracting the adj. p-values from the model
NL1Lon_fdr_corrected <- p.adjust(summary(NL1Lon)$tTable[, "p-value"], method = "fdr")
NL1Lon_fdr_corrected
# VIF (variance inflation factor) to ensure multicollinearity is not an issue. 
# If VIF < 5, it's safe to include both variables in the model.
vif(NL1Lon)
# Grab likelihood-based R2, Lambda, and p values 
RelNL1Lon  <- R2_lik(NL1Lon)
n <- nrow(mydata)
p <- 2  # Number of predictors (L1s and longevity)
adjRelNL1Lon  <- signif(1 - ((1 - RelNL1Lon) * (n - 1)) / (n - p - 1), 2)
adjRelNL1Lon
# Check for normality of residuals 
ols_test_normality(NL1Lon$residuals)
hist(NL1Lon$residuals, main = "NL1Lon Residual Histogram")
qqnorm(NL1Lon$residuals)
qqline(NL1Lon$residuals)
# Check for independence of errors (autocorrelation) 
# Autocorrelation occurs when the residuals of a regression model are correlated
# with each other, indicating that there is some pattern or dependence among the errors.
model <- lm(NeoplasiaPrevalence ~ Transformed_L1_Counts +  maximum_longevity_m , data=mydata)
durbinWatsonTest(model) #Durbin-Watson test using car package
# Check for heteroscedasticity (Breusch-Pagan test)
# Checks if the variance of the errors is the same for every value of x (homoscedasticity). 
# Look for a random scatter of points around zero, with no discernible pattern or trend. 
# If the spread of residuals appears to vary systematically as the predicted
# values or predictor variable(s) change, it suggests potential heteroscedasticity.
plot(NL1Lon, which = 1)  # Residuals vs. Fitted value plot
plot(NL1Lon, which = 3)  # Scale-Location plot (square root of standardized residuals vs. Fitted values)
bptest(model)
### AIC Model Comparison
# Load the data and prepare comparative data object for AIC comparisons
# comparative_data <- comparative.data(phy = tre, data = mydata, 
#                                      names.col = "species", 
#                                      vcv = TRUE, na.omit = FALSE)
# Lambda fixed at 1 (full phylogenetic signal)
model_lambda_1 <- pgls(NeoplasiaPrevalence ~ Transformed_L1_Counts + maximum_longevity_m, data = comparative_data, lambda = 1.00)
# Compare AIC values
cat("AIC with Lambda = 0:", round(AIC(model),2), "\n")
cat("AIC with Estimated Lambda:", round(AIC(NL1Lon),2), "\n")
cat("AIC with Lambda = 1:", round(AIC(model_lambda_1),2), "\n")

#### Neoplasia vs (L1 + SINES) + Longevity
NL1SINEsLon <- pglsSEyPagel(NeoplasiaPrevalence ~ Transformed_L1_SINEs_Counts +  maximum_longevity_m, data = mydata, tree=tre, se=SE, method = "ML")
summary(NL1SINEsLon)
anova(NL1SINEsLon)
# FDR test and extracting the adj. p-values from the model
NL1SINEsLon_fdr_corrected <- p.adjust(summary(NL1SINEsLon)$tTable[, "p-value"], method = "fdr")
NL1SINEsLon_fdr_corrected
# VIF (variance inflation factor) to ensure multicollinearity is not an issue. 
# If VIF < 5, it's safe to include both variables in the model.
vif(NL1SINEsLon)
# Grab likelihood-based R2, Lambda, and p values 
RelNL1SINEsLon  <- R2_lik(NL1SINEsLon)
n <- nrow(mydata)
p <- 2  # Number of predictors (L1s and longevity)
adjRelNL1SINEsLon  <- signif(1 - ((1 - RelNL1SINEsLon) * (n - 1)) / (n - p - 1), 2)
adjRelNL1SINEsLon
# Check for normality of residuals 
ols_test_normality(NL1Lon$residuals)
hist(NL1Lon$residuals, main = "NL1Lon Residual Histogram")
qqnorm(NL1Lon$residuals)
qqline(NL1Lon$residuals)
# Check for independence of errors (autocorrelation) 
# Autocorrelation occurs when the residuals of a regression model are correlated
# with each other, indicating that there is some pattern or dependence among the errors.
model <- lm(NeoplasiaPrevalence ~ Transformed_L1_SINEs_Counts +  maximum_longevity_m , data=mydata)
durbinWatsonTest(model) #Durbin-Watson test using car package
# Check for heteroscedasticity (Breusch-Pagan test)
# Checks if the variance of the errors is the same for every value of x (homoscedasticity). 
# Look for a random scatter of points around zero, with no discernible pattern or trend. 
# If the spread of residuals appears to vary systematically as the predicted
# values or predictor variable(s) change, it suggests potential heteroscedasticity.
plot(NL1Lon, which = 1)  # Residuals vs. Fitted value plot
plot(NL1Lon, which = 3)  # Scale-Location plot (square root of standardized residuals vs. Fitted values)
bptest(model)
### AIC Model Comparison
# Load the data and prepare comparative data object for AIC comparisons
# comparative_data <- comparative.data(phy = tre, data = mydata, 
#                                      names.col = "species", 
#                                      vcv = TRUE, na.omit = FALSE)
# Lambda fixed at 1 (full phylogenetic signal)
model_lambda_1 <- pgls(NeoplasiaPrevalence ~ Transformed_L1_SINEs_Counts + maximum_longevity_m, data = comparative_data, lambda = 1.00)
# Compare AIC values
cat("AIC with Lambda = 0:", round(AIC(model),2), "\n")
cat("AIC with Estimated Lambda:", round(AIC(NL1SINEsLon),2), "\n")
cat("AIC with Lambda = 1:", round(AIC(model_lambda_1),2), "\n")

#### Malignancy vs L1 + Longevity
ML1Lon <- pglsSEyPagel(MalignancyPrevalence ~ Transformed_L1_Counts +  maximum_longevity_m, data = mydata, tree=tre, se=SE, method = "ML")
summary(ML1Lon)
anova(ML1Lon)
# FDR test and extracting the adj. p-values from the model
ML1Lon_fdr_corrected <- p.adjust(summary(ML1Lon)$tTable[, "p-value"], method = "fdr")
ML1Lon_fdr_corrected
# VIF (variance inflation factor) to ensure multicollinearity is not an issue. 
# If VIF < 5, it's safe to include both variables in the model.
vif(ML1Lon)
# Grab likelihood-based R2, Lambda, and p values 
RelML1Lon  <- R2_lik(ML1Lon)
n <- nrow(mydata)
p <- 2  # Number of predictors (L1s and longevity)
adjRelML1Lon  <- signif(1 - ((1 - RelML1Lon) * (n - 1)) / (n - p - 1), 2)
adjRelML1Lon
# Check for normality of residuals 
ols_test_normality(ML1Lon$residuals)
hist(ML1Lon$residuals, main = "ML1Lon Residual Histogram")
qqnorm(ML1Lon$residuals)
qqline(ML1Lon$residuals)
# Check for independence of errors (autocorrelation) 
# Autocorrelation occurs when the residuals of a regression model are correlated
# with each other, indicating that there is some pattern or dependence among the errors.
model <- lm(MalignancyPrevalence ~ Transformed_L1_Counts +  maximum_longevity_m , data=mydata)
durbinWatsonTest(model) #Durbin-Watson test using car package
# Check for heteroscedasticity (Breusch-Pagan test)
# Checks if the variance of the errors is the same for every value of x (homoscedasticity). 
# Look for a random scatter of points around zero, with no discernible pattern or trend. 
# If the spread of residuals appears to vary systematically as the predicted
# values or predictor variable(s) change, it suggests potential heteroscedasticity.
plot(ML1Lon, which = 1)  # Residuals vs. Fitted value plot
plot(ML1Lon, which = 3)  # Scale-Location plot (square root of standardized residuals vs. Fitted values)
bptest(model)
### AIC Model Comparison
# Load the data and prepare comparative data object for AIC comparisons
# comparative_data <- comparative.data(phy = tre, data = mydata, 
#                                      names.col = "species", 
#                                      vcv = TRUE, na.omit = FALSE)
# Lambda fixed at 1 (full phylogenetic signal)
model_lambda_1 <- pgls(MalignancyPrevalence ~ Transformed_L1_Counts + maximum_longevity_m, data = comparative_data, lambda = 1.00)
# Compare AIC values
cat("AIC with Lambda = 0:", round(AIC(model),2), "\n")
cat("AIC with Estimated Lambda:", round(AIC(ML1Lon),2), "\n")
cat("AIC with Lambda = 1:", round(AIC(model_lambda_1),2), "\n")

#### Malignancy vs (L1 + SINES) + Longevity
ML1SINEsLon <- pglsSEyPagel(MalignancyPrevalence ~ Transformed_L1_SINEs_Counts +  maximum_longevity_m, data = mydata, tree=tre, se=SE, method = "ML")
summary(ML1SINEsLon)
anova(ML1SINEsLon)
# FDR test and extracting the adj. p-values from the model
ML1SINEsLon_fdr_corrected <- p.adjust(summary(ML1SINEsLon)$tTable[, "p-value"], method = "fdr")
ML1SINEsLon_fdr_corrected
# VIF (variance inflation factor) to ensure multicollinearity is not an issue. 
# If VIF < 5, it's safe to include both variables in the model.
vif(ML1SINEsLon)
# Grab likelihood-based R2, Lambda, and p values 
RelML1SINEsLon  <- R2_lik(ML1SINEsLon)
n <- nrow(mydata)
p <- 2  # Number of predictors (L1s and longevity)
adjRelML1SINEsLon  <- signif(1 - ((1 - RelML1SINEsLon) * (n - 1)) / (n - p - 1), 2)
adjRelML1SINEsLon
# Check for normality of residuals 
ols_test_normality(ML1SINEsLon$residuals)
hist(ML1SINEsLon$residuals, main = "NL1Lon Residual Histogram")
qqnorm(ML1SINEsLon$residuals)
qqline(ML1SINEsLon$residuals)
# Check for independence of errors (autocorrelation) 
# Autocorrelation occurs when the residuals of a regression model are correlated
# with each other, indicating that there is some pattern or dependence among the errors.
model <- lm(MalignancyPrevalence ~ Transformed_L1_SINEs_Counts +  maximum_longevity_m , data=mydata)
durbinWatsonTest(model) #Durbin-Watson test using car package
# Check for heteroscedasticity (Breusch-Pagan test)
# Checks if the variance of the errors is the same for every value of x (homoscedasticity). 
# Look for a random scatter of points around zero, with no discernible pattern or trend. 
# If the spread of residuals appears to vary systematically as the predicted
# values or predictor variable(s) change, it suggests potential heteroscedasticity.
plot(ML1SINEsLon, which = 1)  # Residuals vs. Fitted value plot
plot(ML1SINEsLon, which = 3)  # Scale-Location plot (square root of standardized residuals vs. Fitted values)
bptest(model)
### AIC Model Comparison
# Load the data and prepare comparative data object for AIC comparisons
# comparative_data <- comparative.data(phy = tre, data = mydata, 
#                                      names.col = "species", 
#                                      vcv = TRUE, na.omit = FALSE)
# Lambda fixed at 1 (full phylogenetic signal)
model_lambda_1 <- pgls(MalignancyPrevalence ~ Transformed_L1_SINEs_Counts + maximum_longevity_m, data = comparative_data, lambda = 1.00)
# Compare AIC values
cat("AIC with Lambda = 0:", round(AIC(model),2), "\n")
cat("AIC with Estimated Lambda:", round(AIC(ML1SINEs),2), "\n")
cat("AIC with Lambda = 1:", round(AIC(model_lambda_1),2), "\n")
```
