## Supplementary Scripts for "Recent Non-LTR Retrotransposon Activity Predicts Cancer Prevalence in Mammals": 01_nLTR_Abundance_Plotting.R

```
################################################################################
####################### nLTR Counts Plotting
########

################################################### 
###  Neoplasia vs L1 counts
################################################### 
# Define the custom color palette for the Orders
order_colors <- c("Carnivora" = "#414487FF",
                  "Artiodactyla" = "#2A788EFF",
                  "Primates" = "#440154FF",
                  "Rodentia" = "#FCE205",
                  "Chiroptera" = "#22A884FF", 
                  "Other Orders" = "#7AD151FF")

# Store Current model and Response variable in temp vars 
Current_model <- NL1
Current_Response <- mydata$NeoplasiaPrevalence

# Calculate fitted values
fitted_values <- predict(Current_model)
mydata$nudge_y <- ifelse(Current_Response > fitted_values, 0.02, -0.02)

# Classify orders
mydata$color_group <- ifelse(mydata$Order %in% c("Rodentia", "Artiodactyla", "Carnivora", "Chiroptera", "Primates"), 
                             mydata$Order, 
                             "Other Orders")

# Clean species names
mydata$species_clean <- gsub("_", " ", mydata$species)

# Dummy variable for legend formatting
mydata$ModelStats <- "Model Stats"

# Final Plot
plot <- ggplot(mydata, aes(x = Transformed_L1_Counts, y = NeoplasiaPrevalence)) +
  
  # Colored dots
  geom_point(aes(color = color_group), size = 3, alpha = 0.9) +
  
  # Species labels (all species)
  geom_text_repel(
    data = mydata,
    aes(label = Common_Name),
    size = 2.5,
    color = "grey20",
    box.padding = 0.3,
    point.padding = 0.8,
    segment.color = "grey60",
    segment.size = 0.3,
    segment.curvature = 0,
    segment.angle = 20,
    min.segment.length = 0.01,
    max.overlaps = Inf
  ) +
  
  # Regression line
  geom_smooth(method = "lm", se = FALSE, color = "grey50", alpha = 0.7, 
              fullrange = TRUE, linewidth = 0.9) +
  
  # Axis labels
  xlab("Number of Active L1")+#&#8201;<span style='font-size:8pt;'>(Tukey-Transformed)</span>") +
  ylab("Neoplasia Prevalence")+#&#8201;<span style='font-size:8pt;'>(%)</span>") +
  
  
  # Color legend with model stats
  scale_color_manual(
    name = paste0(
      "<b style='font-size:11pt;'>Model Stats</b><br>",
      nrow(mydata), " species<br>",
      "Corr = ", corrNeo, "<br>",
      "p value = ", NL1_pvalue, "<span style='color:red;'><b>*</b></span><br>",
      "R² = ", RelNL1, "<br>",
      "λ = ", lambdaNL1, "<br><br>",
      "<b style='font-size:11pt;'>Orders</b>"
    ),
    values = order_colors
  ) +
  
  # Axis limits
  scale_x_continuous(limits = c(-0.08, 1.1), expand = c(0, 0)) +
  scale_y_continuous(limits = c(0.1, 0.97), expand = c(0, 0)) +
  
  # Theme
  theme_minimal(base_size = 6) +
  theme(
    legend.position = "right",
    legend.justification = "bottom",
    legend.direction = "vertical",
    legend.title = element_markdown(size = 10, color = "grey30", lineheight = 1.75),
    legend.text = element_text(size = 9, color = "grey40"),
    plot.title = element_text(size = 12, hjust = 0.5, color = "grey30"),
    axis.title.x = ggtext::element_markdown(size = 12, color = "grey30", margin = margin(t = 10)),
    axis.title.y = ggtext::element_markdown(size = 12, color = "grey30", margin = margin(r = 10)),
    axis.text = element_text(size = 10),
    panel.background = element_rect(fill = "white", color = "white"),
    plot.background = element_rect(fill = "white", color = "white"),
    panel.grid.major = element_blank(),
    panel.grid.minor = element_blank(),
    axis.line = element_line(color = "grey40", linewidth = 0.9),
    panel.border = element_rect(color = "grey40", fill = NA, size = 0.9),
    plot.margin = unit(c(0.5, 0.5, 0.5, 0.5), "cm")
  )

# Print plot
print(plot)

# Save PNG with exact name
ggsave("1_L1_Counts_vs_Neoplasia_N.png", plot, width = 10, height = 8, dpi = 800)
# Save RDS
saveRDS(plot, "1_L1_Counts_vs_Neoplasia_N.rds")

################################################### 
###  Neoplasia vs L1-SINEs counts
################################################### 

# Store Current model and Response variable in temp vars 
Current_model <- NL1SINEs
Current_Response <- mydata$NeoplasiaPrevalence

# Fitted values (optional, but nudge_y is not used)
fitted_values <- predict(Current_model)
mydata$nudge_y <- ifelse(Current_Response > fitted_values, 0.02, -0.02)

# Classify Orders
mydata$color_group <- ifelse(mydata$Order %in% c("Rodentia", "Artiodactyla", "Carnivora", "Chiroptera", "Primates"),
                             mydata$Order, 
                             "Other Orders")

# Clean species name
mydata$species_clean <- gsub("_", " ", mydata$species)

# Dummy variable for legend title
mydata$ModelStats <- "Model Stats"

# Final Plot
plot <- ggplot(mydata, aes(x = Transformed_L1_SINEs_Counts, y = NeoplasiaPrevalence)) +
  
  # Colored points
  geom_point(aes(color = color_group), size = 3, alpha = 0.9) +
  
  # Labels for all species
  geom_text_repel(
    data = mydata,
    aes(label = Common_Name),
    size = 2.5,
    color = "grey20",
    box.padding = 0.3,
    point.padding = 0.8,
    segment.color = "grey60",
    segment.size = 0.3,
    segment.curvature = 0,
    segment.angle = 20,
    min.segment.length = 0.01,
    max.overlaps = Inf
  ) +
  
  # Regression line
  geom_smooth(method = "lm", se = FALSE, color = "grey50", alpha = 0.7,
              fullrange = TRUE, linewidth = 0.9) +
  
  # Axis labels
  xlab("Combined Number of Active L1 and SINEs")+#&#8201;<span style='font-size:8pt;'>(Tukey-Transformed)</span>") +
  ylab("Neoplasia Prevalence")+#&#8201;<span style='font-size:8pt;'>(%)</span>") +
  
  
  # Legend with model stats
  scale_color_manual(
    name = paste0(
      "<b style='font-size:11pt;'>Model Stats</b><br>",
      nrow(mydata), " species<br>",
      "Corr = ", corrNeo2, "<br>",
      "p value = ", NL1SINEs_pvalue, "<span style='color:red;'><b>*</b></span><br>",
      "R² = ", RelNL1SINEs, "<br>",
      "λ = ", lambdaNL1SINEs, "<br><br>",
      "<b style='font-size:11pt;'>Orders</b>"
    ),
    values = order_colors
  ) +
  
  # Axis limits
  scale_x_continuous(limits = c(-0.08, 1.1), expand = c(0, 0)) +
  scale_y_continuous(limits = c(0.1, 0.97), expand = c(0, 0)) +
  
  # Clean theme
  theme_minimal(base_size = 6) +
  theme(
    legend.position = "right",
    legend.justification = "bottom",
    legend.direction = "vertical",
    legend.title = element_markdown(size = 10, color = "grey30", lineheight = 1.75),
    legend.text = element_text(size = 9, color = "grey40"),
    plot.title = element_text(size = 12, hjust = 0.5, color = "grey30"),
    axis.title.x = ggtext::element_markdown(size = 12, color = "grey30", margin = margin(t = 10)),
    axis.title.y = ggtext::element_markdown(size = 12, color = "grey30", margin = margin(r = 10)),
    axis.text = element_text(size = 10),
    panel.background = element_rect(fill = "white", color = "white"),
    plot.background = element_rect(fill = "white", color = "white"),
    panel.grid.major = element_blank(),
    panel.grid.minor = element_blank(),
    axis.line = element_line(color = "grey40", linewidth = 0.9),
    panel.border = element_rect(color = "grey40", fill = NA, size = 0.9),
    plot.margin = unit(c(0.5, 0.5, 0.5, 0.5), "cm")
  )

# Print
print(plot)

# Save with original file name
ggsave("2_L1_SINEs_Counts_vs_Neoplasia_N.png", plot, width = 10, height = 8, dpi = 800)
# Save RDS
saveRDS(plot, "2_L1_SINEs_Counts_vs_Neoplasia_N.rds")

################################################### 
###  Malignancy vs L1 counts
################################################### 

# Store Current model and Response variable in temp vars 
Current_model <- ML1
Current_Response <- mydata$MalignancyPrevalence

# Optional: fitted values (nudge_y not used)
fitted_values <- predict(Current_model)
mydata$nudge_y <- ifelse(Current_Response > fitted_values, 0.02, -0.02)

# Classify orders for coloring
mydata$color_group <- ifelse(mydata$Order %in% c("Rodentia", "Artiodactyla", "Carnivora", "Chiroptera", "Primates"),
                             mydata$Order,
                             "Other Orders")

# Clean species names
mydata$species_clean <- gsub("_", " ", mydata$species)

# Dummy variable for legend formatting
mydata$ModelStats <- "Model Stats"

# Final plot
plot <- ggplot(mydata, aes(x = Transformed_L1_Counts, y = MalignancyPrevalence)) +
  
  # Colored points by order
  geom_point(aes(color = color_group), size = 3, alpha = 0.9) +
  
  # Labels for all species
  geom_text_repel(
    data = mydata,
    aes(label = Common_Name),
    size = 2.5,
    color = "grey20",
    box.padding = 0.3,
    point.padding = 0.8,
    segment.color = "grey60",
    segment.size = 0.3,
    segment.curvature = 0,
    segment.angle = 20,
    min.segment.length = 0.01,
    max.overlaps = Inf
  ) +
  
  # Regression line
  geom_smooth(method = "lm", se = FALSE, color = "grey50", alpha = 0.7,
              fullrange = TRUE, linewidth = 0.9) +
  
  # Axis titles
  xlab("Number of Active L1")+#&#8201;<span style='font-size:8pt;'>(Tukey-Transformed)</span>") +
  ylab("Malignancy Prevalence")+#&#8201;<span style='font-size:8pt;'>(%)</span>") +
  
  
  # Legend with model stats
  scale_color_manual(
    name = paste0(
      "<b style='font-size:11pt;'>Model Stats</b><br>",
      nrow(mydata), " species<br>",
      "Corr = ", corrMal, "<br>",
      "p value = ", ML1_pvalue, "<br>",
      "R² = ", RelML1, "<br>",
      "λ = ", lambdaML1, "<br><br>",
      "<b style='font-size:11pt;'>Orders</b>"
    ),
    values = order_colors
  ) +
  
  # Axis limits
  scale_x_continuous(limits = c(-0.08, 1.1), expand = c(0, 0)) +
  scale_y_continuous(limits = c(0.1, 0.97), expand = c(0, 0)) +
  
  # Theme
  theme_minimal(base_size = 6) +
  theme(
    legend.position = "right",
    legend.justification = "bottom",
    legend.direction = "vertical",
    legend.title = element_markdown(size = 10, color = "grey30", lineheight = 1.75),
    legend.text = element_text(size = 9, color = "grey40"),
    plot.title = element_text(size = 12, hjust = 0.5, color = "grey30"),
    axis.title.x = ggtext::element_markdown(size = 12, color = "grey30", margin = margin(t = 10)),
    axis.title.y = ggtext::element_markdown(size = 12, color = "grey30", margin = margin(r = 10)),
    axis.text = element_text(size = 10),
    panel.background = element_rect(fill = "white", color = "white"),
    plot.background = element_rect(fill = "white", color = "white"),
    panel.grid.major = element_blank(),
    panel.grid.minor = element_blank(),
    axis.line = element_line(color = "grey40", linewidth = 0.9),
    panel.border = element_rect(color = "grey40", fill = NA, size = 0.9),
    plot.margin = unit(c(0.5, 0.5, 0.5, 0.5), "cm")
  )

# Print
print(plot)

# Save with original file names
ggsave("3_L1_Counts_vs_Malignancy_N.png", plot, width = 10, height = 8, dpi = 800)
# Save RDS
saveRDS(plot, "3_L1_Counts_vs_Malignancy_N.rds")


################################################### 
###  Malignancy vs L1-SINEs counts
################################################### 

# Store Current model and Response variable in temp vars 
Current_model <- ML1SINEs
Current_Response <- mydata$MalignancyPrevalence

# (Optional) Predict fitted values
fitted_values <- predict(Current_model)
mydata$nudge_y <- ifelse(Current_Response > fitted_values, 0.02, -0.02)

# Define coloring groups
mydata$color_group <- ifelse(mydata$Order %in% c("Rodentia", "Artiodactyla", "Carnivora", "Chiroptera", "Primates"),
                             mydata$Order, "Other Orders")

# Clean species names
mydata$species_clean <- gsub("_", " ", mydata$species)

# Dummy column for legend title
mydata$ModelStats <- "Model Stats"

# Final Plot
plot <- ggplot(mydata, aes(x = Transformed_L1_SINEs_Counts, y = MalignancyPrevalence)) +
  
  # Plot dots
  geom_point(aes(color = color_group), size = 3, alpha = 0.9) +
  
  # All species labels
  geom_text_repel(
    data = mydata,
    aes(label = Common_Name),
    size = 2.5,
    color = "grey20",
    box.padding = 0.3,
    point.padding = 0.8,
    segment.color = "grey60",
    segment.size = 0.3,
    segment.curvature = 0,
    segment.angle = 20,
    min.segment.length = 0.01,
    max.overlaps = Inf
  ) +
  
  # Regression line
  geom_smooth(method = "lm", se = FALSE, color = "grey50", alpha = 0.7,
              fullrange = TRUE, linewidth = 0.9) +
  
  # Axis labels
  xlab("Combined Number of Active L1 and SINEs")+#&#8201;<span style='font-size:8pt;'>(Tukey-Transformed)</span>") +
  ylab("Malignancy Prevalence")+#&#8201;<span style='font-size:8pt;'>(%)</span>") +
  
  # Manual colors and custom legend
  scale_color_manual(
    name = paste0(
      "<b style='font-size:11pt;'>Model Stats</b><br>",
      nrow(mydata), " species<br>",
      "Corr = ", corrMal2, "<br>",
      "p value = ", ML1SINEs_pvalue, "<span style='color:red;'><b>*</b></span><br>",
      "R² = ", RelML1SINEs, "<br>",
      "λ = ", lambdaML1SINEs, "<br><br>",
      "<b style='font-size:11pt;'>Orders</b>"
    ),
    values = order_colors
  ) +
  
  # Axis limits
  scale_x_continuous(limits = c(-0.08, 1.1), expand = c(0, 0)) +
  scale_y_continuous(limits = c(0.1, 0.97), expand = c(0, 0)) +
  
  # Theme
  theme_minimal(base_size = 6) +
  theme(
    legend.position = "right",
    legend.justification = "bottom",
    legend.direction = "vertical",
    legend.title = element_markdown(size = 10, color = "grey30", lineheight = 1.75),
    legend.text = element_text(size = 9, color = "grey40"),
    plot.title = element_text(size = 12, hjust = 0.5, color = "grey30"),
    axis.title.x = ggtext::element_markdown(size = 12, color = "grey30", margin = margin(t = 10)),
    axis.title.y = ggtext::element_markdown(size = 12, color = "grey30", margin = margin(r = 10)),
    axis.text = element_text(size = 10),
    panel.background = element_rect(fill = "white", color = "white"),
    plot.background = element_rect(fill = "white", color = "white"),
    panel.grid.major = element_blank(),
    panel.grid.minor = element_blank(),
    axis.line = element_line(color = "grey40", linewidth = 0.9),
    panel.border = element_rect(color = "grey40", fill = NA, size = 0.9),
    plot.margin = unit(c(0.5, 0.5, 0.5, 0.5), "cm")
  )

# Print
print(plot)

# Save with original file names
ggsave("4_L1_SINEs_Counts_vs_Malignancy_N.png", plot, width = 10, height = 8, dpi = 800)
# Save RDS
saveRDS(plot, "4_L1_SINEs_Counts_vs_Malignancy_N.rds")
```
