## Supplementary Scripts for "Recent Non-LTR Retrotransposon Activity Predicts Cancer Prevalence in Mammals": 08_CGOs_Analyses.R

```
###################################################################################################
######################################### CGOs Analyses
#################
rm(list=ls(all=TRUE))
ls()

#####################################################################
suppressPackageStartupMessages({
  library(ggplot2)
  library(reshape2)
  library(plyr)
  library(tidyverse)
  library(dplyr)
  library(rcompanion)# Tukey Transformation
  library(car) # Durbin Watson Test
  library(phytools) # PGLS 
  library(olsrr) # Assess the normality of residuals
  library(MASS) # Transformation
  library(lmtest) # Breusch_Pagan test
  library(outliers) # To identify outliers
  library(ggimage) # Plotting
  library(magick) # Plotting
  library(ggtree)
  library(cowplot) # For Plotting
  library(ggtext) # For legends in plots
  library(ggrepel) # To avoid overlaps between labels in plots
  library(viridis) # Color blind palette for plots
  library(caper) # For reading tree
  library(rr2) # For reading tree
  library(devtools) # For RPICR
  #library(ROBRT) # For RPICR > You need to use install_github from devtools
  library(geiger) # for RPICR
})
#####################################################################
########## Load custom PGLS functions into the current session
source("Path to /0_pglsSEyPagel.R")
#####################################################################
setwd("Path to the working directory")
getwd()

# Reading Excel data 
mydata <- read.csv("Path to the Supplementary Data S1/Supplementary Data S1.csv", header = TRUE)

#####################################################################
########## Load the ultrametric phylogenetic tree
tre <- read.tree("Mammals_tree.newick")

#####################################################################
# Data preparation
#####################################################################
rownames(mydata) <- mydata$species

########## Setting SE
SE <- setNames(mydata$SE, mydata$species)[rownames(mydata)]
nrow(mydata)

########### To remove species with peptide BUSCO score <60%
# Identify species to be removed
removed_species <- mydata$species[mydata$Peptide_BUSCO_Scores <= 60]
print(removed_species)

#Removing
mydata <- mydata %>%
  filter(Peptide_BUSCO_Scores >= 60)

nrow(mydata)

########## CGOs Stats
# Top 5 with highest Total_CGOs
top5 <- mydata[order(-mydata$Total_CGOs), c("Common_Name", "Total_CGOs")][1:5, ]
# top5 <- mydata[order(-mydata$Total_CGOs), c("Common_Name", "Oncogene")][1:5, ]
# top5 <- mydata[order(-mydata$Total_CGOs), c("Common_Name", "TSG")][1:5, ]

# Top 5 with lowest Total_CGOs
bottom5 <- mydata[order(mydata$Total_CGOs), c("Common_Name", "Total_CGOs")][1:5, ]
# bottom5 <- mydata[order(mydata$Total_CGOs), c("Common_Name", "Oncogene")][1:5, ]
# bottom5 <- mydata[order(mydata$Total_CGOs), c("Common_Name", "TSG")][1:5, ]

# Print results
print("Top 5 species with highest Total_CGOs:")
print(top5)
print("Top 5 species with lowest Total_CGOs:")
print(bottom5)

####****************************************************************************
#### Mouse and Rat are influential points in this analysis. Mouse also stands as 
#### as oultlier in Oncogene analysis. However, removing it from the analysis does
#### not change the significance of the results.  
#mydata <- subset(mydata, species != "Mus_musculus")
#mydata <- subset(mydata, species != "Rattus_norvegicus")
####****************************************************************************

########## Check for normality in cancer data
hist(mydata$NeoplasiaPrevalence)
hist(mydata$MalignancyPrevalence)

########## Transform data to mitigate skewness and make the data follow a normal distribution
### Tukey's Ladder of Powers transform 
mydata$NeoplasiaPrevalence <- transformTukey(mydata$NeoplasiaPrevalence)
mydata$MalignancyPrevalence <- transformTukey(mydata$MalignancyPrevalence)
hist(mydata$NeoplasiaPrevalence)
hist(mydata$MalignancyPrevalence)

########## Check for the poteintial outliers in cancer data
grubbs_result <- grubbs.test(mydata$NeoplasiaPrevalence)
print(grubbs_result)
grubbs_result <- grubbs.test(mydata$MalignancyPrevalence)
print(grubbs_result)

########## Define predictors (Total_CGOs, Somatic, Germline, Oncogene, TSG, Fusion)

# No transformation needed!
# Just scale data to the 0,1 range

hist(mydata$Total_CGOs)
#mydata$Total_CGOs <- transformTukey(mydata$Total_CGOs)
#hist(mydata$Total_CGOs) # Transforming makes the distribution worse!
mydata$Total_CGOs <- (mydata$Total_CGOs - min(mydata$Total_CGOs)) / (max(mydata$Total_CGOs) - min(mydata$Total_CGOs))
hist(mydata$Total_CGOs)

hist(mydata$Somatic)
#mydata$Somatic <- transformTukey(mydata$Somatic)
#hist(mydata$Somatic)
mydata$Somatic <- (mydata$Somatic - min(mydata$Somatic)) / (max(mydata$Somatic) - min(mydata$Somatic))
hist(mydata$Somatic)

hist(mydata$Germline)
#mydata$Germline <- transformTukey(mydata$Germline)
#hist(mydata$Germline)
mydata$Germline <- (mydata$Germline - min(mydata$Germline)) / (max(mydata$Germline) - min(mydata$Germline))
hist(mydata$Germline)

hist(mydata$Oncogene)
#mydata$Oncogene <- transformTukey(mydata$Oncogene)
#hist(mydata$Oncogene)
mydata$Oncogene <- (mydata$Oncogene - min(mydata$Oncogene)) / (max(mydata$Oncogene) - min(mydata$Oncogene))
hist(mydata$Oncogene)

hist(mydata$TSG)
#mydata$TSG <- transformTukey(mydata$TSG)
#hist(mydata$TSG)
mydata$TSG <- (mydata$TSG - min(mydata$TSG)) / (max(mydata$TSG) - min(mydata$TSG))
hist(mydata$TSG)

hist(mydata$Fusion)
#mydata$Fusion <- transformTukey(mydata$Fusion)
#hist(mydata$Fusion)
mydata$Fusion <- (mydata$Fusion - min(mydata$Fusion)) / (max(mydata$Fusion) - min(mydata$Fusion))
hist(mydata$Fusion)

########### Check for the potential outliers in nLTR counts
grubbs_result <- grubbs.test(mydata$Total_CGOs) # needs
print(grubbs_result)
grubbs_result <- grubbs.test(mydata$Somatic) #needs
print(grubbs_result)
grubbs_result <- grubbs.test(mydata$Germline)
print(grubbs_result)
grubbs_result <- grubbs.test(mydata$Oncogene) #needs
print(grubbs_result)
grubbs_result <- grubbs.test(mydata$TSG) 
print(grubbs_result)
grubbs_result <- grubbs.test(mydata$Fusion) #needs
print(grubbs_result)

# Get the top 10 species with the highest Total_CGOs values
# top_species <- mydata %>%
#   arrange(desc(Total_CGOs)) %>%
#   dplyr::select(species, Total_CGOs) %>%
#   head(10)
# # Print the result
# print(top_species)

########### To remove the potential outliers, if needed
#mydata <- mydata[mydata$Total_CGOs != (max(mydata$Total_CGOs)), ]
#nrow(mydata)

########## Pearson Correlations 
corrNeo1 <- round(cor(mydata$NeoplasiaPrevalence, mydata$Total_CGOs),2)
corrNeo1
corrNeo2 <- round(cor(mydata$NeoplasiaPrevalence, mydata$Somatic),2)
corrNeo2
corrNeo3 <- round(cor(mydata$NeoplasiaPrevalence, mydata$Germline),2)
corrNeo3
corrNeo4 <- round(cor(mydata$NeoplasiaPrevalence, mydata$Oncogene),2)
corrNeo4
corrNeo5 <- round(cor(mydata$NeoplasiaPrevalence, mydata$TSG),2)
corrNeo5
corrNeo6 <- round(cor(mydata$NeoplasiaPrevalence, mydata$Fusion),2)
corrNeo6

corrMal1 <- round(cor(mydata$MalignancyPrevalence, mydata$Total_CGOs),2)
corrMal1
corrMal2 <- round(cor(mydata$MalignancyPrevalence, mydata$Somatic),2)
corrMal2
corrMal3 <- round(cor(mydata$MalignancyPrevalence, mydata$Germline),2)
corrMal3
corrMal4 <- round(cor(mydata$MalignancyPrevalence, mydata$Oncogene),2)
corrMal4
corrMal5 <- round(cor(mydata$MalignancyPrevalence, mydata$TSG),2)
corrMal5
corrMal6 <- round(cor(mydata$MalignancyPrevalence, mydata$Fusion),2)
corrMal6

#######################################################################
### Running PGLS
#######################################################################

########### Simple PGLS (Cancer ~ Number of CGOs of various types + e)

################################
### Neoplasia vs Total_CGOs
################################
NTotal_CGOs <- pglsSEyPagel(NeoplasiaPrevalence ~ Total_CGOs, data = mydata, tree=tre, se=SE, method = "ML")
summary(NTotal_CGOs)
anova(NTotal_CGOs)
# Grab likelihood-based R2, Lambda, and p values for plotting
RelNTotal_CGOs <- R2_lik(NTotal_CGOs)
RelNTotal_CGOs <- format(RelNTotal_CGOs)
RelNTotal_CGOs <- signif(as.numeric(RelNTotal_CGOs), digits = 2)
RelNTotal_CGOs
lambdaNTotal_CGOs <- summary(NTotal_CGOs)$modelStruct$corStruct
lambdaNTotal_CGOs <- signif(lambdaNTotal_CGOs[1], digits = 2)
lambdaNTotal_CGOs
NTotal_CGOs_pvalue <- summary(NTotal_CGOs)$tTable
NTotal_CGOs_pvalue <- signif(NTotal_CGOs_pvalue[2,4], digits = 2)
NTotal_CGOs_pvalue
# Check for normality of residuals 
ols_test_normality(NTotal_CGOs$residuals)
hist(NTotal_CGOs$residuals, main = "NTotal_CGOs Residual Histogram")
qqnorm(NTotal_CGOs$residuals)
qqline(NTotal_CGOs$residuals)
# Check for independence of errors (autocorrelation) 
# Autocorrelation occurs when the residuals of a regression model are correlated
# with each other, indicating that there is some pattern or dependence among the errors.
model <- lm(NeoplasiaPrevalence ~ Total_CGOs , data=mydata)
durbinWatsonTest(model) #Durbin-Watson test using car package
# Check for heteroscedasticity (Breusch-Pagan test)
# Checks if the variance of the errors is the same for every value of x (homoscedasticity). 
# Look for a random scatter of points around zero, with no discernible pattern or trend. 
# If the spread of residuals appears to vary systematically as the predicted
# values or predictor variable(s) change, it suggests potential heteroscedasticity.
plot(NTotal_CGOs, which = 1)  # Residuals vs. Fitted value plot
plot(NTotal_CGOs, which = 3)  # Scale-Location plot (square root of standardized residuals vs. Fitted values)
bptest(lm(NTotal_CGOs$residuals ~ fitted(NTotal_CGOs)))
### AIC Model Comparison
# Load the data and prepare comparative data object for AIC comparisons
comparative_data <- comparative.data(phy = tre, data = mydata, 
                                     names.col = "species", 
                                     vcv = TRUE, na.omit = FALSE)
# Lambda fixed at 1 (full phylogenetic signal)
model_lambda_1 <- pgls(NeoplasiaPrevalence ~ Total_CGOs, data = comparative_data, lambda = 1.00)
# Compare AIC values
cat("AIC with Lambda = 0:", round(AIC(model),2), "\n")
cat("AIC with Estimated Lambda:", round(AIC(NTotal_CGOs),2), "\n")
cat("AIC with Lambda = 1:", round(AIC(model_lambda_1),2), "\n")
#******************************************
# Quadratic form to test for non-linearity
#******************************************
mydata$Total_CGOs2 <- mydata$Total_CGOs^2
NTotal_CGOs_quad <- pglsSEyPagel(NeoplasiaPrevalence ~ Total_CGOs + Total_CGOs2, data = mydata, tree=tre, se=SE, method = "ML")
summary(NTotal_CGOs_quad)
anova(NTotal_CGOs_quad)
plot(NTotal_CGOs_quad, which = 1)
#*******************************


### Neoplasia vs Somatic
NSomatic <- pglsSEyPagel(NeoplasiaPrevalence ~ Somatic, data = mydata, tree=tre, se=SE, method = "ML")
summary(NSomatic)
anova(NSomatic)
# Grab likelihood-based R2, Lambda, and p values for plotting
RelNSomatic <- R2_lik(NSomatic)
RelNSomatic <- format(RelNSomatic)
RelNSomatic <- signif(as.numeric(RelNSomatic), digits = 2)
RelNSomatic
lambdaNSomatic <- summary(NSomatic)$modelStruct$corStruct
lambdaNSomatic <- signif(lambdaNSomatic[1], digits = 2)
lambdaNSomatic
NSomatic_pvalue <- summary(NSomatic)$tTable
NSomatic_pvalue <- signif(NSomatic_pvalue[2,4], digits = 2)
NSomatic_pvalue
# Check for normality of residuals 
ols_test_normality(NSomatic$residuals)
hist(NSomatic$residuals, main = "NSomatic Residual Histogram")
qqnorm(NSomatic$residuals)
qqline(NSomatic$residuals)
# Check for independence of errors (autocorrelation) 
# Autocorrelation occurs when the residuals of a regression model are correlated
# with each other, indicating that there is some pattern or dependence among the errors.
model <- lm(NeoplasiaPrevalence ~ Somatic , data=mydata)
durbinWatsonTest(model) #Durbin-Watson test using car package
# Check for heteroscedasticity (Breusch-Pagan test)
# Checks if the variance of the errors is the same for every value of x (homoscedasticity). 
# Look for a random scatter of points around zero, with no discernible pattern or trend. 
# If the spread of residuals appears to vary systematically as the predicted
# values or predictor variable(s) change, it suggests potential heteroscedasticity.
plot(NSomatic, which = 1)  # Residuals vs. Fitted value plot
plot(NSomatic, which = 3)  # Scale-Location plot (square root of standardized residuals vs. Fitted values)
bptest(lm(NSomatic$residuals ~ fitted(NSomatic)))
### AIC Model Comparison
# Load the data and prepare comparative data object for AIC comparisons
# comparative_data <- comparative.data(phy = tre, data = mydata, 
#                                      names.col = "species", 
#                                      vcv = TRUE, na.omit = FALSE)
# Lambda fixed at 1 (full phylogenetic signal)
model_lambda_1 <- pgls(NeoplasiaPrevalence ~ Somatic, data = comparative_data, lambda = 1.00)
# Compare AIC values
cat("AIC with Lambda = 0:", round(AIC(model),2), "\n")
cat("AIC with Estimated Lambda:", round(AIC(NSomatic),2), "\n")
cat("AIC with Lambda = 1:", round(AIC(model_lambda_1),2), "\n")
#******************************************
# Quadratic form to test for non-linearity
#******************************************
# mydata$Somatic2 <- mydata$Somatic^2
# NSomatic_quad <- pglsSEyPagel(NeoplasiaPrevalence ~ Somatic + Somatic2, data = mydata, tree=tre, se=SE, method = "ML")
# summary(NSomatic_quad)
# anova(NSomatic_quad)
# plot(NSomatic_quad, which = 1)
#*******************************


### Neoplasia vs Germline
NGermline <- pglsSEyPagel(NeoplasiaPrevalence ~ Germline, data = mydata, tree=tre, se=SE, method = "ML")
summary(NGermline)
anova(NGermline)
# Grab likelihood-based R2, Lambda, and p values for plotting
RelNGermline <- R2_lik(NGermline)
RelNGermline <- format(RelNGermline)
RelNGermline <- signif(as.numeric(RelNGermline), digits = 2)
RelNGermline
lambdaNGermline <- summary(NGermline)$modelStruct$corStruct
lambdaNGermline <- signif(lambdaNGermline[1], digits = 2)
lambdaNGermline
NGermline_pvalue <- summary(NGermline)$tTable
NGermline_pvalue <- signif(NGermline_pvalue[2,4], digits = 2)
NGermline_pvalue
# Check for normality of residuals 
ols_test_normality(NGermline$residuals)
hist(NGermline$residuals, main = "NGermline Residual Histogram")
qqnorm(NGermline$residuals)
qqline(NGermline$residuals)
# Check for independence of errors (autocorrelation) 
# Autocorrelation occurs when the residuals of a regression model are correlated
# with each other, indicating that there is some pattern or dependence among the errors.
model <- lm(NeoplasiaPrevalence ~ Germline , data=mydata)
durbinWatsonTest(model) #Durbin-Watson test using car package
# Check for heteroscedasticity (Breusch-Pagan test)
# Checks if the variance of the errors is the same for every value of x (homoscedasticity). 
# Look for a random scatter of points around zero, with no discernible pattern or trend. 
# If the spread of residuals appears to vary systematically as the predicted
# values or predictor variable(s) change, it suggests potential heteroscedasticity.
plot(NGermline, which = 1)  # Residuals vs. Fitted value plot
plot(NGermline, which = 3)  # Scale-Location plot (square root of standardized residuals vs. Fitted values)
bptest(lm(NGermline$residuals ~ fitted(NGermline)))
### AIC Model Comparison
# Load the data and prepare comparative data object for AIC comparisons
# comparative_data <- comparative.data(phy = tre, data = mydata, 
#                                      names.col = "species", 
#                                      vcv = TRUE, na.omit = FALSE)
# Lambda fixed at 1 (full phylogenetic signal)
model_lambda_1 <- pgls(NeoplasiaPrevalence ~ Germline, data = comparative_data, lambda = 1.00)
# Compare AIC values
cat("AIC with Lambda = 0:", round(AIC(model),2), "\n")
cat("AIC with Estimated Lambda:", round(AIC(NGermline),2), "\n")
cat("AIC with Lambda = 1:", round(AIC(model_lambda_1),2), "\n")
#******************************************
# Quadratic form to test for non-linearity
#******************************************
# mydata$Germline2 <- mydata$Germline^2
# NGermline_quad <- pglsSEyPagel(NeoplasiaPrevalence ~ Germline + Germline2, data = mydata, tree=tre, se=SE, method = "ML")
# summary(NGermline_quad)
# anova(NGermline_quad)
# plot(NGermline_quad, which = 1)

# # Both the linear and quadratic components of Germline gene counts were significantly 
# # associated with neoplasia prevalence (p = 0.040 and p = 0.042, respectively), 
# # suggesting a non-linear relationship.
#*******************************


### Neoplasia vs Oncogene
NOncogene <- pglsSEyPagel(NeoplasiaPrevalence ~ Oncogene, data = mydata, tree=tre, se=SE, method = "ML")
summary(NOncogene)
anova(NOncogene)
# Grab likelihood-based R2, Lambda, and p values for plotting
RelNOncogene <- R2_lik(NOncogene)
RelNOncogene <- format(RelNOncogene)
RelNOncogene <- signif(as.numeric(RelNOncogene), digits = 2)
RelNOncogene
lambdaNOncogene <- summary(NOncogene)$modelStruct$corStruct
lambdaNOncogene <- signif(lambdaNOncogene[1], digits = 2)
lambdaNOncogene
NOncogene_pvalue <- summary(NOncogene)$tTable
NOncogene_pvalue <- signif(NOncogene_pvalue[2,4], digits = 2)
NOncogene_pvalue
# Check for normality of residuals 
ols_test_normality(NOncogene$residuals)
hist(NOncogene$residuals, main = "NOncogene Residual Histogram")
qqnorm(NOncogene$residuals)
qqline(NOncogene$residuals)
# Check for independence of errors (autocorrelation) 
# Autocorrelation occurs when the residuals of a regression model are correlated
# with each other, indicating that there is some pattern or dependence among the errors.
model <- lm(NeoplasiaPrevalence ~ Oncogene , data=mydata)
durbinWatsonTest(model) #Durbin-Watson test using car package
# Check for heteroscedasticity (Breusch-Pagan test)
# Checks if the variance of the errors is the same for every value of x (homoscedasticity). 
# Look for a random scatter of points around zero, with no discernible pattern or trend. 
# If the spread of residuals appears to vary systematically as the predicted
# values or predictor variable(s) change, it suggests potential heteroscedasticity.
plot(NOncogene, which = 1)  # Residuals vs. Fitted value plot
plot(NOncogene, which = 3)  # Scale-Location plot (square root of standardized residuals vs. Fitted values)
bptest(lm(NOncogene$residuals ~ fitted(NOncogene)))
### AIC Model Comparison
# Load the data and prepare comparative data object for AIC comparisons
# comparative_data <- comparative.data(phy = tre, data = mydata, 
#                                      names.col = "species", 
#                                      vcv = TRUE, na.omit = FALSE)
# Lambda fixed at 1 (full phylogenetic signal)
model_lambda_1 <- pgls(NeoplasiaPrevalence ~ Oncogene, data = comparative_data, lambda = 1.00)
# Compare AIC values
cat("AIC with Lambda = 0:", round(AIC(model),2), "\n")
cat("AIC with Estimated Lambda:", round(AIC(NOncogene),2), "\n")
cat("AIC with Lambda = 1:", round(AIC(model_lambda_1),2), "\n")
#******************************************
# Quadratic form to test for non-linearity
#******************************************
# mydata$Oncogene2 <- mydata$Oncogene^2
# NOncogene_quad <- pglsSEyPagel(NeoplasiaPrevalence ~ Oncogene + Oncogene2, data = mydata, tree=tre, se=SE, method = "ML")
# summary(NOncogene_quad)
# anova(NOncogene_quad)
# plot(NOncogene_quad, which = 1)
#*******************************


### Neoplasia vs TSG
NTSG <- pglsSEyPagel(NeoplasiaPrevalence ~ TSG, data = mydata, tree=tre, se=SE, method = "ML")
summary(NTSG)
anova(NTSG)
# Grab likelihood-based R2, Lambda, and p values for plotting
RelNTSG <- R2_lik(NTSG)
RelNTSG <- format(RelNTSG)
RelNTSG <- signif(as.numeric(RelNTSG), digits = 2)
RelNTSG
lambdaNTSG <- summary(NTSG)$modelStruct$corStruct
lambdaNTSG <- signif(lambdaNTSG[1], digits = 2)
lambdaNTSG
NTSG_pvalue <- summary(NTSG)$tTable
NTSG_pvalue <- signif(NTSG_pvalue[2,4], digits = 2)
NTSG_pvalue
# Check for normality of residuals 
ols_test_normality(NTSG$residuals)
hist(NTSG$residuals, main = "NTSG Residual Histogram")
qqnorm(NTSG$residuals)
qqline(NTSG$residuals)
# Check for independence of errors (autocorrelation) 
# Autocorrelation occurs when the residuals of a regression model are correlated
# with each other, indicating that there is some pattern or dependence among the errors.
model <- lm(NeoplasiaPrevalence ~ TSG , data=mydata)
durbinWatsonTest(model) #Durbin-Watson test using car package
# Check for heteroscedasticity (Breusch-Pagan test)
# Checks if the variance of the errors is the same for every value of x (homoscedasticity). 
# Look for a random scatter of points around zero, with no discernible pattern or trend. 
# If the spread of residuals appears to vary systematically as the predicted
# values or predictor variable(s) change, it suggests potential heteroscedasticity.
plot(NTSG, which = 1)  # Residuals vs. Fitted value plot
plot(NTSG, which = 3)  # Scale-Location plot (square root of standardized residuals vs. Fitted values)
bptest(lm(NTSG$residuals ~ fitted(NTSG)))
### AIC Model Comparison
# Load the data and prepare comparative data object for AIC comparisons
# comparative_data <- comparative.data(phy = tre, data = mydata, 
#                                      names.col = "species", 
#                                      vcv = TRUE, na.omit = FALSE)
# Lambda fixed at 1 (full phylogenetic signal)
model_lambda_1 <- pgls(NeoplasiaPrevalence ~ TSG, data = comparative_data, lambda = 1.00)
# Compare AIC values
cat("AIC with Lambda = 0:", round(AIC(model),2), "\n")
cat("AIC with Estimated Lambda:", round(AIC(NTSG),2), "\n")
cat("AIC with Lambda = 1:", round(AIC(model_lambda_1),2), "\n")
#******************************************
# Quadratic form to test for non-linearity
#******************************************
# mydata$TSG2 <- mydata$TSG^2
# NTSG_quad <- pglsSEyPagel(NeoplasiaPrevalence ~ TSG + TSG2, data = mydata, tree=tre, se=SE, method = "ML")
# summary(NTSG_quad)
# anova(NTSG_quad)
# plot(NTSG_quad, which = 1)
#*******************************


### Neoplasia vs Fusion
NFusion <- pglsSEyPagel(NeoplasiaPrevalence ~ Fusion, data = mydata, tree=tre, se=SE, method = "ML")
summary(NFusion)
anova(NFusion)
# Grab likelihood-based R2, Lambda, and p values for plotting
RelNFusion <- R2_lik(NFusion)
RelNFusion <- format(RelNFusion)
RelNFusion <- signif(as.numeric(RelNFusion), digits = 2)
RelNFusion
lambdaNFusion <- summary(NFusion)$modelStruct$corStruct
lambdaNFusion <- signif(lambdaNFusion[1], digits = 2)
lambdaNFusion
NFusion_pvalue <- summary(NFusion)$tTable
NFusion_pvalue <- signif(NFusion_pvalue[2,4], digits = 2)
NFusion_pvalue
# Check for normality of residuals 
ols_test_normality(NFusion$residuals)
hist(NFusion$residuals, main = "NFusion Residual Histogram")
qqnorm(NFusion$residuals)
qqline(NFusion$residuals)
# Check for independence of errors (autocorrelation) 
# Autocorrelation occurs when the residuals of a regression model are correlated
# with each other, indicating that there is some pattern or dependence among the errors.
model <- lm(NeoplasiaPrevalence ~ Fusion , data=mydata)
durbinWatsonTest(model) #Durbin-Watson test using car package
# Check for heteroscedasticity (Breusch-Pagan test)
# Checks if the variance of the errors is the same for every value of x (homoscedasticity). 
# Look for a random scatter of points around zero, with no discernible pattern or trend. 
# If the spread of residuals appears to vary systematically as the predicted
# values or predictor variable(s) change, it suggests potential heteroscedasticity.
plot(NFusion, which = 1)  # Residuals vs. Fitted value plot
plot(NFusion, which = 3)  # Scale-Location plot (square root of standardized residuals vs. Fitted values)
bptest(lm(NFusion$residuals ~ fitted(NFusion)))
### AIC Model Comparison
# Load the data and prepare comparative data object for AIC comparisons
# comparative_data <- comparative.data(phy = tre, data = mydata, 
#                                      names.col = "species", 
#                                      vcv = TRUE, na.omit = FALSE)
# Lambda fixed at 1 (full phylogenetic signal)
model_lambda_1 <- pgls(NeoplasiaPrevalence ~ Fusion, data = comparative_data, lambda = 1.00)
# Compare AIC values
cat("AIC with Lambda = 0:", round(AIC(model),2), "\n")
cat("AIC with Estimated Lambda:", round(AIC(NFusion),2), "\n")
cat("AIC with Lambda = 1:", round(AIC(model_lambda_1),2), "\n")
#******************************************
# Quadratic form to test for non-linearity
#******************************************
# mydata$Fusion2 <- mydata$Fusion^2
# NFusion_quad <- pglsSEyPagel(NeoplasiaPrevalence ~ Fusion + Fusion2, data = mydata, tree=tre, se=SE, method = "ML")
# summary(NFusion_quad)
# anova(NFusion_quad)
# plot(NFusion_quad, which = 1)
#*******************************


################################
### Malignancy vs Total_CGOs
################################
MTotal_CGOs <- pglsSEyPagel(MalignancyPrevalence ~ Total_CGOs, data = mydata, tree=tre, se=SE, method = "ML")
summary(MTotal_CGOs)
anova(MTotal_CGOs)
# Grab likelihood-based R2, Lambda, and p values for plotting
RelMTotal_CGOs <- R2_lik(MTotal_CGOs)
RelMTotal_CGOs <- format(RelMTotal_CGOs)
RelMTotal_CGOs <- signif(as.numeric(RelMTotal_CGOs), digits = 2)
RelMTotal_CGOs
lambdaMTotal_CGOs <- summary(MTotal_CGOs)$modelStruct$corStruct
lambdaMTotal_CGOs <- signif(lambdaMTotal_CGOs[1], digits = 2)
lambdaMTotal_CGOs
MTotal_CGOs_pvalue <- summary(MTotal_CGOs)$tTable
MTotal_CGOs_pvalue <- signif(MTotal_CGOs_pvalue[2,4], digits = 2)
MTotal_CGOs_pvalue
# Check for normality of residuals 
ols_test_normality(MTotal_CGOs$residuals)
hist(MTotal_CGOs$residuals, main = "MTotal_CGOs Residual Histogram")
qqnorm(MTotal_CGOs$residuals)
qqline(MTotal_CGOs$residuals)
# Check for independence of errors (autocorrelation) 
# Autocorrelation occurs when the residuals of a regression model are correlated
# with each other, indicating that there is some pattern or dependence among the errors.
model <- lm(MalignancyPrevalence ~ Total_CGOs , data=mydata)
durbinWatsonTest(model) #Durbin-Watson test using car package
# Check for heteroscedasticity (Breusch-Pagan test)
# Checks if the variance of the errors is the same for every value of x (homoscedasticity). 
# Look for a random scatter of points around zero, with no discernible pattern or trend. 
# If the spread of residuals appears to vary systematically as the predicted
# values or predictor variable(s) change, it suggests potential heteroscedasticity.
plot(MTotal_CGOs, which = 1)  # Residuals vs. Fitted value plot
plot(MTotal_CGOs, which = 3)  # Scale-Location plot (square root of standardized residuals vs. Fitted values)
bptest(lm(MTotal_CGOs$residuals ~ fitted(MTotal_CGOs)))
### AIC Model Comparison
# Load the data and prepare comparative data object for AIC comparisons
# comparative_data <- comparative.data(phy = tre, data = mydata, 
#                                      names.col = "species", 
#                                      vcv = TRUE, na.omit = FALSE)
# Lambda fixed at 1 (full phylogenetic signal)
model_lambda_1 <- pgls(MalignancyPrevalence ~ Total_CGOs, data = comparative_data, lambda = 1.00)
# Compare AIC values
cat("AIC with Lambda = 0:", round(AIC(model),2), "\n")
cat("AIC with Estimated Lambda:", round(AIC(NTotal_CGOs),2), "\n")
cat("AIC with Lambda = 1:", round(AIC(model_lambda_1),2), "\n")
#******************************************
# Quadratic form to test for non-linearity
#******************************************
mydata$Total_CGOs2 <- mydata$Total_CGOs^2
MTotal_CGOs_quad <- pglsSEyPagel(MalignancyPrevalence ~ Total_CGOs + Total_CGOs2, data = mydata, tree=tre, se=SE, method = "ML")
summary(MTotal_CGOs_quad)
anova(MTotal_CGOs_quad)
plot(MTotal_CGOs_quad, which = 1)
#*******************************


### Malignancy vs Somatic
MSomatic <- pglsSEyPagel(MalignancyPrevalence ~ Somatic, data = mydata, tree=tre, se=SE, method = "ML")
summary(MSomatic)
anova(MSomatic)
# Grab likelihood-based R2, Lambda, and p values for plotting
RelMSomatic <- R2_lik(MSomatic)
RelMSomatic <- format(RelMSomatic)
RelMSomatic <- signif(as.numeric(RelMSomatic), digits = 2)
RelMSomatic
lambdaMSomatic <- summary(MSomatic)$modelStruct$corStruct
lambdaMSomatic <- signif(lambdaMSomatic[1], digits = 2)
lambdaMSomatic
MSomatic_pvalue <- summary(MSomatic)$tTable
MSomatic_pvalue <- signif(MSomatic_pvalue[2,4], digits = 2)
MSomatic_pvalue
# Check for normality of residuals 
ols_test_normality(MSomatic$residuals)
hist(MSomatic$residuals, main = "MSomatic Residual Histogram")
qqnorm(MSomatic$residuals)
qqline(MSomatic$residuals)
# Check for independence of errors (autocorrelation) 
# Autocorrelation occurs when the residuals of a regression model are correlated
# with each other, indicating that there is some pattern or dependence among the errors.
model <- lm(MalignancyPrevalence ~ Somatic , data=mydata)
durbinWatsonTest(model) #Durbin-Watson test using car package
# Check for heteroscedasticity (Breusch-Pagan test)
# Checks if the variance of the errors is the same for every value of x (homoscedasticity). 
# Look for a random scatter of points around zero, with no discernible pattern or trend. 
# If the spread of residuals appears to vary systematically as the predicted
# values or predictor variable(s) change, it suggests potential heteroscedasticity.
plot(MSomatic, which = 1)  # Residuals vs. Fitted value plot
plot(MSomatic, which = 3)  # Scale-Location plot (square root of standardized residuals vs. Fitted values)
bptest(lm(MSomatic$residuals ~ fitted(MSomatic)))
### AIC Model Comparison
# Load the data and prepare comparative data object for AIC comparisons
# comparative_data <- comparative.data(phy = tre, data = mydata, 
#                                      names.col = "species", 
#                                      vcv = TRUE, na.omit = FALSE)
# Lambda fixed at 1 (full phylogenetic signal)
model_lambda_1 <- pgls(MalignancyPrevalence ~ Somatic, data = comparative_data, lambda = 1.00)
# Compare AIC values
cat("AIC with Lambda = 0:", round(AIC(model),2), "\n")
cat("AIC with Estimated Lambda:", round(AIC(MSomatic),2), "\n")
cat("AIC with Lambda = 1:", round(AIC(model_lambda_1),2), "\n")
#******************************************
# Quadratic form to test for non-linearity
#******************************************
mydata$Somatic2 <- mydata$Somatic^2
MSomatic_quad <- pglsSEyPagel(MalignancyPrevalence ~ Somatic + Somatic2, data = mydata, tree=tre, se=SE, method = "ML")
summary(MSomatic_quad)
anova(MSomatic_quad)
plot(MSomatic_quad, which = 1)
#*******************************

### Malignancy vs Germline
MGermline <- pglsSEyPagel(MalignancyPrevalence ~ Germline, data = mydata, tree=tre, se=SE, method = "ML")
summary(MGermline)
anova(MGermline)
# Grab likelihood-based R2, Lambda, and p values for plotting
RelMGermline <- R2_lik(MGermline)
RelMGermline <- format(RelMGermline)
RelMGermline <- signif(as.numeric(RelMGermline), digits = 2)
RelMGermline
lambdaMGermline <- summary(MGermline)$modelStruct$corStruct
lambdaMGermline <- signif(lambdaMGermline[1], digits = 2)
lambdaMGermline
MGermline_pvalue <- summary(MGermline)$tTable
MGermline_pvalue <- signif(MGermline_pvalue[2,4], digits = 2)
MGermline_pvalue
# Check for normality of residuals 
ols_test_normali
ty(MGermline$residuals)
hist(MGermline$residuals, main = "MGermline Residual Histogram")
qqnorm(MGermline$residuals)
qqline(MGermline$residuals)
# Check for independence of errors (autocorrelation) 
# Autocorrelation occurs when the residuals of a regression model are correlated
# with each other, indicating that there is some pattern or dependence among the errors.
model <- lm(MalignancyPrevalence ~ Germline , data=mydata)
durbinWatsonTest(model) #Durbin-Watson test using car package
# Check for heteroscedasticity (Breusch-Pagan test)
# Checks if the variance of the errors is the same for every value of x (homoscedasticity). 
# Look for a random scatter of points around zero, with no discernible pattern or trend. 
# If the spread of residuals appears to vary systematically as the predicted
# values or predictor variable(s) change, it suggests potential heteroscedasticity.
plot(MGermline, which = 1)  # Residuals vs. Fitted value plot
plot(MGermline, which = 3)  # Scale-Location plot (square root of standardized residuals vs. Fitted values)
bptest(lm(MGermline$residuals ~ fitted(MGermline)))
### AIC Model Comparison
# Load the data and prepare comparative data object for AIC comparisons
# comparative_data <- comparative.data(phy = tre, data = mydata, 
#                                      names.col = "species", 
#                                      vcv = TRUE, na.omit = FALSE)
# Lambda fixed at 1 (full phylogenetic signal)
model_lambda_1 <- pgls(MalignancyPrevalence ~ Germline, data = comparative_data, lambda = 1.00)
# Compare AIC values
cat("AIC with Lambda = 0:", round(AIC(model),2), "\n")
cat("AIC with Estimated Lambda:", round(AIC(MGermline),2), "\n")
cat("AIC with Lambda = 1:", round(AIC(model_lambda_1),2), "\n")
#******************************************
# Quadratic form to test for non-linearity
#******************************************
# mydata$Germline2 <- mydata$Germline^2
# MGermline2_quad <- pglsSEyPagel(MalignancyPrevalence ~ Germline + Germline2, data = mydata, tree=tre, se=SE, method = "ML")
# summary(MGermline2_quad)
# anova(MGermline2_quad)
# plot(MGermline2_quad, which = 1)
#*******************************

### Malignancy vs Oncogene
MOncogene <- pglsSEyPagel(MalignancyPrevalence ~ Oncogene, data = mydata, tree=tre, se=SE, method = "ML")
summary(MOncogene)
anova(MOncogene)
# Grab likelihood-based R2, Lambda, and p values for plotting
RelMOncogene <- R2_lik(MOncogene)
RelMOncogene <- format(RelMOncogene)
RelMOncogene <- signif(as.numeric(RelMOncogene), digits = 2)
RelMOncogene
lambdaMOncogene <- summary(MOncogene)$modelStruct$corStruct
lambdaMOncogene <- signif(lambdaMOncogene[1], digits = 2)
lambdaMOncogene
MOncogene_pvalue <- summary(MOncogene)$tTable
MOncogene_pvalue <- signif(MOncogene_pvalue[2,4], digits = 2)
MOncogene_pvalue
# Check for normality of residuals 
ols_test_normality(MOncogene$residuals)
hist(MOncogene$residuals, main = "MOncogene Residual Histogram")
qqnorm(MOncogene$residuals)
qqline(MOncogene$residuals)
# Check for independence of errors (autocorrelation) 
# Autocorrelation occurs when the residuals of a regression model are correlated
# with each other, indicating that there is some pattern or dependence among the errors.
model <- lm(MalignancyPrevalence ~ Oncogene , data=mydata)
durbinWatsonTest(model) #Durbin-Watson test using car package
# Check for heteroscedasticity (Breusch-Pagan test)
# Checks if the variance of the errors is the same for every value of x (homoscedasticity). 
# Look for a random scatter of points around zero, with no discernible pattern or trend. 
# If the spread of residuals appears to vary systematically as the predicted
# values or predictor variable(s) change, it suggests potential heteroscedasticity.
plot(MOncogene, which = 1)  # Residuals vs. Fitted value plot
plot(MOncogene, which = 3)  # Scale-Location plot (square root of standardized residuals vs. Fitted values)
bptest(lm(MOncogene$residuals ~ fitted(MOncogene)))
### AIC Model Comparison
# Load the data and prepare comparative data object for AIC comparisons
# comparative_data <- comparative.data(phy = tre, data = mydata, 
#                                      names.col = "species", 
#                                      vcv = TRUE, na.omit = FALSE)
# Lambda fixed at 1 (full phylogenetic signal)
model_lambda_1 <- pgls(MalignancyPrevalence ~ Oncogene, data = comparative_data, lambda = 1.00)
# Compare AIC values
cat("AIC with Lambda = 0:", round(AIC(model),2), "\n")
cat("AIC with Estimated Lambda:", round(AIC(MOncogene),2), "\n")
cat("AIC with Lambda = 1:", round(AIC(model_lambda_1),2), "\n")
#******************************************
# Quadratic form to test for non-linearity
#******************************************
# mydata$Oncogene2 <- mydata$Oncogene^2
# MOncogene2_quad <- pglsSEyPagel(MalignancyPrevalence ~ Oncogene + Oncogene2, data = mydata, tree=tre, se=SE, method = "ML")
# summary(MOncogene2_quad)
# anova(MOncogene2_quad)
# plot(MOncogene2_quad, which = 1)
#*******************************

### Malignancy vs TSG
MTSG <- pglsSEyPagel(MalignancyPrevalence ~ TSG, data = mydata, tree=tre, se=SE, method = "ML")
summary(MTSG)
anova(MTSG)
# Grab likelihood-based R2, Lambda, and p values for plotting
RelMTSG <- R2_lik(MTSG)
RelMTSG <- format(RelMTSG)
RelMTSG <- signif(as.numeric(RelMTSG), digits = 2)
RelMTSG
lambdaMTSG <- summary(MTSG)$modelStruct$corStruct
lambdaMTSG <- signif(lambdaMTSG[1], digits = 2)
lambdaMTSG
MTSG_pvalue <- summary(MTSG)$tTable
MTSG_pvalue <- signif(MTSG_pvalue[2,4], digits = 2)
MTSG_pvalue
# Check for normality of residuals 
ols_test_normality(MTSG$residuals)
hist(MTSG$residuals, main = "MTSG Residual Histogram")
qqnorm(MTSG$residuals)
qqline(MTSG$residuals)
# Check for independence of errors (autocorrelation) 
# Autocorrelation occurs when the residuals of a regression model are correlated
# with each other, indicating that there is some pattern or dependence among the errors.
model <- lm(MalignancyPrevalence ~ TSG , data=mydata)
durbinWatsonTest(model) #Durbin-Watson test using car package
# Check for heteroscedasticity (Breusch-Pagan test)
# Checks if the variance of the errors is the same for every value of x (homoscedasticity). 
# Look for a random scatter of points around zero, with no discernible pattern or trend. 
# If the spread of residuals appears to vary systematically as the predicted
# values or predictor variable(s) change, it suggests potential heteroscedasticity.
plot(MTSG, which = 1)  # Residuals vs. Fitted value plot
plot(MTSG, which = 3)  # Scale-Location plot (square root of standardized residuals vs. Fitted values)
bptest(lm(MTSG$residuals ~ fitted(MTSG)))
### AIC Model Comparison
# Load the data and prepare comparative data object for AIC comparisons
# comparative_data <- comparative.data(phy = tre, data = mydata, 
#                                      names.col = "species", 
#                                      vcv = TRUE, na.omit = FALSE)
# Lambda fixed at 1 (full phylogenetic signal)
model_lambda_1 <- pgls(MalignancyPrevalence ~ TSG, data = comparative_data, lambda = 1.00)
# Compare AIC values
cat("AIC with Lambda = 0:", round(AIC(model),2), "\n")
cat("AIC with Estimated Lambda:", round(AIC(MTSG),2), "\n")
cat("AIC with Lambda = 1:", round(AIC(model_lambda_1),2), "\n")
#******************************************
# Quadratic form to test for non-linearity
#******************************************
# mydata$TSG2 <- mydata$TSG^2
# MTSG2_quad <- pglsSEyPagel(MalignancyPrevalence ~ TSG + TSG2, data = mydata, tree=tre, se=SE, method = "ML")
# summary(MTSG2_quad)
# anova(MTSG2_quad)
# plot(MTSG2_quad, which = 1)
#*******************************

### Malignancy vs Fusion
MFusion <- pglsSEyPagel(MalignancyPrevalence ~ Fusion, data = mydata, tree=tre, se=SE, method = "ML")
summary(MFusion)
anova(MFusion)
# Grab likelihood-based R2, Lambda, and p values for plotting
RelMFusion <- R2_lik(MFusion)
RelMFusion <- format(RelMFusion)
RelMFusion <- signif(as.numeric(RelMFusion), digits = 2)
RelMFusion
lambdaMFusion <- summary(MFusion)$modelStruct$corStruct
lambdaMFusion <- signif(lambdaMFusion[1], digits = 2)
lambdaMFusion
MFusion_pvalue <- summary(MFusion)$tTable
MFusion_pvalue <- signif(MFusion_pvalue[2,4], digits = 2)
MFusion_pvalue
# Check for normality of residuals 
ols_test_normality(MFusion$residuals)
hist(MFusion$residuals, main = "MFusion Residual Histogram")
qqnorm(MFusion$residuals)
qqline(MFusion$residuals)
# Check for independence of errors (autocorrelation) 
# Autocorrelation occurs when the residuals of a regression model are correlated
# with each other, indicating that there is some pattern or dependence among the errors.
model <- lm(MalignancyPrevalence ~ Fusion , data=mydata)
durbinWatsonTest(model) #Durbin-Watson test using car package
# Check for heteroscedasticity (Breusch-Pagan test)
# Checks if the variance of the errors is the same for every value of x (homoscedasticity). 
# Look for a random scatter of points around zero, with no discernible pattern or trend. 
# If the spread of residuals appears to vary systematically as the predicted
# values or predictor variable(s) change, it suggests potential heteroscedasticity.
plot(MFusion, which = 1)  # Residuals vs. Fitted value plot
plot(MFusion, which = 3)  # Scale-Location plot (square root of standardized residuals vs. Fitted values)
bptest(lm(MFusion$residuals ~ fitted(MFusion)))
### AIC Model Comparison
# Load the data and prepare comparative data object for AIC comparisons
# comparative_data <- comparative.data(phy = tre, data = mydata, 
#                                      names.col = "species", 
#                                      vcv = TRUE, na.omit = FALSE)
# Lambda fixed at 1 (full phylogenetic signal)
model_lambda_1 <- pgls(MalignancyPrevalence ~ Fusion, data = comparative_data, lambda = 1.00)
# Compare AIC values
cat("AIC with Lambda = 0:", round(AIC(model),2), "\n")
cat("AIC with Estimated Lambda:", round(AIC(MFusion),2), "\n")
cat("AIC with Lambda = 1:", round(AIC(model_lambda_1),2), "\n")
#******************************************
# Quadratic form to test for non-linearity
#******************************************
# mydata$Fusion2 <- mydata$Fusion^2
# MFusion2_quad <- pglsSEyPagel(MalignancyPrevalence ~ Fusion + Fusion2, data = mydata, tree=tre, se=SE, method = "ML")
# summary(MFusion2_quad)
# anova(MFusion2_quad)
# plot(MFusion2_quad, which = 1)
#*******************************


########### Multiple PGLS Controlling for Longevity 
########### Neoplasia or Malignancy ~ Number of CGOs of Various Types + Longevity + e 

hist(mydata$maximum_longevity_m)

########### Check for the potential outliers in Longevity data
grubbs_result <- grubbs.test(mydata$maximum_longevity_m)
print(grubbs_result)

# We did not remove two longevity influential points in our analyses. 
# See Methods for more detail ... 
# To remove two influential points (elephant and chimpanzee)
#mydata <- mydata[mydata$maximum_longevity_m != (max(mydata$maximum_longevity_m)), ]
#mydata <- mydata[mydata$maximum_longevity_m != (max(mydata$maximum_longevity_m)), ]
#nrow(mydata)

########### Pearson Correlation between Neoplasia/Malignancy and Longevity
cor(mydata$NeoplasiaPrevalence, mydata$maximum_longevity_m)
cor(mydata$MalignancyPrevalence, mydata$maximum_longevity_m)

# nLTR activities are also shown to be associated with longevity
# Test for any potential correlation 
#cor(mydata$maximum_longevity_m, mydata$Transformed_L1_Counts)
#cor(mydata$maximum_longevity_m, mydata$Transformed_L1_SINEs_Counts)


##########################################
### Neoplasia vs Total_CGOs + Longevity
##########################################
NTotal_CGOsLongevity <- pglsSEyPagel(NeoplasiaPrevalence ~ Total_CGOs + maximum_longevity_m, data = mydata, tree=tre, se=SE, method = "ML")
summary(NTotal_CGOsLongevity)
anova(NTotal_CGOsLongevity)
# FDR test and extracting the adj. p-values from the model
NTotal_CGOsLongevity_fdr_corrected <- p.adjust(summary(NTotal_CGOsLongevity)$tTable[, "p-value"], method = "fdr")
NTotal_CGOsLongevity_fdr_corrected
# VIF (variance inflation factor) to ensure multicollinearity is not an issue. 
# If VIF < 5, it's safe to include both variables in the model.
vif(NTotal_CGOsLongevity)
# Grab likelihood-based R2, Lambda, and p values 
RelNTotal_CGOsLongevity  <- R2_lik(NTotal_CGOsLongevity)
n <- nrow(mydata)
p <- 2  # Number of predictors (CGO and longevity)
adjRelNTotal_CGOsLongevity  <- signif(1 - ((1 - RelNTotal_CGOsLongevity) * (n - 1)) / (n - p - 1), 2)
adjRelNTotal_CGOsLongevity
# Check for normality of residuals 
ols_test_normality(NTotal_CGOsLongevity$residuals)
hist(NTotal_CGOsLongevity$residuals, main = "NTotal_CGOsLongevity Residual Histogram")
qqnorm(NTotal_CGOsLongevity$residuals)
qqline(NTotal_CGOsLongevity$residuals)
# Check for independence of errors (autocorrelation) 
# Autocorrelation occurs when the residuals of a regression model are correlated
# with each other, indicating that there is some pattern or dependence among the errors.
model <- lm(NeoplasiaPrevalence ~ Total_CGOs + maximum_longevity_m, data=mydata)
durbinWatsonTest(model) #Durbin-Watson test using car package
# Check for heteroscedasticity (Breusch-Pagan test)
# Checks if the variance of the errors is the same for every value of x (homoscedasticity). 
# Look for a random scatter of points around zero, with no discernible pattern or trend. 
# If the spread of residuals appears to vary systematically as the predicted
# values or predictor variable(s) change, it suggests potential heteroscedasticity.
plot(NTotal_CGOsLongevity, which = 1)  # Residuals vs. Fitted value plot
plot(NTotal_CGOsLongevity, which = 3)  # Scale-Location plot (square root of standardized residuals vs. Fitted values)
bptest(model)
### AIC Model Comparison
# Load the data and prepare comparative data object for AIC comparisons
# comparative_data <- comparative.data(phy = tre, data = mydata, 
#                                      names.col = "species", 
#                                      vcv = TRUE, na.omit = FALSE)
# Lambda fixed at 1 (full phylogenetic signal)
model_lambda_1 <- pgls(NeoplasiaPrevalence ~ Total_CGOs + maximum_longevity_m, data = comparative_data, lambda = 1.00)
# Compare AIC values
cat("AIC with Lambda = 0:", round(AIC(model),2), "\n")
cat("AIC with Estimated Lambda:", round(AIC(NTotal_CGOsLongevity),2), "\n")
cat("AIC with Lambda = 1:", round(AIC(model_lambda_1),2), "\n")
#******************************************
# Quadratic form to test for non-linearity
#******************************************
mydata$Total_CGOs2 <- mydata$Total_CGOs^2
NTotal_CGOsLongevity_quad <- pglsSEyPagel(NeoplasiaPrevalence ~ Total_CGOs + Total_CGOs2, data = mydata, tree=tre, se=SE, method = "ML")
summary(NTotal_CGOsLongevity_quad)
anova(NTotal_CGOsLongevity_quad)
plot(NTotal_CGOsLongevity_quad, which = 1)
#*******************************


### Neoplasia vs Somatic + Longevity
NSomaticLongevity <- pglsSEyPagel(NeoplasiaPrevalence ~ Somatic + maximum_longevity_m, data = mydata, tree=tre, se=SE, method = "ML")
summary(NSomaticLongevity)
anova(NSomaticLongevity)
# FDR test and extracting the adj. p-values from the model
NSomaticLongevity_fdr_corrected <- p.adjust(summary(NSomaticLongevity)$tTable[, "p-value"], method = "fdr")
NSomaticLongevity_fdr_corrected
# VIF (variance inflation factor) to ensure multicollinearity is not an issue. 
# If VIF < 5, it's safe to include both variables in the model.
vif(NSomaticLongevity)
# Grab likelihood-based R2, Lambda, and p values 
RelNSomaticLongevity  <- R2_lik(NSomaticLongevity)
n <- nrow(mydata)
p <- 2  # Number of predictors (CGO and longevity)
adjRelNSomaticLongevity  <- signif(1 - ((1 - RelNSomaticLongevity) * (n - 1)) / (n - p - 1), 2)
adjRelNSomaticLongevity
# Check for normality of residuals 
ols_test_normality(NSomaticLongevity$residuals)
hist(NSomaticLongevity$residuals, main = "NSomaticLongevity Residual Histogram")
qqnorm(NSomaticLongevity$residuals)
qqline(NSomaticLongevity$residuals)
# Check for independence of errors (autocorrelation) 
# Autocorrelation occurs when the residuals of a regression model are correlated
# with each other, indicating that there is some pattern or dependence among the errors.
model <- lm(NeoplasiaPrevalence ~ Somatic + maximum_longevity_m, data=mydata)
durbinWatsonTest(model) #Durbin-Watson test using car package
# Check for heteroscedasticity (Breusch-Pagan test)
# Checks if the variance of the errors is the same for every value of x (homoscedasticity). 
# Look for a random scatter of points around zero, with no discernible pattern or trend. 
# If the spread of residuals appears to vary systematically as the predicted
# values or predictor variable(s) change, it suggests potential heteroscedasticity.
plot(NSomaticLongevity, which = 1)  # Residuals vs. Fitted value plot
plot(NSomaticLongevity, which = 3)  # Scale-Location plot (square root of standardized residuals vs. Fitted values)
bptest(model)
### AIC Model Comparison
# Load the data and prepare comparative data object for AIC comparisons
# comparative_data <- comparative.data(phy = tre, data = mydata, 
#                                      names.col = "species", 
#                                      vcv = TRUE, na.omit = FALSE)
# Lambda fixed at 1 (full phylogenetic signal)
model_lambda_1 <- pgls(NeoplasiaPrevalence ~ Somatic + maximum_longevity_m, data = comparative_data, lambda = 1.00)
# Compare AIC values
cat("AIC with Lambda = 0:", round(AIC(model),2), "\n")
cat("AIC with Estimated Lambda:", round(AIC(NSomaticLongevity),2), "\n")
cat("AIC with Lambda = 1:", round(AIC(model_lambda_1),2), "\n")
#******************************************
# Quadratic form to test for non-linearity
#******************************************
mydata$Somatic2 <- mydata$Somatic^2
# NSomaticLongevity_quad <- pglsSEyPagel(NeoplasiaPrevalence ~ Somatic + Somatic2, data = mydata, tree=tre, se=SE, method = "ML")
# summary(NSomaticLongevity_quad)
# anova(NSomaticLongevity_quad)
# plot(NSomaticLongevity_quad, which = 1)
#*******************************

### Neoplasia vs Germline + Longevity
NGermlineLongevity <- pglsSEyPagel(NeoplasiaPrevalence ~ Germline + maximum_longevity_m, data = mydata, tree=tre, se=SE, method = "ML")
summary(NGermlineLongevity)
anova(NGermlineLongevity)
# FDR test and extracting the adj. p-values from the model
NGermlineLongevity_fdr_corrected <- p.adjust(summary(NGermlineLongevity)$tTable[, "p-value"], method = "fdr")
NGermlineLongevity_fdr_corrected
# VIF (variance inflation factor) to ensure multicollinearity is not an issue. 
# If VIF < 5, it's safe to include both variables in the model.
vif(NGermlineLongevity)
# Grab likelihood-based R2, Lambda, and p values 
RelNGermlineLongevity  <- R2_lik(NGermlineLongevity)
n <- nrow(mydata)
p <- 2  # Number of predictors (CGO and longevity)
adjRelNGermlineLongevity  <- signif(1 - ((1 - RelNGermlineLongevity) * (n - 1)) / (n - p - 1), 2)
adjRelNGermlineLongevity
# Check for normality of residuals 
ols_test_normality(NGermlineLongevity$residuals)
hist(NGermlineLongevity$residuals, main = "NGermlineLongevity Residual Histogram")
qqnorm(NGermlineLongevity$residuals)
qqline(NGermlineLongevity$residuals)
# Check for independence of errors (autocorrelation) 
# Autocorrelation occurs when the residuals of a regression model are correlated
# with each other, indicating that there is some pattern or dependence among the errors.
model <- lm(NeoplasiaPrevalence ~ Germline + maximum_longevity_m, data=mydata)
durbinWatsonTest(model) #Durbin-Watson test using car package
# Check for heteroscedasticity (Breusch-Pagan test)
# Checks if the variance of the errors is the same for every value of x (homoscedasticity). 
# Look for a random scatter of points around zero, with no discernible pattern or trend. 
# If the spread of residuals appears to vary systematically as the predicted
# values or predictor variable(s) change, it suggests potential heteroscedasticity.
plot(NGermlineLongevity, which = 1)  # Residuals vs. Fitted value plot
plot(NGermlineLongevity, which = 3)  # Scale-Location plot (square root of standardized residuals vs. Fitted values)
bptest(model)
### AIC Model Comparison
# Load the data and prepare comparative data object for AIC comparisons
# comparative_data <- comparative.data(phy = tre, data = mydata, 
#                                      names.col = "species", 
#                                      vcv = TRUE, na.omit = FALSE)
# Lambda fixed at 1 (full phylogenetic signal)
model_lambda_1 <- pgls(NeoplasiaPrevalence ~ Germline + maximum_longevity_m, data = comparative_data, lambda = 1.00)
# Compare AIC values
cat("AIC with Lambda = 0:", round(AIC(model),2), "\n")
cat("AIC with Estimated Lambda:", round(AIC(NGermlineLongevity),2), "\n")
cat("AIC with Lambda = 1:", round(AIC(model_lambda_1),2), "\n")
#******************************************
# Quadratic form to test for non-linearity
#******************************************
# mydata$Germline2 <- mydata$Germline^2
# NGermlineLongevity_quad <- pglsSEyPagel(NeoplasiaPrevalence ~ Germline + Germline2, data = mydata, tree=tre, se=SE, method = "ML")
# summary(NGermlineLongevity_quad)
# anova(NGermlineLongevity_quad)
# plot(NGermlineLongevity_quad, which = 1)
#*******************************

### Neoplasia vs Oncogene + Longevity
NOncogeneLongevity <- pglsSEyPagel(NeoplasiaPrevalence ~ Oncogene + maximum_longevity_m, data = mydata, tree=tre, se=SE, method = "ML")
summary(NOncogeneLongevity)
anova(NOncogeneLongevity)
# FDR test and extracting the adj. p-values from the model
NOncogeneLongevity_fdr_corrected <- p.adjust(summary(NOncogeneLongevity)$tTable[, "p-value"], method = "fdr")
NOncogeneLongevity_fdr_corrected
# VIF (variance inflation factor) to ensure multicollinearity is not an issue. 
# If VIF < 5, it's safe to include both variables in the model.
vif(NOncogeneLongevity)
# Grab likelihood-based R2, Lambda, and p values 
RelNOncogeneLongevity  <- R2_lik(NOncogeneLongevity)
n <- nrow(mydata)
p <- 2  # Number of predictors (CGO and longevity)
adjRelNOncogeneLongevity  <- signif(1 - ((1 - RelNOncogeneLongevity) * (n - 1)) / (n - p - 1), 2)
adjRelNOncogeneLongevity
# Check for normality of residuals 
ols_test_normality(NOncogeneLongevity$residuals)
hist(NOncogeneLongevity$residuals, main = "NOncogeneLongevity Residual Histogram")
qqnorm(NOncogeneLongevity$residuals)
qqline(NOncogeneLongevity$residuals)
# Check for independence of errors (autocorrelation) 
# Autocorrelation occurs when the residuals of a regression model are correlated
# with each other, indicating that there is some pattern or dependence among the errors.
model <- lm(NeoplasiaPrevalence ~ Oncogene + maximum_longevity_m, data=mydata)
durbinWatsonTest(model) #Durbin-Watson test using car package
# Check for heteroscedasticity (Breusch-Pagan test)
# Checks if the variance of the errors is the same for every value of x (homoscedasticity). 
# Look for a random scatter of points around zero, with no discernible pattern or trend. 
# If the spread of residuals appears to vary systematically as the predicted
# values or predictor variable(s) change, it suggests potential heteroscedasticity.
plot(NOncogeneLongevity, which = 1)  # Residuals vs. Fitted value plot
plot(NOncogeneLongevity, which = 3)  # Scale-Location plot (square root of standardized residuals vs. Fitted values)
bptest(model)
### AIC Model Comparison
# Load the data and prepare comparative data object for AIC comparisons
# comparative_data <- comparative.data(phy = tre, data = mydata, 
#                                      names.col = "species", 
#                                      vcv = TRUE, na.omit = FALSE)
# Lambda fixed at 1 (full phylogenetic signal)
model_lambda_1 <- pgls(NeoplasiaPrevalence ~ Oncogene + maximum_longevity_m, data = comparative_data, lambda = 1.00)
# Compare AIC values
cat("AIC with Lambda = 0:", round(AIC(model),2), "\n")
cat("AIC with Estimated Lambda:", round(AIC(NOncogeneLongevity),2), "\n")
cat("AIC with Lambda = 1:", round(AIC(model_lambda_1),2), "\n")
#******************************************
# Quadratic form to test for non-linearity
#******************************************
# mydata$Oncogene2 <- mydata$Oncogene^2
# NOncogeneLongevity_quad <- pglsSEyPagel(NeoplasiaPrevalence ~ Oncogene + Oncogene2, data = mydata, tree=tre, se=SE, method = "ML")
# summary(NOncogeneLongevity_quad)
# anova(NOncogeneLongevity_quad)
# plot(NOncogeneLongevity_quad, which = 1)
#*******************************

### Neoplasia vs TSG + Longevity
NTSGLongevity <- pglsSEyPagel(NeoplasiaPrevalence ~ TSG + maximum_longevity_m, data = mydata, tree=tre, se=SE, method = "ML")
summary(NTSGLongevity)
anova(NTSGLongevity)
# FDR test and extracting the adj. p-values from the model
NTSGLongevity_fdr_corrected <- p.adjust(summary(NTSGLongevity)$tTable[, "p-value"], method = "fdr")
NTSGLongevity_fdr_corrected
# VIF (variance inflation factor) to ensure multicollinearity is not an issue. 
# If VIF < 5, it's safe to include both variables in the model.
vif(NTSGLongevity)
# Grab likelihood-based R2, Lambda, and p values 
RelNTSGLongevity  <- R2_lik(NTSGLongevity)
n <- nrow(mydata)
p <- 2  # Number of predictors (CGO and longevity)
adjRelNTSGLongevity  <- signif(1 - ((1 - RelNTSGLongevity) * (n - 1)) / (n - p - 1), 2)
adjRelNTSGLongevity
# Check for normality of residuals 
ols_test_normality(NTSGLongevity$residuals)
hist(NTSGLongevity$residuals, main = "NTSGLongevity Residual Histogram")
qqnorm(NTSGLongevity$residuals)
qqline(NTSGLongevity$residuals)
# Check for independence of errors (autocorrelation) 
# Autocorrelation occurs when the residuals of a regression model are correlated
# with each other, indicating that there is some pattern or dependence among the errors.
model <- lm(NeoplasiaPrevalence ~ TSG + maximum_longevity_m, data=mydata)
durbinWatsonTest(model) #Durbin-Watson test using car package
# Check for heteroscedasticity (Breusch-Pagan test)
# Checks if the variance of the errors is the same for every value of x (homoscedasticity). 
# Look for a random scatter of points around zero, with no discernible pattern or trend. 
# If the spread of residuals appears to vary systematically as the predicted
# values or predictor variable(s) change, it suggests potential heteroscedasticity.
plot(NTSGLongevity, which = 1)  # Residuals vs. Fitted value plot
plot(NTSGLongevity, which = 3)  # Scale-Location plot (square root of standardized residuals vs. Fitted values)
bptest(model)
### AIC Model Comparison
# Load the data and prepare comparative data object for AIC comparisons
# comparative_data <- comparative.data(phy = tre, data = mydata, 
#                                      names.col = "species", 
#                                      vcv = TRUE, na.omit = FALSE)
# Lambda fixed at 1 (full phylogenetic signal)
model_lambda_1 <- pgls(NeoplasiaPrevalence ~ TSG + maximum_longevity_m, data = comparative_data, lambda = 1.00)
# Compare AIC values
cat("AIC with Lambda = 0:", round(AIC(model),2), "\n")
cat("AIC with Estimated Lambda:", round(AIC(NTSGLongevity),2), "\n")
cat("AIC with Lambda = 1:", round(AIC(model_lambda_1),2), "\n")
#******************************************
# Quadratic form to test for non-linearity
#******************************************
# mydata$TSG2 <- mydata$TSG^2
# NTSGLongevity_quad <- pglsSEyPagel(NeoplasiaPrevalence ~ TSG + TSG2, data = mydata, tree=tre, se=SE, method = "ML")
# summary(NTSGLongevity_quad)
# anova(NTSGLongevity_quad)
# plot(NTSGLongevity_quad, which = 1)
#*******************************

### Neoplasia vs Fusion + Longevity
NFusionLongevity <- pglsSEyPagel(NeoplasiaPrevalence ~ Fusion + maximum_longevity_m, data = mydata, tree=tre, se=SE, method = "ML")
summary(NFusionLongevity)
anova(NFusionLongevity)
# FDR test and extracting the adj. p-values from the model
NFusionLongevity_fdr_corrected <- p.adjust(summary(NFusionLongevity)$tTable[, "p-value"], method = "fdr")
NFusionLongevity_fdr_corrected
# VIF (variance inflation factor) to ensure multicollinearity is not an issue. 
# If VIF < 5, it's safe to include both variables in the model.
vif(NFusionLongevity)
# Grab likelihood-based R2, Lambda, and p values 
RelNFusionLongevity  <- R2_lik(NFusionLongevity)
n <- nrow(mydata)
p <- 2  # Number of predictors (CGO and longevity)
adjRelNFusionLongevity  <- signif(1 - ((1 - RelNFusionLongevity) * (n - 1)) / (n - p - 1), 2)
adjRelNFusionLongevity
# Check for normality of residuals 
ols_test_normality(NFusionLongevity$residuals)
hist(NFusionLongevity$residuals, main = "NFusionLongevity Residual Histogram")
qqnorm(NFusionLongevity$residuals)
qqline(NFusionLongevity$residuals)
# Check for independence of errors (autocorrelation) 
# Autocorrelation occurs when the residuals of a regression model are correlated
# with each other, indicating that there is some pattern or dependence among the errors.
model <- lm(NeoplasiaPrevalence ~ Fusion + maximum_longevity_m, data=mydata)
durbinWatsonTest(model) #Durbin-Watson test using car package
# Check for heteroscedasticity (Breusch-Pagan test)
# Checks if the variance of the errors is the same for every value of x (homoscedasticity). 
# Look for a random scatter of points around zero, with no discernible pattern or trend. 
# If the spread of residuals appears to vary systematically as the predicted
# values or predictor variable(s) change, it suggests potential heteroscedasticity.
plot(NFusionLongevity, which = 1)  # Residuals vs. Fitted value plot
plot(NFusionLongevity, which = 3)  # Scale-Location plot (square root of standardized residuals vs. Fitted values)
bptest(model)
### AIC Model Comparison
# Load the data and prepare comparative data object for AIC comparisons
# comparative_data <- comparative.data(phy = tre, data = mydata, 
#                                      names.col = "species", 
#                                      vcv = TRUE, na.omit = FALSE)
# Lambda fixed at 1 (full phylogenetic signal)
model_lambda_1 <- pgls(NeoplasiaPrevalence ~ Fusion + maximum_longevity_m, data = comparative_data, lambda = 1.00)
# Compare AIC values
cat("AIC with Lambda = 0:", round(AIC(model),2), "\n")
cat("AIC with Estimated Lambda:", round(AIC(NFusionLongevity),2), "\n")
cat("AIC with Lambda = 1:", round(AIC(model_lambda_1),2), "\n")
#******************************************
# Quadratic form to test for non-linearity
#******************************************
# mydata$Fusion2 <- mydata$Fusion^2
# NFusionLongevity_quad <- pglsSEyPagel(NeoplasiaPrevalence ~ Fusion + Fusion2, data = mydata, tree=tre, se=SE, method = "ML")
# summary(NFusionLongevity_quad)
# anova(NFusionLongevity_quad)
# plot(NFusionLongevity_quad, which = 1)
#*******************************

###########################################
### Malignancy vs Total_CGOs + Longevity
###########################################
MTotal_CGOsLongevity <- pglsSEyPagel(MalignancyPrevalence ~ Total_CGOs + maximum_longevity_m, data = mydata, tree=tre, se=SE, method = "ML")
summary(MTotal_CGOsLongevity)
anova(MTotal_CGOsLongevity)
# FDR test and extracting the adj. p-values from the model
MTotal_CGOsLongevity_fdr_corrected <- p.adjust(summary(MTotal_CGOsLongevity)$tTable[, "p-value"], method = "fdr")
MTotal_CGOsLongevity_fdr_corrected
# VIF (variance inflation factor) to ensure multicollinearity is not an issue. 
# If VIF < 5, it's safe to include both variables in the model.
vif(MTotal_CGOsLongevity)
# Grab likelihood-based R2, Lambda, and p values 
RelMTotal_CGOsLongevity  <- R2_lik(MTotal_CGOsLongevity)
n <- nrow(mydata)
p <- 2  # Number of predictors (CGO and longevity)
adjRelMTotal_CGOsLongevity  <- signif(1 - ((1 - RelMTotal_CGOsLongevity) * (n - 1)) / (n - p - 1), 2)
adjRelMTotal_CGOsLongevity
# Check for normality of residuals 
ols_test_normality(MTotal_CGOsLongevity$residuals)
hist(MTotal_CGOsLongevity$residuals, main = "MTotal_CGOsLongevity Residual Histogram")
qqnorm(MTotal_CGOsLongevity$residuals)
qqline(MTotal_CGOsLongevity$residuals)
# Check for independence of errors (autocorrelation) 
# Autocorrelation occurs when the residuals of a regression model are correlated
# with each other, indicating that there is some pattern or dependence among the errors.
model <- lm(MalignancyPrevalence ~ Total_CGOs + maximum_longevity_m, data=mydata)
durbinWatsonTest(model) #Durbin-Watson test using car package
# Check for heteroscedasticity (Breusch-Pagan test)
# Checks if the variance of the errors is the same for every value of x (homoscedasticity). 
# Look for a random scatter of points around zero, with no discernible pattern or trend. 
# If the spread of residuals appears to vary systematically as the predicted
# values or predictor variable(s) change, it suggests potential heteroscedasticity.
plot(MTotal_CGOsLongevity, which = 1)  # Residuals vs. Fitted value plot
plot(MTotal_CGOsLongevity, which = 3)  # Scale-Location plot (square root of standardized residuals vs. Fitted values)
bptest(model)
### AIC Model Comparison
# Load the data and prepare comparative data object for AIC comparisons
# comparative_data <- comparative.data(phy = tre, data = mydata, 
#                                      names.col = "species", 
#                                      vcv = TRUE, na.omit = FALSE)
# Lambda fixed at 1 (full phylogenetic signal)
model_lambda_1 <- pgls(MalignancyPrevalence ~ Total_CGOs + maximum_longevity_m, data = comparative_data, lambda = 1.00)
# Compare AIC values
cat("AIC with Lambda = 0:", round(AIC(model),2), "\n")
cat("AIC with Estimated Lambda:", round(AIC(MTotal_CGOsLongevity),2), "\n")
cat("AIC with Lambda = 1:", round(AIC(model_lambda_1),2), "\n")
#******************************************
# Quadratic form to test for non-linearity
#******************************************
# mydata$Total_CGOs2 <- mydata$Total_CGOs^2
# MTotal_CGOsLongevity_quad <- pglsSEyPagel(MalignancyPrevalence ~ Total_CGOs + Total_CGOs2, data = mydata, tree=tre, se=SE, method = "ML")
# summary(MTotal_CGOsLongevity_quad)
# anova(MTotal_CGOsLongevity_quad)
# plot(MTotal_CGOsLongevity_quad, which = 1)
#*******************************

### Malignancy vs Somatic + Longevity
MSomaticLongevity <- pglsSEyPagel(MalignancyPrevalence ~ Somatic + maximum_longevity_m, data = mydata, tree=tre, se=SE, method = "ML")
summary(MSomaticLongevity)
anova(MSomaticLongevity)
# FDR test and extracting the adj. p-values from the model
MSomaticLongevity_fdr_corrected <- p.adjust(summary(MSomaticLongevity)$tTable[, "p-value"], method = "fdr")
MSomaticLongevity_fdr_corrected
# VIF (variance inflation factor) to ensure multicollinearity is not an issue. 
# If VIF < 5, it's safe to include both variables in the model.
vif(MSomaticLongevity)
# Grab likelihood-based R2, Lambda, and p values 
RelMSomaticLongevity  <- R2_lik(MSomaticLongevity)
n <- nrow(mydata)
p <- 2  # Number of predictors (CGO and longevity)
adjRelMSomaticLongevity  <- signif(1 - ((1 - RelMSomaticLongevity) * (n - 1)) / (n - p - 1), 2)
adjRelMSomaticLongevity
# Check for normality of residuals 
ols_test_normality(MSomaticLongevity$residuals)
hist(MSomaticLongevity$residuals, main = "MSomaticLongevity Residual Histogram")
qqnorm(MSomaticLongevity$residuals)
qqline(MSomaticLongevity$residuals)
# Check for independence of errors (autocorrelation) 
# Autocorrelation occurs when the residuals of a regression model are correlated
# with each other, indicating that there is some pattern or dependence among the errors.
model <- lm(MalignancyPrevalence ~ Somatic + maximum_longevity_m, data=mydata)
durbinWatsonTest(model) #Durbin-Watson test using car package
# Check for heteroscedasticity (Breusch-Pagan test)
# Checks if the variance of the errors is the same for every value of x (homoscedasticity). 
# Look for a random scatter of points around zero, with no discernible pattern or trend. 
# If the spread of residuals appears to vary systematically as the predicted
# values or predictor variable(s) change, it suggests potential heteroscedasticity.
plot(MSomaticLongevity, which = 1)  # Residuals vs. Fitted value plot
plot(MSomaticLongevity, which = 3)  # Scale-Location plot (square root of standardized residuals vs. Fitted values)
bptest(model)
### AIC Model Comparison
# Load the data and prepare comparative data object for AIC comparisons
# comparative_data <- comparative.data(phy = tre, data = mydata, 
#                                      names.col = "species", 
#                                      vcv = TRUE, na.omit = FALSE)
# Lambda fixed at 1 (full phylogenetic signal)
model_lambda_1 <- pgls(MalignancyPrevalence ~ Somatic + maximum_longevity_m, data = comparative_data, lambda = 1.00)
# Compare AIC values
cat("AIC with Lambda = 0:", round(AIC(model),2), "\n")
cat("AIC with Estimated Lambda:", round(AIC(MSomaticLongevity),2), "\n")
cat("AIC with Lambda = 1:", round(AIC(model_lambda_1),2), "\n")
#******************************************
# Quadratic form to test for non-linearity
#******************************************
# mydata$Somatic2 <- mydata$Somatic^2
# MSomaticLongevity_quad <- pglsSEyPagel(MalignancyPrevalence ~ Somatic + Somatic2, data = mydata, tree=tre, se=SE, method = "ML")
# summary(MSomaticLongevity_quad)
# anova(MSomaticLongevity_quad)
# plot(MSomaticLongevity_quad, which = 1)
#*******************************

### Malignancy vs Germline + Longevity
MGermlineLongevity <- pglsSEyPagel(MalignancyPrevalence ~ Germline + maximum_longevity_m, data = mydata, tree=tre, se=SE, method = "ML")
summary(MGermlineLongevity)
anova(MGermlineLongevity)
# FDR test and extracting the adj. p-values from the model
MGermlineLongevity_fdr_corrected <- p.adjust(summary(MGermlineLongevity)$tTable[, "p-value"], method = "fdr")
MGermlineLongevity_fdr_corrected
# VIF (variance inflation factor) to ensure multicollinearity is not an issue. 
# If VIF < 5, it's safe to include both variables in the model.
vif(MGermlineLongevity)
# Grab likelihood-based R2, Lambda, and p values 
RelMGermlineLongevity  <- R2_lik(MGermlineLongevity)
n <- nrow(mydata)
p <- 2  # Number of predictors (CGO and longevity)
adjRelMGermlineLongevity  <- signif(1 - ((1 - RelMGermlineLongevity) * (n - 1)) / (n - p - 1), 2)
adjRelMGermlineLongevity
# Check for normality of residuals 
ols_test_normality(MGermlineLongevity$residuals)
hist(MGermlineLongevity$residuals, main = "MGermlineLongevity Residual Histogram")
qqnorm(MGermlineLongevity$residuals)
qqline(MGermlineLongevity$residuals)
# Check for independence of errors (autocorrelation) 
# Autocorrelation occurs when the residuals of a regression model are correlated
# with each other, indicating that there is some pattern or dependence among the errors.
model <- lm(MalignancyPrevalence ~ Germline + maximum_longevity_m, data=mydata)
durbinWatsonTest(model) #Durbin-Watson test using car package
# Check for heteroscedasticity (Breusch-Pagan test)
# Checks if the variance of the errors is the same for every value of x (homoscedasticity). 
# Look for a random scatter of points around zero, with no discernible pattern or trend. 
# If the spread of residuals appears to vary systematically as the predicted
# values or predictor variable(s) change, it suggests potential heteroscedasticity.
plot(MGermlineLongevity, which = 1)  # Residuals vs. Fitted value plot
plot(MGermlineLongevity, which = 3)  # Scale-Location plot (square root of standardized residuals vs. Fitted values)
bptest(model)
### AIC Model Comparison
# Load the data and prepare comparative data object for AIC comparisons
# comparative_data <- comparative.data(phy = tre, data = mydata, 
#                                      names.col = "species", 
#                                      vcv = TRUE, na.omit = FALSE)
# Lambda fixed at 1 (full phylogenetic signal)
model_lambda_1 <- pgls(MalignancyPrevalence ~ Germline + maximum_longevity_m, data = comparative_data, lambda = 1.00)
# Compare AIC values
cat("AIC with Lambda = 0:", round(AIC(model),2), "\n")
cat("AIC with Estimated Lambda:", round(AIC(MGermlineLongevity),2), "\n")
cat("AIC with Lambda = 1:", round(AIC(model_lambda_1),2), "\n")
#******************************************
# Quadratic form to test for non-linearity
#******************************************
# mydata$Germline2 <- mydata$Germline^2
# MGermlineLongevity_quad <- pglsSEyPagel(MalignancyPrevalence ~ Germline + Germline2, data = mydata, tree=tre, se=SE, method = "ML")
# summary(MGermlineLongevity_quad)
# anova(MGermlineLongevity_quad)
# plot(MGermlineLongevity_quad, which = 1)
#*******************************

### Malignancy vs Oncogene + Longevity
MOncogeneLongevity <- pglsSEyPagel(MalignancyPrevalence ~ Oncogene + maximum_longevity_m, data = mydata, tree=tre, se=SE, method = "ML")
summary(MOncogeneLongevity)
anova(MOncogeneLongevity)
# FDR test and extracting the adj. p-values from the model
MOncogeneLongevity_fdr_corrected <- p.adjust(summary(MOncogeneLongevity)$tTable[, "p-value"], method = "fdr")
MOncogeneLongevity_fdr_corrected
# VIF (variance inflation factor) to ensure multicollinearity is not an issue. 
# If VIF < 5, it's safe to include both variables in the model.
vif(MOncogeneLongevity)
# Grab likelihood-based R2, Lambda, and p values 
RelMOncogeneLongevity  <- R2_lik(MOncogeneLongevity)
n <- nrow(mydata)
p <- 2  # Number of predictors (CGO and longevity)
adjRelMOncogeneLongevity  <- signif(1 - ((1 - RelMOncogeneLongevity) * (n - 1)) / (n - p - 1), 2)
adjRelMOncogeneLongevity
# Check for normality of residuals 
ols_test_normality(MOncogeneLongevity$residuals)
hist(MOncogeneLongevity$residuals, main = "MOncogeneLongevity Residual Histogram")
qqnorm(MOncogeneLongevity$residuals)
qqline(MOncogeneLongevity$residuals)
# Check for independence of errors (autocorrelation) 
# Autocorrelation occurs when the residuals of a regression model are correlated
# with each other, indicating that there is some pattern or dependence among the errors.
model <- lm(MalignancyPrevalence ~ Oncogene + maximum_longevity_m, data=mydata)
durbinWatsonTest(model) #Durbin-Watson test using car package
# Check for heteroscedasticity (Breusch-Pagan test)
# Checks if the variance of the errors is the same for every value of x (homoscedasticity). 
# Look for a random scatter of points around zero, with no discernible pattern or trend. 
# If the spread of residuals appears to vary systematically as the predicted
# values or predictor variable(s) change, it suggests potential heteroscedasticity.
plot(MOncogeneLongevity, which = 1)  # Residuals vs. Fitted value plot
plot(MOncogeneLongevity, which = 3)  # Scale-Location plot (square root of standardized residuals vs. Fitted values)
bptest(model)
### AIC Model Comparison
# Load the data and prepare comparative data object for AIC comparisons
# comparative_data <- comparative.data(phy = tre, data = mydata, 
#                                      names.col = "species", 
#                                      vcv = TRUE, na.omit = FALSE)
# Lambda fixed at 1 (full phylogenetic signal)
model_lambda_1 <- pgls(MalignancyPrevalence ~ Oncogene + maximum_longevity_m, data = comparative_data, lambda = 1.00)
# Compare AIC values
cat("AIC with Lambda = 0:", round(AIC(model),2), "\n")
cat("AIC with Estimated Lambda:", round(AIC(MOncogeneLongevity),2), "\n")
cat("AIC with Lambda = 1:", round(AIC(model_lambda_1),2), "\n")
#******************************************
# Quadratic form to test for non-linearity
#******************************************
# mydata$Oncogene2 <- mydata$Oncogene^2
# MOncogeneLongevity_quad <- pglsSEyPagel(MalignancyPrevalence ~ Oncogene + Oncogene2, data = mydata, tree=tre, se=SE, method = "ML")
# summary(MOncogeneLongevity_quad)
# anova(MOncogeneLongevity_quad)
# plot(MOncogeneLongevity_quad, which = 1)
#*******************************

### Malignancy vs TSG + Longevity
MTSGLongevity <- pglsSEyPagel(MalignancyPrevalence ~ TSG + maximum_longevity_m, data = mydata, tree=tre, se=SE, method = "ML")
summary(MTSGLongevity)
anova(MTSGLongevity)
# FDR test and extracting the adj. p-values from the model
MTSGLongevity_fdr_corrected <- p.adjust(summary(MTSGLongevity)$tTable[, "p-value"], method = "fdr")
MTSGLongevity_fdr_corrected
# VIF (variance inflation factor) to ensure multicollinearity is not an issue. 
# If VIF < 5, it's safe to include both variables in the model.
vif(MTSGLongevity)
# Grab likelihood-based R2, Lambda, and p values 
RelMTSGLongevity  <- R2_lik(MTSGLongevity)
n <- nrow(mydata)
p <- 2  # Number of predictors (CGO and longevity)
adjRelMTSGLongevity  <- signif(1 - ((1 - RelMTSGLongevity) * (n - 1)) / (n - p - 1), 2)
adjRelMTSGLongevity
# Check for normality of residuals 
ols_test_normality(MTSGLongevity$residuals)
hist(MTSGLongevity$residuals, main = "MTSGLongevity Residual Histogram")
qqnorm(MTSGLongevity$residuals)
qqline(MTSGLongevity$residuals)
# Check for independence of errors (autocorrelation) 
# Autocorrelation occurs when the residuals of a regression model are correlated
# with each other, indicating that there is some pattern or dependence among the errors.
model <- lm(MalignancyPrevalence ~ TSG + maximum_longevity_m, data=mydata)
durbinWatsonTest(model) #Durbin-Watson test using car package
# Check for heteroscedasticity (Breusch-Pagan test)
# Checks if the variance of the errors is the same for every value of x (homoscedasticity). 
# Look for a random scatter of points around zero, with no discernible pattern or trend. 
# If the spread of residuals appears to vary systematically as the predicted
# values or predictor variable(s) change, it suggests potential heteroscedasticity.
plot(MTSGLongevity, which = 1)  # Residuals vs. Fitted value plot
plot(MTSGLongevity, which = 3)  # Scale-Location plot (square root of standardized residuals vs. Fitted values)
bptest(model)
### AIC Model Comparison
# Load the data and prepare comparative data object for AIC comparisons
# comparative_data <- comparative.data(phy = tre, data = mydata, 
#                                      names.col = "species", 
#                                      vcv = TRUE, na.omit = FALSE)
# Lambda fixed at 1 (full phylogenetic signal)
model_lambda_1 <- pgls(MalignancyPrevalence ~ TSG + maximum_longevity_m, data = comparative_data, lambda = 1.00)
# Compare AIC values
cat("AIC with Lambda = 0:", round(AIC(model),2), "\n")
cat("AIC with Estimated Lambda:", round(AIC(MTSGLongevity),2), "\n")
cat("AIC with Lambda = 1:", round(AIC(model_lambda_1),2), "\n")
#******************************************
# Quadratic form to test for non-linearity
#******************************************
# mydata$TSG2 <- mydata$TSG^2
# MTSGLongevity_quad <- pglsSEyPagel(MalignancyPrevalence ~ TSG + TSG2, data = mydata, tree=tre, se=SE, method = "ML")
# summary(MTSGLongevity_quad)
# anova(MTSGLongevity_quad)
# plot(MTSGLongevity_quad, which = 1)
#*******************************

### Malignancy vs Fusion + Longevity
MFusionLongevity <- pglsSEyPagel(MalignancyPrevalence ~ Fusion + maximum_longevity_m, data = mydata, tree=tre, se=SE, method = "ML")
summary(MFusionLongevity)
anova(MFusionLongevity)
# FDR test and extracting the adj. p-values from the model
MFusionLongevity_fdr_corrected <- p.adjust(summary(MFusionLongevity)$tTable[, "p-value"], method = "fdr")
MFusionLongevity_fdr_corrected
# VIF (variance inflation factor) to ensure multicollinearity is not an issue. 
# If VIF < 5, it's safe to include both variables in the model.
vif(MFusionLongevity)
# Grab likelihood-based R2, Lambda, and p values 
RelMFusionLongevity  <- R2_lik(MFusionLongevity)
n <- nrow(mydata)
p <- 2  # Number of predictors (CGO and longevity)
adjRelMFusionLongevity  <- signif(1 - ((1 - RelMFusionLongevity) * (n - 1)) / (n - p - 1), 2)
adjRelMFusionLongevity
# Check for normality of residuals 
ols_test_normality(MFusionLongevity$residuals)
hist(MFusionLongevity$residuals, main = "MFusionLongevity Residual Histogram")
qqnorm(MFusionLongevity$residuals)
qqline(MFusionLongevity$residuals)
# Check for independence of errors (autocorrelation) 
# Autocorrelation occurs when the residuals of a regression model are correlated
# with each other, indicating that there is some pattern or dependence among the errors.
model <- lm(MalignancyPrevalence ~ Fusion + maximum_longevity_m, data=mydata)
durbinWatsonTest(model) #Durbin-Watson test using car package
# Check for heteroscedasticity (Breusch-Pagan test)
# Checks if the variance of the errors is the same for every value of x (homoscedasticity). 
# Look for a random scatter of points around zero, with no discernible pattern or trend. 
# If the spread of residuals appears to vary systematically as the predicted
# values or predictor variable(s) change, it suggests potential heteroscedasticity.
plot(MFusionLongevity, which = 1)  # Residuals vs. Fitted value plot
plot(MFusionLongevity, which = 3)  # Scale-Location plot (square root of standardized residuals vs. Fitted values)
bptest(model)
### AIC Model Comparison
# Load the data and prepare comparative data object for AIC comparisons
# comparative_data <- comparative.data(phy = tre, data = mydata, 
#                                      names.col = "species", 
#                                      vcv = TRUE, na.omit = FALSE)
# Lambda fixed at 1 (full phylogenetic signal)
model_lambda_1 <- pgls(MalignancyPrevalence ~ Fusion + maximum_longevity_m, data = comparative_data, lambda = 1.00)
# Compare AIC values
cat("AIC with Lambda = 0:", round(AIC(model),2), "\n")
cat("AIC with Estimated Lambda:", round(AIC(MFusionLongevity),2), "\n")
cat("AIC with Lambda = 1:", round(AIC(model_lambda_1),2), "\n")
#******************************************
# Quadratic form to test for non-linearity
#******************************************
# mydata$Fusion2 <- mydata$Fusion^2
# MFusionLongevity_quad <- pglsSEyPagel(MalignancyPrevalence ~ Fusion + Fusion2, data = mydata, tree=tre, se=SE, method = "ML")
# summary(MFusionLongevity_quad)
# anova(MFusionLongevity_quad)
# plot(MFusionLongevity_quad, which = 1)
#*******************************
```
