## Supplementary Scripts for "Recent Non-LTR Retrotransposon Activity Predicts Cancer Prevalence in Mammals": 08_CGOs_Analyses_Plotting.R

```
################################################################################
####################### CGOs Plotting
########
################################################### 
###  Neoplasia vs Total CGOs
################################################### 
# Define the custom color palette for the Orders
order_colors <- c("Carnivora" = "#414487FF",
                  "Artiodactyla" = "#2A788EFF",
                  "Primates" = "#440154FF",
                  "Rodentia" = "#FCE205",
                  "Chiroptera" = "#22A884FF", 
                  "Other Orders" = "#7AD151FF")


# Store Current model and Response variable in temp vars 
Current_model <- NTotal_CGOs
Current_Response <- mydata$NeoplasiaPrevalence

# Calculate fitted values
fitted_values <- predict(Current_model)
mydata$nudge_y <- ifelse(Current_Response > fitted_values, 0.02, -0.02)

# Classify orders
mydata$color_group <- ifelse(mydata$Order %in% c("Rodentia", "Artiodactyla", "Carnivora", "Chiroptera", "Primates"), 
                             mydata$Order, 
                             "Other Orders")

# Clean species names
mydata$species_clean <- gsub("_", " ", mydata$species)

# Dummy variable for legend formatting
mydata$ModelStats <- "Model Stats"

# Final Plot
plot <- ggplot(mydata, aes(x = Total_CGOs, y = NeoplasiaPrevalence)) +
  
  # Colored dots
  geom_point(aes(color = color_group), size = 3, alpha = 0.9) +
  
  # Species labels (all species)
  geom_text_repel(
    data = mydata,
    aes(label = Common_Name),
    size = 2.5,
    color = "grey20",
    box.padding = 0.3,
    point.padding = 0.8,
    segment.color = "grey60",
    segment.size = 0.3,
    segment.curvature = 0,
    segment.angle = 20,
    min.segment.length = 0.01,
    max.overlaps = Inf) +

  # Regression line
  geom_smooth(method = "lm", se = FALSE, color = "grey50", alpha = 0.7, 
              fullrange = TRUE, linewidth = 0.9) +
  
  # Axis labels
  xlab("Total Number of Cancer Gene Orthologs")+
  ylab("Neoplasia Prevalence")+
  
  # Color legend with model stats
  scale_color_manual(
    name = paste0(
      "<b style='font-size:11pt;'>Model Stats</b><br>",
      nrow(mydata), " species<br>",
      "Corr = ", corrNeo1, "<br>",
      "p value = ", NTotal_CGOs_pvalue, "<span style='color:red;'><b>*</b></span><br>",
      "R² = ", RelNTotal_CGOs, "<br>",
      "λ = ", lambdaNTotal_CGOs, "<br><br>",
      "<b style='font-size:11pt;'>Orders</b>"
    ),
    values = order_colors
  ) +
  
  # Axis limits
  scale_x_continuous(limits = c(-0.08, 1.1), expand = c(0, 0)) +
  scale_y_continuous(limits = c(0.1, 0.97), expand = c(0, 0)) +
  
  # Theme
  theme_minimal(base_size = 6) +
  theme(
    legend.position = "right",
    legend.justification = "bottom",
    legend.direction = "vertical",
    legend.title = element_markdown(size = 10, color = "grey30", lineheight = 1.75),
    legend.text = element_text(size = 9, color = "grey40"),
    plot.title = element_text(size = 12, hjust = 0.5, color = "grey30"),
    axis.title.x = ggtext::element_markdown(size = 12, color = "grey30", margin = margin(t = 10)),
    axis.title.y = ggtext::element_markdown(size = 12, color = "grey30", margin = margin(r = 10)),
    axis.text = element_text(size = 10),
    panel.background = element_rect(fill = "white", color = "white"),
    plot.background = element_rect(fill = "white", color = "white"),
    panel.grid.major = element_blank(),
    panel.grid.minor = element_blank(),
    axis.line = element_line(color = "grey40", linewidth = 0.9),
    panel.border = element_rect(color = "grey40", fill = NA, size = 0.9),
    plot.margin = unit(c(0.5, 0.5, 0.5, 0.5), "cm")
  )

# Print plot
print(plot)

# Save PNG with exact name
ggsave("1_Total_CGOs_vs_Neoplasia_N.png", plot, width = 10, height = 8, dpi = 800)
# Save RDS
saveRDS(plot, "1_Total_CGOs_vs_Neoplasia_N.rds")

################################################### 
###  Neoplasia vs Somatic counts
################################################### 

# Store Current model and Response variable in temp vars 
Current_model <- NSomatic
Current_Response <- mydata$NeoplasiaPrevalence

# Fitted values (optional, but nudge_y is not used)
fitted_values <- predict(Current_model)
mydata$nudge_y <- ifelse(Current_Response > fitted_values, 0.02, -0.02)

# Classify Orders
mydata$color_group <- ifelse(mydata$Order %in% c("Rodentia", "Artiodactyla", "Carnivora", "Chiroptera", "Primates"),
                             mydata$Order, 
                             "Other Orders")

# Clean species name
mydata$species_clean <- gsub("_", " ", mydata$species)

# Dummy variable for legend title
mydata$ModelStats <- "Model Stats"

# Final Plot
plot <- ggplot(mydata, aes(x = Somatic, y = NeoplasiaPrevalence)) +
  
  # Colored points
  geom_point(aes(color = color_group), size = 3, alpha = 0.9) +
  
  # Labels for all species
  geom_text_repel(
    data = mydata,
    aes(label = Common_Name),
    size = 2.5,
    color = "grey20",
    box.padding = 0.3,
    point.padding = 0.8,
    segment.color = "grey60",
    segment.size = 0.3,
    segment.curvature = 0,
    segment.angle = 20,
    min.segment.length = 0.01,
    max.overlaps = Inf
  ) +
  
  # Regression line
  geom_smooth(method = "lm", se = FALSE, color = "grey50", alpha = 0.7,
              fullrange = TRUE, linewidth = 0.9) +
  
  # Axis labels
  xlab("Number of Somatic Cancer Gene Orthologs")+
  ylab("Neoplasia Prevalence")+
  
  # Legend with model stats
  scale_color_manual(
    name = paste0(
      "<b style='font-size:11pt;'>Model Stats</b><br>",
      nrow(mydata), " species<br>",
      "Corr = ", corrNeo2, "<br>",
      "p value = ", NSomatic_pvalue, "<span style='color:red;'><b>*</b></span><br>",
      "R² = ", RelNSomatic, "<br>",
      "λ = ", lambdaNSomatic, "<br><br>",
      "<b style='font-size:11pt;'>Orders</b>"
    ),
    values = order_colors
  ) +
  
  # Axis limits
  scale_x_continuous(limits = c(-0.08, 1.1), expand = c(0, 0)) +
  scale_y_continuous(limits = c(0.1, 0.97), expand = c(0, 0)) +
  
  # Clean theme
  theme_minimal(base_size = 6) +
  theme(
    legend.position = "right",
    legend.justification = "bottom",
    legend.direction = "vertical",
    legend.title = element_markdown(size = 10, color = "grey30", lineheight = 1.75),
    legend.text = element_text(size = 9, color = "grey40"),
    plot.title = element_text(size = 12, hjust = 0.5, color = "grey30"),
    axis.title.x = ggtext::element_markdown(size = 12, color = "grey30", margin = margin(t = 10)),
    axis.title.y = ggtext::element_markdown(size = 12, color = "grey30", margin = margin(r = 10)),
    axis.text = element_text(size = 10),
    panel.background = element_rect(fill = "white", color = "white"),
    plot.background = element_rect(fill = "white", color = "white"),
    panel.grid.major = element_blank(),
    panel.grid.minor = element_blank(),
    axis.line = element_line(color = "grey40", linewidth = 0.9),
    panel.border = element_rect(color = "grey40", fill = NA, size = 0.9),
    plot.margin = unit(c(0.5, 0.5, 0.5, 0.5), "cm")
  )

# Print
print(plot)

# Save with original file name
ggsave("2_Number_of_Somatic_Cancer_Gene_Orthologs_vs_Neoplasia_N.png", plot, width = 10, height = 8, dpi = 800)
saveRDS(plot, "2_Number_of_Somatic_Cancer_Gene_Orthologs_vs_Neoplasia_N.rds")

################################################### 
###  Neoplasia vs Germline counts
################################################### 

# Store Current model and Response variable in temp vars 
Current_model <- NGermline
Current_Response <- mydata$NeoplasiaPrevalence

# Fitted values (optional, but nudge_y is not used)
fitted_values <- predict(Current_model)
mydata$nudge_y <- ifelse(Current_Response > fitted_values, 0.02, -0.02)

# Classify Orders
mydata$color_group <- ifelse(mydata$Order %in% c("Rodentia", "Artiodactyla", "Carnivora", "Chiroptera", "Primates"),
                             mydata$Order, 
                             "Other Orders")

# Clean species name
mydata$species_clean <- gsub("_", " ", mydata$species)

# Dummy variable for legend title
mydata$ModelStats <- "Model Stats"

# Final Plot
plot <- ggplot(mydata, aes(x = Germline, y = NeoplasiaPrevalence)) +
  
  # Colored points
  geom_point(aes(color = color_group), size = 3, alpha = 0.9) +
  
  # Labels for all species
  geom_text_repel(
    data = mydata,
    aes(label = Common_Name),
    size = 2.5,
    color = "grey20",
    box.padding = 0.3,
    point.padding = 0.8,
    segment.color = "grey60",
    segment.size = 0.3,
    segment.curvature = 0,
    segment.angle = 20,
    min.segment.length = 0.01,
    max.overlaps = Inf
  ) +
  
  # Regression line
  geom_smooth(method = "lm", se = FALSE, color = "grey50", alpha = 0.7,
              fullrange = TRUE, linewidth = 0.9) +
  
  # Axis labels
  xlab("Number of Germline Cancer Gene Orthologs")+
  ylab("Neoplasia Prevalence")+
  
  # Legend with model stats
  scale_color_manual(
    name = paste0(
      "<b style='font-size:11pt;'>Model Stats</b><br>",
      nrow(mydata), " species<br>",
      "Corr = ", corrNeo3, "<br>",
      "p value = ", NGermline_pvalue, "<span style='color:red;'><b>*</b></span><br>",
      "R² = ", RelNGermline, "<br>",
      "λ = ", lambdaNGermline, "<br><br>",
      "<b style='font-size:11pt;'>Orders</b>"
    ),
    values = order_colors
  ) +
  
  # Axis limits
  scale_x_continuous(limits = c(-0.08, 1.1), expand = c(0, 0)) +
  scale_y_continuous(limits = c(0.1, 0.97), expand = c(0, 0)) +
  
  # Clean theme
  theme_minimal(base_size = 6) +
  theme(
    legend.position = "right",
    legend.justification = "bottom",
    legend.direction = "vertical",
    legend.title = element_markdown(size = 10, color = "grey30", lineheight = 1.75),
    legend.text = element_text(size = 9, color = "grey40"),
    plot.title = element_text(size = 12, hjust = 0.5, color = "grey30"),
    axis.title.x = ggtext::element_markdown(size = 12, color = "grey30", margin = margin(t = 10)),
    axis.title.y = ggtext::element_markdown(size = 12, color = "grey30", margin = margin(r = 10)),
    axis.text = element_text(size = 10),
    panel.background = element_rect(fill = "white", color = "white"),
    plot.background = element_rect(fill = "white", color = "white"),
    panel.grid.major = element_blank(),
    panel.grid.minor = element_blank(),
    axis.line = element_line(color = "grey40", linewidth = 0.9),
    panel.border = element_rect(color = "grey40", fill = NA, size = 0.9),
    plot.margin = unit(c(0.5, 0.5, 0.5, 0.5), "cm")
  )

# Print
print(plot)

# Save with original file name
ggsave("3_Number_of_Germline_Cancer_Gene_Orthologs_vs_Neoplasia_N_1.png", plot, width = 10, height = 8, dpi = 800)
saveRDS(plot, "3_Number_of_Germline_Cancer_Gene_Orthologs_vs_Neoplasia_N_1.rds")

################################################### 
###  Neoplasia vs Oncogene counts
################################################### 

# Store Current model and Response variable in temp vars 
Current_model <- NOncogene
Current_Response <- mydata$NeoplasiaPrevalence

# Fitted values (optional, but nudge_y is not used)
fitted_values <- predict(Current_model)
mydata$nudge_y <- ifelse(Current_Response > fitted_values, 0.02, -0.02)

# Classify Orders
mydata$color_group <- ifelse(mydata$Order %in% c("Rodentia", "Artiodactyla", "Carnivora", "Chiroptera", "Primates"),
                             mydata$Order, 
                             "Other Orders")

# Clean species name
mydata$species_clean <- gsub("_", " ", mydata$species)

# Dummy variable for legend title
mydata$ModelStats <- "Model Stats"

# Final Plot
plot <- ggplot(mydata, aes(x = Oncogene, y = NeoplasiaPrevalence)) +
  
  # Colored points
  geom_point(aes(color = color_group), size = 3, alpha = 0.9) +
  
  # Labels for all species
  geom_text_repel(
    data = mydata,
    aes(label = Common_Name),
    size = 2.5,
    color = "grey20",
    box.padding = 0.3,
    point.padding = 0.8,
    segment.color = "grey60",
    segment.size = 0.3,
    segment.curvature = 0,
    segment.angle = 20,
    min.segment.length = 0.01,
    max.overlaps = Inf
  ) +
  
  # Regression line
  geom_smooth(method = "lm", se = FALSE, color = "grey50", alpha = 0.7,
              fullrange = TRUE, linewidth = 0.9) +
  
  # Axis labels
  xlab("Number of Oncogene Orthologs")+
  ylab("Neoplasia Prevalence")+
  
  # Legend with model stats
  scale_color_manual(
    name = paste0(
      "<b style='font-size:11pt;'>Model Stats</b><br>",
      nrow(mydata), " species<br>",
      "Corr = ", corrNeo4, "<br>",
      "p value = ", NOncogene_pvalue, "<span style='color:red;'><b>*</b></span><br>",
      "R² = ", RelNOncogene, "<br>",
      "λ = ", lambdaNOncogene, "<br><br>",
      "<b style='font-size:11pt;'>Orders</b>"
    ),
    values = order_colors
  ) +
  
  # Axis limits
  scale_x_continuous(limits = c(-0.08, 1.1), expand = c(0, 0)) +
  scale_y_continuous(limits = c(0.1, 0.97), expand = c(0, 0)) +
  
  # Clean theme
  theme_minimal(base_size = 6) +
  theme(
    legend.position = "right",
    legend.justification = "bottom",
    legend.direction = "vertical",
    legend.title = element_markdown(size = 10, color = "grey30", lineheight = 1.75),
    legend.text = element_text(size = 9, color = "grey40"),
    plot.title = element_text(size = 12, hjust = 0.5, color = "grey30"),
    axis.title.x = ggtext::element_markdown(size = 12, color = "grey30", margin = margin(t = 10)),
    axis.title.y = ggtext::element_markdown(size = 12, color = "grey30", margin = margin(r = 10)),
    axis.text = element_text(size = 10),
    panel.background = element_rect(fill = "white", color = "white"),
    plot.background = element_rect(fill = "white", color = "white"),
    panel.grid.major = element_blank(),
    panel.grid.minor = element_blank(),
    axis.line = element_line(color = "grey40", linewidth = 0.9),
    panel.border = element_rect(color = "grey40", fill = NA, size = 0.9),
    plot.margin = unit(c(0.5, 0.5, 0.5, 0.5), "cm")
  )

# Print
print(plot)

# Save with original file name
ggsave("4_Number_of_Oncogene_Orthologs_vs_Neoplasia_N_1.png", plot, width = 10, height = 8, dpi = 800)
saveRDS(plot, "4_Number_of_Oncogene_Orthologs_vs_Neoplasia_N_1.rds")

################################################### 
###  Neoplasia vs TSG counts
################################################### 

# Store Current model and Response variable in temp vars 
Current_model <- NTSG
Current_Response <- mydata$NeoplasiaPrevalence

# Fitted values (optional, but nudge_y is not used)
fitted_values <- predict(Current_model)
mydata$nudge_y <- ifelse(Current_Response > fitted_values, 0.02, -0.02)

# Classify Orders
mydata$color_group <- ifelse(mydata$Order %in% c("Rodentia", "Artiodactyla", "Carnivora", "Chiroptera", "Primates"),
                             mydata$Order, 
                             "Other Orders")

# Clean species name
mydata$species_clean <- gsub("_", " ", mydata$species)

# Dummy variable for legend title
mydata$ModelStats <- "Model Stats"

# Final Plot
plot <- ggplot(mydata, aes(x = TSG, y = NeoplasiaPrevalence)) +
  
  # Colored points
  geom_point(aes(color = color_group), size = 3, alpha = 0.9) +
  
  # Labels for all species
  geom_text_repel(
    data = mydata,
    aes(label = Common_Name),
    size = 2.5,
    color = "grey20",
    box.padding = 0.3,
    point.padding = 0.8,
    segment.color = "grey60",
    segment.size = 0.3,
    segment.curvature = 0,
    segment.angle = 20,
    min.segment.length = 0.01,
    max.overlaps = Inf
  ) +
  
  # Regression line
  geom_smooth(method = "lm", se = FALSE, color = "grey50", alpha = 0.7,
              fullrange = TRUE, linewidth = 0.9) +
  
  # Axis labels
  xlab("Number of TSG Orthologs")+
  ylab("Neoplasia Prevalence")+
  
  # Legend with model stats
  scale_color_manual(
    name = paste0(
      "<b style='font-size:11pt;'>Model Stats</b><br>",
      nrow(mydata), " species<br>",
      "Corr = ", corrNeo5, "<br>",
      "p value = ", NTSG_pvalue, "<span style='color:red;'><b>*</b></span><br>",
      "R² = ", RelNTSG, "<br>",
      "λ = ", lambdaNTSG, "<br><br>",
      "<b style='font-size:11pt;'>Orders</b>"
    ),
    values = order_colors
  ) +
  
  # Axis limits
  scale_x_continuous(limits = c(-0.08, 1.1), expand = c(0, 0)) +
  scale_y_continuous(limits = c(0.1, 0.97), expand = c(0, 0)) +
  
  # Clean theme
  theme_minimal(base_size = 6) +
  theme(
    legend.position = "right",
    legend.justification = "bottom",
    legend.direction = "vertical",
    legend.title = element_markdown(size = 10, color = "grey30", lineheight = 1.75),
    legend.text = element_text(size = 9, color = "grey40"),
    plot.title = element_text(size = 12, hjust = 0.5, color = "grey30"),
    axis.title.x = ggtext::element_markdown(size = 12, color = "grey30", margin = margin(t = 10)),
    axis.title.y = ggtext::element_markdown(size = 12, color = "grey30", margin = margin(r = 10)),
    axis.text = element_text(size = 10),
    panel.background = element_rect(fill = "white", color = "white"),
    plot.background = element_rect(fill = "white", color = "white"),
    panel.grid.major = element_blank(),
    panel.grid.minor = element_blank(),
    axis.line = element_line(color = "grey40", linewidth = 0.9),
    panel.border = element_rect(color = "grey40", fill = NA, size = 0.9),
    plot.margin = unit(c(0.5, 0.5, 0.5, 0.5), "cm")
  )

# Print
print(plot)

# Save with original file name
ggsave("5_Number_of_TSG_Orthologs_vs_Neoplasia_N.png", plot, width = 10, height = 8, dpi = 800)
saveRDS(plot, "5_Number_of_TSG_Orthologs_vs_Neoplasia_N.rds")

################################################### 
###  Neoplasia vs Fusion counts
################################################### 

# Store Current model and Response variable in temp vars 
Current_model <- NFusion
Current_Response <- mydata$NeoplasiaPrevalence

# Fitted values (optional, but nudge_y is not used)
fitted_values <- predict(Current_model)
mydata$nudge_y <- ifelse(Current_Response > fitted_values, 0.02, -0.02)

# Classify Orders
mydata$color_group <- ifelse(mydata$Order %in% c("Rodentia", "Artiodactyla", "Carnivora", "Chiroptera", "Primates"),
                             mydata$Order, 
                             "Other Orders")

# Clean species name
mydata$species_clean <- gsub("_", " ", mydata$species)

# Dummy variable for legend title
mydata$ModelStats <- "Model Stats"

# Final Plot
plot <- ggplot(mydata, aes(x = Fusion, y = NeoplasiaPrevalence)) +
  
  # Colored points
  geom_point(aes(color = color_group), size = 3, alpha = 0.9) +
  
  # Labels for all species
  geom_text_repel(
    data = mydata,
    aes(label = Common_Name),
    size = 2.5,
    color = "grey20",
    box.padding = 0.3,
    point.padding = 0.8,
    segment.color = "grey60",
    segment.size = 0.3,
    segment.curvature = 0,
    segment.angle = 20,
    min.segment.length = 0.01,
    max.overlaps = Inf
  ) +
  
  # Regression line
  geom_smooth(method = "lm", se = FALSE, color = "grey50", alpha = 0.7,
              fullrange = TRUE, linewidth = 0.9) +
  
  # Axis labels
  xlab("Number of Fusion Cancer Gene Orthologs")+
  ylab("Neoplasia Prevalence")+
  
  # Legend with model stats
  scale_color_manual(
    name = paste0(
      "<b style='font-size:11pt;'>Model Stats</b><br>",
      nrow(mydata), " species<br>",
      "Corr = ", corrNeo6, "<br>",
      "p value = ", NFusion_pvalue, "<span style='color:red;'><b>*</b></span><br>",
      "R² = ", RelNFusion, "<br>",
      "λ = ", lambdaNFusion, "<br><br>",
      "<b style='font-size:11pt;'>Orders</b>"
    ),
    values = order_colors
  ) +
  
  # Axis limits
  scale_x_continuous(limits = c(-0.08, 1.1), expand = c(0, 0)) +
  scale_y_continuous(limits = c(0.1, 0.97), expand = c(0, 0)) +
  
  # Clean theme
  theme_minimal(base_size = 6) +
  theme(
    legend.position = "right",
    legend.justification = "bottom",
    legend.direction = "vertical",
    legend.title = element_markdown(size = 10, color = "grey30", lineheight = 1.75),
    legend.text = element_text(size = 9, color = "grey40"),
    plot.title = element_text(size = 12, hjust = 0.5, color = "grey30"),
    axis.title.x = ggtext::element_markdown(size = 12, color = "grey30", margin = margin(t = 10)),
    axis.title.y = ggtext::element_markdown(size = 12, color = "grey30", margin = margin(r = 10)),
    axis.text = element_text(size = 10),
    panel.background = element_rect(fill = "white", color = "white"),
    plot.background = element_rect(fill = "white", color = "white"),
    panel.grid.major = element_blank(),
    panel.grid.minor = element_blank(),
    axis.line = element_line(color = "grey40", linewidth = 0.9),
    panel.border = element_rect(color = "grey40", fill = NA, size = 0.9),
    plot.margin = unit(c(0.5, 0.5, 0.5, 0.5), "cm")
  )

# Print
print(plot)

# Save with original file name
ggsave("6_Number_of_Fusion_Cancer_Gene_Orthologs_vs_Neoplasia_N.png", plot, width = 10, height = 8, dpi = 800)
saveRDS(plot, "6_Number_of_Fusion_Cancer_Gene_Orthologs_vs_Neoplasia_N.rds")

########################################################################################
########################################################################################
########################################################################################
########################################################################################
########################################################################################
########################################################################################
########################################################################################
########################################################################################
########################################################################################
########################################################################################

################################################### 
###  Malignancy vs Total CGOs
################################################### 

# Store Current model and Response variable in temp vars 
Current_model <- MTotal_CGOs
Current_Response <- mydata$MalignancyPrevalence

# Calculate fitted values
fitted_values <- predict(Current_model)
mydata$nudge_y <- ifelse(Current_Response > fitted_values, 0.02, -0.02)

# Classify orders
mydata$color_group <- ifelse(mydata$Order %in% c("Rodentia", "Artiodactyla", "Carnivora", "Chiroptera", "Primates"), 
                             mydata$Order, 
                             "Other Orders")

# Clean species names
mydata$species_clean <- gsub("_", " ", mydata$species)

# Dummy variable for legend formatting
mydata$ModelStats <- "Model Stats"

# Final Plot
plot <- ggplot(mydata, aes(x = Total_CGOs, y = MalignancyPrevalence)) +
  
  # Colored dots
  geom_point(aes(color = color_group), size = 3, alpha = 0.9) +
  
  # Species labels (all species)
  geom_text_repel(
    data = mydata,
    aes(label = Common_Name),
    size = 2.5,
    color = "grey20",
    box.padding = 0.3,
    point.padding = 0.8,
    segment.color = "grey60",
    segment.size = 0.3,
    segment.curvature = 0,
    segment.angle = 20,
    min.segment.length = 0.01,
    max.overlaps = Inf) +
  
  # Regression line
  geom_smooth(method = "lm", se = FALSE, color = "grey50", alpha = 0.7, 
              fullrange = TRUE, linewidth = 0.9) +
  
  # Axis labels
  xlab("Total Number of Cancer Gene Orthologs")+
  ylab("Malignancy Prevalence")+
  
  # Color legend with model stats
  scale_color_manual(
    name = paste0(
      "<b style='font-size:11pt;'>Model Stats</b><br>",
      nrow(mydata), " species<br>",
      "Corr = ", corrMal1, "<br>",
      #"p value = ", MTotal_CGOs_pvalue, "<span style='color:red;'><b>*</b></span><br>",
      "p value = ", MTotal_CGOs_pvalue, "<br>",
      "R² = ", RelMTotal_CGOs, "<br>",
      "λ = ", lambdaMTotal_CGOs, "<br><br>",
      "<b style='font-size:11pt;'>Orders</b>"
    ),
    values = order_colors
  ) +
  
  # Axis limits
  scale_x_continuous(limits = c(-0.08, 1.1), expand = c(0, 0)) +
  scale_y_continuous(limits = c(0.1, 0.97), expand = c(0, 0)) +
  
  # Theme
  theme_minimal(base_size = 6) +
  theme(
    legend.position = "right",
    legend.justification = "bottom",
    legend.direction = "vertical",
    legend.title = element_markdown(size = 10, color = "grey30", lineheight = 1.75),
    legend.text = element_text(size = 9, color = "grey40"),
    plot.title = element_text(size = 12, hjust = 0.5, color = "grey30"),
    axis.title.x = ggtext::element_markdown(size = 12, color = "grey30", margin = margin(t = 10)),
    axis.title.y = ggtext::element_markdown(size = 12, color = "grey30", margin = margin(r = 10)),
    axis.text = element_text(size = 10),
    panel.background = element_rect(fill = "white", color = "white"),
    plot.background = element_rect(fill = "white", color = "white"),
    panel.grid.major = element_blank(),
    panel.grid.minor = element_blank(),
    axis.line = element_line(color = "grey40", linewidth = 0.9),
    panel.border = element_rect(color = "grey40", fill = NA, size = 0.9),
    plot.margin = unit(c(0.5, 0.5, 0.5, 0.5), "cm")
  )

# Print plot
print(plot)

# Save PNG with exact name
ggsave("7_Total_CGOs_vs_Malignancy_N.png", plot, width = 10, height = 8, dpi = 800)
# Save RDS
saveRDS(plot, "7_Total_CGOs_vs_Malignancy_N.rds")

################################################### 
###  Malignancy vs Somatic counts
################################################### 

# Store Current model and Response variable in temp vars 
Current_model <- MSomatic
Current_Response <- mydata$MalignancyPrevalence

# Fitted values (optional, but nudge_y is not used)
fitted_values <- predict(Current_model)
mydata$nudge_y <- ifelse(Current_Response > fitted_values, 0.02, -0.02)

# Classify Orders
mydata$color_group <- ifelse(mydata$Order %in% c("Rodentia", "Artiodactyla", "Carnivora", "Chiroptera", "Primates"),
                             mydata$Order, 
                             "Other Orders")

# Clean species name
mydata$species_clean <- gsub("_", " ", mydata$species)

# Dummy variable for legend title
mydata$ModelStats <- "Model Stats"

# Final Plot
plot <- ggplot(mydata, aes(x = Somatic, y = MalignancyPrevalence)) +
  
  # Colored points
  geom_point(aes(color = color_group), size = 3, alpha = 0.9) +
  
  # Labels for all species
  geom_text_repel(
    data = mydata,
    aes(label = Common_Name),
    size = 2.5,
    color = "grey20",
    box.padding = 0.3,
    point.padding = 0.8,
    segment.color = "grey60",
    segment.size = 0.3,
    segment.curvature = 0,
    segment.angle = 20,
    min.segment.length = 0.01,
    max.overlaps = Inf
  ) +
  
  # Regression line
  geom_smooth(method = "lm", se = FALSE, color = "grey50", alpha = 0.7,
              fullrange = TRUE, linewidth = 0.9) +
  
  # Axis labels
  xlab("Number of Somatic Cancer Gene Orthologs")+
  ylab("Malignancy Prevalence")+
  
  # Legend with model stats
  scale_color_manual(
    name = paste0(
      "<b style='font-size:11pt;'>Model Stats</b><br>",
      nrow(mydata), " species<br>",
      "Corr = ", corrMal2, "<br>",
      #"p value = ", MSomatic_pvalue, "<span style='color:red;'><b>*</b></span><br>",
      "p value = ", MSomatic_pvalue, "<br>",
      "R² = ", RelMSomatic, "<br>",
      "λ = ", lambdaMSomatic, "<br><br>",
      "<b style='font-size:11pt;'>Orders</b>"
    ),
    values = order_colors
  ) +
  
  # Axis limits
  scale_x_continuous(limits = c(-0.08, 1.1), expand = c(0, 0)) +
  scale_y_continuous(limits = c(0.1, 0.97), expand = c(0, 0)) +
  
  # Clean theme
  theme_minimal(base_size = 6) +
  theme(
    legend.position = "right",
    legend.justification = "bottom",
    legend.direction = "vertical",
    legend.title = element_markdown(size = 10, color = "grey30", lineheight = 1.75),
    legend.text = element_text(size = 9, color = "grey40"),
    plot.title = element_text(size = 12, hjust = 0.5, color = "grey30"),
    axis.title.x = ggtext::element_markdown(size = 12, color = "grey30", margin = margin(t = 10)),
    axis.title.y = ggtext::element_markdown(size = 12, color = "grey30", margin = margin(r = 10)),
    axis.text = element_text(size = 10),
    panel.background = element_rect(fill = "white", color = "white"),
    plot.background = element_rect(fill = "white", color = "white"),
    panel.grid.major = element_blank(),
    panel.grid.minor = element_blank(),
    axis.line = element_line(color = "grey40", linewidth = 0.9),
    panel.border = element_rect(color = "grey40", fill = NA, size = 0.9),
    plot.margin = unit(c(0.5, 0.5, 0.5, 0.5), "cm")
  )

# Print
print(plot)

# Save with original file name
ggsave("8_Number_of_Somatic_Cancer_Gene_Orthologs_vs_Malignancy_N.png", plot, width = 10, height = 8, dpi = 800)
saveRDS(plot, "8_Number_of_Somatic_Cancer_Gene_Orthologs_vs_Malignancy_N.rds")

################################################### 
###  Malignancy vs Germline counts
################################################### 

# Store Current model and Response variable in temp vars 
Current_model <- MGermline
Current_Response <- mydata$MalignancyPrevalence

# Fitted values (optional, but nudge_y is not used)
fitted_values <- predict(Current_model)
mydata$nudge_y <- ifelse(Current_Response > fitted_values, 0.02, -0.02)

# Classify Orders
mydata$color_group <- ifelse(mydata$Order %in% c("Rodentia", "Artiodactyla", "Carnivora", "Chiroptera", "Primates"),
                             mydata$Order, 
                             "Other Orders")

# Clean species name
mydata$species_clean <- gsub("_", " ", mydata$species)

# Dummy variable for legend title
mydata$ModelStats <- "Model Stats"

# Final Plot
plot <- ggplot(mydata, aes(x = Germline, y = MalignancyPrevalence)) +
  
  # Colored points
  geom_point(aes(color = color_group), size = 3, alpha = 0.9) +
  
  # Labels for all species
  geom_text_repel(
    data = mydata,
    aes(label = Common_Name),
    size = 2.5,
    color = "grey20",
    box.padding = 0.3,
    point.padding = 0.8,
    segment.color = "grey60",
    segment.size = 0.3,
    segment.curvature = 0,
    segment.angle = 20,
    min.segment.length = 0.01,
    max.overlaps = Inf
  ) +
  
  # Regression line
  geom_smooth(method = "lm", se = FALSE, color = "grey50", alpha = 0.7,
              fullrange = TRUE, linewidth = 0.9) +
  
  # Axis labels
  xlab("Number of Germline Cancer Gene Orthologs")+
  ylab("Malignancy Prevalence")+
  
  # Legend with model stats
  scale_color_manual(
    name = paste0(
      "<b style='font-size:11pt;'>Model Stats</b><br>",
      nrow(mydata), " species<br>",
      "Corr = ", corrMal3, "<br>",
      #"p value = ", MGermline_pvalue, "<span style='color:red;'><b>*</b></span><br>",
      "p value = ", MGermline_pvalue, "<br>",
      "R² = ", RelMGermline, "<br>",
      "λ = ", lambdaMGermline, "<br><br>",
      "<b style='font-size:11pt;'>Orders</b>"
    ),
    values = order_colors
  ) +
  
  # Axis limits
  scale_x_continuous(limits = c(-0.08, 1.1), expand = c(0, 0)) +
  scale_y_continuous(limits = c(0.1, 0.97), expand = c(0, 0)) +
  
  # Clean theme
  theme_minimal(base_size = 6) +
  theme(
    legend.position = "right",
    legend.justification = "bottom",
    legend.direction = "vertical",
    legend.title = element_markdown(size = 10, color = "grey30", lineheight = 1.75),
    legend.text = element_text(size = 9, color = "grey40"),
    plot.title = element_text(size = 12, hjust = 0.5, color = "grey30"),
    axis.title.x = ggtext::element_markdown(size = 12, color = "grey30", margin = margin(t = 10)),
    axis.title.y = ggtext::element_markdown(size = 12, color = "grey30", margin = margin(r = 10)),
    axis.text = element_text(size = 10),
    panel.background = element_rect(fill = "white", color = "white"),
    plot.background = element_rect(fill = "white", color = "white"),
    panel.grid.major = element_blank(),
    panel.grid.minor = element_blank(),
    axis.line = element_line(color = "grey40", linewidth = 0.9),
    panel.border = element_rect(color = "grey40", fill = NA, size = 0.9),
    plot.margin = unit(c(0.5, 0.5, 0.5, 0.5), "cm")
  )

# Print
print(plot)

# Save with original file name
ggsave("9_Number_of_Germline_Cancer_Gene_Orthologs_vs_Malignancy_N_1.png", plot, width = 10, height = 8, dpi = 800)
saveRDS(plot, "9_Number_of_Germline_Cancer_Gene_Orthologs_vs_Malignancy_N_1.rds")

################################################### 
###  Malignancy vs Oncogene counts
################################################### 

# Store Current model and Response variable in temp vars 
Current_model <- MOncogene
Current_Response <- mydata$MalignancyPrevalence

# Fitted values (optional, but nudge_y is not used)
fitted_values <- predict(Current_model)
mydata$nudge_y <- ifelse(Current_Response > fitted_values, 0.02, -0.02)

# Classify Orders
mydata$color_group <- ifelse(mydata$Order %in% c("Rodentia", "Artiodactyla", "Carnivora", "Chiroptera", "Primates"),
                             mydata$Order, 
                             "Other Orders")

# Clean species name
mydata$species_clean <- gsub("_", " ", mydata$species)

# Dummy variable for legend title
mydata$ModelStats <- "Model Stats"

# Final Plot
plot <- ggplot(mydata, aes(x = Oncogene, y = MalignancyPrevalence)) +
  
  # Colored points
  geom_point(aes(color = color_group), size = 3, alpha = 0.9) +
  
  # Labels for all species
  geom_text_repel(
    data = mydata,
    aes(label = Common_Name),
    size = 2.5,
    color = "grey20",
    box.padding = 0.3,
    point.padding = 0.8,
    segment.color = "grey60",
    segment.size = 0.3,
    segment.curvature = 0,
    segment.angle = 20,
    min.segment.length = 0.01,
    max.overlaps = Inf
  ) +
  
  # Regression line
  geom_smooth(method = "lm", se = FALSE, color = "grey50", alpha = 0.7,
              fullrange = TRUE, linewidth = 0.9) +
  
  # Axis labels
  xlab("Number of Oncogene Orthologs")+
  ylab("Malignancy Prevalence")+
  
  # Legend with model stats
  scale_color_manual(
    name = paste0(
      "<b style='font-size:11pt;'>Model Stats</b><br>",
      nrow(mydata), " species<br>",
      "Corr = ", corrMal4, "<br>",
      #"p value = ", MOncogene_pvalue, "<span style='color:red;'><b>*</b></span><br>",
      "p value = ", MOncogene_pvalue, "<br>",
      "R² = ", RelMOncogene, "<br>",
      "λ = ", lambdaMOncogene, "<br><br>",
      "<b style='font-size:11pt;'>Orders</b>"
    ),
    values = order_colors
  ) +
  
  # Axis limits
  scale_x_continuous(limits = c(-0.08, 1.1), expand = c(0, 0)) +
  scale_y_continuous(limits = c(0.1, 0.97), expand = c(0, 0)) +
  
  # Clean theme
  theme_minimal(base_size = 6) +
  theme(
    legend.position = "right",
    legend.justification = "bottom",
    legend.direction = "vertical",
    legend.title = element_markdown(size = 10, color = "grey30", lineheight = 1.75),
    legend.text = element_text(size = 9, color = "grey40"),
    plot.title = element_text(size = 12, hjust = 0.5, color = "grey30"),
    axis.title.x = ggtext::element_markdown(size = 12, color = "grey30", margin = margin(t = 10)),
    axis.title.y = ggtext::element_markdown(size = 12, color = "grey30", margin = margin(r = 10)),
    axis.text = element_text(size = 10),
    panel.background = element_rect(fill = "white", color = "white"),
    plot.background = element_rect(fill = "white", color = "white"),
    panel.grid.major = element_blank(),
    panel.grid.minor = element_blank(),
    axis.line = element_line(color = "grey40", linewidth = 0.9),
    panel.border = element_rect(color = "grey40", fill = NA, size = 0.9),
    plot.margin = unit(c(0.5, 0.5, 0.5, 0.5), "cm")
  )

# Print
print(plot)

# Save with original file name
ggsave("10_Number_of_Oncogene_Orthologs_vs_Malignancy_N_1.png", plot, width = 10, height = 8, dpi = 800)
saveRDS(plot, "10_Number_of_Oncogene_Orthologs_vs_Malignancy_N_1.rds")

################################################### 
###  Malignancy vs TSG counts
################################################### 

# Store Current model and Response variable in temp vars 
Current_model <- MTSG
Current_Response <- mydata$MalignancyPrevalence

# Fitted values (optional, but nudge_y is not used)
fitted_values <- predict(Current_model)
mydata$nudge_y <- ifelse(Current_Response > fitted_values, 0.02, -0.02)

# Classify Orders
mydata$color_group <- ifelse(mydata$Order %in% c("Rodentia", "Artiodactyla", "Carnivora", "Chiroptera", "Primates"),
                             mydata$Order, 
                             "Other Orders")

# Clean species name
mydata$species_clean <- gsub("_", " ", mydata$species)

# Dummy variable for legend title
mydata$ModelStats <- "Model Stats"

# Final Plot
plot <- ggplot(mydata, aes(x = TSG, y = MalignancyPrevalence)) +
  
  # Colored points
  geom_point(aes(color = color_group), size = 3, alpha = 0.9) +
  
  # Labels for all species
  geom_text_repel(
    data = mydata,
    aes(label = Common_Name),
    size = 2.5,
    color = "grey20",
    box.padding = 0.3,
    point.padding = 0.8,
    segment.color = "grey60",
    segment.size = 0.3,
    segment.curvature = 0,
    segment.angle = 20,
    min.segment.length = 0.01,
    max.overlaps = Inf
  ) +
  
  # Regression line
  geom_smooth(method = "lm", se = FALSE, color = "grey50", alpha = 0.7,
              fullrange = TRUE, linewidth = 0.9) +
  
  # Axis labels
  xlab("Number of TSG Orthologs")+
  ylab("Malignancy Prevalence")+
  
  # Legend with model stats
  scale_color_manual(
    name = paste0(
      "<b style='font-size:11pt;'>Model Stats</b><br>",
      nrow(mydata), " species<br>",
      "Corr = ", corrMal5, "<br>",
      #"p value = ", MTSG_pvalue, "<span style='color:red;'><b>*</b></span><br>",
      "p value = ", MTSG_pvalue, "<br>",
      "R² = ", RelMTSG, "<br>",
      "λ = ", lambdaMTSG, "<br><br>",
      "<b style='font-size:11pt;'>Orders</b>"
    ),
    values = order_colors
  ) +
  
  # Axis limits
  scale_x_continuous(limits = c(-0.08, 1.1), expand = c(0, 0)) +
  scale_y_continuous(limits = c(0.1, 0.97), expand = c(0, 0)) +
  
  # Clean theme
  theme_minimal(base_size = 6) +
  theme(
    legend.position = "right",
    legend.justification = "bottom",
    legend.direction = "vertical",
    legend.title = element_markdown(size = 10, color = "grey30", lineheight = 1.75),
    legend.text = element_text(size = 9, color = "grey40"),
    plot.title = element_text(size = 12, hjust = 0.5, color = "grey30"),
    axis.title.x = ggtext::element_markdown(size = 12, color = "grey30", margin = margin(t = 10)),
    axis.title.y = ggtext::element_markdown(size = 12, color = "grey30", margin = margin(r = 10)),
    axis.text = element_text(size = 10),
    panel.background = element_rect(fill = "white", color = "white"),
    plot.background = element_rect(fill = "white", color = "white"),
    panel.grid.major = element_blank(),
    panel.grid.minor = element_blank(),
    axis.line = element_line(color = "grey40", linewidth = 0.9),
    panel.border = element_rect(color = "grey40", fill = NA, size = 0.9),
    plot.margin = unit(c(0.5, 0.5, 0.5, 0.5), "cm")
  )

# Print
print(plot)

# Save with original file name
ggsave("11_Number_of_TSG_Orthologs_vs_Malignancy_N.png", plot, width = 10, height = 8, dpi = 800)
saveRDS(plot, "11_Number_of_TSG_Orthologs_vs_Malignancy_N.rds")

################################################### 
###  Malignancy vs Fusion counts
################################################### 

# Store Current model and Response variable in temp vars 
Current_model <- MFusion
Current_Response <- mydata$MalignancyPrevalence

# Fitted values (optional, but nudge_y is not used)
fitted_values <- predict(Current_model)
mydata$nudge_y <- ifelse(Current_Response > fitted_values, 0.02, -0.02)

# Classify Orders
mydata$color_group <- ifelse(mydata$Order %in% c("Rodentia", "Artiodactyla", "Carnivora", "Chiroptera", "Primates"),
                             mydata$Order, 
                             "Other Orders")

# Clean species name
mydata$species_clean <- gsub("_", " ", mydata$species)

# Dummy variable for legend title
mydata$ModelStats <- "Model Stats"

# Final Plot
plot <- ggplot(mydata, aes(x = Fusion, y = MalignancyPrevalence)) +
  
  # Colored points
  geom_point(aes(color = color_group), size = 3, alpha = 0.9) +
  
  # Labels for all species
  geom_text_repel(
    data = mydata,
    aes(label = Common_Name),
    size = 2.5,
    color = "grey20",
    box.padding = 0.3,
    point.padding = 0.8,
    segment.color = "grey60",
    segment.size = 0.3,
    segment.curvature = 0,
    segment.angle = 20,
    min.segment.length = 0.01,
    max.overlaps = Inf
  ) +
  
  # Regression line
  geom_smooth(method = "lm", se = FALSE, color = "grey50", alpha = 0.7,
              fullrange = TRUE, linewidth = 0.9) +
  
  # Axis labels
  xlab("Number of Fusion Cancer Gene Orthologs")+
  ylab("Malignancy Prevalence")+
  
  # Legend with model stats
  scale_color_manual(
    name = paste0(
      "<b style='font-size:11pt;'>Model Stats</b><br>",
      nrow(mydata), " species<br>",
      "Corr = ", corrMal6, "<br>",
      #"p value = ", MFusion_pvalue, "<span style='color:red;'><b>*</b></span><br>",
      "p value = ", MFusion_pvalue, "<br>",
      "R² = ", RelMFusion, "<br>",
      "λ = ", lambdaMFusion, "<br><br>",
      "<b style='font-size:11pt;'>Orders</b>"
    ),
    values = order_colors
  ) +
  
  # Axis limits
  scale_x_continuous(limits = c(-0.08, 1.1), expand = c(0, 0)) +
  scale_y_continuous(limits = c(0.1, 0.97), expand = c(0, 0)) +
  
  # Clean theme
  theme_minimal(base_size = 6) +
  theme(
    legend.position = "right",
    legend.justification = "bottom",
    legend.direction = "vertical",
    legend.title = element_markdown(size = 10, color = "grey30", lineheight = 1.75),
    legend.text = element_text(size = 9, color = "grey40"),
    plot.title = element_text(size = 12, hjust = 0.5, color = "grey30"),
    axis.title.x = ggtext::element_markdown(size = 12, color = "grey30", margin = margin(t = 10)),
    axis.title.y = ggtext::element_markdown(size = 12, color = "grey30", margin = margin(r = 10)),
    axis.text = element_text(size = 10),
    panel.background = element_rect(fill = "white", color = "white"),
    plot.background = element_rect(fill = "white", color = "white"),
    panel.grid.major = element_blank(),
    panel.grid.minor = element_blank(),
    axis.line = element_line(color = "grey40", linewidth = 0.9),
    panel.border = element_rect(color = "grey40", fill = NA, size = 0.9),
    plot.margin = unit(c(0.5, 0.5, 0.5, 0.5), "cm")
  )

# Print
print(plot)

# Save with original file name
ggsave("12_Number_of_Fusion_Cancer_Gene_Orthologs_vs_Malignancy_N.png", plot, width = 10, height = 8, dpi = 800)
saveRDS(plot, "12_Number_of_Fusion_Cancer_Gene_Orthologs_vs_Malignancy_N.rds")
```
