## Supplementary Scripts for "Recent Non-LTR Retrotransposon Activity Predicts Cancer Prevalence in Mammals": 09_Interrelationships_Among_Predictors.R

```
###################################################################################################
######################################### Longevity vs nLTR Abundance Analyses (n = 55)
#################
rm(list=ls(all=TRUE))
ls()

#####################################################################
suppressPackageStartupMessages({
  library(ggplot2)
  library(reshape2)
  library(plyr)
  library(tidyverse)
  library(dplyr)
  library(rcompanion)# Tukey Transformation
  library(car) # Durbin Watson Test
  library(phytools) # PGLS 
  library(olsrr) # Assess the normality of residuals
  library(MASS) # Transformation
  library(lmtest) # Breusch_Pagan test
  library(outliers) # To identify outliers
  library(ggimage) # Plotting
  library(magick) # Plotting
  library(ggtree)
  library(cowplot) # For Plotting
  library(ggtext) # For legends in plots
  library(ggrepel) # To avoid overlaps between labels in plots
  library(viridis) # Color blind palette for plots
  library(caper) # For reading tree
  library(rr2) # For reading tree
  library(devtools) # For RPICR
  #library(ROBRT) # For RPICR > You need to use install_github from devtools
  library(geiger) # for RPICR
})
#####################################################################
########## Load custom PGLS functions into the current session
source("Path to /0_pglsSEyPagel.R")
#####################################################################
setwd("Path to the working directory")
getwd()

# Reading Excel data 
mydata <- read.csv("Path to the Supplementary Data S1/Supplementary Data S1.csv", header = TRUE)

#####################################################################
########## Load the ultrametric phylogenetic tree
tre <- read.tree("Mammals_tree.newick")

########## See the orders and superorders
species_count <- as.data.frame(table(mydata$SuperOrder))
colnames(species_count) <- c("SuperOrder","Count")  # Rename the columns
print(species_count)
#View(mydata[mydata$SuperOrder == "Marsupialia", ])

species_count <- as.data.frame(table(mydata$Order))
colnames(species_count) <- c("Order","Count")  # Rename the columns
print(species_count)
#View(mydata[mydata$Order == "Diprotodontia", ])

#####################################################################
# Extract the top 5 and bottom 5 species for each of the variables 

# Top and bottom 5 for Neoplasia
mydata[order(-mydata$NeoplasiaPrevalence), ][1:5, c("Common_Name", "NeoplasiaPrevalence")]  # Top 5
mydata[order(mydata$NeoplasiaPrevalence), ][1:5, c("Common_Name", "NeoplasiaPrevalence")]   # Bottom 5

# Top and bottom 5 for Malignancy
mydata[order(-mydata$MalignancyPrevalence), ][1:5, c("Common_Name", "MalignancyPrevalence")]  # Top 5
mydata[order(mydata$MalignancyPrevalence), ][1:5, c("Common_Name", "MalignancyPrevalence")]   # Bottom 5

# Top and bottom 5 for L1 counts
mydata[order(-mydata$L1_Counts), ][1:5, c("Common_Name", "L1_Counts")]  # Top 5
mydata[order(mydata$L1_Counts), ][1:10, c("Common_Name", "L1_Counts")]   # Bottom 5

# Top and bottom 5 for SINEs counts
mydata[order(-mydata$SINEs_Counts), ][1:5, c("Common_Name", "SINEs_Counts")]  # Top 5
mydata[order(mydata$SINEs_Counts), ][1:5, c("Common_Name", "SINEs_Counts")]   # Bottom 5


######################################
### Top and bottom 5 cancer/nLTR
######################################
# # Get top 5 species by Neoplasia prevalence
# top_neoplasia <- mydata[order(-mydata$NeoplasiaPrevalence), ][1:10, "Common_Name"]
# top_neoplasia
# # Get top 5 species by L1 counts
# top_L1 <- mydata[order(-mydata$L1_Counts), ][1:10, "Common_Name"]
# top_L1
# # Find intersection
# intersect(top_neoplasia, top_L1)
# 
# # Get bottom 5 species by Neoplasia prevalence
# bottom_neoplasia <- mydata[order(mydata$NeoplasiaPrevalence), ][1:10, "Common_Name"]
# bottom_neoplasia
# # Get bottom 5 species by L1 counts
# bottom_L1 <- mydata[order(mydata$L1_Counts), ][1:10, "Common_Name"]
# bottom_L1
# # Find intersection
# intersect(bottom_neoplasia, bottom_L1)

#####################################################################
# Conducting PGLS: Longevity vs nLTR Abundance
#####################################################################
rownames(mydata) <- mydata$species

########## setting SE
SE <- setNames(mydata$SE, mydata$species)[rownames(mydata)]
nrow(mydata)

########## Transform data to make them follow a normal distribution 
###******************************************************************************
### NOTE: Even though it appears normal, We need to transform the longevity data 
### because longevity is the response variable here and the model without  
### transformation does not meet the assumption of residual normality. 
###******************************************************************************

hist(mydata$maximum_longevity_m)
mydata$maximum_longevity_m <- transformTukey(mydata$maximum_longevity_m)
# Scale data to the 0,1 range
mydata$maximum_longevity_m <- (mydata$maximum_longevity_m - min(mydata$maximum_longevity_m)) / (max(mydata$maximum_longevity_m) - min(mydata$maximum_longevity_m))
hist(mydata$maximum_longevity_m)

########### Check for the potential outliers in Longevity data
grubbs_result <- grubbs.test(mydata$maximum_longevity_m)
print(grubbs_result)

################################################################################
################################################################################
### Define Predictors: nLTR Counts (Abundance Model) 
################################################################################
################################################################################
########## Longevity vs Counts and intersections
mydata$Transformed_L1_Counts <- mydata$L1_Counts
mydata$Transformed_L1_SINEs_Counts <- mydata$L1_SINEs_Counts

########## Transform nLTRs data to make them follow a normal distribution 
hist(mydata$Transformed_L1_Counts)
mydata$Transformed_L1_Counts <- transformTukey(mydata$Transformed_L1_Counts)
# Scale data to the 0,1 range
mydata$Transformed_L1_Counts <- (mydata$Transformed_L1_Counts - min(mydata$Transformed_L1_Counts)) / (max(mydata$Transformed_L1_Counts) - min(mydata$Transformed_L1_Counts))
hist(mydata$Transformed_L1_Counts)

hist(mydata$Transformed_L1_SINEs_Counts)
mydata$Transformed_L1_SINEs_Counts <- transformTukey(mydata$Transformed_L1_SINEs_Counts)
# Scale data to the 0,1 range
mydata$Transformed_L1_SINEs_Counts <- (mydata$Transformed_L1_SINEs_Counts - min(mydata$Transformed_L1_SINEs_Counts)) / (max(mydata$Transformed_L1_SINEs_Counts) - min(mydata$Transformed_L1_SINEs_Counts))
hist(mydata$Transformed_L1_SINEs_Counts)

########### Check for the potential outliers in nLTR counts
grubbs_result <- grubbs.test(mydata$Transformed_L1_Counts)
print(grubbs_result)
grubbs_result <- grubbs.test(mydata$Transformed_L1_SINEs_Counts)
print(grubbs_result)

########### Pearson Correlation between Predictors and Longevity
corrLon <- round(cor(mydata$Transformed_L1_Counts, mydata$maximum_longevity_m),2)
corrLon
corrLon2 <- round(cor(mydata$Transformed_L1_SINEs_Counts, mydata$maximum_longevity_m),2)
corrLon2

#######################################################################
### Conducting PGLS
#######################################################################
### Longevity vs L1 Counts
LL1 <- pglsSEyPagel(maximum_longevity_m ~ Transformed_L1_Counts, data = mydata, tree=tre, se=SE, method = "ML")
summary(LL1)
anova(LL1)
# Grab likelihood-based R2, Lambda, and p values for plotting
RelLL1 <- R2_lik(LL1)
RelLL1 <- format(RelLL1)
RelLL1 <- signif(as.numeric(RelLL1), digits = 2)
RelLL1
lambdaLL1 <- summary(LL1)$modelStruct$corStruct
lambdaLL1 <- signif(lambdaLL1[1], digits = 2)
lambdaLL1
LL1_pvalue <- summary(LL1)$tTable
LL1_pvalue <- signif(LL1_pvalue[2,4], digits = 2)
LL1_pvalue
# Check for normality of residuals 
ols_test_normality(LL1$residuals)
hist(LL1$residuals, main = "LL1 Residual Histogram")
qqnorm(LL1$residuals)
qqline(LL1$residuals)
# Check for independence of errors (autocorrelation) 
# Autocorrelation occurs when the residuals of a regression model are correlated
# with each other, indicating that there is some pattern or dependence among the errors.
model <- lm(maximum_longevity_m ~ Transformed_L1_Counts , data=mydata)
durbinWatsonTest(model) #Durbin-Watson test using car package
# Check for heteroscedasticity (Breusch-Pagan test)
# Checks if the variance of the errors is the same for every value of x (homoscedasticity). 
# Look for a random scatter of points around zero, with no discernible pattern or trend. 
# If the spread of residuals appears to vary systematically as the predicted
# values or predictor variable(s) change, it suggests potential heteroscedasticity.
plot(LL1, which = 1)  # Residuals vs. Fitted value plot
plot(LL1, which = 3)  # Scale-Location plot (square root of standardized residuals vs. Fitted values)
bptest(lm(LL1$residuals ~ fitted(LL1)))


### Longevity vs L1+SINEs Counts
LL1SINEs <- pglsSEyPagel(maximum_longevity_m ~ Transformed_L1_SINEs_Counts, data = mydata, tree=tre, se=SE, method = "ML")
summary(LL1SINEs)
anova(LL1SINEs)
# Grab likelihood-based R2, Lambda, and p values for plotting
RelLL1SINEs <- R2_lik(LL1SINEs)
RelLL1SINEs <- format(RelLL1SINEs)
RelLL1SINEs <- signif(as.numeric(RelLL1SINEs), digits = 2)
RelLL1SINEs
lambdaLL1SINEs <- summary(LL1SINEs)$modelStruct$corStruct
lambdaLL1SINEs <- signif(lambdaLL1SINEs[1], digits = 2)
lambdaLL1SINEs
LL1SINEs_pvalue <- summary(LL1SINEs)$tTable
LL1SINEs_pvalue <- signif(LL1SINEs_pvalue[2,4], digits = 2)
LL1SINEs_pvalue
# Check for normality of residuals 
ols_test_normality(LL1SINEs$residuals)
hist(LL1SINEs$residuals, main = "LL1SINEs Residual Histogram")
qqnorm(LL1SINEs$residuals)
qqline(LL1SINEs$residuals)
# Check for independence of errors (autocorrelation) 
# Autocorrelation occurs when the residuals of a regression model are correlated
# with each other, indicating that there is some pattern or dependence among the errors.
model <- lm(maximum_longevity_m ~ Transformed_L1_Counts , data=mydata)
durbinWatsonTest(model) #Durbin-Watson test using car package
# Check for heteroscedasticity (Breusch-Pagan test)
# Checks if the variance of the errors is the same for every value of x (homoscedasticity). 
# Look for a random scatter of points around zero, with no discernible pattern or trend. 
# If the spread of residuals appears to vary systematically as the predicted
# values or predictor variable(s) change, it suggests potential heteroscedasticity.
plot(LL1SINEs, which = 1)  # Residuals vs. Fitted value plot
plot(LL1SINEs, which = 3)  # Scale-Location plot (square root of standardized residuals vs. Fitted values)
bptest(lm(LL1SINEs$residuals ~ fitted(LL1SINEs)))


#####################################################################
# Conducting PGLS: Longevity vs Neoplasia and Malignancy
#####################################################################
########## Check for normality in cancer data
hist(mydata$NeoplasiaPrevalence)
hist(mydata$MalignancyPrevalence)

########## Transform cancer data to make it follow a normal distribution 
### Tukey's Ladder of Powers transform 
mydata$NeoplasiaPrevalence <- transformTukey(mydata$NeoplasiaPrevalence)
mydata$MalignancyPrevalence <- transformTukey(mydata$MalignancyPrevalence)
hist(mydata$NeoplasiaPrevalence)
hist(mydata$MalignancyPrevalence)

########## Check for the poteintial outliers in cancer data
# grubbs_result <- grubbs.test(mydata$NeoplasiaPrevalence)
# print(grubbs_result)
# grubbs_result <- grubbs.test(mydata$MalignancyPrevalence)
# print(grubbs_result)

########## Pearson Correlations Between Neoplasia/Malignancy and Longevity
corrNeo <- round(cor(mydata$NeoplasiaPrevalence, mydata$maximum_longevity_m),2)
corrNeo
corrMal <- round(cor(mydata$MalignancyPrevalence, mydata$maximum_longevity_m),2)
corrMal

### Neoplasia vs L1
NL1 <- pglsSEyPagel(maximum_longevity_m ~ NeoplasiaPrevalence, data = mydata, tree=tre, se=SE, method = "ML")
summary(NL1)
anova(NL1)
# Grab likelihood-based R2, Lambda, and p values for plotting
RelNL1 <- R2_lik(NL1)
RelNL1 <- format(RelNL1)
RelNL1 <- signif(as.numeric(RelNL1), digits = 2)
RelNL1
lambdaNL1 <- summary(NL1)$modelStruct$corStruct
lambdaNL1 <- signif(lambdaNL1[1], digits = 2)
lambdaNL1
NL1_pvalue <- summary(NL1)$tTable
NL1_pvalue <- signif(NL1_pvalue[2,4], digits = 2)
NL1_pvalue
# Check for normality of residuals 
ols_test_normality(NL1$residuals)
hist(NL1$residuals, main = "NL1 Residual Histogram")
qqnorm(NL1$residuals)
qqline(NL1$residuals)
# Check for independence of errors (autocorrelation) 
# Autocorrelation occurs when the residuals of a regression model are correlated
# with each other, indicating that there is some pattern or dependence among the errors.
model <- lm(maximum_longevity_m ~ NeoplasiaPrevalence , data=mydata)
durbinWatsonTest(model) #Durbin-Watson test using car package
# Check for heteroscedasticity (Breusch-Pagan test)
# Checks if the variance of the errors is the same for every value of x (homoscedasticity). 
# Look for a random scatter of points around zero, with no discernible pattern or trend. 
# If the spread of residuals appears to vary systematically as the predicted
# values or predictor variable(s) change, it suggests potential heteroscedasticity.
plot(NL1, which = 1)  # Residuals vs. Fitted value plot
plot(NL1, which = 3)  # Scale-Location plot (square root of standardized residuals vs. Fitted values)
bptest(lm(NL1$residuals ~ fitted(NL1)))


#### Malignancy vs L1
ML1 <- pglsSEyPagel(maximum_longevity_m ~ MalignancyPrevalence, data = mydata, tree=tre, se=SE, method = "ML")
summary(ML1)
anova(ML1)
# Grab likelihood-based R2, Lambda, and p values for plotting
RelML1 <- R2_lik(ML1)
RelML1 <- format(RelML1)
RelML1 <- signif(as.numeric(RelML1), digits= 2)
RelML1
lambdaML1 <- summary(ML1)$modelStruct$corStruct
lambdaML1 <- signif(lambdaML1[1], digits = 2)
lambdaML1
ML1_pvalue <- summary(ML1)$tTable
ML1_pvalue <- signif(ML1_pvalue[2,4], digits = 2)
ML1_pvalue
# Check for normality of residuals 
ols_test_normality(ML1$residuals)
hist(ML1$residuals, main = "ML1 Residual Histogram")
qqnorm(ML1$residuals)
qqline(ML1$residuals)
# Check for independence of errors (autocorrelation) 
# Autocorrelation occurs when the residuals of a regression model are correlated
# with each other, indicating that there is some pattern or dependence among the errors.
model <- lm(maximum_longevity_m ~ MalignancyPrevalence , data=mydata)
durbinWatsonTest(model) #Durbin-Watson test using car package
# Check for heteroscedasticity (Breusch-Pagan test)
# Checks if the variance of the errors is the same for every value of x (homoscedasticity). 
# Look for a random scatter of points around zero, with no discernible pattern or trend. 
# If the spread of residuals appears to vary systematically as the predicted
# values or predictor variable(s) change, it suggests potential heteroscedasticity.
plot(ML1, which = 1)  # Residuals vs. Fitted value plot
plot(ML1, which = 3)  # Scale-Location plot (square root of standardized residuals vs. Fitted values)
bptest(lm(ML1$residuals ~ fitted(ML1)))


###################################################################################################
######################################### Analyses that Need Proteome Data 
################# In these analyses, we remove species with BUSCO score < 0.6 
rm(list=ls(all=TRUE))
ls()

#####################################################################
### nLTR Insertions within PC Genes vs Longevity
#####################################################################
suppressPackageStartupMessages({
  library(ggplot2)
  library(reshape2)
  library(plyr)
  library(tidyverse)
  library(dplyr)
  library(rcompanion)# Tukey Transformation
  library(car) # Durbin Watson Test
  library(phytools) # PGLS 
  library(olsrr) # Assess the normality of residuals
  library(MASS) # Transformation
  library(lmtest) # Breusch_Pagan test
  library(outliers) # To identify outliers
  library(ggimage) # Plotting
  library(magick) # Plotting
  library(ggtree)
  library(cowplot) # For Plotting
  library(ggtext) # For legends in plots
  library(ggrepel) # To avoid overlaps between labels in plots
  library(viridis) # Color blind palette for plots
  library(caper) # For reading tree
  library(rr2) # For reading tree
  library(devtools) # For RPICR
  #library(ROBRT) # For RPICR > You need to use install_github from devtools
  library(geiger) # for RPICR
})
#####################################################################
########## Load custom PGLS functions into the current session
source("Path to /0_pglsSEyPagel.R")
#####################################################################
setwd("Path to the working directory")
getwd()

# Reading Excel data 
mydata <- read.csv("Path to the Supplementary Data S1/Supplementary Data S1.csv", header = TRUE)

#####################################################################
########## Load the ultrametric phylogenetic tree
tre <- read.tree("Mammals_tree.newick")

#####################################################################
# Conducting PGLS: Longevity vs nLTR Abundance
#####################################################################
rownames(mydata) <- mydata$species

########## setting SE
SE <- setNames(mydata$SE, mydata$species)[rownames(mydata)]
nrow(mydata)

########### To remove species with peptide BUSCO score <60%
# Identify species to be removed
removed_species <- mydata$species[mydata$Peptide_BUSCO_Scores <= 60]
print(removed_species)

mydata <- mydata %>%
  filter(Peptide_BUSCO_Scores >= 60)

nrow(mydata)

#####################################################################
# Extract the top 5 and bottom 5 species for CGOs 

# Top and bottom 5 for CGOs
mydata[order(-mydata$Total_CGOs), ][1:5, c("Common_Name", "Total_CGOs")]  # Top 5
mydata[order(mydata$Total_CGOs), ][1:5, c("Common_Name", "Total_CGOs")]   # Bottom 5


########## Transform data to make them follow a normal distribution 
###******************************************************************************
### NOTE: Even though it appears normal, We need to transform the longevity data 
### because the model without transformation does not meet the assumption 
### of residual normality. 
###******************************************************************************

hist(mydata$maximum_longevity_m)
mydata$maximum_longevity_m <- transformTukey(mydata$maximum_longevity_m)
# Scale data to the 0,1 range
mydata$maximum_longevity_m <- (mydata$maximum_longevity_m - min(mydata$maximum_longevity_m)) / (max(mydata$maximum_longevity_m) - min(mydata$maximum_longevity_m))
hist(mydata$maximum_longevity_m)

########### Check for the potential outliers in Longevity data
grubbs_result <- grubbs.test(mydata$maximum_longevity_m)
print(grubbs_result)

################################################################################
### Define Predictors: nLTR-PC Insertions (Genic-Insertion Model) 
################################################################################
### Note: We skipped the Longevity vs nLTR-CGOs insertion analysis due to 
### numerous zeros in that data

########## PC Intersection Variables
mydata$Transformed_L1_PC_Intersections <- mydata$L1_PC_Intersections
mydata$Transformed_L1_SINEs_PC_Intersections <- mydata$L1_SINEs_PC_Intersections

########## Transform data to make them follow a normal distribution 
hist(mydata$Transformed_L1_PC_Intersections)
mydata$Transformed_L1_PC_Intersections <- transformTukey(mydata$Transformed_L1_PC_Intersections)
# Scale data to the 0,1 range
mydata$Transformed_L1_PC_Intersections <- (mydata$Transformed_L1_PC_Intersections - min(mydata$Transformed_L1_PC_Intersections)) / (max(mydata$Transformed_L1_PC_Intersections) - min(mydata$Transformed_L1_PC_Intersections))
hist(mydata$Transformed_L1_PC_Intersections)


hist(mydata$Transformed_L1_SINEs_PC_Intersections)
mydata$Transformed_L1_SINEs_PC_Intersections <- transformTukey(mydata$Transformed_L1_SINEs_PC_Intersections)
# Scale data to the 0,1 range
mydata$Transformed_L1_SINEs_PC_Intersections <- (mydata$Transformed_L1_SINEs_PC_Intersections - min(mydata$Transformed_L1_SINEs_PC_Intersections)) / (max(mydata$Transformed_L1_SINEs_PC_Intersections) - min(mydata$Transformed_L1_SINEs_PC_Intersections))
hist(mydata$Transformed_L1_SINEs_PC_Intersections)

########### Check for the potential outliers 
grubbs_result <- grubbs.test(mydata$Transformed_L1_PC_Intersections)
print(grubbs_result)
grubbs_result <- grubbs.test(mydata$Transformed_L1_SINEs_PC_Intersections)
print(grubbs_result)

########### Pearson Correlation between Predictors and Longevity
corrInt <- round(cor(mydata$Transformed_L1_PC_Intersections, mydata$maximum_longevity_m),2)
corrInt
corrInt2 <- round(cor(mydata$Transformed_L1_SINEs_PC_Intersections, mydata$maximum_longevity_m),2)
corrInt2

#######################################################################
### Conducting PGLS
#######################################################################
### Longevity vs L1 Intersections with PC Genes
LL1Intersection <- pglsSEyPagel(maximum_longevity_m ~ Transformed_L1_PC_Intersections, data = mydata, tree=tre, se=SE, method = "ML")
summary(LL1Intersection)
anova(LL1Intersection)
# Grab likelihood-based R2, Lambda, and p values for plotting
RelLL1Intersection <- R2_lik(LL1Intersection)
RelLL1Intersection <- format(RelLL1Intersection)
RelLL1Intersection <- signif(as.numeric(RelLL1Intersection), digits = 2)
RelLL1Intersection
lambdaLL1Intersection <- summary(LL1Intersection)$modelStruct$corStruct
lambdaLL1Intersection <- signif(lambdaLL1Intersection[1], digits = 2)
lambdaLL1Intersection
LL1Intersection_pvalue <- summary(LL1Intersection)$tTable
LL1Intersection_pvalue <- signif(LL1Intersection_pvalue[2,4], digits = 2)
LL1Intersection_pvalue
# Check for normality of residuals 
ols_test_normality(LL1Intersection$residuals)
hist(LL1Intersection$residuals, main = "LL1Intersection Residual Histogram")
qqnorm(LL1Intersection$residuals)
qqline(LL1Intersection$residuals)
# Check for independence of errors (autocorrelation) 
# Autocorrelation occurs when the residuals of a regression model are correlated
# with each other, indicating that there is some pattern or dependence among the errors.
model <- lm(maximum_longevity_m ~ Transformed_L1_PC_Intersections , data=mydata)
durbinWatsonTest(model) #Durbin-Watson test using car package
# Check for heteroscedasticity (Breusch-Pagan test)
# Checks if the variance of the errors is the same for every value of x (homoscedasticity). 
# Look for a random scatter of points around zero, with no discernible pattern or trend. 
# If the spread of residuals appears to vary systematically as the predicted
# values or predictor variable(s) change, it suggests potential heteroscedasticity.
plot(LL1Intersection, which = 1)  # Residuals vs. Fitted value plot
plot(LL1Intersection, which = 3)  # Scale-Location plot (square root of standardized residuals vs. Fitted values)
bptest(lm(LL1Intersection$residuals ~ fitted(LL1Intersection)))
#******************************************
# Quadratic form to test for non-linearity
#******************************************
# mydata$Transformed_L1_PC_Intersections2 <- mydata$Transformed_L1_PC_Intersections^2
# MTransformed_L1_PC_Intersections2_quad <- pglsSEyPagel(maximum_longevity_m ~ Transformed_L1_PC_Intersections + 
#                                                        Transformed_L1_PC_Intersections2, 
#                                                        data = mydata, tree=tre, se=SE, method = "ML")
# summary(MTransformed_L1_PC_Intersections2_quad)
# anova(MTransformed_L1_PC_Intersections2_quad)
# plot(MTransformed_L1_PC_Intersections2_quad, which = 1)


### Longevity vs L1+SINEs Intersections with PC Genes
LL1SINEsIntersection <- pglsSEyPagel(maximum_longevity_m ~ Transformed_L1_SINEs_PC_Intersections, data = mydata, tree=tre, se=SE, method = "ML")
summary(LL1SINEsIntersection)
anova(LL1SINEsIntersection)
# Grab likelihood-based R2, Lambda, and p values for plotting
RelLL1SINEsIntersection <- R2_lik(LL1SINEsIntersection)
RelLL1SINEsIntersection <- format(RelLL1SINEsIntersection)
RelLL1SINEsIntersection <- signif(as.numeric(RelLL1SINEsIntersection), digits = 2)
RelLL1SINEsIntersection
lambdaLL1SINEsIntersection <- summary(LL1SINEsIntersection)$modelStruct$corStruct
lambdaLL1SINEsIntersection <- signif(lambdaLL1SINEsIntersection[1], digits = 2)
lambdaLL1SINEsIntersection
LL1SINEsIntersection_pvalue <- summary(LL1SINEsIntersection)$tTable
LL1SINEsIntersection_pvalue <- signif(LL1SINEsIntersection_pvalue[2,4], digits = 2)
LL1SINEsIntersection_pvalue
# Check for normality of residuals 
ols_test_normality(LL1SINEsIntersection$residuals)
hist(LL1SINEsIntersection$residuals, main = "LL1SINEsIntersection Residual Histogram")
qqnorm(LL1SINEsIntersection$residuals)
qqline(LL1SINEsIntersection$residuals)
# Check for independence of errors (autocorrelation) 
# Autocorrelation occurs when the residuals of a regression model are correlated
# with each other, indicating that there is some pattern or dependence among the errors.
model <- lm(maximum_longevity_m ~ Transformed_L1_SINEs_PC_Intersections , data=mydata)
durbinWatsonTest(model) #Durbin-Watson test using car package
# Check for heteroscedasticity (Breusch-Pagan test)
# Checks if the variance of the errors is the same for every value of x (homoscedasticity). 
# Look for a random scatter of points around zero, with no discernible pattern or trend. 
# If the spread of residuals appears to vary systematically as the predicted
# values or predictor variable(s) change, it suggests potential heteroscedasticity.
plot(LL1SINEsIntersection, which = 1)  # Residuals vs. Fitted value plot
plot(LL1SINEsIntersection, which = 3)  # Scale-Location plot (square root of standardized residuals vs. Fitted values)
bptest(lm(LL1SINEsIntersection$residuals ~ fitted(LL1SINEsIntersection)))
#******************************************
# Quadratic form to test for non-linearity
#******************************************
# mydata$Transformed_L1_SINEs_PC_Intersections2 <- mydata$Transformed_L1_SINEs_PC_Intersections^2
# MTransformed_L1_SINEs_PC_Intersections2_quad <- pglsSEyPagel(maximum_longevity_m ~ Transformed_L1_SINEs_PC_Intersections +
#                                                        Transformed_L1_SINEs_PC_Intersections2,
#                                                        data = mydata, tree=tre, se=SE, method = "ML")
# summary(MTransformed_L1_SINEs_PC_Intersections2_quad)
# anova(MTransformed_L1_SINEs_PC_Intersections2_quad)
# plot(MTransformed_L1_SINEs_PC_Intersections2_quad, which = 1)


###################################################################################################
######################################### Analyses that Need Proteome Data 
################# In these analyses, we remove species with BUSCO score < 0.6 
rm(list=ls(all=TRUE))
ls()

#####################################################################
### nLTRs vs Fusion Genes
#####################################################################
suppressPackageStartupMessages({
  library(ggplot2)
  library(reshape2)
  library(plyr)
  library(tidyverse)
  library(dplyr)
  library(rcompanion)# Tukey Transformation
  library(car) # Durbin Watson Test
  library(phytools) # PGLS 
  library(olsrr) # Assess the normality of residuals
  library(MASS) # Transformation
  library(lmtest) # Breusch_Pagan test
  library(outliers) # To identify outliers
  library(ggimage) # Plotting
  library(magick) # Plotting
  library(ggtree)
  library(cowplot) # For Plotting
  library(ggtext) # For legends in plots
  library(ggrepel) # To avoid overlaps between labels in plots
  library(viridis) # Color blind palette for plots
  library(caper) # For reading tree
  library(rr2) # For reading tree
  library(devtools) # For RPICR
  #library(ROBRT) # For RPICR > You need to use install_github from devtools
  library(geiger) # for RPICR
})
#####################################################################
########## Load custom PGLS functions into the current session
source("Path to /0_pglsSEyPagel.R")
#####################################################################
setwd("Path to the working directory")
getwd()

# Reading Excel data 
mydata <- read.csv("Path to the Supplementary Data S1/Supplementary Data S1.csv", header = TRUE)

#####################################################################
########## Load the ultrametric phylogenetic tree
tre <- read.tree("Mammals_tree.newick")

#####################################################################
# Conducting PGLS: Longevity vs nLTR Abundance
#####################################################################
rownames(mydata) <- mydata$species

########## setting SE
SE <- setNames(mydata$SE, mydata$species)[rownames(mydata)]
nrow(mydata)

########### To remove species with peptide BUSCO score <60%
# Identify species to be removed
removed_species <- mydata$species[mydata$Peptide_BUSCO_Scores <= 60]
print(removed_species)

mydata <- mydata %>%
  filter(Peptide_BUSCO_Scores >= 60)

nrow(mydata)


########## Response Variable
hist(mydata$Fusion)
mydata$Fusion <- transformTukey(mydata$Fusion)
#hist(mydata$Fusion)
mydata$Fusion <- (mydata$Fusion - min(mydata$Fusion)) / (max(mydata$Fusion) - min(mydata$Fusion))
hist(mydata$Fusion)

grubbs_result <- grubbs.test(mydata$Fusion) #needs
print(grubbs_result)

########## Define predictors (number of L1 and SINEs)
mydata$Transformed_L1_Counts <- mydata$L1_Counts
mydata$Transformed_L1_SINEs_Counts <- mydata$L1_SINEs_Counts

########## Transform nLTRs data to make them follow a normal distribution 
hist(mydata$Transformed_L1_Counts)
mydata$Transformed_L1_Counts <- transformTukey(mydata$Transformed_L1_Counts)
# Scale data to the 0,1 range
mydata$Transformed_L1_Counts <- (mydata$Transformed_L1_Counts - min(mydata$Transformed_L1_Counts)) / (max(mydata$Transformed_L1_Counts) - min(mydata$Transformed_L1_Counts))
hist(mydata$Transformed_L1_Counts)


hist(mydata$Transformed_L1_SINEs_Counts)
mydata$Transformed_L1_SINEs_Counts <- transformTukey(mydata$Transformed_L1_SINEs_Counts)
# Scale data to the 0,1 range
mydata$Transformed_L1_SINEs_Counts <- (mydata$Transformed_L1_SINEs_Counts - min(mydata$Transformed_L1_SINEs_Counts)) / (max(mydata$Transformed_L1_SINEs_Counts) - min(mydata$Transformed_L1_SINEs_Counts))
hist(mydata$Transformed_L1_SINEs_Counts)


########### Check for the potential outliers in nLTR counts
grubbs_result <- grubbs.test(mydata$Transformed_L1_Counts)
print(grubbs_result)
grubbs_result <- grubbs.test(mydata$Transformed_L1_SINEs_Counts)
print(grubbs_result)

########## Pearson Correlations Between Neoplasia/Malignancy and nLTR measurements
corrFus <- round(cor(mydata$Fusion, mydata$Transformed_L1_Counts),2)
corrFus
corrFus2 <- round(cor(mydata$Fusion, mydata$Transformed_L1_SINEs_Counts),2)
corrFus2

#######################################################################
### Conducting PGLS
#######################################################################

########### Fusion Genes ~ Number of L1s + e

### Fusion Genes vs L1
FL1 <- pglsSEyPagel(Fusion ~ Transformed_L1_Counts, data = mydata, tree=tre, se=SE, method = "ML")
summary(FL1)
anova(FL1)
# Grab likelihood-based R2, Lambda, and p values for plotting
RelFL1 <- R2_lik(FL1)
RelFL1 <- format(RelFL1)
RelFL1 <- signif(as.numeric(RelFL1), digits = 2)
RelFL1
lambdaFL1 <- summary(FL1)$modelStruct$corStruct
lambdaFL1 <- signif(lambdaFL1[1], digits = 2)
lambdaFL1
FL1_pvalue <- summary(FL1)$tTable
FL1_pvalue <- signif(FL1_pvalue[2,4], digits = 2)
FL1_pvalue
# Check for normality of residuals 
ols_test_normality(FL1$residuals)
hist(FL1$residuals, main = "FL1 Residual Histogram")
qqnorm(FL1$residuals)
qqline(FL1$residuals)
# Check for independence of errors (autocorrelation) 
# Autocorrelation occurs when the residuals of a regression model are correlated
# with each other, indicating that there is some pattern or dependence among the errors.
model <- lm(Fusion ~ Transformed_L1_Counts , data=mydata)
durbinWatsonTest(model) #Durbin-Watson test using car package
# Check for heteroscedasticity (Breusch-Pagan test)
# Checks if the variance of the errors is the same for every value of x (homoscedasticity). 
# Look for a random scatter of points around zero, with no discernible pattern or trend. 
# If the spread of residuals appears to vary systematically as the predicted
# values or predictor variable(s) change, it suggests potential heteroscedasticity.
plot(FL1, which = 1)  # Residuals vs. Fitted value plot
plot(FL1, which = 3)  # Scale-Location plot (square root of standardized residuals vs. Fitted values)
bptest(lm(FL1$residuals ~ fitted(FL1)))


########### Fusion Genes ~ Number of L1 and SINEs + e

#### Fusion Genes vs (L1 + SINES)
FL1SINEs <- pglsSEyPagel(Fusion ~ Transformed_L1_SINEs_Counts, data = mydata, tree=tre, se=SE, method = "ML")
summary(FL1SINEs)
anova(FL1SINEs)
# Grab likelihood-based R2, Lambda, and p values for plotting
RelFL1SINEs <- R2_lik(FL1SINEs)
RelFL1SINEs <- format(RelFL1SINEs)
RelFL1SINEs <- signif(as.numeric(RelFL1SINEs), digits= 2)
RelFL1SINEs
lambdaFL1SINEs <- summary(FL1SINEs)$modelStruct$corStruct
lambdaFL1SINEs <- signif(lambdaFL1SINEs[1], digits = 2)
lambdaFL1SINEs
FL1SINEs_pvalue <- summary(FL1SINEs)$tTable
FL1SINEs_pvalue <- signif(FL1SINEs_pvalue[2,4], digits = 2)
FL1SINEs_pvalue
# Check for normality of residuals 
ols_test_normality(FL1SINEs$residuals)
hist(FL1SINEs$residuals, main = "FL1SINEs Residual Histogram")
qqnorm(FL1SINEs$residuals)
qqline(FL1SINEs$residuals)
# Check for independence of errors (autocorrelation) 
# Autocorrelation occurs when the residuals of a regression model are correlated
# with each other, indicating that there is some pattern or dependence among the errors.
model <- lm(Fusion ~ Transformed_L1_SINEs_Counts , data=mydata)
durbinWatsonTest(model) #Durbin-Watson test using car package
# Check for heteroscedasticity (Breusch-Pagan test)
# Checks if the variance of the errors is the same for every value of x (homoscedasticity). 
# Look for a random scatter of points around zero, with no discernible pattern or trend. 
# If the spread of residuals appears to vary systematically as the predicted
# values or predictor variable(s) change, it suggests potential heteroscedasticity.
plot(FL1SINEs, which = 1)  # Residuals vs. Fitted value plot
plot(FL1SINEs, which = 3)  # Scale-Location plot (square root of standardized residuals vs. Fitted values)
bptest(lm(FL1SINEs$residuals ~ fitted(FL1SINEs)))


#####################################################################
### nLTRs vs nLTR Insertions within PC Genes and CGOs
#####################################################################

mydata$Transformed_L1_PC_Intersections <- mydata$L1_PC_Intersections
mydata$Transformed_L1_SINEs_PC_Intersections <- mydata$L1_SINEs_PC_Intersections

########## Transform nLTRs data to make them follow a normal distribution 
hist(mydata$Transformed_L1_PC_Intersections)
mydata$Transformed_L1_PC_Intersections <- transformTukey(mydata$Transformed_L1_PC_Intersections)
# Scale data to the 0,1 range
mydata$Transformed_L1_PC_Intersections <- (mydata$Transformed_L1_PC_Intersections - min(mydata$Transformed_L1_PC_Intersections)) / (max(mydata$Transformed_L1_PC_Intersections) - min(mydata$Transformed_L1_PC_Intersections))
hist(mydata$Transformed_L1_PC_Intersections)


hist(mydata$Transformed_L1_SINEs_PC_Intersections)
mydata$Transformed_L1_SINEs_PC_Intersections <- transformTukey(mydata$Transformed_L1_SINEs_PC_Intersections)
# Scale data to the 0,1 range
mydata$Transformed_L1_SINEs_PC_Intersections <- (mydata$Transformed_L1_SINEs_PC_Intersections - min(mydata$Transformed_L1_SINEs_PC_Intersections)) / (max(mydata$Transformed_L1_SINEs_PC_Intersections) - min(mydata$Transformed_L1_SINEs_PC_Intersections))
hist(mydata$Transformed_L1_SINEs_PC_Intersections)


########### Check for the potential outliers in nLTR counts
grubbs_result <- grubbs.test(mydata$Transformed_L1_PC_Intersections)
print(grubbs_result)
grubbs_result <- grubbs.test(mydata$Transformed_L1_SINEs_PC_Intersections)
print(grubbs_result)

########## Pearson Correlations 
corr1 <- round(cor(mydata$Transformed_L1_Counts, mydata$Transformed_L1_PC_Intersections),2)
corr1
corr2 <- round(cor(mydata$Transformed_L1_SINEs_Counts, mydata$Transformed_L1_SINEs_PC_Intersections),2)
corr2

### L1 Insertions within PC Genes vs L1
L1L1PC <- pglsSEyPagel(Transformed_L1_PC_Intersections ~ Transformed_L1_Counts, data = mydata, tree=tre, se=SE, method = "ML")
summary(L1L1PC)
anova(L1L1PC)
# Grab likelihood-based R2, Lambda, and p values for plotting
RelL1L1PC <- R2_lik(L1L1PC)
RelL1L1PC <- format(RelL1L1PC)
RelL1L1PC <- signif(as.numeric(RelL1L1PC), digits = 2)
RelL1L1PC
lambdaL1L1PC <- summary(L1L1PC)$modelStruct$corStruct
lambdaL1L1PC <- signif(lambdaL1L1PC[1], digits = 2)
lambdaL1L1PC
L1L1PC_pvalue <- summary(L1L1PC)$tTable
L1L1PC_pvalue <- signif(L1L1PC_pvalue[2,4], digits = 2)
L1L1PC_pvalue
# Check for normality of residuals 
ols_test_normality(L1L1PC$residuals)
hist(L1L1PC$residuals, main = "L1L1PC Residual Histogram")
qqnorm(L1L1PC$residuals)
qqline(L1L1PC$residuals)
# Check for independence of errors (autocorrelation) 
# Autocorrelation occurs when the residuals of a regression model are correlated
# with each other, indicating that there is some pattern or dependence among the errors.
model <- lm(Transformed_L1_PC_Intersections ~ Transformed_L1_Counts , data=mydata)
durbinWatsonTest(model) #Durbin-Watson test using car package
# Check for heteroscedasticity (Breusch-Pagan test)
# Checks if the variance of the errors is the same for every value of x (homoscedasticity). 
# Look for a random scatter of points around zero, with no discernible pattern or trend. 
# If the spread of residuals appears to vary systematically as the predicted
# values or predictor variable(s) change, it suggests potential heteroscedasticity.
plot(L1L1PC, which = 1)  # Residuals vs. Fitted value plot
plot(L1L1PC, which = 3)  # Scale-Location plot (square root of standardized residuals vs. Fitted values)
bptest(lm(L1L1PC$residuals ~ fitted(L1L1PC)))


#### L1 and SINEs Insertions within PC Genes vs L1 + SINES
L1SINEsL1SINEsPC <- pglsSEyPagel(Transformed_L1_SINEs_PC_Intersections ~ Transformed_L1_SINEs_Counts, data = mydata, tree=tre, se=SE, method = "ML")
summary(L1SINEsL1SINEsPC)
anova(L1SINEsL1SINEsPC)
# Grab likelihood-based R2, Lambda, and p values for plotting
RelL1SINEsL1SINEsPC <- R2_lik(L1SINEsL1SINEsPC)
RelL1SINEsL1SINEsPC <- format(RelL1SINEsL1SINEsPC)
RelL1SINEsL1SINEsPC <- signif(as.numeric(RelL1SINEsL1SINEsPC), digits= 2)
RelL1SINEsL1SINEsPC
lambdaL1SINEsL1SINEsPC <- summary(L1SINEsL1SINEsPC)$modelStruct$corStruct
lambdaL1SINEsL1SINEsPC <- signif(lambdaL1SINEsL1SINEsPC[1], digits = 2)
lambdaL1SINEsL1SINEsPC
L1SINEsL1SINEsPC_pvalue <- summary(L1SINEsL1SINEsPC)$tTable
L1SINEsL1SINEsPC_pvalue <- signif(L1SINEsL1SINEsPC_pvalue[2,4], digits = 2)
L1SINEsL1SINEsPC_pvalue
# Check for normality of residuals 
ols_test_normality(L1SINEsL1SINEsPC$residuals)
hist(L1SINEsL1SINEsPC$residuals, main = "L1SINEsL1SINEsPC Residual Histogram")
qqnorm(L1SINEsL1SINEsPC$residuals)
qqline(L1SINEsL1SINEsPC$residuals)
# Check for independence of errors (autocorrelation) 
# Autocorrelation occurs when the residuals of a regression model are correlated
# with each other, indicating that there is some pattern or dependence among the errors.
model <- lm(Transformed_L1_SINEs_PC_Intersections ~ Transformed_L1_SINEs_Counts, data=mydata)
durbinWatsonTest(model) #Durbin-Watson test using car package
# Check for heteroscedasticity (Breusch-Pagan test)
# Checks if the variance of the errors is the same for every value of x (homoscedasticity). 
# Look for a random scatter of points around zero, with no discernible pattern or trend. 
# If the spread of residuals appears to vary systematically as the predicted
# values or predictor variable(s) change, it suggests potential heteroscedasticity.
plot(L1SINEsL1SINEsPC, which = 1)  # Residuals vs. Fitted value plot
plot(L1SINEsL1SINEsPC, which = 3)  # Scale-Location plot (square root of standardized residuals vs. Fitted values)
bptest(lm(L1SINEsL1SINEsPC$residuals ~ fitted(L1SINEsL1SINEsPC)))


############### Multiple PGLS model including counts and intersections to check 
### if PC insertions are orthogonal to nLTR abundance   

#### Neoplasia vs L1 + L1-PC Insertions
NL1L1PCinsertions <- pglsSEyPagel(NeoplasiaPrevalence ~ Transformed_L1_Counts + Transformed_L1_PC_Intersections, 
                                  data = mydata, tree=tre, se=SE, method = "ML")
summary(NL1L1PCinsertions)
anova(NL1L1PCinsertions)
# FDR test and extracting the adj. p-values from the model
NL1L1PCinsertions_fdr_corrected <- p.adjust(summary(NL1L1PCinsertions)$tTable[, "p-value"], method = "fdr")
NL1L1PCinsertions_fdr_corrected
# VIF (variance inflation factor) to ensure multicollinearity is not an issue. 
# If VIF < 5, it's safe to include both variables in the model.
vif(NL1L1PCinsertions)
# Grab likelihood-based R2, Lambda, and p values 
RelNL1L1PCinsertions  <- R2_lik(NL1L1PCinsertions)
n <- nrow(mydata)
p <- 2  # Number of predictors (L1s and longevity)
adjRelNL1L1PCinsertions  <- signif(1 - ((1 - RelNL1L1PCinsertions) * (n - 1)) / (n - p - 1), 2)
adjRelNL1L1PCinsertions
# Check for normality of residuals 
ols_test_normality(NL1L1PCinsertions$residuals)
hist(NL1L1PCinsertions$residuals, main = "NL1L1PCinsertions Residual Histogram")
qqnorm(NL1L1PCinsertions$residuals)
qqline(NL1L1PCinsertions$residuals)
# Check for independence of errors (autocorrelation) 
# Autocorrelation occurs when the residuals of a regression model are correlated
# with each other, indicating that there is some pattern or dependence among the errors.
model <- lm(NeoplasiaPrevalence ~ Transformed_L1_Counts + Transformed_L1_PC_Intersections, data=mydata)
durbinWatsonTest(model) #Durbin-Watson test using car package
# Check for heteroscedasticity (Breusch-Pagan test)
# Checks if the variance of the errors is the same for every value of x (homoscedasticity). 
# Look for a random scatter of points around zero, with no discernible pattern or trend. 
# If the spread of residuals appears to vary systematically as the predicted
# values or predictor variable(s) change, it suggests potential heteroscedasticity.
plot(NL1L1PCinsertions, which = 1)  # Residuals vs. Fitted value plot
plot(NL1L1PCinsertions, which = 3)  # Scale-Location plot (square root of standardized residuals vs. Fitted values)
bptest(model)


#### Neoplasia vs (L1+SINEs) + (L1_SINEs-PC Insertions)
NL1SINEsL1SINEsPCinsertions <- pglsSEyPagel(NeoplasiaPrevalence ~ Transformed_L1_SINEs_Counts + Transformed_L1_SINEs_PC_Intersections, 
                                  data = mydata, tree=tre, se=SE, method = "ML")
summary(NL1SINEsL1SINEsPCinsertions)
anova(NL1SINEsL1SINEsPCinsertions)
# FDR test and extracting the adj. p-values from the model
NL1SINEsL1SINEsPCinsertions_fdr_corrected <- p.adjust(summary(NL1SINEsL1SINEsPCinsertions)$tTable[, "p-value"], method = "fdr")
NL1SINEsL1SINEsPCinsertions_fdr_corrected
# VIF (variance inflation factor) to ensure multicollinearity is not an issue. 
# If VIF < 5, it's safe to include both variables in the model.
vif(NL1SINEsL1SINEsPCinsertions)
# Grab likelihood-based R2, Lambda, and p values 
RelNL1SINEsL1SINEsPCinsertions  <- R2_lik(NL1SINEsL1SINEsPCinsertions)
n <- nrow(mydata)
p <- 2  # Number of predictors (L1s and longevity)
adjRelNL1SINEsL1SINEsPCinsertions  <- signif(1 - ((1 - RelNL1SINEsL1SINEsPCinsertions) * (n - 1)) / (n - p - 1), 2)
adjRelNL1SINEsL1SINEsPCinsertions
# Check for normality of residuals 
ols_test_normality(NL1SINEsL1SINEsPCinsertions$residuals)
hist(NL1SINEsL1SINEsPCinsertions$residuals, main = "NL1SINEsL1SINEsPCinsertions Residual Histogram")
qqnorm(NL1SINEsL1SINEsPCinsertions$residuals)
qqline(NL1SINEsL1SINEsPCinsertions$residuals)
# Check for independence of errors (autocorrelation) 
# Autocorrelation occurs when the residuals of a regression model are correlated
# with each other, indicating that there is some pattern or dependence among the errors.
model <- lm(NeoplasiaPrevalence ~ Transformed_L1_SINEs_Counts + Transformed_L1_SINEs_PC_Intersections, data=mydata)
durbinWatsonTest(model) #Durbin-Watson test using car package
# Check for heteroscedasticity (Breusch-Pagan test)
# Checks if the variance of the errors is the same for every value of x (homoscedasticity). 
# Look for a random scatter of points around zero, with no discernible pattern or trend. 
# If the spread of residuals appears to vary systematically as the predicted
# values or predictor variable(s) change, it suggests potential heteroscedasticity.
plot(NL1SINEsL1SINEsPCinsertions, which = 1)  # Residuals vs. Fitted value plot
plot(NL1SINEsL1SINEsPCinsertions, which = 3)  # Scale-Location plot (square root of standardized residuals vs. Fitted values)
bptest(model)


#####################################################################
### TSGs vs Oncogenes
#####################################################################

hist(mydata$Oncogene)
#mydata$Oncogene <- transformTukey(mydata$Oncogene)
#hist(mydata$Oncogene)
mydata$Oncogene <- (mydata$Oncogene - min(mydata$Oncogene)) / (max(mydata$Oncogene) - min(mydata$Oncogene))
hist(mydata$Oncogene)

hist(mydata$TSG)
#mydata$TSG <- transformTukey(mydata$TSG)
#hist(mydata$TSG)
mydata$TSG <- (mydata$TSG - min(mydata$TSG)) / (max(mydata$TSG) - min(mydata$TSG))
hist(mydata$TSG)


########### Check for the potential outliers in nLTR counts
grubbs_result <- grubbs.test(mydata$Oncogene)
print(grubbs_result)
grubbs_result <- grubbs.test(mydata$TSG)
print(grubbs_result)

########## Pearson Correlations 
corrTSGOnco <- round(cor(mydata$Oncogene, mydata$TSG),2)
corrTSGOnco

### L1 Insertions within PC Genes vs L1
TSGOnco <- pglsSEyPagel(TSG ~ Oncogene, data = mydata, tree=tre, se=SE, method = "ML")
summary(TSGOnco)
anova(TSGOnco)
# Grab likelihood-based R2, Lambda, and p values for plotting
RelTSGOnco <- R2_lik(TSGOnco)
RelTSGOnco <- format(RelTSGOnco)
RelTSGOnco <- signif(as.numeric(RelTSGOnco), digits = 2)
RelTSGOnco
lambdaTSGOnco <- summary(TSGOnco)$modelStruct$corStruct
lambdaTSGOnco <- signif(lambdaTSGOnco[1], digits = 2)
lambdaTSGOnco
TSGOnco_pvalue <- summary(TSGOnco)$tTable
TSGOnco_pvalue <- signif(TSGOnco_pvalue[2,4], digits = 2)
TSGOnco_pvalue
# Check for normality of residuals 
ols_test_normality(TSGOnco$residuals)
hist(TSGOnco$residuals, main = "TSGOnco Residual Histogram")
qqnorm(TSGOnco$residuals)
qqline(TSGOnco$residuals)
# Check for independence of errors (autocorrelation) 
# Autocorrelation occurs when the residuals of a regression model are correlated
# with each other, indicating that there is some pattern or dependence among the errors.
model <- lm(TSG ~ Oncogene , data=mydata)
durbinWatsonTest(model) #Durbin-Watson test using car package
# Check for heteroscedasticity (Breusch-Pagan test)
# Checks if the variance of the errors is the same for every value of x (homoscedasticity). 
# Look for a random scatter of points around zero, with no discernible pattern or trend. 
# If the spread of residuals appears to vary systematically as the predicted
# values or predictor variable(s) change, it suggests potential heteroscedasticity.
plot(TSGOnco, which = 1)  # Residuals vs. Fitted value plot
plot(TSGOnco, which = 3)  # Scale-Location plot (square root of standardized residuals vs. Fitted values)
bptest(lm(TSGOnco$residuals ~ fitted(TSGOnco)))
```
