## Supplementary Scripts for "Recent Non-LTR Retrotransposon Activity Predicts Cancer Prevalence in Mammals": 09_Interrelationships_Among_Predictors_Plotting.R

```
################################################################################
####################### Plotting Interrelationships among Variables

################################################################################
### L1 Counts vs Longevity
################################################################################
# Define the custom color palette for the Orders
order_colors <- c("Carnivora" = "#414487FF",
                  "Artiodactyla" = "#2A788EFF",
                  "Primates" = "#440154FF",
                  "Rodentia" = "#FCE205",
                  "Chiroptera" = "#22A884FF", 
                  "Other Orders" = "#7AD151FF")

# Store Current model and Response variable in temp vars 
Current_model <- LL1
Current_Response <- mydata$maximum_longevity_m

# Calculate fitted values
fitted_values <- predict(Current_model)
mydata$nudge_y <- ifelse(Current_Response > fitted_values, 0.02, -0.02)

# Classify orders
mydata$color_group <- ifelse(mydata$Order %in% c("Rodentia", "Artiodactyla", "Carnivora", "Chiroptera", "Primates"), 
                             mydata$Order, 
                             "Other Orders")

# Clean species names
mydata$species_clean <- gsub("_", " ", mydata$species)

# Dummy variable for legend formatting
mydata$ModelStats <- "Model Stats"

# Final Plot
plot <- ggplot(mydata, aes(x = Transformed_L1_Counts, y = maximum_longevity_m)) +
  
  # Colored dots
  geom_point(aes(color = color_group), size = 3, alpha = 0.9) +
  
  # Species labels (all species)
  geom_text_repel(
    data = mydata,
    aes(label = Common_Name),
    size = 2.5,
    color = "grey20",
    box.padding = 0.3,
    point.padding = 0.8,
    segment.color = "grey60",
    segment.size = 0.3,
    segment.curvature = 0,
    segment.angle = 20,
    min.segment.length = 0.01,
    max.overlaps = Inf
  ) +
  
  # Regression line
  geom_smooth(method = "lm", se = FALSE, color = "grey50", alpha = 0.7, 
              fullrange = TRUE, linewidth = 0.9) +
  
  # Axis labels
  xlab("Number of Active L1")+
  ylab("Longevity")+
  
  # Color legend with model stats
  scale_color_manual(
    name = paste0(
      "<b style='font-size:11pt;'>Model Stats</b><br>",
      nrow(mydata), " species<br>",
      "Corr = ", corrLon, "<br>",
      "p value = ", LL1_pvalue, "<br>",
      #"p value = ", LL1_pvalue, "<span style='color:red;'><b>*</b></span><br>",
      "R² = ", RelLL1, "<br>",
      "λ = ", lambdaLL1, "<br><br>",
      "<b style='font-size:11pt;'>Orders</b>"
    ),
    values = order_colors
  ) +
  
  # Axis limits
  scale_x_continuous(limits = c(-0.1, 1.05), expand = c(0, 0)) +
  scale_y_continuous(limits = c(-0.1, 1.05), expand = c(0, 0)) +
  
  # Theme
  theme_minimal(base_size = 6) +
  theme(
    legend.position = "right",
    legend.justification = "bottom",
    legend.direction = "vertical",
    legend.title = element_markdown(size = 10, color = "grey30", lineheight = 1.75),
    legend.text = element_text(size = 9, color = "grey40"),
    plot.title = element_text(size = 12, hjust = 0.5, color = "grey30"),
    axis.title.x = ggtext::element_markdown(size = 12, color = "grey30", margin = margin(t = 10)),
    axis.title.y = ggtext::element_markdown(size = 12, color = "grey30", margin = margin(r = 10)),
    axis.text = element_text(size = 10),
    panel.background = element_rect(fill = "white", color = "white"),
    plot.background = element_rect(fill = "white", color = "white"),
    panel.grid.major = element_blank(),
    panel.grid.minor = element_blank(),
    axis.line = element_line(color = "grey40", linewidth = 0.9),
    panel.border = element_rect(color = "grey40", fill = NA, size = 0.9),
    plot.margin = unit(c(0.5, 0.5, 0.5, 0.5), "cm")
  )

# Print plot
print(plot)

# Save PNG with exact name
ggsave("1_L1_Counts_vs_Longevity_N.png", plot, width = 10, height = 8, dpi = 800)
# Save RDS
saveRDS(plot, "1_L1_Counts_vs_Longevity_N.rds")


################################################################################
### L1 and SINE Counts vs Longevity
################################################################################

# Store Current model and Response variable in temp vars 
Current_model <- LL1SINEs
Current_Response <- mydata$maximum_longevity_m

# Calculate fitted values
fitted_values <- predict(Current_model)
mydata$nudge_y <- ifelse(Current_Response > fitted_values, 0.02, -0.02)

# Classify orders
mydata$color_group <- ifelse(mydata$Order %in% c("Rodentia", "Artiodactyla", "Carnivora", "Chiroptera", "Primates"), 
                             mydata$Order, 
                             "Other Orders")

# Clean species names
mydata$species_clean <- gsub("_", " ", mydata$species)

# Dummy variable for legend formatting
mydata$ModelStats <- "Model Stats"

# Final Plot
plot <- ggplot(mydata, aes(x = Transformed_L1_SINEs_Counts, y = maximum_longevity_m)) +
  
  # Colored dots
  geom_point(aes(color = color_group), size = 3, alpha = 0.9) +
  
  # Species labels (all species)
  geom_text_repel(
    data = mydata,
    aes(label = Common_Name),
    size = 2.5,
    color = "grey20",
    box.padding = 0.3,
    point.padding = 0.8,
    segment.color = "grey60",
    segment.size = 0.3,
    segment.curvature = 0,
    segment.angle = 20,
    min.segment.length = 0.01,
    max.overlaps = Inf
  ) +
  
  # Regression line
  geom_smooth(method = "lm", se = FALSE, color = "grey50", alpha = 0.7, 
              fullrange = TRUE, linewidth = 0.9) +
  
  # Axis labels
  xlab("Number of Active L1 and SINEs")+
  ylab("Longevity")+
  
  # Color legend with model stats
  scale_color_manual(
    name = paste0(
      "<b style='font-size:11pt;'>Model Stats</b><br>",
      nrow(mydata), " species<br>",
      "Corr = ", corrLon, "<br>",
      "p value = ", LL1SINEs_pvalue, "<br>",
      #"p value = ", LL1SINEs_pvalue, "<span style='color:red;'><b>*</b></span><br>",
      "R² = ", RelLL1SINEs, "<br>",
      "λ = ", lambdaLL1SINEs, "<br><br>",
      "<b style='font-size:11pt;'>Orders</b>"
    ),
    values = order_colors
  ) +
  
  # Axis limits
  scale_x_continuous(limits = c(-0.1, 1.05), expand = c(0, 0)) +
  scale_y_continuous(limits = c(-0.1, 1.05), expand = c(0, 0)) +
  
  # Theme
  theme_minimal(base_size = 6) +
  theme(
    legend.position = "right",
    legend.justification = "bottom",
    legend.direction = "vertical",
    legend.title = element_markdown(size = 10, color = "grey30", lineheight = 1.75),
    legend.text = element_text(size = 9, color = "grey40"),
    plot.title = element_text(size = 12, hjust = 0.5, color = "grey30"),
    axis.title.x = ggtext::element_markdown(size = 12, color = "grey30", margin = margin(t = 10)),
    axis.title.y = ggtext::element_markdown(size = 12, color = "grey30", margin = margin(r = 10)),
    axis.text = element_text(size = 10),
    panel.background = element_rect(fill = "white", color = "white"),
    plot.background = element_rect(fill = "white", color = "white"),
    panel.grid.major = element_blank(),
    panel.grid.minor = element_blank(),
    axis.line = element_line(color = "grey40", linewidth = 0.9),
    panel.border = element_rect(color = "grey40", fill = NA, size = 0.9),
    plot.margin = unit(c(0.5, 0.5, 0.5, 0.5), "cm")
  )

# Print plot
print(plot)

# Save PNG with exact name
ggsave("2_L1_SINEs_Counts_vs_Longevity_N.png", plot, width = 10, height = 8, dpi = 800)
# Save RDS
saveRDS(plot, "2_L1_SINEs_Counts_vs_Longevity_N.rds")


################################################################################
### Longevity vs Neoplasia
################################################################################
# Define the custom color palette for the Orders
order_colors <- c("Carnivora" = "#414487FF",
                  "Artiodactyla" = "#2A788EFF",
                  "Primates" = "#440154FF",
                  "Rodentia" = "#FCE205",
                  "Chiroptera" = "#22A884FF", 
                  "Other Orders" = "#7AD151FF")

# Store Current model and Response variable in temp vars 
Current_model <- NL1
Current_Response <- mydata$maximum_longevity_m

# Calculate fitted values
fitted_values <- predict(Current_model)
mydata$nudge_y <- ifelse(Current_Response > fitted_values, 0.02, -0.02)

# Classify orders
mydata$color_group <- ifelse(mydata$Order %in% c("Rodentia", "Artiodactyla", "Carnivora", "Chiroptera", "Primates"), 
                             mydata$Order, 
                             "Other Orders")

# Clean species names
mydata$species_clean <- gsub("_", " ", mydata$species)

# Dummy variable for legend formatting
mydata$ModelStats <- "Model Stats"

# Final Plot
plot <- ggplot(mydata, aes(x = NeoplasiaPrevalence, y = maximum_longevity_m)) +
  
  # Colored dots
  geom_point(aes(color = color_group), size = 3, alpha = 0.9) +
  
  # Species labels (all species)
  geom_text_repel(
    data = mydata,
    aes(label = Common_Name),
    size = 2.5,
    color = "grey20",
    box.padding = 0.3,
    point.padding = 0.8,
    segment.color = "grey60",
    segment.size = 0.3,
    segment.curvature = 0,
    segment.angle = 20,
    min.segment.length = 0.01,
    max.overlaps = Inf
  ) +
  
  # Regression line
  geom_smooth(method = "lm", se = FALSE, color = "grey50", alpha = 0.7, 
              fullrange = TRUE, linewidth = 0.9) +
  
  # Axis labels
  xlab("Neoplasia Prevalence")+
  ylab("Longevity")+
  
  # Color legend with model stats
  scale_color_manual(
    name = paste0(
      "<b style='font-size:11pt;'>Model Stats</b><br>",
      nrow(mydata), " species<br>",
      "Corr = ", corrNeo, "<br>",
      "p value = ", NL1_pvalue, "<br>",
      #"p value = ", NL1_pvalue, "<span style='color:red;'><b>*</b></span><br>",
      "R² = ", RelNL1, "<br>",
      "λ = ", lambdaNL1, "<br><br>",
      "<b style='font-size:11pt;'>Orders</b>"
    ),
    values = order_colors
  ) +
  
  # Axis limits
  scale_x_continuous(limits = c(-0.1, 1.05), expand = c(0, 0)) +
  scale_y_continuous(limits = c(-0.1, 1.05), expand = c(0, 0)) +
  
  # Theme
  theme_minimal(base_size = 6) +
  theme(
    legend.position = "right",
    legend.justification = "bottom",
    legend.direction = "vertical",
    legend.title = element_markdown(size = 10, color = "grey30", lineheight = 1.75),
    legend.text = element_text(size = 9, color = "grey40"),
    plot.title = element_text(size = 12, hjust = 0.5, color = "grey30"),
    axis.title.x = ggtext::element_markdown(size = 12, color = "grey30", margin = margin(t = 10)),
    axis.title.y = ggtext::element_markdown(size = 12, color = "grey30", margin = margin(r = 10)),
    axis.text = element_text(size = 10),
    panel.background = element_rect(fill = "white", color = "white"),
    plot.background = element_rect(fill = "white", color = "white"),
    panel.grid.major = element_blank(),
    panel.grid.minor = element_blank(),
    axis.line = element_line(color = "grey40", linewidth = 0.9),
    panel.border = element_rect(color = "grey40", fill = NA, size = 0.9),
    plot.margin = unit(c(0.5, 0.5, 0.5, 0.5), "cm")
  )

# Print plot
print(plot)

# Save PNG with exact name
ggsave("9_Neoplasia_vs_Longevity_N.png", plot, width = 10, height = 8, dpi = 800)
# Save RDS
saveRDS(plot, "9_Neoplasia_vs_Longevity_N.rds")


################################################################################
### Longevity vs Malignancy
################################################################################

# Store Current model and Response variable in temp vars 
Current_model <- ML1
Current_Response <- mydata$maximum_longevity_m

# Calculate fitted values
fitted_values <- predict(Current_model)
mydata$nudge_y <- ifelse(Current_Response > fitted_values, 0.02, -0.02)

# Classify orders
mydata$color_group <- ifelse(mydata$Order %in% c("Rodentia", "Artiodactyla", "Carnivora", "Chiroptera", "Primates"), 
                             mydata$Order, 
                             "Other Orders")

# Clean species names
mydata$species_clean <- gsub("_", " ", mydata$species)

# Dummy variable for legend formatting
mydata$ModelStats <- "Model Stats"

# Final Plot
plot <- ggplot(mydata, aes(x = MalignancyPrevalence, y = maximum_longevity_m)) +
  
  # Colored dots
  geom_point(aes(color = color_group), size = 3, alpha = 0.9) +
  
  # Species labels (all species)
  geom_text_repel(
    data = mydata,
    aes(label = Common_Name),
    size = 2.5,
    color = "grey20",
    box.padding = 0.3,
    point.padding = 0.8,
    segment.color = "grey60",
    segment.size = 0.3,
    segment.curvature = 0,
    segment.angle = 20,
    min.segment.length = 0.01,
    max.overlaps = Inf
  ) +
  
  # Regression line
  geom_smooth(method = "lm", se = FALSE, color = "grey50", alpha = 0.7, 
              fullrange = TRUE, linewidth = 0.9) +
  
  # Axis labels
  xlab("Malignancy")+
  ylab("Longevity")+
  
  # Color legend with model stats
  scale_color_manual(
    name = paste0(
      "<b style='font-size:11pt;'>Model Stats</b><br>",
      nrow(mydata), " species<br>",
      "Corr = ", corrMal, "<br>",
      "p value = ", ML1_pvalue, "<br>",
      #"p value = ", ML1SINEs_pvalue, "<span style='color:red;'><b>*</b></span><br>",
      "R² = ", RelML1, "<br>",
      "λ = ", lambdaML1, "<br><br>",
      "<b style='font-size:11pt;'>Orders</b>"
    ),
    values = order_colors
  ) +
  
  # Axis limits
  scale_x_continuous(limits = c(-0.1, 1.05), expand = c(0, 0)) +
  scale_y_continuous(limits = c(-0.1, 1.05), expand = c(0, 0)) +
  
  # Theme
  theme_minimal(base_size = 6) +
  theme(
    legend.position = "right",
    legend.justification = "bottom",
    legend.direction = "vertical",
    legend.title = element_markdown(size = 10, color = "grey30", lineheight = 1.75),
    legend.text = element_text(size = 9, color = "grey40"),
    plot.title = element_text(size = 12, hjust = 0.5, color = "grey30"),
    axis.title.x = ggtext::element_markdown(size = 12, color = "grey30", margin = margin(t = 10)),
    axis.title.y = ggtext::element_markdown(size = 12, color = "grey30", margin = margin(r = 10)),
    axis.text = element_text(size = 10),
    panel.background = element_rect(fill = "white", color = "white"),
    plot.background = element_rect(fill = "white", color = "white"),
    panel.grid.major = element_blank(),
    panel.grid.minor = element_blank(),
    axis.line = element_line(color = "grey40", linewidth = 0.9),
    panel.border = element_rect(color = "grey40", fill = NA, size = 0.9),
    plot.margin = unit(c(0.5, 0.5, 0.5, 0.5), "cm")
  )

# Print plot
print(plot)

# Save PNG with exact name
ggsave("10_Malignancy_vs_Longevity_N.png", plot, width = 10, height = 8, dpi = 800)
# Save RDS
saveRDS(plot, "10_Malignancy_vs_Longevity_N.rds")

#********************************************************************************
#********************************************************************************
#********************************************************************************
#********************************************************************************
#********************************************************************************
#********************************************************************************
#********************************************************************************
#********************************************************************************
#********************************************************************************
#********************************************************************************

################################################################################
### L1 Insertions within PC Genes vs Longevity
################################################################################

# Define the custom color palette for the Orders
order_colors <- c("Carnivora" = "#414487FF",
                  "Artiodactyla" = "#2A788EFF",
                  "Primates" = "#440154FF",
                  "Rodentia" = "#FCE205",
                  "Chiroptera" = "#22A884FF", 
                  "Other Orders" = "#7AD151FF")


# Store Current model and Response variable in temp vars 
Current_model <- LL1Intersection
Current_Response <- mydata$maximum_longevity_m

# Calculate fitted values
fitted_values <- predict(Current_model)
mydata$nudge_y <- ifelse(Current_Response > fitted_values, 0.02, -0.02)

# Classify orders
mydata$color_group <- ifelse(mydata$Order %in% c("Rodentia", "Artiodactyla", "Carnivora", "Chiroptera", "Primates"), 
                             mydata$Order, 
                             "Other Orders")

# Clean species names
mydata$species_clean <- gsub("_", " ", mydata$species)

# Dummy variable for legend formatting
mydata$ModelStats <- "Model Stats"

# Final Plot
plot <- ggplot(mydata, aes(x = Transformed_L1_PC_Intersections, y = maximum_longevity_m)) +
  
  # Colored dots
  geom_point(aes(color = color_group), size = 3, alpha = 0.9) +
  
  # Species labels (all species)
  geom_text_repel(
    data = mydata,
    aes(label = Common_Name),
    size = 2.5,
    color = "grey20",
    box.padding = 0.3,
    point.padding = 0.8,
    segment.color = "grey60",
    segment.size = 0.3,
    segment.curvature = 0,
    segment.angle = 20,
    min.segment.length = 0.01,
    max.overlaps = Inf
  ) +
  
  # Regression line
  geom_smooth(method = "lm", se = FALSE, color = "grey50", alpha = 0.7, 
              fullrange = TRUE, linewidth = 0.9) +
  
  # Axis labels
  xlab("Number of L1 Insertions within PC Genes")+#&#8201;<span style='font-size:8pt;'>(Tukey-Transformed)</span>") +
  ylab("Longevity")+#&#8201;<span style='font-size:8pt;'>(zero-one Normalized)</span>") +
  
  # Color legend with model stats
  scale_color_manual(
    name = paste0(
      "<b style='font-size:11pt;'>Model Stats</b><br>",
      nrow(mydata), " species<br>",
      "Corr = ", corrInt, "<br>",
      "p value = ", LL1Intersection_pvalue, "<br>",
      #"p value = ", LL1Intersection_pvalue, "<span style='color:red;'><b>*</b></span><br>",
      "R² = ", RelLL1Intersection, "<br>",
      "λ = ", lambdaLL1Intersection, "<br><br>",
      "<b style='font-size:11pt;'>Orders</b>"
    ),
    values = order_colors
  ) +
  
  # Axis limits
  scale_x_continuous(limits = c(-0.1, 1.05), expand = c(0, 0)) +
  scale_y_continuous(limits = c(-0.1, 1.05), expand = c(0, 0)) +
  
  # Theme
  theme_minimal(base_size = 6) +
  theme(
    legend.position = "right",
    legend.justification = "bottom",
    legend.direction = "vertical",
    legend.title = element_markdown(size = 10, color = "grey30", lineheight = 1.75),
    legend.text = element_text(size = 9, color = "grey40"),
    plot.title = element_text(size = 12, hjust = 0.5, color = "grey30"),
    axis.title.x = ggtext::element_markdown(size = 12, color = "grey30", margin = margin(t = 10)),
    axis.title.y = ggtext::element_markdown(size = 12, color = "grey30", margin = margin(r = 10)),
    axis.text = element_text(size = 10),
    panel.background = element_rect(fill = "white", color = "white"),
    plot.background = element_rect(fill = "white", color = "white"),
    panel.grid.major = element_blank(),
    panel.grid.minor = element_blank(),
    axis.line = element_line(color = "grey40", linewidth = 0.9),
    panel.border = element_rect(color = "grey40", fill = NA, size = 0.9),
    plot.margin = unit(c(0.5, 0.5, 0.5, 0.5), "cm")
  )

# Print plot
print(plot)

# Save PNG with exact name
ggsave("3_L1_Insertions_PC_vs_Longevity_N.png", plot, width = 10, height = 8, dpi = 800)
# Save RDS
saveRDS(plot, "3_L1_Insertions_PC_vs_Longevity_N.rds")


################################################################################
### L1 and SINEs Insertions within PC Genes vs Longevity
################################################################################

# Store Current model and Response variable in temp vars 
Current_model <- LL1SINEsIntersection
Current_Response <- mydata$maximum_longevity_m

# Calculate fitted values
fitted_values <- predict(Current_model)
mydata$nudge_y <- ifelse(Current_Response > fitted_values, 0.02, -0.02)

# Classify orders
mydata$color_group <- ifelse(mydata$Order %in% c("Rodentia", "Artiodactyla", "Carnivora", "Chiroptera", "Primates"), 
                             mydata$Order, 
                             "Other Orders")

# Clean species names
mydata$species_clean <- gsub("_", " ", mydata$species)

# Dummy variable for legend formatting
mydata$ModelStats <- "Model Stats"

# Final Plot
plot <- ggplot(mydata, aes(x = Transformed_L1_SINEs_PC_Intersections, y = maximum_longevity_m)) +
  
  # Colored dots
  geom_point(aes(color = color_group), size = 3, alpha = 0.9) +
  
  # Species labels (all species)
  geom_text_repel(
    data = mydata,
    aes(label = Common_Name),
    size = 2.5,
    color = "grey20",
    box.padding = 0.3,
    point.padding = 0.8,
    segment.color = "grey60",
    segment.size = 0.3,
    segment.curvature = 0,
    segment.angle = 20,
    min.segment.length = 0.01,
    max.overlaps = Inf
  ) +
  
  # Regression line
  geom_smooth(method = "lm", se = FALSE, color = "grey50", alpha = 0.7, 
              fullrange = TRUE, linewidth = 0.9) +
  
  # Axis labels
  xlab("Number of L1 and SINEs Insertions within PC Genes")+
  ylab("Longevity")+
  
  # Color legend with model stats
  scale_color_manual(
    name = paste0(
      "<b style='font-size:11pt;'>Model Stats</b><br>",
      nrow(mydata), " species<br>",
      "Corr = ", corrInt2, "<br>",
      "p value = ", LL1SINEsIntersection_pvalue, "<br>",
      #"p value = ", LL1SINEsIntersection_pvalue, "<span style='color:red;'><b>*</b></span><br>",
      "R² = ", RelLL1SINEsIntersection, "<br>",
      "λ = ", lambdaLL1SINEsIntersection, "<br><br>",
      "<b style='font-size:11pt;'>Orders</b>"
    ),
    values = order_colors
  ) +
  
  # Axis limits
  scale_x_continuous(limits = c(-0.1, 1.05), expand = c(0, 0)) +
  scale_y_continuous(limits = c(-0.1, 1.05), expand = c(0, 0)) +
  
  # Theme
  theme_minimal(base_size = 6) +
  theme(
    legend.position = "right",
    legend.justification = "bottom",
    legend.direction = "vertical",
    legend.title = element_markdown(size = 10, color = "grey30", lineheight = 1.75),
    legend.text = element_text(size = 9, color = "grey40"),
    plot.title = element_text(size = 12, hjust = 0.5, color = "grey30"),
    axis.title.x = ggtext::element_markdown(size = 12, color = "grey30", margin = margin(t = 10)),
    axis.title.y = ggtext::element_markdown(size = 12, color = "grey30", margin = margin(r = 10)),
    axis.text = element_text(size = 10),
    panel.background = element_rect(fill = "white", color = "white"),
    plot.background = element_rect(fill = "white", color = "white"),
    panel.grid.major = element_blank(),
    panel.grid.minor = element_blank(),
    axis.line = element_line(color = "grey40", linewidth = 0.9),
    panel.border = element_rect(color = "grey40", fill = NA, size = 0.9),
    plot.margin = unit(c(0.5, 0.5, 0.5, 0.5), "cm")
  )

# Print plot
print(plot)

# Save PNG with exact name
ggsave("4_L1_SINEs_Insertions_PC_vs_Longevity_N.png", plot, width = 10, height = 8, dpi = 800)
# Save RDS
saveRDS(plot, "4_L1_SINEs_Insertions_PC_vs_Longevity_N.rds")


################################################################################
### L1 Counts vs Fusion Gene Counts
################################################################################

# Store Current model and Response variable in temp vars 
Current_model <- FL1
Current_Response <- mydata$Fusion

# Calculate fitted values
fitted_values <- predict(Current_model)
mydata$nudge_y <- ifelse(Current_Response > fitted_values, 0.02, -0.02)

# Classify orders
mydata$color_group <- ifelse(mydata$Order %in% c("Rodentia", "Artiodactyla", "Carnivora", "Chiroptera", "Primates"), 
                             mydata$Order, 
                             "Other Orders")

# Clean species names
mydata$species_clean <- gsub("_", " ", mydata$species)

# Dummy variable for legend formatting
mydata$ModelStats <- "Model Stats"

# Final Plot
plot <- ggplot(mydata, aes(x = Transformed_L1_Counts, y = Fusion)) +
  
  # Colored dots
  geom_point(aes(color = color_group), size = 3, alpha = 0.9) +
  
  # Species labels (all species)
  geom_text_repel(
    data = mydata,
    aes(label = Common_Name),
    size = 2.5,
    color = "grey20",
    box.padding = 0.3,
    point.padding = 0.8,
    segment.color = "grey60",
    segment.size = 0.3,
    segment.curvature = 0,
    segment.angle = 20,
    min.segment.length = 0.01,
    max.overlaps = Inf
  ) +
  
  # Regression line
  geom_smooth(method = "lm", se = FALSE, color = "grey50", alpha = 0.7, 
              fullrange = TRUE, linewidth = 0.9) +
  
  # Axis labels
  xlab("Number of Active L1")+
  ylab("Number of Fusion Cancer Gene Orthologs")+
  
  # Color legend with model stats
  scale_color_manual(
    name = paste0(
      "<b style='font-size:11pt;'>Model Stats</b><br>",
      nrow(mydata), " species<br>",
      "Corr = ", corrFus, "<br>",
      #"p value = ", FL1_pvalue, "<br>",
      "p value = ", FL1_pvalue, "<span style='color:red;'><b>*</b></span><br>",
      "R² = ", RelFL1, "<br>",
      "λ = ", lambdaFL1, "<br><br>",
      "<b style='font-size:11pt;'>Orders</b>"
    ),
    values = order_colors
  ) +
  
  # Axis limits
  scale_x_continuous(limits = c(-0.1, 1.05), expand = c(0, 0)) +
  scale_y_continuous(limits = c(-0.1, 1.05), expand = c(0, 0)) +
  
  # Theme
  theme_minimal(base_size = 6) +
  theme(
    legend.position = "right",
    legend.justification = "bottom",
    legend.direction = "vertical",
    legend.title = element_markdown(size = 10, color = "grey30", lineheight = 1.75),
    legend.text = element_text(size = 9, color = "grey40"),
    plot.title = element_text(size = 12, hjust = 0.5, color = "grey30"),
    axis.title.x = ggtext::element_markdown(size = 12, color = "grey30", margin = margin(t = 10)),
    axis.title.y = ggtext::element_markdown(size = 12, color = "grey30", margin = margin(r = 10)),
    axis.text = element_text(size = 10),
    panel.background = element_rect(fill = "white", color = "white"),
    plot.background = element_rect(fill = "white", color = "white"),
    panel.grid.major = element_blank(),
    panel.grid.minor = element_blank(),
    axis.line = element_line(color = "grey40", linewidth = 0.9),
    panel.border = element_rect(color = "grey40", fill = NA, size = 0.9),
    plot.margin = unit(c(0.5, 0.5, 0.5, 0.5), "cm")
  )

# Print plot
print(plot)

# Save PNG with exact name
ggsave("5_L1_Counts_vs_Fusion_N.png", plot, width = 10, height = 8, dpi = 800)
# Save RDS
saveRDS(plot, "5_L1_Counts_vs_Fusion_N.rds")


################################################################################
### L1 and SINE Counts vs Fusion Gene Counts
################################################################################

# Store Current model and Response variable in temp vars 
Current_model <- FL1SINEs
Current_Response <- mydata$Fusion

# Calculate fitted values
fitted_values <- predict(Current_model)
mydata$nudge_y <- ifelse(Current_Response > fitted_values, 0.02, -0.02)

# Classify orders
mydata$color_group <- ifelse(mydata$Order %in% c("Rodentia", "Artiodactyla", "Carnivora", "Chiroptera", "Primates"), 
                             mydata$Order, 
                             "Other Orders")

# Clean species names
mydata$species_clean <- gsub("_", " ", mydata$species)

# Dummy variable for legend formatting
mydata$ModelStats <- "Model Stats"

# Final Plot
plot <- ggplot(mydata, aes(x = Transformed_L1_SINEs_Counts, y = Fusion)) +
  
  # Colored dots
  geom_point(aes(color = color_group), size = 3, alpha = 0.9) +
  
  # Species labels (all species)
  geom_text_repel(
    data = mydata,
    aes(label = Common_Name),
    size = 2.5,
    color = "grey20",
    box.padding = 0.3,
    point.padding = 0.8,
    segment.color = "grey60",
    segment.size = 0.3,
    segment.curvature = 0,
    segment.angle = 20,
    min.segment.length = 0.01,
    max.overlaps = Inf
  ) +
  
  # Regression line
  geom_smooth(method = "lm", se = FALSE, color = "grey50", alpha = 0.7, 
              fullrange = TRUE, linewidth = 0.9) +
  
  # Axis labels
  xlab("Number of Active L1 and SINEs")+
  ylab("Number of Fusion Cancer Gene Orthologs")+
  
  # Color legend with model stats
  scale_color_manual(
    name = paste0(
      "<b style='font-size:11pt;'>Model Stats</b><br>",
      nrow(mydata), " species<br>",
      "Corr = ", corrFus2, "<br>",
      "p value = ", FL1SINEs_pvalue, "<br>",
      #"p value = ", FL1SINEs_pvalue, "<span style='color:red;'><b>*</b></span><br>",
      "R² = ", RelFL1SINEs, "<br>",
      "λ = ", lambdaFL1SINEs, "<br><br>",
      "<b style='font-size:11pt;'>Orders</b>"
    ),
    values = order_colors
  ) +
  
  # Axis limits
  scale_x_continuous(limits = c(-0.1, 1.05), expand = c(0, 0)) +
  scale_y_continuous(limits = c(-0.1, 1.05), expand = c(0, 0)) +
  
  # Theme
  theme_minimal(base_size = 6) +
  theme(
    legend.position = "right",
    legend.justification = "bottom",
    legend.direction = "vertical",
    legend.title = element_markdown(size = 10, color = "grey30", lineheight = 1.75),
    legend.text = element_text(size = 9, color = "grey40"),
    plot.title = element_text(size = 12, hjust = 0.5, color = "grey30"),
    axis.title.x = ggtext::element_markdown(size = 12, color = "grey30", margin = margin(t = 10)),
    axis.title.y = ggtext::element_markdown(size = 12, color = "grey30", margin = margin(r = 10)),
    axis.text = element_text(size = 10),
    panel.background = element_rect(fill = "white", color = "white"),
    plot.background = element_rect(fill = "white", color = "white"),
    panel.grid.major = element_blank(),
    panel.grid.minor = element_blank(),
    axis.line = element_line(color = "grey40", linewidth = 0.9),
    panel.border = element_rect(color = "grey40", fill = NA, size = 0.9),
    plot.margin = unit(c(0.5, 0.5, 0.5, 0.5), "cm")
  )

# Print plot
print(plot)

# Save PNG with exact name
ggsave("6_L1_SINEs_Counts_vs_Fusion_N.png", plot, width = 10, height = 8, dpi = 800)
# Save RDS
saveRDS(plot, "6_L1_SINEs_Counts_vs_Fusion_N.rds")


################################################################################
### L1 Counts vs L1 Insertions within PC Genes
################################################################################

# Store Current model and Response variable in temp vars 
Current_model <- L1L1PC
Current_Response <- mydata$Transformed_L1_PC_Intersections

# Calculate fitted values
fitted_values <- predict(Current_model)
mydata$nudge_y <- ifelse(Current_Response > fitted_values, 0.02, -0.02)

# Classify orders
mydata$color_group <- ifelse(mydata$Order %in% c("Rodentia", "Artiodactyla", "Carnivora", "Chiroptera", "Primates"), 
                             mydata$Order, 
                             "Other Orders")

# Clean species names
mydata$species_clean <- gsub("_", " ", mydata$species)

# Dummy variable for legend formatting
mydata$ModelStats <- "Model Stats"

# Final Plot
plot <- ggplot(mydata, aes(x = Transformed_L1_Counts, y = Transformed_L1_PC_Intersections)) +
  
  # Colored dots
  geom_point(aes(color = color_group), size = 3, alpha = 0.9) +
  
  # Species labels (all species)
  geom_text_repel(
    data = mydata,
    aes(label = Common_Name),
    size = 2.5,
    color = "grey20",
    box.padding = 0.3,
    point.padding = 0.8,
    segment.color = "grey60",
    segment.size = 0.3,
    segment.curvature = 0,
    segment.angle = 20,
    min.segment.length = 0.01,
    max.overlaps = Inf
  ) +
  
  # Regression line
  geom_smooth(method = "lm", se = FALSE, color = "grey50", alpha = 0.7, 
              fullrange = TRUE, linewidth = 0.9) +
  
  # Axis labels
  xlab("Number of Active L1")+#&#8201;<span style='font-size:8pt;'>(Tukey-Transformed)</span>") +
  ylab("Number of L1 Insertions within PC Genes")+#&#8201;<span style='font-size:8pt;'>(Tukey-Transformed)</span>") +
  
  # Color legend with model stats
  scale_color_manual(
    name = paste0(
      "<b style='font-size:11pt;'>Model Stats</b><br>",
      nrow(mydata), " species<br>",
      "Corr = ", corr1, "<br>",
      #"p value = ", L1L1PC_pvalue, "<br>",
      "p value = ", L1L1PC_pvalue, "<span style='color:red;'><b>*</b></span><br>",
      "R² = ", RelL1L1PC, "<br>",
      "λ = ", lambdaL1L1PC, "<br><br>",
      "<b style='font-size:11pt;'>Orders</b>"
    ),
    values = order_colors
  ) +
  
  # Axis limits
  scale_x_continuous(limits = c(-0.1, 1.05), expand = c(0, 0)) +
  scale_y_continuous(limits = c(-0.1, 1.05), expand = c(0, 0)) +
  
  # Theme
  theme_minimal(base_size = 6) +
  theme(
    legend.position = "right",
    legend.justification = "bottom",
    legend.direction = "vertical",
    legend.title = element_markdown(size = 10, color = "grey30", lineheight = 1.75),
    legend.text = element_text(size = 9, color = "grey40"),
    plot.title = element_text(size = 12, hjust = 0.5, color = "grey30"),
    axis.title.x = ggtext::element_markdown(size = 12, color = "grey30", margin = margin(t = 10)),
    axis.title.y = ggtext::element_markdown(size = 12, color = "grey30", margin = margin(r = 10)),
    axis.text = element_text(size = 10),
    panel.background = element_rect(fill = "white", color = "white"),
    plot.background = element_rect(fill = "white", color = "white"),
    panel.grid.major = element_blank(),
    panel.grid.minor = element_blank(),
    axis.line = element_line(color = "grey40", linewidth = 0.9),
    panel.border = element_rect(color = "grey40", fill = NA, size = 0.9),
    plot.margin = unit(c(0.5, 0.5, 0.5, 0.5), "cm")
  )

# Print plot
print(plot)

# Save PNG with exact name
ggsave("7_L1_Counts_vs_L1_Insertions_within_PC_Genes_N.png", plot, width = 10, height = 8, dpi = 800)
# Save RDS
saveRDS(plot, "7_L1_Counts_vs_L1_Insertions_within_PC_Genes_N.rds")


################################################################################
### L1 and SINE Counts vs L1 and SINE Insertions within PC Genes
################################################################################

# Store Current model and Response variable in temp vars 
Current_model <- L1SINEsL1SINEsPC
Current_Response <- mydata$Transformed_L1_SINEs_PC_Intersections

# Calculate fitted values
fitted_values <- predict(Current_model)
mydata$nudge_y <- ifelse(Current_Response > fitted_values, 0.02, -0.02)

# Classify orders
mydata$color_group <- ifelse(mydata$Order %in% c("Rodentia", "Artiodactyla", "Carnivora", "Chiroptera", "Primates"), 
                             mydata$Order, 
                             "Other Orders")

# Clean species names
mydata$species_clean <- gsub("_", " ", mydata$species)

# Dummy variable for legend formatting
mydata$ModelStats <- "Model Stats"

# Final Plot
plot <- ggplot(mydata, aes(x = Transformed_L1_SINEs_Counts, y = Transformed_L1_SINEs_PC_Intersections)) +
  
  # Colored dots
  geom_point(aes(color = color_group), size = 3, alpha = 0.9) +
  
  # Species labels (all species)
  geom_text_repel(
    data = mydata,
    aes(label = Common_Name),
    size = 2.5,
    color = "grey20",
    box.padding = 0.3,
    point.padding = 0.8,
    segment.color = "grey60",
    segment.size = 0.3,
    segment.curvature = 0,
    segment.angle = 20,
    min.segment.length = 0.01,
    max.overlaps = Inf
  ) +
  
  # Regression line
  geom_smooth(method = "lm", se = FALSE, color = "grey50", alpha = 0.7, 
              fullrange = TRUE, linewidth = 0.9) +
  
  # Axis labels
  xlab("Number of Active L1 and SINEs")+
  ylab("Number of L1 and SINE Insertions within PC Genes")+
  
  # Color legend with model stats
  scale_color_manual(
    name = paste0(
      "<b style='font-size:11pt;'>Model Stats</b><br>",
      nrow(mydata), " species<br>",
      "Corr = ", corr2, "<br>",
      #"p value = ", L1SINEsL1SINEsPC_pvalue, "<br>",
      "p value = ", L1SINEsL1SINEsPC_pvalue, "<span style='color:red;'><b>*</b></span><br>",
      "R² = ", RelL1SINEsL1SINEsPC, "<br>",
      "λ = ", lambdaL1SINEsL1SINEsPC, "<br><br>",
      "<b style='font-size:11pt;'>Orders</b>"
    ),
    values = order_colors
  ) +
  
  # Axis limits
  scale_x_continuous(limits = c(-0.1, 1.05), expand = c(0, 0)) +
  scale_y_continuous(limits = c(-0.1, 1.05), expand = c(0, 0)) +
  
  # Theme
  theme_minimal(base_size = 6) +
  theme(
    legend.position = "right",
    legend.justification = "bottom",
    legend.direction = "vertical",
    legend.title = element_markdown(size = 10, color = "grey30", lineheight = 1.75),
    legend.text = element_text(size = 9, color = "grey40"),
    plot.title = element_text(size = 12, hjust = 0.5, color = "grey30"),
    axis.title.x = ggtext::element_markdown(size = 12, color = "grey30", margin = margin(t = 10)),
    axis.title.y = ggtext::element_markdown(size = 12, color = "grey30", margin = margin(r = 10)),
    axis.text = element_text(size = 10),
    panel.background = element_rect(fill = "white", color = "white"),
    plot.background = element_rect(fill = "white", color = "white"),
    panel.grid.major = element_blank(),
    panel.grid.minor = element_blank(),
    axis.line = element_line(color = "grey40", linewidth = 0.9),
    panel.border = element_rect(color = "grey40", fill = NA, size = 0.9),
    plot.margin = unit(c(0.5, 0.5, 0.5, 0.5), "cm")
  )

# Print plot
print(plot)

# Save PNG with exact name
ggsave("8_L1_SINE_Counts_vs_L1_SINE_Insertions_within_PC_Genes_N.png", plot, width = 10, height = 8, dpi = 800)
# Save RDS
saveRDS(plot, "8_L1_SINE_Counts_vs_L1_SINE_Insertions_within_PC_Genes_N.rds")


################################################################################
### TSG vs Oncogenes
################################################################################

# Store Current model and Response variable in temp vars 
Current_model <- TSGOnco
Current_Response <- mydata$TSG

# Calculate fitted values
fitted_values <- predict(Current_model)
mydata$nudge_y <- ifelse(Current_Response > fitted_values, 0.02, -0.02)

# Classify orders
mydata$color_group <- ifelse(mydata$Order %in% c("Rodentia", "Artiodactyla", "Carnivora", "Chiroptera", "Primates"), 
                             mydata$Order, 
                             "Other Orders")

# Clean species names
mydata$species_clean <- gsub("_", " ", mydata$species)

# Dummy variable for legend formatting
mydata$ModelStats <- "Model Stats"

# Final Plot
plot <- ggplot(mydata, aes(x = Oncogene, y = TSG)) +
  
  # Colored dots
  geom_point(aes(color = color_group), size = 3, alpha = 0.9) +
  
  # Species labels (all species)
  geom_text_repel(
    data = mydata,
    aes(label = Common_Name),
    size = 2.5,
    color = "grey20",
    box.padding = 0.3,
    point.padding = 0.8,
    segment.color = "grey60",
    segment.size = 0.3,
    segment.curvature = 0,
    segment.angle = 20,
    min.segment.length = 0.01,
    max.overlaps = Inf
  ) +
  
  # Regression line
  geom_smooth(method = "lm", se = FALSE, color = "grey50", alpha = 0.7, 
              fullrange = TRUE, linewidth = 0.9) +
  
  # Axis labels
  xlab("Number of Oncogenes") +
  ylab("Number of TSGs") +
  
  
  # Color legend with model stats
  scale_color_manual(
    name = paste0(
      "<b style='font-size:11pt;'>Model Stats</b><br>",
      nrow(mydata), " species<br>",
      "Corr = ", corrTSGOnco, "<br>",
      #"p value = ", TSGOnco_pvalue, "<br>",
      "p value = ", TSGOnco_pvalue, "<span style='color:red;'><b>*</b></span><br>",
      "R² = ", RelTSGOnco, "<br>",
      "λ = ", lambdaTSGOnco, "<br><br>",
      "<b style='font-size:11pt;'>Orders</b>"
    ),
    values = order_colors
  ) +
  
  # Axis limits
  scale_x_continuous(limits = c(-0.1, 1.05), expand = c(0, 0)) +
  scale_y_continuous(limits = c(-0.1, 1.05), expand = c(0, 0)) +
  
  # Theme
  theme_minimal(base_size = 6) +
  theme(
    legend.position = "right",
    legend.justification = "bottom",
    legend.direction = "vertical",
    legend.title = element_markdown(size = 10, color = "grey30", lineheight = 1.75),
    legend.text = element_text(size = 9, color = "grey40"),
    plot.title = element_text(size = 12, hjust = 0.5, color = "grey30"),
    axis.title.x = ggtext::element_markdown(size = 12, color = "grey30", margin = margin(t = 10)),
    axis.title.y = ggtext::element_markdown(size = 12, color = "grey30", margin = margin(r = 10)),
    axis.text = element_text(size = 10),
    panel.background = element_rect(fill = "white", color = "white"),
    plot.background = element_rect(fill = "white", color = "white"),
    panel.grid.major = element_blank(),
    panel.grid.minor = element_blank(),
    axis.line = element_line(color = "grey40", linewidth = 0.9),
    panel.border = element_rect(color = "grey40", fill = NA, size = 0.9),
    plot.margin = unit(c(0.5, 0.5, 0.5, 0.5), "cm")
  )

# Print plot
print(plot)

# Save PNG with exact name
ggsave("11_TSG_Oncogene.png", plot, width = 10, height = 8, dpi = 800)
# Save RDS
saveRDS(plot, "11_TSG_Oncogene.rds")
```
