## Supplementary Scripts for "Recent Non-LTR Retrotransposon Activity Predicts Cancer Prevalence in Mammals": 10_Model Comparisons.R

```
################################################################################
###### Comparing Different Regressions via SEy
###
################################################################################
############ Univariate Regression Statistical Models
################################################################################
###  Loading required packages
#library(caper)

# Load the data and prepare comparative data object
comparative_data <- comparative.data(phy = tre, data = mydata, 
                                     names.col = "species", # replace with your species column name
                                     vcv = TRUE, na.omit = FALSE)

############ comparing lmbda=0, calcuated Lambda, and lambda=1

############ Neoplasia
# Fit a model with lambda fixed at 0 (no phylogenetic signal)
model_lambda_0 <- lm(NeoplasiaPrevalence ~ Transformed_L1_Counts, data = mydata)

# Fit a model with calculated lambda
model_lambda_1 <- ML1

# Fit a model with lambda fixed at 1 (full phylogenetic signal)
model_lambda_2 <- pgls(NeoplasiaPrevalence ~ Transformed_L1_Counts, 
                       data = comparative_data, 
                       lambda = 1.00)
# Compare AIC values
cat(
  round(AIC(model_lambda_0), 2), "\t",
  round(AIC(model_lambda_1), 2), "\t",
  round(AIC(model_lambda_2), 2), "\n"
)
cat("AIC with Lambda = 0:", round(AIC(model_lambda_0),2), "\n")
cat("AIC with Estimated Lambda:", round(AIC(model_lambda_1),2), "\n")
cat("AIC with Lambda = 1:", round(AIC(model_lambda_2),2), "\n")

############ Malignancy
# Fit a model with lambda fixed at 0 (no phylogenetic signal)
model_lambda_0 <- lm(MalignancyPrevalence ~ Transformed_L1_SINEs_Counts, data = mydata)

# Fit a model with calculated lambda
model_lambda_1 <- ML1SINEs

# Fit a model with lambda fixed at 1 (full phylogenetic signal)
model_lambda_2 <- pgls(MalignancyPrevalence ~ Transformed_L1_SINEs_Counts, 
                       data = comparative_data, 
                       lambda = 1.00)

# Compare AIC values
cat(
  round(AIC(model_lambda_0), 2), "\t",
  round(AIC(model_lambda_1), 2), "\t",
  round(AIC(model_lambda_2), 2), "\n"
)

################################################################################
############ Multivariate Regression Statistical Models
################################################################################
### Neoplasia

# Fit a model with lambda fixed at 0 (no phylogenetic signal)
model_lambda_0 <- lm(NeoplasiaPrevalence ~ Transformed_L1_Counts, data = mydata)

# Fit a model with calculated lambda
model_lambda_1 <- ML1

# Fit a model with lambda fixed at 1 (full phylogenetic signal)
model_lambda_2 <- pgls(NeoplasiaPrevalence ~ Transformed_L1_Counts, 
                       data = comparative_data, 
                       lambda = 1.00)
# Compare AIC values
cat(
  round(AIC(model_lambda_0), 2), "\t",
  round(AIC(model_lambda_1), 2), "\t",
  round(AIC(model_lambda_2), 2), "\n"
)
#cat("AIC with Lambda = 0:", round(AIC(model_lambda_0),2), "\n")
#cat("AIC with Estimated Lambda:", round(AIC(model_lambda_1),2), "\n")
#cat("AIC with Lambda = 0:", round(AIC(model_lambda_2),2), "\n")

### Malignancy

# Fit a model with lambda fixed at 0 (no phylogenetic signal)
model_lambda_0 <- lm(MalignancyPrevalence ~ Transformed_L1_SINEs_Counts, data = mydata)

# Fit a model with calculated lambda
model_lambda_1 <- ML1SINEs

# Fit a model with lambda fixed at 1 (full phylogenetic signal)
model_lambda_2 <- pgls(MalignancyPrevalence ~ Transformed_L1_SINEs_Counts, 
                       data = comparative_data, 
                       lambda = 1.00)

# Compare AIC values
cat(
  round(AIC(model_lambda_0), 2), "\t",
  round(AIC(model_lambda_1), 2), "\t",
  round(AIC(model_lambda_2), 2), "\n"
   )
```
