## Supplementary Scripts for "Recent Non-LTR Retrotransposon Activity Predicts Cancer Prevalence in Mammals": 12 - phylogeny plot.R

```
################################################
######### Plotting the Phylogeny

library(ggtree)
library(ggplot2)
library(dplyr)
library(cowplot)
library(grid) # Needed for using grid functionality with annotation_custom
library(ggtext)

# Step 1: Transform and Normalize Cancer Data
mydata$NeoplasiaPrevalence <- transformTukey(mydata$NeoplasiaPrevalence)
mydata$NeoplasiaPrevalence <- round((mydata$NeoplasiaPrevalence - min(mydata$NeoplasiaPrevalence)) / (max(mydata$NeoplasiaPrevalence) - min(mydata$NeoplasiaPrevalence)),2) 

mydata$MalignancyPrevalence <- transformTukey(mydata$MalignancyPrevalence)
mydata$MalignancyPrevalence <- round((mydata$MalignancyPrevalence - min(mydata$MalignancyPrevalence)) / (max(mydata$MalignancyPrevalence) - min(mydata$MalignancyPrevalence)),2) 


# Step 2: Transform and Normalize L1 and SINEs
mydata$Autonomous <- mydata$L1_Counts #
mydata$Autonomous <- round((transformTukey(mydata$Autonomous)),2)
mydata$Autonomous <- round((mydata$Autonomous - min(mydata$Autonomous)) / (max(mydata$Autonomous) - min(mydata$Autonomous)),2) 

mydata$Non_Autonomous <- mydata$SINEs_Counts
mydata$Non_Autonomous <- round((transformTukey(mydata$Non_Autonomous)) + 1, 2) # Adding a constant value to make the data points positive
mydata$Non_Autonomous <- round((mydata$Non_Autonomous - min(mydata$Non_Autonomous)) / (max(mydata$Non_Autonomous) - min(mydata$Non_Autonomous)),2) 

# Ensure species names in your data frame match the tree's tip labels (convert spaces to underscores)
mydata$species <- gsub(" ", "_", mydata$species)

# Step 2: Filter data to include only species present in the tree
species_in_tree <- tre$tip.label
filtered_data <- mydata %>% filter(species %in% species_in_tree)

# Step 3: Reorder filtered data to match the order of tree tip labels
filtered_data <- filtered_data %>% 
  mutate(species = factor(species, levels = species_in_tree)) %>% 
  arrange(species)

# Define bar_data based on filtered_data
bar_data <- filtered_data

# Ensure the "Order" column is a factor with correct levels
bar_data <- bar_data %>%
  mutate(Order = ifelse(Order %in% c("Carnivora", "Artiodactyla", "Primates", "Rodentia", "Chiroptera"), Order, "Other Orders")) %>%
  mutate(Order = factor(Order, levels = c("Carnivora", "Artiodactyla", "Primates", "Rodentia", "Chiroptera", "Other Orders")))

# Define the custom color palette for the Orders
order_colors <- c("Carnivora" = "#414487FF",
                  "Artiodactyla" = "#2A788EFF",
                  "Primates" = "#440154FF",
                  "Rodentia" = "#FCE205",
                  "Chiroptera" = "#22A884FF", 
                  "Other Orders" = "#7AD151FF")


# Use a distinct dataset for the legend (this step ensures the legend has unique categories)
legend_data <- bar_data %>% distinct(Order)

# Step 4: Create the tree plot
# ladderize = FALSE, 
tree_plot <- ggtree(tre, ladderize = TRUE, layout = "rectangular", size = 0.7, color = "grey20") %<+% bar_data +  
  geom_tiplab(aes(label = gsub("_", " ", label), color = Order), offset = 75, size = 2.5, 
                          hjust = TRUE, show.legend = FALSE, fontface = "italic") +
  xlim(0, 440) +  
  scale_color_manual(values = order_colors, na.translate = FALSE) +  
  geom_point(data = legend_data, aes(x = Inf, y = Inf, color = Order), size = 0, show.legend = TRUE) +  
  theme(
    legend.position = c(0.07, 1.01),
    legend.justification = c(0.1, 0.3),
    legend.direction = "horizontal",  
    legend.text = element_text(size = 8, color = "grey20"),
    legend.title = element_text(size = 10, face = "bold", color = "grey20", vjust = 1),  # Adjust title size if needed
    legend.box.margin = margin(t = 0, r = 0, b = 0, l = 0),
    plot.margin = margin(t = 10, r = 10, b = 10, l = 10)
  ) +
  guides(color = guide_legend(override.aes = list(size = 3.5, shape = 15))) +
  labs(color = "A")  # Add your custom title here

# Define the end point of all branches (maximum distance to the root)
branch_end <- max(tree_plot$data$x, na.rm = TRUE)

# Add secondary x-axis with reversed values
tree_plot <- tree_plot + 
  annotate("segment", x = 0, xend = branch_end, y = -1, yend = -1, color = "grey20", size = 0.8) +
  annotate("text", x = seq(0, branch_end, length.out = 6), y = -2, 
           label = rev(seq(0, branch_end, length.out = 6)), color = "grey20", size = 3) +
  annotate("segment", x = seq(0, branch_end, length.out = 6), xend = seq(0, branch_end, length.out = 6), 
           y = -1, yend = -1.4, color = "grey20", size = 0.8) +
  annotate("text", x = (branch_end / 2), y = -4, label = "Millions of Years Ago", size = 3.5, 
           color =  "grey20", hjust = 0.5) +
  coord_cartesian(clip = "off") +
  theme(
    plot.margin = margin(t = 50, r = 1, b = 10, l = 1),
    axis.line.x.bottom = element_blank(),
    axis.ticks.x.bottom = element_blank()
  )


# Extract the new label order from the ladderized tree
new_order <- tree_plot$data %>%
  filter(isTip) %>%
  arrange(y) %>%  # Arrange by the y-position to get the order from top to bottom
  pull(label)

# Reorder `bar_data` to match the ladderized tree order
bar_data <- bar_data %>%
  mutate(species = factor(species, levels = new_order)) %>%
  arrange(species)

# Function to create bar plot grobs for given variables with specified y-limits
create_bar_grob <- function(data, y_var, order_colors, y_limits = NULL) {
  bar_plot <- ggplot(data, aes(y = species, x = !!sym(y_var), fill = Order)) +
    geom_bar(stat = "identity", width = 0.6) +
    scale_fill_manual(values = order_colors, na.translate = FALSE) +
    theme_void() +  # Remove all axes, gridlines, and background
    theme(legend.position = "none")
  
  if (!is.null(y_limits)) {
    bar_plot <- bar_plot + coord_cartesian(xlim = y_limits)
  }
  
  return(ggplotGrob(bar_plot))
}

#### To adjust the limits for ys (actually xs :))
#y_limits_vector <- c(0.8, 0.8, max(bar_data$Autonomous), max(bar_data$Non_Autonomous))
y_limits_vector <- c(1, 1, 1, 1)


# Step 5: Create grobs for all bar plots with individual y-limits
bar_grob_neoplasia <- create_bar_grob(bar_data, "NeoplasiaPrevalence", order_colors, y_limits = c(0, y_limits_vector[1]))
bar_grob_malignancy <- create_bar_grob(bar_data, "MalignancyPrevalence", order_colors, y_limits = c(0, y_limits_vector[2]))
bar_grob_autonomous <- create_bar_grob(bar_data, "Autonomous", order_colors, y_limits = c(0, y_limits_vector[3]))
bar_grob_Non_Autonomous <- create_bar_grob(bar_data, "Non_Autonomous", order_colors, y_limits = c(0, y_limits_vector[4]))

# Step 6: Add an adjustable offset between the tree and bars
offset_value <- 80  # Adjust this value to control the distance between the bars and the tree labels
bar_width <- 45  # Width of each bar plot
bar_spacing <- 6  # Space between each bar plot
y_offset <- 0.3  #To make the bars stand right in front of the branches/labels

# Step 7: Combine the tree plot and the bar plots using annotation_custom
tree_plot_with_bars <- tree_plot + 
  annotation_custom(grob = bar_grob_neoplasia, 
                    xmin = max(tree_plot$data$x) + offset_value,  
                    xmax = max(tree_plot$data$x) + offset_value + bar_width,  
                    ymin = 0 + y_offset, ymax = length(tre$tip.label) + y_offset) +
  annotation_custom(grob = bar_grob_malignancy, 
                    xmin = max(tree_plot$data$x) + offset_value + bar_width + bar_spacing,  
                    xmax = max(tree_plot$data$x) + offset_value + 2 * bar_width + bar_spacing,  
                    ymin = 0 + y_offset, ymax = length(tre$tip.label) + y_offset) +
  annotation_custom(grob = bar_grob_autonomous, 
                    xmin = max(tree_plot$data$x) + offset_value + 2 * (bar_width + bar_spacing),  
                    xmax = max(tree_plot$data$x) + offset_value + 3 * bar_width + 2 * bar_spacing,  
                    ymin = 0 + y_offset, ymax = length(tre$tip.label) + y_offset) +
  annotation_custom(grob = bar_grob_Non_Autonomous, 
                    xmin = max(tree_plot$data$x) + offset_value + 3 * (bar_width + bar_spacing),  
                    xmax = max(tree_plot$data$x) + offset_value + 4 * bar_width + 3 * bar_spacing,  
                    ymin = 0 + y_offset, ymax = length(tre$tip.label) + y_offset) +
  coord_cartesian(clip = "off")

# Step 9: Add the x-axis for each bar plot manually
bar_variables <- c("NeoplasiaPrevalence", "MalignancyPrevalence", "Autonomous", "Non_Autonomous")

# Step 9: Add the x-axis for each bar plot manually with vertical numbers
for (i in seq_along(bar_variables)) {
  bar_max_value <- max(bar_data[[bar_variables[i]]], na.rm = TRUE)  # Get the maximum value of the current variable
  x_start <- max(tree_plot$data$x) + offset_value + (i - 1) * (bar_width + bar_spacing)
  x_end <- x_start + bar_width
  
  tree_plot_with_bars <- tree_plot_with_bars +
    # Add the main axis line for the bar plot
    annotate("segment", x = x_start,  # Starting x-position
             xend = x_end,  # Ending x-position (bar plot width)
             y = -1, yend = -1,  # Y-position (axis line placement)
             color = "grey20", size = 0.7) +  # Line color and thickness
    
    # Add the tick labels (variable values) with vertical alignment
    annotate("text", x = seq(x_start, x_end, length.out = 6),  # X-positions for the labels
             y = -2.5,  # Y-position for the labels (below the axis line)
             label = round(seq(0, y_limits_vector[i], length.out = 6), 1),  # Correctly use the maximum value for each bar
             size = 2.5,  # Font size of the labels
             angle = 90,  # Set angle to 90 to make labels vertical
             vjust = 0.5,
             color = "grey20") +  # Adjust vertical alignment of the labels
    
    # Add the tick marks for the bar plot x-axis
    annotate("segment", x = seq(x_start, x_end, length.out = 6),  # X-positions for the tick marks
             xend = seq(x_start, x_end, length.out = 6),  # End points for the tick marks (same as start x)
             y = -1, yend = -1.4,  # Y-positions for the tick marks (slightly below the axis line)
             color = "grey20", size = 0.7)  # Line color and thickness for the tick marks
}

bar_labels <- c(
  "<span style='font-size:10pt;'><b>B</b></span><br>Neoplasia<br>Prevalence", 
  "<span style='font-size:10pt;'><b>C</b></span><br>Malignancy<br>Prevalence", 
  "<span style='font-size:10pt;'><b>D</b></span><br>Number<br>of L1s", 
  "<span style='font-size:10pt;'><b>E</b></span><br>Number<br>of SINEs"
)

for (i in seq_along(bar_labels)) {
  x_center <- max(tree_plot$data$x) + offset_value + (i - 1) * (bar_width + bar_spacing) + (bar_width / 2)
  tree_plot_with_bars <- tree_plot_with_bars +
    annotate("richtext", x = x_center,  # Center of the bar plot
             y = length(tre$tip.label) + 1.8,  # Position above the top of the plot
             label = bar_labels[i], size = 3,  # Default text size for non-bold parts
             hjust = 0.55,  # Center it horizontally
             angle = 0,  # Keep text horizontal
             vjust = 0.05,
             color = "grey20",
             fill = NA,  # No background color
             label.color = NA)  # No border
}

# Print the final plot
print(tree_plot_with_bars)  # Output the final combined tree and bar plot
ggsave("tree_plot_with_bars.png", tree_plot_with_bars, width = 8, height = 10, dpi = 1000)
# Save with original file name
saveRDS(plot, "tree_plot_with_bars.rds")
```
