## Supplementary material for "Recent Non-LTR Retrotransposon Activity Predicts Cancer Prevalence in Mammals": README file

#### Pipeline and Data Repository for 
### **Recent Non-LTR Retrotransposon Activity Predicts Cancer Prevalence in Mammals**
This Readme file provides a step-by-step guide for reproducing all analyses and results presented in the study entitled **"Recent Non-LTR Retrotransposon Activity Predicts Cancer Prevalence in Mammals"**. Please note that this pipeline relies heavily on self-developed codes, and our coding strategy may not be the most advanced and optimized way to parse the data. One may prefer different coding approaches or strategies, but the methods described here worked correctly and efficiently for our purposes and reflect the exact pipeline used in this study. All items referenced here with numbering, including shell, Python, and R scripts as well as datasheets, are provided in the supplementary "Shell Scripts & Commands" folder.

#### **Repeat annotation**

We employed [RepeatModeler v2.0](https://www.repeatmasker.org/RepeatModeler/) to perform *de novo* identification of repetitive elements and to create species-specific repeat libraries. We then queried each species library against the respective reference genome to generate repeat annotations via [RepeatMasker v4.1](https://www.repeatmasker.org/). The output files were converted to `bed` format using `rmsk2bed` from [BEDOPS v2.4.41](https://github.com/bedops/bedops). The length of each repeat element was then calculated with the custom script `0_calculate_lengths.py`, and the values in column 10 of the bed files were replaced with these lengths. By default, column 10 records the number of query bases outside the repeat match (as reported by RepeatMasker), a metric not required for our analyses. We substituted these values with element lengths to simplify downstream processing.

#### **Abundance Model**

We used the `1_subset_BED.sh` script to subset the bed files into separate files containing only L1 or SINE elements, which can be saved in their own subdirectories. Next, we used two shell scripts,  `2_length_Divergence_LINE1s.sh` and `3_length_Divergence_ALL_SINES.sh`, to extract entries from the L1 and SINE bed files that matched both length and divergence criteria. Lengths, stored in column 10, and element-specific divergence values, stored in column 7, were used to apply the following thresholds:

- L1s within ±10% of the 6.1 kb consensus length and with ≤5% sequence divergence.

- SINEs between 100 and 400 bp in length and with ≤5% sequence divergence.

In the provided scripts, we extracted data at four different divergence cutoffs (i.e., 2%, 5%, 10%, and 20%), with each subset saved in a separate column. Elements that met these criteria were saved into separate files for L1s and SINEs. The resulting output files are formatted with species names in the first column, followed by counts of elements meeting the divergence thresholds in separate columns. For example:

<pre>---
Species            SINEs_Div_2   SINEs_Div_5   SINEs_Div_10  SINEs_Div_20

Acinonyx_jubatus        0            36            807          30531

Acomys_cahirinus        260         9269          109256        597939

Addax_nasomaculatus     0            4             38           24015
---</pre>

We used only elements with divergence rates of ≤5% for our analyses. The resulting data were compiled into the `Supplementary Data S1` Excel sheet for downstream analyses.

#### **Proximity Model**

We used the script `4_nLTRs_to_Genes_distance_script.sh` to compute the distance from each TE insertion (L1 or SINE) to the nearest protein-coding (PC) gene or cancer gene ortholog (CGO). The script requires the raw `bed` files generated from `rmsk2bed` from [BEDOPS v2.4.41](https://github.com/bedops/bedops) (i.e., files created from RepeatMasker `.out` files) and a gene annotation file in bed format. We provided a bed annotation file containing only PC genes for the proximity to PC genes analyses and a bed annotation file containing only CGOs for the proximity to CGOs analysis (see below). This script also depends on one additional file named `species_list.txt`, which must be located in the same directory. This plain text file enables submission as a job array as well as access to each species' subdirectories in respective analyses, with species names (i.e., Genus_species) listed in a single column without extensions, as follows:

<pre>---
Acinonyx_jubatus

Addax_nasomaculatus

Aepyceros_melampus

...

---</pre>

Each command in the script is explained with detailed comments. The flags used in the [BEDTools](https://bedtools.readthedocs.io/en/latest/) `closest` command are:

<code>`bedtools closest -d -a "TE_file.bed" -b "gene_annotation_file.bed" -t all -io -D a > gene_te_closest_distances.txt`</code>

`-a`: input file A (your TE annotation file in `bed` format)

`-b`: input file B (the gene annotation file in `bed` format)

`-d`: report the distance to the closest feature in B for each feature in A (adds an extra column with distance values)

`-t` all: when there are multiple equally close features, report all of them (instead of just one using `-t first` or `-t last`)

`-io`: ignore overlaps (if a TE overlaps a gene, it will not be counted as distance 0; the tool will instead look for the nearest non-overlapping gene)

`-D a`: report signed distance relative to A (negative if the gene is upstream of the TE, positive if downstream)


The outputs for each species are two text files, one for L1s and one for SINEs. Each includes three columns representing the superfamily, family, and the distance to the nearest gene (i.e., either PC gene or CGOs) as follows:

<pre>---
SINE    B4      6650

SINE    Alu     7129

SINE    Alu     1

SINE    B2      33918
---</pre>

Please note that this script relies on `awk` and `bedtools`, and differences in software versions may affect behavior and results. Thus, please use the same versions applied in this study to regenerate our results exactly. We then used the script `5_Combining_results.sh` to merge the L1 and SINE distance files for each species into a single output file named `Genus_species.txt` (e.g., `Elephas_maximus.txt`). For convenience, the individual L1 and SINE files from all species were first moved into a single directory, from which the merging script was run. Each merged file contains two columns: the first representing distances in bp for potentially active L1s and the second for potentially active SINEs. Each row corresponds to one element found within the species' genome, as follows:
<pre>---
Mus_musculus_L1   Mus_musculus_SINEs

    871856               730457

    807744               112685

    764709               6650

    94541                7129
---</pre>

In total, 55 files for each proximity scenario were transferred to the local machine, where IQR means and medians were calculated using the `14_calculate_means_medians` for all species and saved into an Excel sheet. This file has seven columns, including the species name and the IQR means and medians for L1s, SINEs, and combined L1 and SINE elements, which further merged into the `Supplementary Data S1`.

#### **Genic Insertion Model**

The same workflow was used to identify intersections of nLTRs within PG genes or CGOs, except that the command `bedtools intersect` was used in `6_intersects_counts_script.sh` with the following flags:

`-a`: input file A (TE annotation file in `bed` format). Each feature in this file will be tested for overlap with features in file B.

`-b`: input file B (the gene annotation file in `bed` format).

`-sorted`: tells `bedtools` that both input files are already sorted by chromosome and start coordinate. This speeds up the intersection and reduces memory usage.

`-c`: instead of outputting the actual overlaps, it appends a new column to each record in `-a`, showing the number of overlapping features in `-b`

The output of this script is a text file named `Genus_species.txt`, saved within the species subdirectory, which reports the number of active nLTR insertions within either PC genes or CGOs. The file has the following structure:
<pre>---
                L1      SINE

Mus_musculus    41      126
---</pre>

We then used the script `7_single_final_intersections_file.sh` to combine the intersection counts from all species into a single file, `all_species_gene_intersections.txt`. This file summarizes the number of nLTR insertions within PC genes or CGOs across all species and has the following structure:
<pre>---
species                 L1      SINE

Acinonyx_jubatus        32      19

Acomys_cahirinus        21      3894

---</pre>

These results were then directly pasted to the Supplementary Data S1 Excel sheet.

#### **Cancer Gene Load Model**

We first downloaded a list of 743 human cancer genes from the [Cancer Gene Census](https://www.sanger.ac.uk/data/cancer-gene-census) within the [COSMIC](https://cancer.sanger.ac.uk/cosmic) database. We retrieved their protein sequences from [Ensembl v112](https://www.ensembl.org/index.html) using [BioMart](https://www.ensembl.org/biomart/martview), resulting in protein sequences for 721 cancer genes. Next, we used this peptide file and the complete proteomes of 46 mammals with peptide BUSCO scores of over 60% in [OrthoFinder](https://github.com/davidemms/OrthoFinder). Two OrthoFinder output files, including `Orthogroups.GeneCount.tsv` and `Orthogroups.tsv` were used to 1) extract the CGO data and 2) create the CGO annotation files for the `bedtools` `closest` and `intersect` functions used for Proximity and Genic-insertion data extractions. Because the codes and commands used in this step are complex, please refer to the script named `8_commands_to_extract_CGO_data.sh` and follow the comments to regenerate the results. There is another shell script, `9_extract_CG_orthogroups_IDs.sh`, and a Python script, `10_counting_orthologs_for_each_gene.py`, which are required to complete this step. The final file generated from the `8_commands_to_extract_CGO_data.sh` is used as input for `10_counting_orthologs_for_each_gene.py` to create the final gene count file named `15_HCG_counts.xlsx` (see the shell scripts & commands supplementary folder). In this file, cancer gene names are represented as rows and species as columns, with each cell showing the count of the corresponding ortholog in the corresponding species.

We moved the `15_HCG_counts.xlsx` file to the local machine and merged it with the `16_CGO_Status.xlsx`, which includes the functional category status of each gene, into the new Excel sheet named `17_CGOs_counts_status.xlsx` (see the shell scripts & commands supplementary folder). The R script named `18_CGO_data_processing.R` was used to merge these two files, processing and counting the orthologs in each functional category, and ultimately generating the data that was incorporated directly into the `Supplementary Data S1`.

#### **Annotation Files**

We used the `13_extract_PC_genes.sh` script to generate annotation files containing only PC genes, based on the `gene_biotype=protein_coding` information in the attribute column of the main `gff` annotation file. To create the same annotation file for CGOs, we first used a shell script, `11_make_species_orthologs_list.sh`, to create `species_orthologs.txt`, which lists all cancer gene orthologs in rows and species in columns. This script uses one of the intermediate files generated from `8_commands_to_extract_CGO_data.sh` named `filtered_HCG_orthogroups.tsv`. Next, `species_orthologs.txt` was used as input together with the complete `gff` annotations of each species in the `script 12_making_HCG_gff.sh`. This script extracts annotation entries related to CGOs and generates a `gff` file containing only these genes. Because `gff` quality varies across species and some genes may not be annotated appropriately, the script first searches for gene entries, then searches for CDS if a gene entry is not found, and finally searches for available exons if neither is available.

We also used [Liftoff v1.6.3](https://github.com/agshumate/Liftoff) to generate gene annotation files for species lacking annotations by transferring information from closely related species with high-quality annotations. Proteomes for these species were then generated from the annotation files using the `convert` function in [GFFtk v23.11.2](https://github.com/nextgenusfs/gfftk). As these steps are straightforward and fully described in their respective documentations, we do not provide additional details here.

The statistical analyses were conducted in the R environment. All R scripts are thoroughly commented and self-explanatory. Please refer to the folder `2-R Scripts for Statistical Analyses` for the scripts corresponding to each analytical model used in this study.

Please contact us for additional questions or clarification of any ambiguities.
